# Unravelling cell type specific response to Parkinson’s Disease at single cell resolution

**DOI:** 10.1101/2023.01.04.522691

**Authors:** Araks Martirosyan, Francisco Pestana, Katja Hebestreit, Hayk Gasparyan, Razmik Aleksanyan, Suresh Poovathingal, Catherine Marneffe, Dietmar R. Thal, Andrew Kottick, Victor Hanson-Smith, Sebastian Guelfi, Emmanouil Metzakopian, T. Grant Belgard

**Affiliations:** VIB Center for Brain & Disease Research, KU Leuven, Leuven, Belgium; Verge Genomics, South San Francisco, CA, USA; Armenian Bioinformatics Institute, Yerevan, Armenia; Yerevan State University, Yerevan, Armenia; Laboratory for Neuropathology, Department of Imaging and Pathology and Leuven Brain Institute, KU-Leuven, and Department of Pathology, UZ Leuven, Leuven, Belgium; UK Dementia Research Institute, Department of Clinical Neurosciences, Cambridge Biomedical Campus, University of Cambridge, Cambridge, UK; The Bioinformatics CRO, Orlando, Florida, USA

## Abstract

Parkinson’s Disease (PD) is the second most common neurodegenerative disorder and is generally characterized by impaired motor functions. It currently affects 6.3 million people aged 60 years and more, worldwide. The pathological hallmarks of PD are Lewy bodies (abnormal aggregation of α-synuclein inside cells), which are observed primarily in the substantia nigra (SN) region of the midbrain. It is yet not known how different cell types in SN respond during PD and what are the molecular mechanisms underlying neurodegeneration. To address this question, we generated a large-scale single cell transcriptomics dataset from human post-mortem SN tissue of 29 donors including 15 sporadic cases and 14 controls. We obtained data for a total of ∼80K nuclei, representing major cell types of the brain (including neurons, astrocytes, microglia and oligodendrocytes). Pathway and differential gene expression analysis revealed multicellular character of PD pathology involving major cellular response from neuronal and glial cells.

## Introduction

Parkinson’s disease (PD) is a progressive neurodegenerative disorder predominantly affecting the elderly. Clinically PD is characterized by resting tremor, slowness of movement, rigidity and postural instability^1^. Loss of the neurotransmitter dopamine and dopaminergic neurons within the substantia nigra pars compacta (SNpc) was recognized as underlying the pathophysiology of the motor dysfunction. In the surviving neurons, accumulation of so-called Lewy bodies (protein aggregates composed primarily of alpha-synuclein, *SNCA*) is observed which is recognized as a primary hallmark for PD^1,2^.

Our understanding of molecular mechanisms underlying PD pathology remains poor. Here a major advance is expected from high-throughput single cell/nuclei RNA sequencing (RNA-seq) technologies that provide a unique opportunity to systematically study cell-type specific response to disease^3^. In recent years RNA-seq technologies are used more extensively in neuroscience, however they are just beginning to be applied to human models of PD, with a few applications *in vitro*^4,5^ and in post-mortem human brain tissues^6^. Obtaining a high-quality post-mortem dataset remains a challenge due to scarcity of the human brain tissue and inevitable RNA-degradation leading to poor RNA quality especially among neurons in PD condition, that have a high degradation rate.

To allow the development of *in vivo* models of cellular response to PD at improved resolution, in this study we provide a unique large-scale single nuclei RNA-seq dataset of 29 post-mortem human SNpc (15 sporadic PD and 14 control samples, see Table 1 for full pathological reports). The dataset includes the transcriptomes of ∼80K high quality nuclei at >40% sequencing saturation rate and on average ∼40K average read depth (see **Supplementary Table 1**). Our analyses were able to detect all the major cell types, including a large population of neurons derived from PD condition, see **Table 2**. Our analysis identified a strong neuronal association with PD pathology with significant degradation among dopaminergic and GABAergic neuronal populations as well as upregulation in major glia populations of genes associated with the response to unfolded proteins and oxidative stress.

## Results

### Single nuclei RNA-seq reveals cell type heterogeneity in human substantia nigra pars compacta

To characterize cell types in human substantia nigra pars compacta (SNpc), 15 PD and 14 Control human samples balanced by age and sex were selected from the Oregon Brain Bank to sequence the whole transcriptome from the nuclei located in SNpc using 10x Chromium technology, see Methods and **Table 1**. More than ∼80K high quality nuclei were harvested from these samples and clustering analysis were performed to identify residing cell types, see Methods and **Figure 1**.

**Figure 1:**
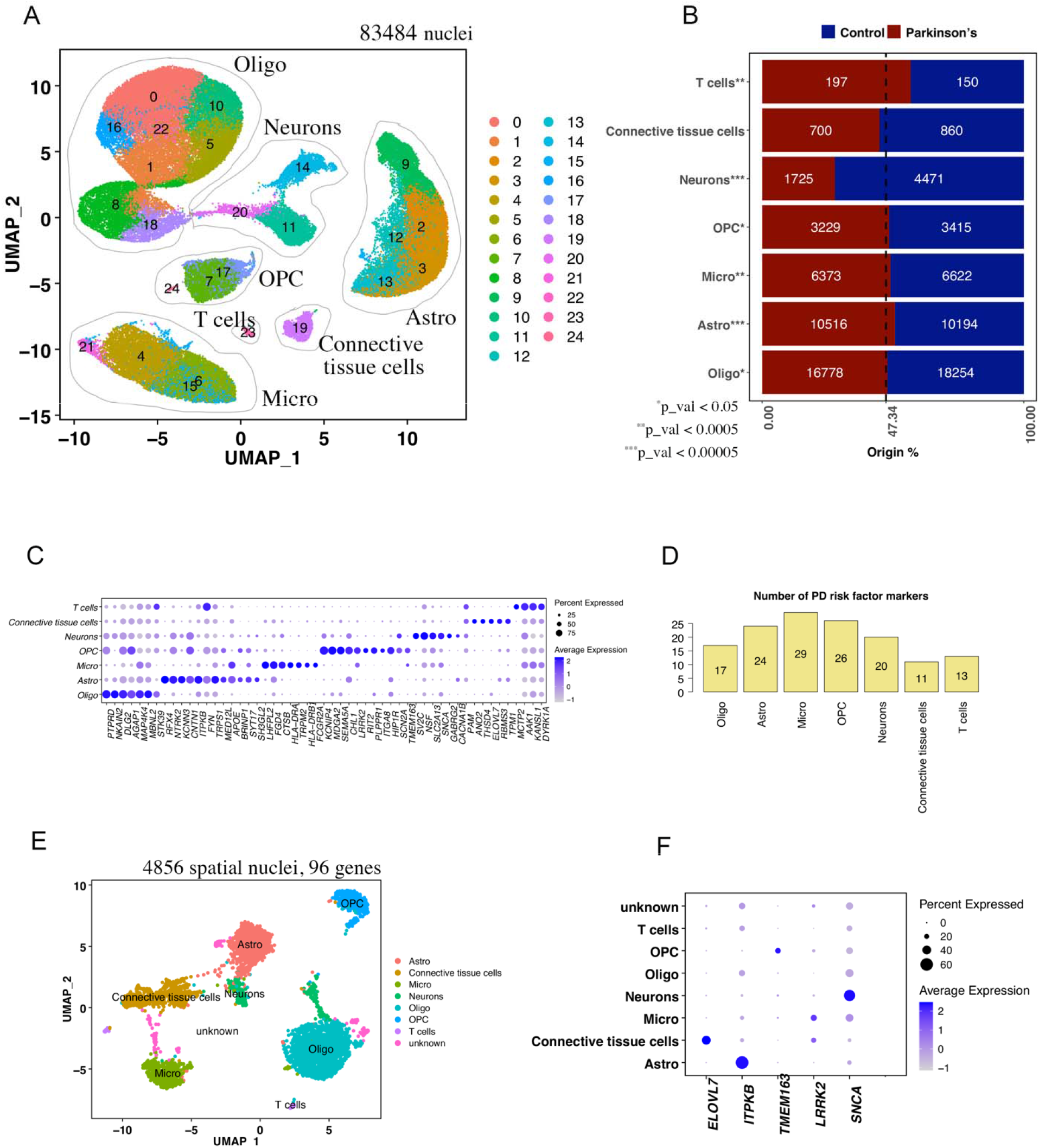
Cell-types in human substantia nigra and their susceptibility to PD. **(A)** Clustering 83,484 high quality nuclei obtained from 14 PD and 15 Control samples. Clusters representing Neurons, Oligodendrocytes (Oligo), Astrocytes (Astro), Microglia (Micro), Oligodendrocyte Progenitor Cells (OPC), T cells and Connective tissue cells were identified based on corresponding marker gene expression patterns, see Supplementary Figure 1. **(B)** A comparison of cell proportions deriving from PD and Control brains with the expected fraction of 47.34% cells representing PD condition among all the nuclei included in the clustering analysis. A binomial test was performed to see if there is any significant divergence of cell proportions from this value, *p-value < 0.05, ** p-value < 0.0005, *** p-value < 0.0005. **(C)** Specificity of PD risk associated genes collected from the EMBL GWAS catalog to cell types identified in panel (A). **(D)** The number of PD-associated genes specifically expressed in cell populations identified in panel (A). **(E)** Independent clustering of the spatial transcriptomics dataset generated using the tissue of 1 control patient to validate cell-type specific expression of PD risk factors reported in panel (C). **(F)** Confirmation of cell-type specific expression of PD-associated genes based on the spatial transcriptomics dataset: *ELOVL7* is mostly found in Connective tissue cells, *ITPKB* in Astrocytes, *TMEM163* in OPCs, *LRRK2* in Microglia and Connective tissue cells, *SNCA* has the highest expression in neurons, but is found also in other cells, such as Oligodendrocytes, Microglia and OPC.

Clustering pooled PD and Control data with Seurat^7^ identified the following populations: Neurons (*SYT1+SNAP25+*), Oligodendrocytes (Oligo, *MOBP+MBP+*), Microglia (Micro, *CD74+ITGAM+*), Connective tissue cells (*FLT1+DCN+*), Oligodendrocyte Progenitor Cells (OPC, *VCAN+PDGFRA+*), Astrocytes (Astro, AQP4+SLC1A3+), T cells (*THEMIS+CD2+*), see **Figure 1A** and **Supplementary Figure 1**.

Evaluation of cell proportions showed that neurons are heavily lost in PD samples as expected, while relative proportions of glial and T cells increase, as shown in **Figure 1B** and **Table 2**. This observation aligns with previous reports showing that the number of T cells increases in the aged brain^8^, therefore this effect might be a hallmark of brain aging, which is a major risk factor for developing PD^9^.

### Cell-type specific expression of genes are associated with PD GWAS loci

Next, we asked if PD genetic risk factors are linked to specific cell types or not. We overlayed the list of known genetic factors obtained from GWAS studies reported in the NHGRI-EBI catalog^10^ with population specific markers, see Methods and **Table 3**. We observed a pattern of cell-type specificity in key genes like *SNCA*^11^ (mostly expressed in Neurons), *LRRK2*^12^ (mostly expressed in Microglia and OPC), *ITPKB*^13^ (mostly expressed in Astrocytes), *ELOVL7*^14^ (specific to Connective tissue cells), *TMEM163*^15^ (specific to OPC), see **Figure 1 C, D**. To confirm these observations, we performed multiplex spatial transcriptomics staining 96 markers in tissue using molecular cartography of Resolve Biosciences, see Methods, **Table 4** and **Figure 1E**. Clustering analysis based on those 96 markers allowed us to detect all the cell-types observed in single nuclei dataset, see **Figure 1E**. Spatial co-localization of *SNCA* with neuronal markers *SYT1* and *SNAP25, LRRK2* with microglial marker *ITGAM, ITPKB* with astrocytic marker *AQP4, ELOVL7* with connective tissue cell marker *FLT1*, and *TMEM163* with OPC marker *VCAN* confirmed the expected specificity of these genes to corresponding cell types, see **Figure 1F**.

These results are consistent with a multicellular nature of PD origin and a complex involvement of multiple cell types in its pathology. Next, we explored PD-related molecular changes among neuronal and major glial populations – astrocytes, microglia, and oligodendrocytes.

### Neuronal subtypes in human substantia nigra and their response to PD

Re-clustering neurons separately revealed 6 neuronal subtypes, which we call neurons0 - neurons5, with well-defined markers, see **Figure 2A, B** and **Table 5**. All subpopulations have representation from multiple PD and Control samples, see **Figure 2C**.

**Figure 2:**
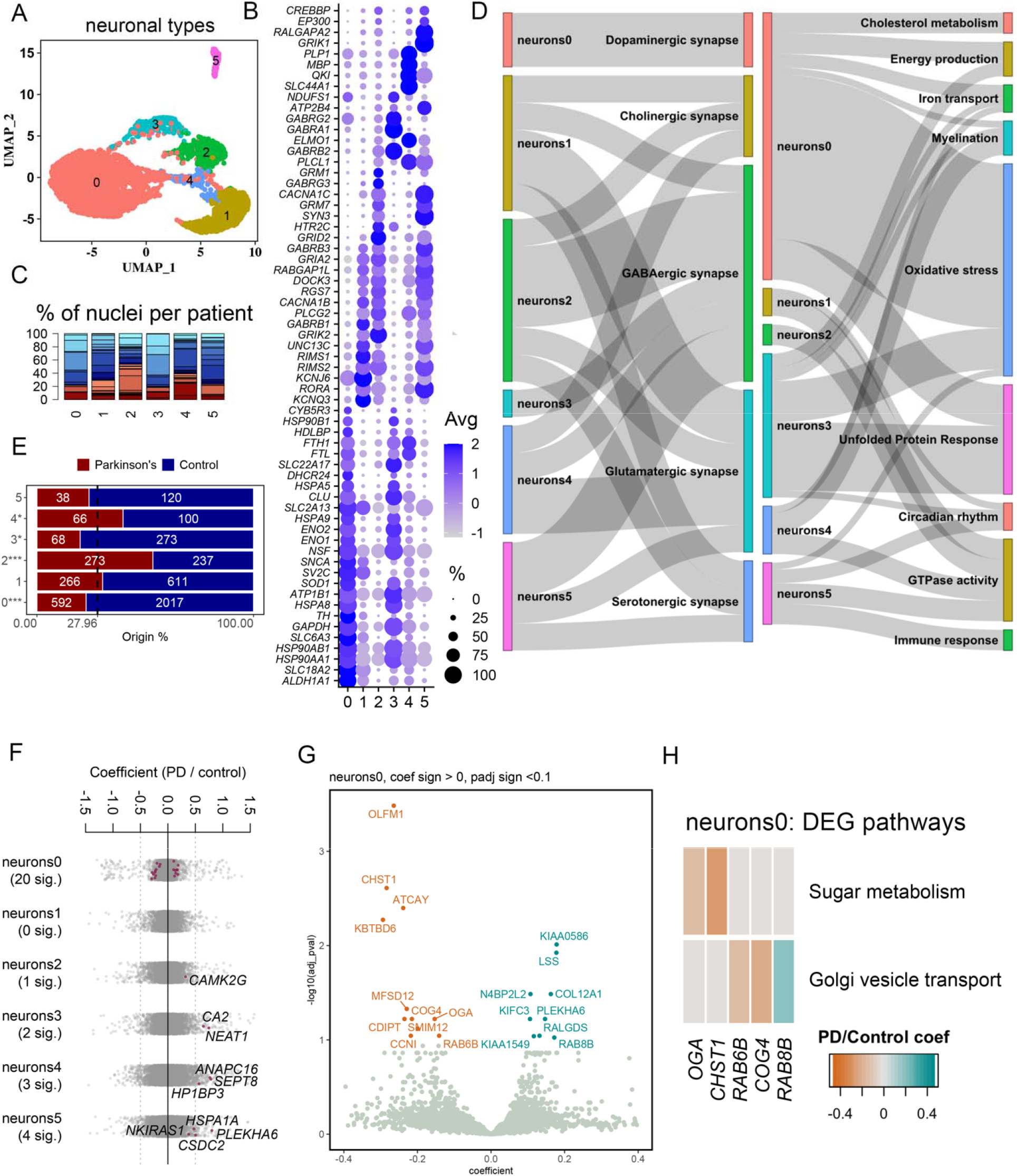
Neuronal composition of human substantia nigra pars-compacta and the response to PD: **(A)** Re-clustering neurons found in Figure 1A. **(B)** Marker genes of neuronal populations reported in Figure 2A. **(C)** Proportion of nuclei coming from Control (blue color-scheme) and PD (red color-scheme) patients per neuronal type reported in Figure 2A. **(D)** A summary of pathway enrichment analysis on the marker genes of neuronal population neurons0-5. **(E)** A comparison of cell proportions deriving from PD and Control brains with the expected fraction of 27.96% cells representing PD condition among all the nuclei included in the neuronal clusters. A binomial test was performed to see if there is any significant divergence of cell proportions from this value, *p-value < 0.05, ** p-value < 0.0005, *** p-value < 0.0005. **(F)** A strip chart reporting the number of significantly up/down regulated genes in PD per neuronal populations found by fitting a linear mixed model, red dots correspond to genes that are up (coefficient > 0) or down (coefficient<0) regulated with the adjusted p value < 0.1 (ANOVA test with Benjamini-Hochberg correction) **(G)** A volcano plot reporting gene up (cyan) or down (orange) regulated in PD neuronal population neurons0. **(F)** GO and KEGG pathways overrepresented in PD up/down regulated genes in neuronal population neurons0.

Pathway overrepresentation analysis on the full list of marker genes (significantly enriched genes with log_2_ Fold change > 0.25 and adjusted p-value < 0.05), reported in **Table 6** and **Supplementary Figure 2**, showed that one of the populations - neurons0, is characterized by markers of dopaminergic neurons (positive for *TH, SLC6A3, SNCA*), see **Figure 2B, D** and **Table 5**. The markers of this population are involved in key processes known to be linked to the PD pathology, such as energy production^16^ (*ATP1B1, ENO1, ENO2, etc*.), cholesterol metabolism^17^ (*DHCR24, CYB5R3, HDLBP, etc*.), iron transport^18^ (*FTL, FTH1, SLC22A17, etc*.), oxidative stress^19^ (CHCHD10, CLU, SOD1, etc.). As expected, this population expressed transcripts linked to unfolded protein response (involving protein-folding assistant chaperons *HSPA8, HSP90AA1, etc*.), which is expected upon aggregation of alpha-synuclein in dopaminergic neurons after the onset of PD, see **Figure 2D** and **Table5**.

Observation of markers shows that the second population positive for members of *HSPA, HSP90* heat shock protein proteins is neurons3, which has a clear GABAergic signature (positive for *GABRG2, GABRA1, GABRB2*), see **Figure 2B, D**. With dopaminergic neurons0, GABAergic neurons3 share many markers associated with unfolded protein response, oxidative stress, energy production and iron transport, see **Figure 2D** and **Table 5**. Therefore, it is not surprising that dopaminergic neurons0 and GABAergic neurons3 are both significantly depleted in numbers after the onset of PD, demonstrated in the cell proportion comparisons in PD vs control brains in **Figure 2E**.

The remaining populations of neurons show involvement in GABAergic (*GABRB1, GABRB3, GABRG3*), glutamatergic (*GRIA2, GRM1, GRM7, GRID2, GRIK2*), cholinergic, and serotonergic signalling, with neurons4 having the strongest expression of choline transporter-like protein 1 *SLC44A1*, and neurons2 the highest expression of serotonin 5-HT-2C receptor *HTR2C*, see **Figure 2B, D**. In contrast to degenerating dopaminergic neurons0 and GABAergic neurons3, the GO and KEGG over-enrichment analysis on markers of remaining neuronal populations (i) do not show enrichment in unfolded protein response, (ii) show enrichment in GTPase activity^20^ (*RIMS1/2, RALGAP(1L/A2), RGS7*, see **Figure 2B**), (iii) show high expression of transcripts associated with theimmune response (such as *PLCG2, CREBBP, EP300* with significant GO/KEGG term enrichment only in neuronal population neurons5).

Next, we examined differentially expressed genes (DEGs) between PD and control condition within each neuronal subtype by fitting a linear mixed model, see Methods and **Table 7**. A summary stripchart of these analyses is shown in **Figure 2F**. We can see that no DEGs are detected in neuronal population neurons1, neurons3,4,5 have a mild response with a few upregulated genes. The highest response is triggered in neurons0 dopaminergic neuronal population, see **Figure 2G, F** and **Table 8**. Pathway overrepresentation analysis on up/down regulated genes within this population suggest perturbations in genes involved in sugar/glucose metabolism (due to the downregulation of O-GlcNAcase *OGA* and Carbohydrate sulfotransferase 1-*CHST1*), which is reported to be an early event in the sporadic PD^21^. Furthermore, dysregulation of Ras-related *RAB6B* and *RAB8B* together with Component of Oligomeric Golgi Complex 4 - *COG4* genes is consistent with dysfunction of Golgi to endoplasmic reticulum vesicle trafficking, in line with recent reports of the disruption of vesicle trafficking caused by the accumulation of alpa-synuclein^22^ and the potential of RAB GTPase to rescue neurons from death^23^.

### Astrocytic response to PD

Re-clustering astrocytes separately revealed 6 distinguishable states astrocytes0 - astrocytes5 that were represented across all samples, see **Figure 3 A-C**. Pathway overrepresentation analysis showed that all the populations except for astrocytes3 show high expression of transcripts linked to synaptic functions (*LRRC4C*^24^, *VAV3*^25^, *CASK*^26^), ion (Solute Carrier family members *SLC6A11, SLC39A12, SLC1A2*) and calcium transport (Calmodulin *CALM1*, Calcium Voltage-Gated Channel Auxiliary Subunit *CACNB2*), see **Figure 3B, D, Tables 5-6** and **Supplementary Figure 3**.

**Figure 3:**
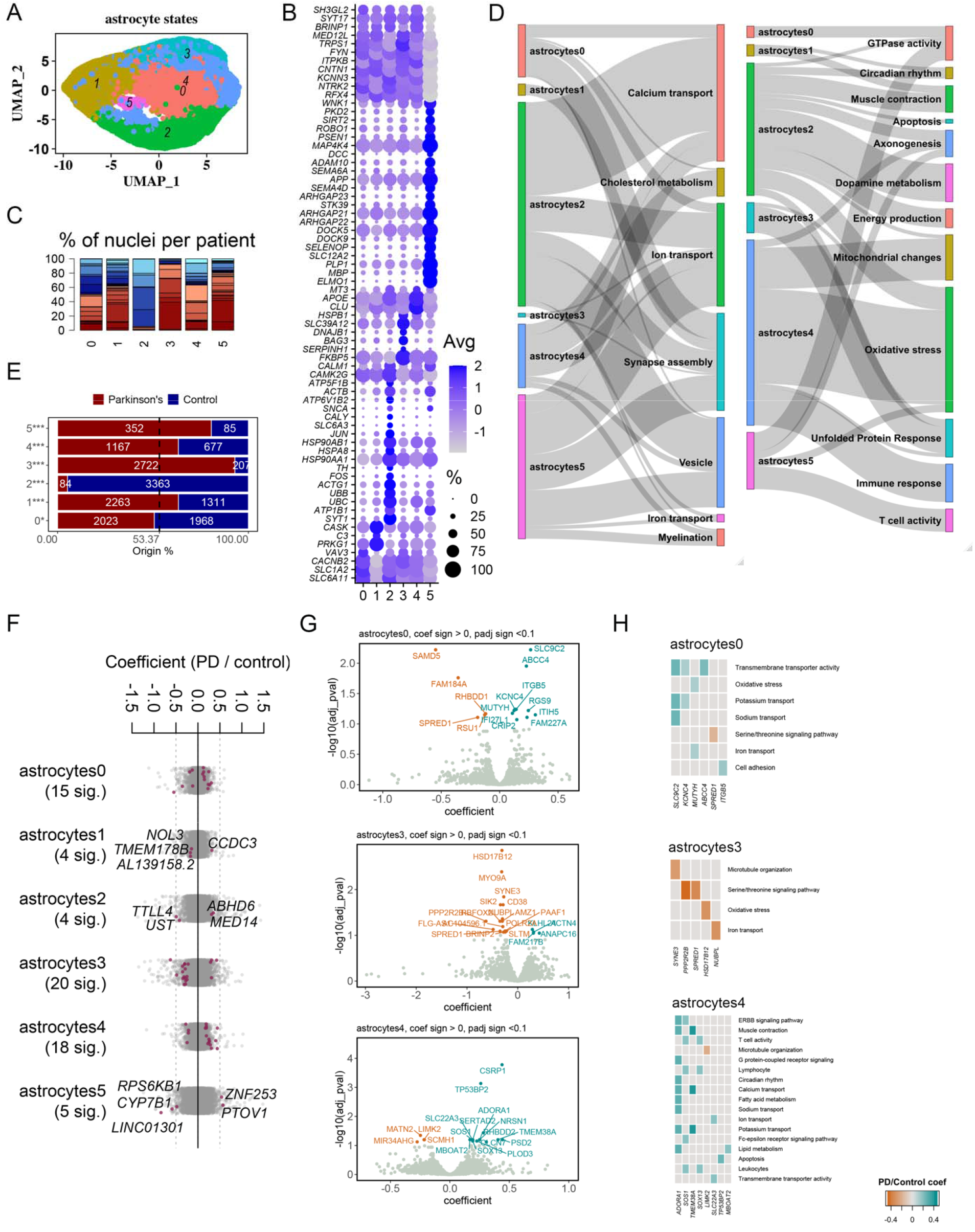
Astrocytic states found in human substantia nigra pars-compacta and their response to PD: **(A)** Re-clustering astrocytes found in Figure1A. **(B)** Marker genes of astrocytic states shown in Figure3A and related pathways. **(C)** Proportion of nuclei coming from Control (blue color-scheme) and PD (red color-scheme) patients per astrocytic state reported in Figure3A. **(D)** A summary of pathway enrichment analysis on the marker genes of astrocytic population astrocytes0-5. **(E)** A comparison of cell proportions deriving from PD and Control brains with the expected fraction of 53.37% cells representing PD condition among all the nuclei included in the astrocytic populations. A binomial test was performed to see if there is any significant divergence of cell proportions from this value, *p-value < 0.05, ** p-value < 0.0005, *** p-value < 0.0005. **(F)** A strip chart reporting the number of significantly up/down regulated genes in PD per astrocytic states found by fitting a linear mixed model, red dots correspond to genes that are up (coefficient > 0) or down (coefficient<0) regulated with the adjusted p value < 0.1 (ANOVA test with Benjamini-Hochberg correction) **(G)** Volcano plots reporting gene up (cyan) or down (orange) regulated in PD in the most responsive astrocytic states astrocytes0,3,4. **(H)** GO and KEGG pathways enriched in PD up/down regulated genes within astrocytes0,3,4.

In contrast, pathways overrepresented among the markers of subpopulation astrocytes3 are predominantly associated with unfolded protein response (driven by transcripts *BAG3, SERPINH1, DNAJB1, HSPB1*) **Figure 3D** and **Table 6**. Interestingly, recent reports suggest that astrocyte reactive state generated in response to unfolded proteins is fatal for neurons^27^. Comparison of cell proportions within the astrocytes shows that astrocytes3 were mostly derived from patients diagnosed with PD, see **Figure 3E**, suggesting that this reactive astrocyte state may be PD-specific.

Among all the astrocyte states astrocytes2 is the only one largely enriched in control brains. It is also the only state of astrocytes that has markers linked to dopamine metabolism (positive in *TH, SLC6A3, CALY, SNCA*) and involved in energy production (*ATP5F1B, ATP6V1B2, ATP1B1*), see **Figure 3B, D, Tables 5-6** and **Supplementary Figure 3**. Previous reports propose that TH expressing astrocytes can induce a functional recovery in an animal model of PD^28,29^. We speculate that astrocytes2 therefore may be a natural way to prevent the development of Parkinsonism in the human brain. At the same time astrocytes2 is positive for Ubiquitin membrane protein transcripts *UBB* and *UBC* that control endocytic vesicle trafficking, *SNCA*, protein chaperons assisting in protein folding *HSP90AA1, HSP90AB1, HSPA8*, which suggest that this population may suffer from oxidative stress upon aggregation of unfolded α-synuclein. As a result, astrocytes2 shows high expression of *JUN, FOS* transcripts that suggest activation of apoptosis^30^, **Figure 3B, D, Tables 5-6** and **Supplementary Figure 3**, which may explain why they are absent in the PD brain.

The well-known reactive astrocytic marker *C3* is more enriched in astrocytes1, that is likely to be another state of astrocyte reactivity^31^, see **Figure 3B, D, Tables 5-6**.

Astrocytes4 have the highest expression of *APOE, MT3, CLU* and is enriched in PD brains, which suggests that mitochondrial dysfunction invokes oxidative stress and immune response in this population^32^, see **Figure 3B, D** and **Table 5-6**.

Astrocytes0 seems to be more resistant to PD: it is lost in PD in less significant numbers and has a high expression of markers linked to synapse growth and function *SLC1A2*^33^, *LRRC4C*^24^ and *VAV3*^25^. Unlike astrocytes2, this population does not have transcripts related to the regulation of dopamine metabolism which may explain the resistance of these cells to PD pathology, see **Figure 3B, D** and **Table 5-6**.

At the level of differential gene expression in PD vs Control condition all the astrocytic states demonstrate a response, see **Figure 3F**, with the highest number of significant DEG genes found in astrocytes0,3,4. These genes are linked mostly to mineral transport in astrocytes0 (*SLC9C2, KCNC4, MUTYH, ABCC4, ITGB5*), serine/threonine signaling pathway (*PPP2R2B, SPRED1*) and oxidative stress (*HSD17B12*) in astrocytes3; and T-cell activity/immune in astrocytes4 (*SOS1*) as shown in **Figure 3G, H** and **Tables 7, 8**.

### Microglial response to PD

Re-clustering analysis of microglia revealed 6 states microglia0 - microglia5, see **Figure4 A, B, C**. One of them largely representing control condition – microglia1, which is the only population with markers involved in dopamine metabolism (positive for *TH, SLC6A3, SNCG*), see **Figure 4, Tables 5, 6** and **Supplementary Figure 4**. It is unknown to us if microglia express TH under stress or if this is a technical artifact from synaptic pruning or phagocytosis. This population has high levels of transcripts linked to HSP90 (*HSP90AA1, HSP90AB1*) and HSPA (*HSPA8*) protein families – protein chaperones assisting in protein folding – consistent with oxidative stress associated with unfolded protein aggregation, a scenario that we saw for astrocytes2. Like astrocytes2, microglia1 also positive for markers associated with apoptosis (positive for *FOS, ACTG1*).

**Figure 4:**
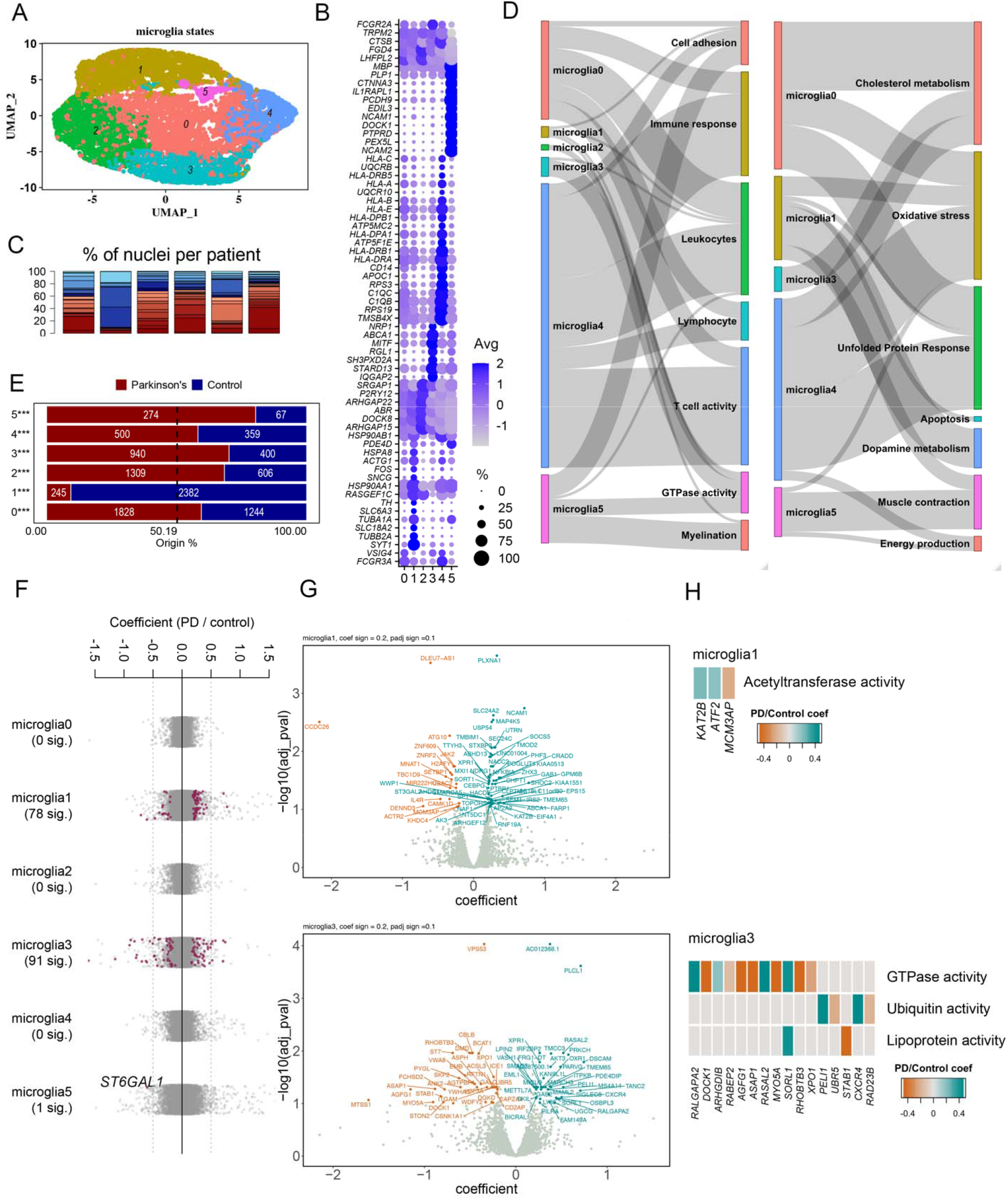
Microglia states found in human substantia nigra pars-compacta and their response to PD: **(A)** Re-clustering microglia found in Figure1A. **(B)** Marker genes of microglial states shown in Figure4A. **(C)** Proportion of nuclei coming from Control (blue color-scheme) and PD (red color-scheme) patients per states reported in Figure 4A. **(D)** A summary of pathway enrichment analysis on the marker genes of microglia states microglia0-5. **(E)** A comparison of cell proportions deriving from PD and Control brains with the expected fraction of 50.19% cells representing PD condition among all the nuclei included in the microglial populations. A binomial test was performed to see if there is any significant divergence of cell proportions from this value, *p-value < 0.05, ** p-value < 0.0005, *** p-value < 0.0005. **(F)** A strip chart reporting the number of significantly up/down regulated genes in PD per microglial states found by fitting a linear mixed model, red dots correspond to genes that are up (coefficient > 0) or down (coefficient<0) regulated with the adjusted p value < 0.1 (ANOVA test with Benjamini-Hochberg correction) **(G)** Volcano plots reporting gene up (cyan) or down (orange) regulated in PD in microglial states microglia1,3. **(F)** GO and KEGG pathways overrepresented in PD up/down regulated genes within microglia1,3.

Among the markers of microglia2 *P2RY12* is standing out as a well investigated P2Y receptor involved in microglial motility and migration towards (damaged) cells releasing ATP, an early event in neuroinflammation process^34^, see **Figure 4B**. This population is also positive for *ARHGAP* family transcripts (*ARHGAP22* and *ARHGAP15*) that are known to be linked to the microglia activation state upon aging^35^. On the other hand, microglia2 lacks markers linked to protein misfolding, **Figure 4D, Tables 5-6** and **Supplementary Figure 4**. Together this data suggests that microglia2 might be an early stage of activated microglia that is typical also for aging, which may explain the absence of PD-related up/down regulated genes in this population, see **Figure 4F**.

Microglia4 seems to represent an activated state of the microglia (expressing markers of complement cascade *C1QC, C1QB, C1QA*^36^), see **Figure 4B**. This population is enriched in human leukocyte antigen (HLA) genes^37,38^ consistent with a concurrent unfolded protein response (mediated by *HSP90* and *HSPA* chaperones) and a large Immune response, **Figure 4B, D, Tables 5, 6** and **Supplementary Figure 4**.

Interestingly, microglia0 seems to express most of the markers of microglia4 at a lower level, as shown in **Figure 4B** and **Supplementary Figure 4**, suggesting that it may be a state forming a continuum with microglia4.

Like microglia2, microglia0 and microglia4 also do not show PD-related up/down regulated genes, however they show a response to unfolded proteins. Therefore, we hypothesize that microglia0 and microglia4 may represent later activation stages of microglia2 driven by age-related misfolded, unfolded, or aggregated proteins^39^.

Microglia5 expresses transcripts linked to cell adhesion (*NCAM1/2, PCDH9, DOCK1/2*) and has a large enrichment of PD-samples, suggesting that it is a distinct activated state of microglia. A key regulator of immunological processes *ST6GAL1*^40^ appears to be downregulated in microglia5 after onset of PD, see **Figure 4F**.

At the level of differentially expressed genes between PD and control conditions microglia1 and microglia3 show a strong response, see **Figure 4F, G** and **Table 7, 8**. Microglia1 shows acetyltransferase activity (regulated by *KAT2B, ATF2, MCM3AP*), which is known to cause inflammatory response in microglia^41^.

### Oligodendrocytic response to PD

Re-clustering of oligodendrocytes revealed 6 states oligos0 – oligos5, see **Figure 5A, B, C**. Like astrocytes2 and microglia1, one of oligodendrocytic subpopulations - oligos2 – is positive for *TH, SLC6A3, SNCG* (transcripts associated with dopamine metabolism) and is largely represented in controls, see **Figure 5B-E**. This population is enriched in transcripts linked to axon development and synapse organization (*UCHL1, NEFL, MAP1B, NRXN3, THY1, CNTN1, GAP43, ANK3, NEFH, ROBO2*), as well as markers involved in ion transport (*CNTN1, ANK3*) and synaptic vesicle cycle (*SLC18A2* - Vesicular monoamine transporter 2; and CALY - Calcyon Neuron Specific Vesicular Protein), see **Figure 5B, D, Supplementary Figure 5, Tables 5-6**.

**Figure 5:**
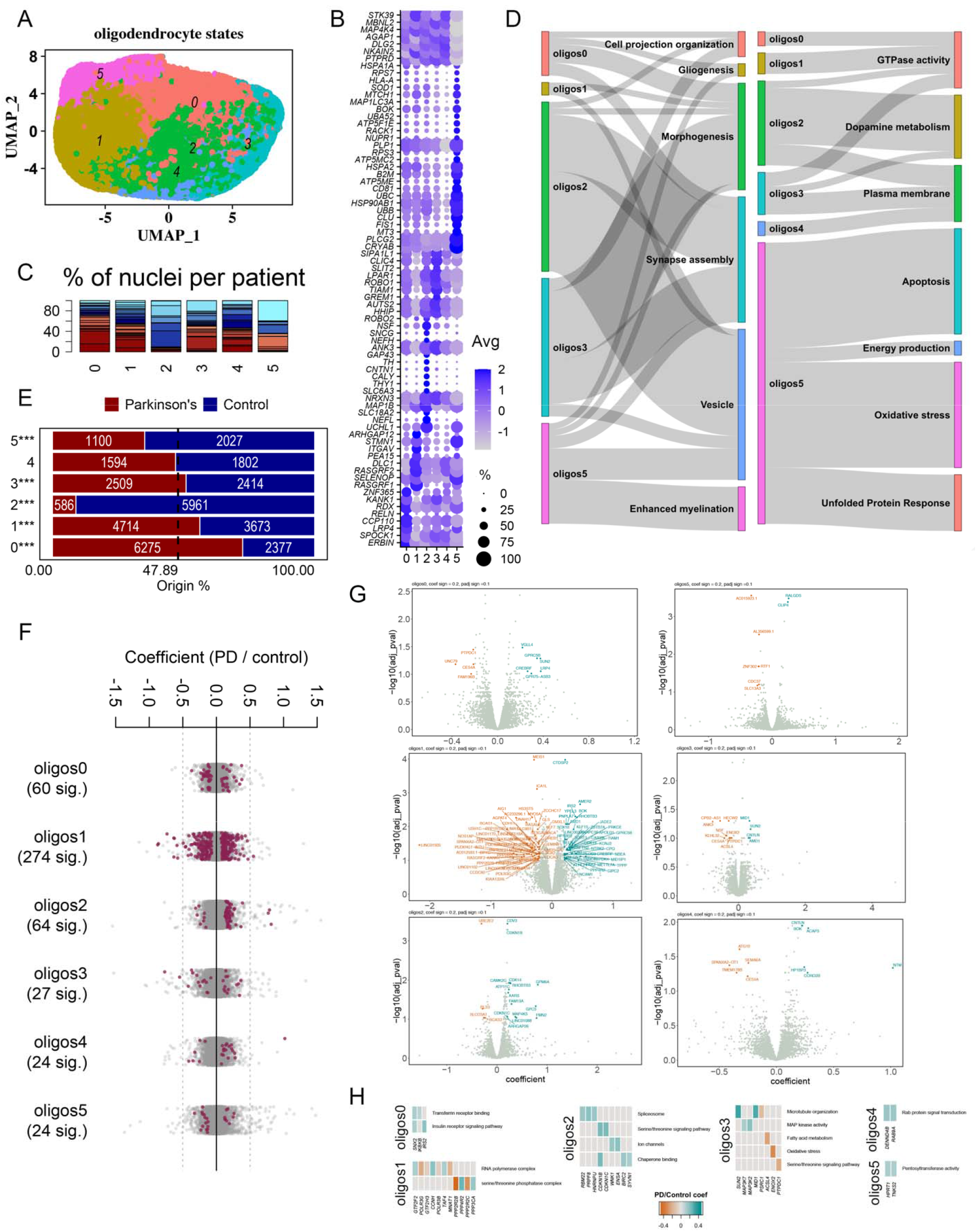
Oligodendrocyte states found in human substantia nigra pars-compacta and their response to PD: **(A)** Re-clustering oligodendrocytes found in Figure1A. **(B)** Marker genes of oligodendrocyte states shown in Figure5A. **(C)** proportion of nuclei coming from Control (blue color-scheme) and PD (red color-scheme) patients per states reported in Figure5A. **(D)** A summary of pathway enrichment analysis on the marker genes of Oligodendrocyte states Oligos0-5. **(E)** A comparison of cell proportions deriving from PD and Control brains with the expected fraction of 47.89% cells representing PD condition among all the nuclei included in the oligodendrocyte populations. A binomial test was performed to see if there is any significant divergence of cell proportions from this value, *p-value < 0.05, ** p-value < 0.0005, *** p-value < 0.0005. **(F)** A strip chart reporting the number of significantly up/down regulated genes in PD per oligodendrocyte states found by fitting a linear mixed model, red dots correspond to genes that are up (coefficient > 0) or down (coefficient<0) regulated with the adjusted p value < 0.1 (ANOVA test with Benjamini-Hochberg correction) **(G)** Volcano plots reporting gene up (cyan) or down (orange) regulated in PD in oligodendrocyte states. **(F)** GO and KEGG pathways enriched in PD up/down regulated genes within oligodendrocyte states reported in Figure 5A.

The second population of oligodendrocytes more enriched in control condition is oligos5. Pathway overrepresentation analysis on the markers of this population show enrichment of terms related to oxidative stress (*CRYAB, MT3, SELENOP, MAP1LC3A, HSPA1A, SOD, MT3, RACK1*), response to protein aggregates (*CLU, HSPA2, HSPA1A, HSP90AB1, BOK*), ATP biosynthesis (*ATP5ME, ATP5F1E, ATP5MC2*), mitochondrial function (*MT3, UBB, UBC, UBA52, MTCH1*) and apoptosis (*FIS1, UBB, RACK1, RPS3, NUPR1, BOK, RPS7, PEA15, SOD1*), see **Figure 5B, D, Supplementary Figure 5, Tables 5-6**. Interestingly, oligos5 shows a higher expression of *PLP1, MBP* linked to myelination, and markers linked to immune response (*PLCG2, CD81, B2M, HLA-A*).

Among oligodendrocytes oligos0 is the subpopulation largely representing the PD condition. Pathway overrepresentation analysis on the markers of this population shows enrichment of terms linked to cell projection organization (*SPOCK1, CCP110, KANK1*), dendrite (*LRP4, ZNF365, RELN*) and epithelium (*CLIC4, SLIT2, GREM1, HHIP*) morphogenesis, glial cell differentiation (*NFIB, ZNF365, NDRG1, RELN*), and GTPase activity (ERBIN, RELN, RDX, KANK1), **Figure 5B, D, Supplementary Figure 5, Tables 5-6**.

GTPase activity is a term enriched also in oligos1 markers (*RASGRF1, RASGRF2, DLC1, ITGAV, STMN1, ARHGAP12*) and oligos3 (AUTS2, LPAR1, SLIT2, ROBO1, SIPA1L1, KANK1, TIAM1), that have a higher representation in PD brains, see **Figure 5, Supplementary Figure 5, Tables 5-6**.

All the states of oligodendrocytes show a very strong PD-related response with a large number of up/down regulated genes, see **Figure5 F-H, Tables 7-8**. Among them oligos2 demonstrates up/downregulation of genes linked to spliceosome (*RBM22, PRPF8, HNRNPU*) and ion channels (*WNK1, ENSA*), oligos5 shows upregulation of Hypoxanthine Guanine Phosphoribosyltransferase HPRT1 and TRF1-interacting ankyrin-related ADP-ribose polymerase 2 *TNKS2* regulating pentosyltransferase activity.

GO/KEGG terms overrepresented within oligos1 up/downregulated transcripts are RNA polymerase complex (*GTF2F2, POLR3G, GTF2H3, CCNH, POLR3B, TAF4, MNAT1*) and serine/threonine phosphatase complex (*PPP2R2B, PPP4R2, PPP2R2C, PPP3CA*). Within oligos3 enriched GO and KEGG terms are liked to MAP kinase activity (*MAP3K7, MAP3K2*). Finally, in oligos4 DENND4B and *RAB9A* genes are upregulated that show an overrepresentation in Rab protein signal transduction, see **Figure5 F-H, Tables 7-8**.

## Discussion

Here we provide a large resource for PD research: a snRNA-seq dataset of human post-mortem SN samples collected from 14 PD and 15 control brains. Our dataset reports the transcriptomics of ∼80K high quality nuclei that allows to identify PD induced cell-type specific changes in all major cell types of human substantia nigra – neurons, astrocytes, microglia and oligodendrocytes.

While we detected various classes of neurons, the major effect appears to be on the dopaminergic system. Dopaminergic neurons, as well as populations of astrocytes, microglia and oligodendrocytes regulating dopamine metabolism, were lost in PD at statistically significant levels, consistent with the known role of glia cells in protecting neurons^42^, see below.

Dopaminergic neurons show high expression of protein chaperons *HSPA8, HSP90AA1*, that are known to regulate protein folding and stability^43,44^. GO/KEGG term over-presentation analysis of up/down regulated genes in PD condition suggests sugar metabolism and Golgi to endoplasmic reticulum (ER) vesicle trafficking as main processes impacted within dopaminergic neurons. This is consistent with reports that glycosylation in Golgi and ER are critical quality control steps to monitor the status of protein folding^45,46^. On the other hand, N-acetyl-glucosamine modification makes alpha-synuclein less toxic for neurons^47^. These observations with our data are consistent with glycosylation as a cellular mechanism reducing aggregation of misfolded alpha-synuclein within dopaminergic neurons. We speculate that this may be related to abnormal sugar uptake observed within PD patients^48,49^.

A high degree of heterogeneity among disease-responsive states of all glial cells has been observed. Interestingly, the microglia activation state positive for complement cascade molecules *C1QC, C1QB, C1QA*^50^ (microglia4) does not differ in abundance between PD and control conditions, suggesting that this mechanism of the innate immune system is equally active in aged brain to clear pathogens and dead or dying cells, contributing to the creation of a vulnerable pre-parkinsonism state^51^. In contrast, among astrocytes a PD specific reactive astrocyte state (astrocytes3) was detected, which is enriched for genes associated with unfolded protein response (UPR). This observation aligns with recent reports of a UPR induced response in astrocytes causing neurodegeneration^27^ suggesting that this population of astrocytes may be directly toxic for neurons.

Among astrocytes, microglia and oligodendrocytes very distinct states (astrocytes2, microglia1 and oligos2) were observed that were typical for control brains. These glial states are positive for dopamine transporter *SLC1A3* (also known as DAT) and tyrosine to dopamine converter enzyme - tyrosine hydroxylase *TH*. To our knowledge there are no reports of endogenous TH expression in glia cells, and this observation may be a technical artifact, however previous studies suggest that inducing TH expression in astrocytes may prevent parkinsonism in rat^28,29^.

All *TH*+ glia populations are almost absent in PD brains, where dopaminergic neurons are heavily lost. Interestingly, astrocytes2 and microglia1 express transcripts involved in apoptosis, oxidative stress and unfolded protein aggregates, while oligos2 does not. This observation suggests that the reduction of astrocytes2 and microglia1 numbers in PD may be a result of direct cell death, while oligos2 may convert to another state.

Our analyses suggest a multi-cellular response to PD. Our data provide a resource for future hypothesis-driven experiments aimed at modifying PD prognosis.

## Methods

### Selection of donors

We selected a total of 29 individuals from the **Oregon Brain Bank**, a human tissue repository for brain research, to perform single nuclei sequencing. The Ethics Committee of UZ Leuven/KU Leuven has approved the current study under the S-number **S64182**. Fresh frozen tissue together with the clinical reports providing information about donor’s clinical diagnosis, age, sex, as well as details about Post-Mortem Interval (PMI) before the tissue was collected, the quality of RNA (RNA integrity number, RIN measure), the presence of Lewy bodies in midbrain, limbic (amygdala) and neocortical (frontal cortex) regions, AD-related changes in cortex (presence of neuritic plaques and neurofibrillary tangles), please see **Table 1**. The presence of Lewy bodies, plaques and tangles is estimated as follows:

1. Lewy bodies are estimated to be present (1) or absent (0) based on the immunostains for alpha-synuclein on the midbrain, amygdala, and frontal cortex. The tissue was labeled positive (1) if at least a few Lewy bodies were detected in the respective locations.
2. Neuritic plaques (the density of tau-positive neurites in the setting of an at least equivalent amount of beta-amyloid immunostaining) are estimated to be absent (CERAD score 0), sparse (CERAD score 1), moderate (CERAD score 2), or abundant (CERAD score 3) based on the evaluation of frontal cortex with immunostains to both tau and beta-amyloid.
3. Neurofibrillary tangles are scored using the measure of Braak staging (present versus absent in various regions of brain). This is calculated as a summary of a series of tau immunostains that are performed on (at least) frontal cortex, hippocampus/entorhinal cortex/inferior temporal cortex, and occipital cortex. Braak stage 1-3 reflects involvement of various parts of the hippocampus and entorhinal cortex with sparing of the rest of the temporal cortex and frontal cortex. Braak stage 4 implies tangles in multiple regions of the hippocampus as well as in the temporal cortex. Braak stage 5 implies tangles in all the previously mentioned areas plus frontal cortex. Braak stage 6 implies tangles in all the previously mentioned areas plus occipital cortex. In classic descriptions of Alzheimer’s disease, tangles are said to spread temporally through the brain following the pattern described above.

15 PD and 14 Controls were selected for the single nuclei RNA-seq experiment. We selected individuals from both sexes (male and female), the age range is dominantly 60+. However, we included 2 young controls (30 and 38 years old), as well as 4 individuals (3 controls and 1 diagnosed with PD) aged 50-60 years old.

### Isolation of nuclei from frozen post-mortem brain tissue and single nuclei RNA-seq

To identify the substantia nigra in control and PD cases we cut cryostat sections (5 μm thickness). These sections were hematoxylin and eosin (H&E) stained and viewed under the microscope by a neuropathologist (DRT). The substantia nigra was indicated with a waterproof pen. H&E-guided 2 mm biopsy punches were collected from the SN of selected samples, nuclei were isolated and sequencing libraries were prepared using the Chromium Single Cell 3⍰ Reagent Kits v.3 according to the manufacturer’s protocol (10x Genomics). The generated scRNA-seq libraries were sequenced using NovaSeq6000 system: 1% PhiX, paired-end sequencing with 10X v3 parameters (28-8-0-91 cycles). 13 libraries having a saturation rate lower than 60% were re-sequenced with the same 10x parameters to obtain more reads and improve the quality of the database. Combining reads from both runs a >40% saturation rate for all the samples was achieved, see **Supplementary Table 1**.

### Pre-processing single nuclei RNA-seq data

Gene counts were obtained by aligning reads to the human hg38 genome (GRCh38/Ensemble 93 pre-built by 10x) using CellRanger software (v.3.0.2) (10x Genomics). To account for unspliced nuclear transcripts, reads mapping to pre-mRNA were counted (as recommended by CellRanger v.3.0.2). The quantification of pre-mRNA was done using the CellRanger count pipeline on each of the 29 individual libraries providing double fastq files as an input for the libraries that have been re-sequenced. Default CellRanger parameters have been used throughout the pipeline.

### Quality control

The initial dataset contained 173,197 nuclei reporting the transcriptome profile of 33,538 genes. At the first step of filtering thresholds to exclude degrading nuclei having low RNA-content (< 500 transcripts) and doublets having high RNA-content (> 6000 transcripts) were imposed. *MALAT1* was excluded from the transcript list since it was highly enriched and could drive the clustering.

Next, to make sure only singlets are left in the database Scrublet algorithm^52^ with default parameters was run on each donor’s dataset to identify nuclei representing doublets. In total 2496 doublets were identified (see **Supplementary Table 1**) and removed from the database.

Finally, nuclei having a high mitochondrial RNA content were considered. A first round of clustering was performed using Seurat v3.1.1 R package^7^ limiting the percentage of mitochondrial genes to 1%, 2% and 5% (data not shown), that has shown a clear bias towards grouping cells having high level of mitochondrial transcriptome together, which was very apparent in the case of 5% threshold and became less relevant for the cases of 1% and 2%. Hence, to keep maximum number of cells, but exclude the mitochondrial gene bias, nuclei having more than 2% representation of mitochondrial transcripts were filtered out leaving 82,222 nuclei with the profile of 33,537 genes for downstream analysis.

### Integration of the datasets and clustering

Data normalization and clustering were done with the Seurat package^7^. First each dataset was SCT normalized using *SCTransform()* function. The most variable 6000 features were selected for the downstream integration using *SelectIntegrationFeatures()* function. PCA analysis was performed on each dataset separately with *RunPCA()* function, with the top 50 PC coordinates evaluated. Integration anchors were found by *FindIntegrationAnchors()* method using sample s.0096 as a reference. Datasets were integrated together by *IntegrateData ()* function on top 20 dimensions. PCA analysis was repeated on the integrated database and top 10 PC components were used for UMAP analysis using *RunUMAP()* function. Finally, top 10 principal components (PC) were used to build a k-nearest-neighbours graph using *FindNeighbors()* function, 0.6 resolution was used to group nuclei in the clusters. Populations positive for well-known cell-type specific markers were grouped together to define broad cell types. Sub-clustering of neuronal and glial populations was done based on top 4 PC components at 0.2 resolution based on SCT normalized data.

### Marker identification

For each subcluster, marker genes were identified by differential gene expression analysis between the nuclei within the given subcluster and remaining nuclei in the population in consideration using Wilcoxon rank-sum test with a FDR-corrected *P* value ≤ 0.05 and a log_2_(mean gene expression across cells in sub-cluster/mean gene expression across cells in other sub-clusters) of 0.25. Transcripts detected in at least 25% of the nuclei within the given subcluster were considered. The full list of markers is reported in **Table 5**.

### Datasets of genetic risk factor associates to Parkinson’s disease

The list of genetic risk factors was extracted from GWAS EMBL catalog on 23.11.21. This is a publicly available resource of Genome Wide Association Studies (GWAS) and their results. The full list of GWAS mapping is presented in **Table 3**.

### Differential gene-expression analysis

snRNAseq-based differential expression analysis was assessed by a linear mixed model accounting for the individual of origin for nuclei using the R packages lme4. One way ANOVA test with Benjamini-Hochberg correction was performed to assess the significance of transcript’s up/down regulation using R package car. The full list of significantly up/down regulated genes detected by this analysis is reported in **Table 7**.

### Pathway analysis

Gene-enrichment and functional annotation analysis in GO and KEGG databases of subtype specific and up/down regulated genes genes were performed using a) clusterProfiler version 4.4.4 ^53^.

### Spatial transcriptomics

Spatial transcriptomics was performed by Resolve Biosciences commercially available Molcular Cartography platform. In total, 8 tissue sections from a control patient were stained with 96 highly variable transcripts at the single cell resolution (see **Tables 4**). Correspondingly images with DAPI staining and the coordinates of each transcript on the image were provided. These images were processed by QuPath 0.2.3 software to segment single nuclei based on DAPI staining. Segmented ROI were transferred to ImageJ 2.0.0-rc-43/1.52n and the transcript count per cell/ROI was extracted using Polylux_V1.6.1 ImageJ plugin developed by Resolve Biosciences. ROI coordinates were extracted using ImageJ 2.0.0-rc-43/1.52n ROI manager. Clustering was done using the default UMAP dimensionality reduction pipeline of R Seurat package v 3.2.0.

