## Supplementary material for "Unravelling cell type specific response to Parkinson’s Disease at single cell resolution": Tables

**Table 1:** The description of donors.

| N | Brain Bank ID | Sample ID | Clinical diagnosis | Age | Sex | PMI hours | RIN measure | Lewy bodies presence in midbrain | Lewy bodies presence in limbic regions (amygdala) | Lewy bodies presence in neocortical regions (frontal cortex) | CERAD score for neuritic plaques | Braak stage for neurofibrillary tangles |
|---|---|---|---|---|---|---|---|---|---|---|---|---|
| **1** | 2020 | s.0096 | Parkinson's | 90 | female | 3 | 7.8 | 1 | 1 | 1 | 1 | 4 |
| **2** | 2140 | s.0098 | Parkinson's | 99 | male | 3 | 7.0 | 1 | 1 | 1 | 1 | 4 |
| **3** | 2122 | s.0100 | Parkinson's | 89 | male | 16.5 | 6.9 | 1 | 1 | 1 | 2 | 4 |
| **4** | 2228 | s.0102 | Parkinson's | 57 | female | 3 | 6.5 | 1 | 1 | 1 | 0 | 1 |
| **5** | 2291 | s.0103 | Parkinson's | 68 | male | 61 | 6.4 | 1 | 1 | 1 | 1 | 4 |
| **6** | 2466 | s.0104 | Parkinson's | 81 | male | 48 | 6.9 | 1 | 1 | 1 | 1 | 4 |
| **7** | 2647 | s.0105 | Parkinson's | 78 | male | 12 | 6.5 | 1 | 1 | 0 | 0 | 1 |
| **8** | 2825 | s.0107 | Parkinson's | 82 | male | 24 | 7.1 | 1 | 1 | 1 | 0 | 2 |
| **9** | 2910 | s.0109 | Parkinson's | 98 | male | 20.5 | 7.0 | 1 | 1 | 1 | 0 | 3 |
| **10** | 3268 | s.0110 | Parkinson's | 82 | male | 4 | 7.3 | 1 | 1 | 1 | 0 | 3 |
| **11** | 3440 | s.0111 | Parkinson's | 80 | male | 46 | 7.5 | 1 | 1 | 1 | 1 | 2 |
| **12** | 3659 | s.0116 | Parkinson's | 77 | male | 24 | 6.3 | 1 | 1 | 0 | 0 | 1 |
| **13** | 3688 | s.0118 | Parkinson's | 85 | female | 40 | 7.7 | 1 | 1 | 1 | 1 | 4 |
| **14** | 3708 | s.0119 | Parkinson's | 84 | male | 21 | 6.4 | 1 | 1 | 1 | 2 | 4 |
| **15** | 3791 | s.0120 | Parkinson's | 85 | female | 24 | 7.5 | 1 | 1 | 1 | 0 | 3 |
| **16** | 2286 | s.0127 | Control | 65 | male | 9 | 6.8 | NA | NA | NA | NA | NA |
| **17** | 2293 | s.0128 | Control | 81 | female | 24 | 6.9 | NA | NA | NA | NA | NA |
| **18** | 2298 | s.0129 | Control | 86 | male | 13 | 6.7 | NA | NA | NA | NA | NA |
| **19** | 2309 | s.0130 | Control | 61 | male | 22 | 7.6 | NA | NA | NA | NA | NA |
| **20** | 2311 | s.0131 | Control | 65 | male | 22 | 7.0 | NA | NA | NA | NA | NA |
| **21** | 2919 | s.0142 | Control | 62 | male | 52 | 7.6 | NA | NA | NA | NA | NA |
| **22** | 2992 | s.0147 | Control | 30 | female | 12 | 8.0 | NA | NA | NA | NA | NA |
| **23** | 3337 | s.0151 | Control | 77 | female | 11 | 6.7 | NA | NA | NA | NA | NA |
| **24** | 3345 | s.0152 | Control | 93 | female | 8 | 7.7 | NA | NA | NA | NA | NA |
| **25** | 3424 | s.0153 | Control | 91 | male | 3.25 | 6.6 | NA | NA | NA | NA | NA |
| **26** | 3624 | s.0154 | Control | 93 | male | 7 | 7.1 | NA | NA | NA | NA | NA |
| **27** | 3649 | s.0158 | Control | 89 | male | 18.75 | 7.7 | NA | NA | NA | NA | NA |
| **28** | 3651 | s.0159 | Control | 73 | female | 21 | 7.7 | NA | NA | NA | NA | NA |
| **29** | 3823 | s.0165 | Control | 91 | male | 17.75 | 7.2 | NA | NA | NA | NA | NA |

**Table 2:** The output of binomial test for cell proportion analysis (major cell types, subtypes of neurons, states of astrocytes, states of microglia, states of oligodendrocytes).

| Cluster | Number of Cells | % Control origin | % Parkinson's origin | P-value (Binomial test) | Expected PD/Control ratio |
|---|---|---|---|---|---|
| Oligos | 18632 | 52.10 | 47.90 | 8,22E+02 | **0.47** |
| Astrocytes | 20710 | 49.22 | 50.78 | 1,46E−21 | **0.47** |
| Microglia | 12995 | 50.96 | 49.04 | 3,19E+00 | **0.47** |
| OPC | 6644 | 51.40 | 48.60 | 9,17E+03 | **0.47** |
| Neurons | 6196 | 72.16 | 27.84 | 4,38E−203 | **0.47** |
| Connective tissue cells | 780 | 55.13 | 44.87 | 9,41E+04 | **0.47** |
| T cells | 347 | 43.23 | 56.77 | 3,03E+02 | **0.47** |
| neurons0 | 2609 | 77.31 | 22.69 | 7,48E-04 | **0.28** |
| neurons1 | 877 | 69.66 | 30.33 | 1,32E+05 | **0.28** |
| neurons2 | 510 | 46.47 | 53.52 | 1,22E-27 | **0.28** |
| neurons3 | 341 | 80.06 | 19.94 | 7,14E+02 | **0.28** |
| neurons4 | 166 | 60.24 | 39.76 | 7,14E+02 | **0.28** |
| neurons5 | 158 | 75.95 | 24.05 | 2,89E+05 | **0.28** |
| astrocytes0 | 3991 | 49.31 | 50.69 | 3,52E+03 | **0.53** |
| astrocytes1 | 3574 | 36.68 | 63.31 | 1,49E-29 | **0.53** |
| astrocytes2 | 3547 | 94.81 | 51.88 | 0,00E+00 | **0.53** |
| astrocytes3 | 2929 | 7.06 | 92.93 | 0,00E+00 | **0.53** |
| astrocytes4 | 1844 | 36.71 | 63.29 | 5,75E-13 | **0.53** |
| astrocytes5 | 437 | 19.45 | 80.55 | 4,32E-27 | **0.53** |
| microglia0 | 3072 | 40.50 | 59.50 | 5,06E-20 | **0.52** |
| microglia1 | 2627 | 90.67 | 9.33 | 0,00E+00 | **0.52** |
| microglia2 | 1915 | 31.65 | 68.35 | 3,09E-53 | **0.52** |
| microglia3 | 1340 | 29.85 | 70.15 | 1,97E-44 | **0.52** |
| microglia4 | 859 | 41.80 | 58.20 | 1,69E+00 | **0.52** |
| microglia5 | 341 | 19.64 | 80.35 | 7,65E-25 | **0.52** |
| oligos0 | 8652 | 27.47 | 72.53 | 5,06E-20 | **0.48** |
| oligos1 | 8387 | 43.79 | 56.21 | 0,00E+00 | **0.48** |
| oligos2 | 6547 | 91.05 | 8.95 | 3,09E-53 | **0.48** |
| oligos3 | 4923 | 49.04 | 50.96 | 1,97E-44 | **0.48** |
| oligos4 | 3396 | 53.06 | 46.94 | 1,69E+00 | **0.48** |
| oligos5 | 3127 | 64.82 | 35.18 | 7,65E-25 | **0.48** |

**Table 3:** EMBL GWAS catalog for PD-associated risk factors.

| Variant and risk allele | Mapped gene | PubMed ID | First Author | Location |
|---|---|---|---|---|
| rs356182 | SNCA | 27182965 | Pickrell JK | 4:89704960 |
| rs28903073 | LRRK2 | 27182965 | Pickrell JK | 12:40259708 |
| rs365825 | LINC02210, LINC02210-CRHR1 | 27182965 | Pickrell JK | 17:45628235 |
| rs144847051 | | 27182965 | Pickrell JK | 12:33148871 |
| rs17425622 | HLA-DRB1, HLA-DQA1 | 27182965 | Pickrell JK | 6:32604184 |
| rs2230288 | GBA | 27182965 | Pickrell JK | 1:155236376 |
| rs13016703 | STK39 | 27182965 | Pickrell JK | 2:168299085 |
| rs35541465 | GAK | 27182965 | Pickrell JK | 4:879499 |
| rs11158026 | GCH1 | 27182965 | Pickrell JK | 14:54882151 |
| rs9858038 | MCCC1, DCUN1D1 | 27182965 | Pickrell JK | 3:183012503 |
| rs10256359 | KLHL7-DT, FAM126A | 27182965 | Pickrell JK | 7:23084258 |
| rs823118 | RAB29, NUCKS1 | 27182965 | Pickrell JK | 1:205754444 |
| rs188789342 | RAD1P1, TIAL1 | 27182965 | Pickrell JK | 10:119612816 |
| rs6812193 | FAM47E-STBD1, FAM47E | 27182965 | Pickrell JK | 4:76277833 |
| rs4266290 | BST1 | 27182965 | Pickrell JK | 4:15735495 |
| rs4130047 | RIT2 | 27182965 | Pickrell JK | 18:43098270 |
| rs4713118 | LINC01012 | 27182965 | Pickrell JK | 6:27709015 |
| rs11060180 | CCDC62 | 27182965 | Pickrell JK | 12:122819039 |
| rs687432 | OR5AZ1P, OR5BD1P | 27182965 | Pickrell JK | 11:57926788 |
| rs148294058 | LINC02451 | 27182965 | Pickrell JK | 12:42655580 |
| rs2209440 | SH3GL2 | 27182965 | Pickrell JK | 9:17736344 |
| rs11343 | SYT17 | 27182965 | Pickrell JK | 16:19268142 |
| rs162227 | NDUFAF2 | 27182965 | Pickrell JK | 5:60964716 |
| rs144074972 | CA8 | 27182965 | Pickrell JK | 8:60251336 |
| rs601999 | HSD17B1P1, NAGLU | 27182965 | Pickrell JK | 17:42546140 |
| rs849898 | WNT9A, CICP26 | 26227905 | Hu Y | 1:227966216 |
| rs11186 | COL5A2 | 26227905 | Hu Y | 2:189032668 |
| rs1879553 | IGSF11 | 26227905 | Hu Y | 3:118896616 |
| rs13153459 | LINC02224 | 26227905 | Hu Y | 5:44515833 |
| rs9323124 | MDGA2, RPA2P1 | 26227905 | Hu Y | 14:46996974 |
| rs1362858 | ZNF396, INO80C | 26227905 | Hu Y | 18:35406636 |
| rs10929159 | AGAP1 | 28892059 | Chang D | 2:236024319 |
| rs117896735 | INPP5F | 28892059 | Chang D | 10:119776815 |
| rs3793947 | DLG2 | 28892059 | Chang D | 11:83833429 |
| rs329648 | IGSF9B, SPATA19 | 28892059 | Chang D | 11:133895472 |
| rs76904798 | LRRK2-DT, LRRK2 | 28892059 | Chang D | 12:40220632 |
| rs943437 | LINC02527 | 28892059 | Chang D | 6:111921050 |
| rs2921073 | PRAG1 | 28892059 | Chang D | 8:8450133 |
| rs2296887 | GBF1 | 28892059 | Chang D | 10:102245653 |
| rs14235 | BCKDK | 28892059 | Chang D | 16:31110472 |
| rs17649553 | MAPT | 28892059 | Chang D | 17:45917282 |
| rs12456492 | RIT2 | 28892059 | Chang D | 18:43093415 |
| rs5910 | ITGA2B | 28892059 | Chang D | 17:44372421 |
| rs6416935 | MED13, Y_RNA | 28892059 | Chang D | 17:62107553 |
| rs823118 | RAB29, NUCKS1 | 28892059 | Chang D | 1:205754444 |
| rs10797576 | SIPA1L2 | 28892059 | Chang D | 1:232528865 |
| rs6430538 | CCNT2,AS1 | 28892059 | Chang D | 2:134782397 |
| rs12497850 | IP6K2 | 28892059 | Chang D | 3:48711556 |
| rs143918452 | ITIH1 | 28892059 | Chang D | 3:52782824 |
| rs78738012 | CAMK2D | 28892059 | Chang D | 4:113439216 |
| rs2694528 | NDUFAF2 | 28892059 | Chang D | 5:60978096 |
| rs9468199 | LINC01012, GPR89P | 28892059 | Chang D | 6:27713436 |
| rs2740594 | CTSB | 28892059 | Chang D | 8:11849665 |
| rs9275326 | HLA-DQB1, MTCO3P1 | 28892059 | Chang D | 6:32698883 |
| rs199347 | GPNMB | 28892059 | Chang D | 7:23254127 |
| rs591323 | RN7SL474P | 28892059 | Chang D | 8:16839582 |
| rs67460515 | SPTSSB | 28892059 | Chang D | 3:161350778 |
| rs10463554 | PAM | 28892059 | Chang D | 5:102983270 |

| | | | | |
|---|---|---|---|---|
| rs17767294 | ZSCAN16-AS1, ZNF165 | 28892059 | Chang D | 6:28086420 |
| rs11060180 | CCDC62 | 28892059 | Chang D | 12:122819039 |
| rs11158026 | GCH1 | 28892059 | Chang D | 14:54882151 |
| rs1555399 | TMEM229B | 28892059 | Chang D | 14:67517653 |
| rs2414739 | LINC02349 | 28892059 | Chang D | 15:61701935 |
| rs9568188 | CAB39L | 28892059 | Chang D | 13:49353596 |
| rs8017172 | RPL13AP3, KTN1 | 28892059 | Chang D | 14:55732330 |
| rs316619 | LTK | 28892059 | Chang D | 15:41506416 |
| rs62120679 | TMPRSS9 | 28892059 | Chang D | 19:2363321 |
| rs8118008 | DDRGK1, LZTS3 | 28892059 | Chang D | 20:3187520 |
| rs4653767 | ITPKB | 28892059 | Chang D | 1:226728377 |
| rs34043159 | MAP4K4 | 28892059 | Chang D | 2:101796654 |
| rs353116 | SCN2A | 28892059 | Chang D | 2:165277122 |
| rs4073221 | TBC1D5 | 28892059 | Chang D | 3:18235996 |
| rs1474055 | STK39 | 28892059 | Chang D | 2:168253884 |
| rs12637471 | MCCC1 | 28892059 | Chang D | 3:183044649 |
| rs34311866 | TMEM175 | 28892059 | Chang D | 4:958159 |
| rs11724635 | BST1 | 28892059 | Chang D | 4:15735478 |
| rs6812193 | FAM47E-STBD1, FAM47E | 28892059 | Chang D | 4:76277833 |
| rs356182 | SNCA | 28892059 | Chang D | 4:89704960 |
| rs2280104 | BIN3 | 28892059 | Chang D | 8:22668467 |
| rs13294100 | SH3GL2 | 28892059 | Chang D | 9:17579692 |
| rs10906923 | ITGA8 | 28892059 | Chang D | 10:15527599 |
| rs8005172 | GPR65 | 28892059 | Chang D | 14:88006268 |
| rs11343 | SYT17 | 28892059 | Chang D | 16:19268142 |
| rs4784227 | CASC16 | 28892059 | Chang D | 16:52565276 |
| rs12600861 | CHRNB1 | 31701892 | Nalls MA | 17:7452302 |
| rs12951632 | RETREG3 | 31701892 | Nalls MA | 17:42588995 |
| rs2269906 | UBTF | 31701892 | Nalls MA | 17:44216969 |
| rs850738 | FAM171A2 | 31701892 | Nalls MA | 17:44357262 |
| rs62053943 | LINC02210, CRHR1 | 31701892 | Nalls MA | 17:45666837 |
| rs117615688 | LINC02210, CRHR1 | 31701892 | Nalls MA | 17:45720942 |
| rs11658976 | WNT3 | 31701892 | Nalls MA | 17:46789439 |
| rs61169879 | BRIP1 | 31701892 | Nalls MA | 17:61840005 |
| rs666463 | DNAH17 | 31701892 | Nalls MA | 17:78429399 |
| rs1941685 | ASXL3 | 31701892 | Nalls MA | 18:33724354 |
| rs12456492 | RIT2 | 31701892 | Nalls MA | 18:43093415 |
| rs8087969 | SRSF10P1, SMAD4 | 31701892 | Nalls MA | 18:51157219 |
| rs55818311 | SPPL2B | 31701892 | Nalls MA | 19:2341049 |
| rs77351827 | CRLS1 | 31701892 | Nalls MA | 20:6025395 |
| rs2248244 | DYRK1A | 31701892 | Nalls MA | 21:37480059 |
| rs35643925 | SEMA4A, SLC25A44 | 31701892 | Nalls MA | 1:156185069 |
| rs4954162 | TMEM163 | 31701892 | Nalls MA | 2:134681219 |
| rs356228 | SNCA | 31701892 | Nalls MA | 4:89685975 |
| rs356203 | SNCA | 31701892 | Nalls MA | 4:89744890 |
| rs6875262 | LINC02240, ZNF608 | 31701892 | Nalls MA | 5:124774580 |
| rs9267659 | SLC44A4, EHMT2-AS1 | 31701892 | Nalls MA | 6:31878457 |
| rs181609621 | DNM1L, FGD4 | 31701892 | Nalls MA | 12:32657825 |
| rs117073808 | MUC19 | 31701892 | Nalls MA | 12:40528770 |
| rs141128804 | MUC19 | 31701892 | Nalls MA | 12:40542837 |
| rs896435 | ITGA8 | 31701892 | Nalls MA | 10:15515407 |
| rs10748818 | GBF1 | 31701892 | Nalls MA | 10:102255522 |
| rs72840788 | BAG3 | 31701892 | Nalls MA | 10:119656173 |
| rs117896735 | INPP5F | 31701892 | Nalls MA | 10:119776815 |
| rs7938782 | RNF141 | 31701892 | Nalls MA | 11:10537230 |
| rs12283611 | DLG2 | 31701892 | Nalls MA | 11:83776234 |
| rs3802920 | IGSF9B | 31701892 | Nalls MA | 11:133917106 |
| rs76904798 | LRRK2-DT, LRRK2 | 31701892 | Nalls MA | 12:40220632 |
| rs34637584 | LRRK2 | 31701892 | Nalls MA | 12:40340400 |
| rs7134559 | NCKAP5L, SCAF11 | 31701892 | Nalls MA | 12:46025303 |
| rs10847864 | HIP1R | 31701892 | Nalls MA | 12:122842051 |
| rs11610045 | FBRSL1 | 31701892 | Nalls MA | 12:132487182 |
| rs9568188 | CAB39L | 31701892 | Nalls MA | 13:49353596 |

| | | | | |
|---|---|---|---|---|
| rs4771268 | MBNL2, LINC00456 | 31701892 | Nalls MA | 13:97212767 |
| rs12147950 | MIPOL1 | 31701892 | Nalls MA | 14:37520065 |
| rs11158026 | GCH1 | 31701892 | Nalls MA | 14:54882151 |
| rs3742785 | RPS6KL1 | 31701892 | Nalls MA | 14:74906331 |
| rs979812 | GPR65, RNU6-835P | 31701892 | Nalls MA | 14:87997920 |
| rs2251086 | LINC02349 | 31701892 | Nalls MA | 15:61705186 |
| rs6497339 | SYT17 | 31701892 | Nalls MA | 16:19266171 |
| rs2904880 | RABEP2, CD19 | 31701892 | Nalls MA | 16:28933075 |
| rs11150601 | SETD1A | 31701892 | Nalls MA | 16:30966478 |
| rs6500328 | NOD2 | 31701892 | Nalls MA | 16:50702745 |
| rs3104783 | CASC16 | 31701892 | Nalls MA | 16:52602330 |
| rs10221156 | PHBP21 | 31701892 | Nalls MA | 16:52935514 |
| rs6825004 | SCARB2 | 31701892 | Nalls MA | 4:76189212 |
| rs6854006 | FAM47E-STBD1, FAM47E | 31701892 | Nalls MA | 4:76276901 |
| rs356182 | SNCA | 31701892 | Nalls MA | 4:89704960 |
| rs5019538 | SNCA | 31701892 | Nalls MA | 4:89715479 |
| rs13117519 | CAMK2D | 31701892 | Nalls MA | 4:113447909 |
| rs62333164 | CLCN3 | 31701892 | Nalls MA | 4:169662006 |
| rs1867598 | ELOVL7 | 31701892 | Nalls MA | 5:60842132 |
| rs26431 | EIF3KP1, PAM | 31701892 | Nalls MA | 5:103030090 |
| rs11950533 | C5orf24, TXNDC15 | 31701892 | Nalls MA | 5:134863415 |
| rs4140646 | RSL24D1P1, GPR89P | 31701892 | Nalls MA | 6:27771022 |
| rs9261484 | TRIM40 | 31701892 | Nalls MA | 6:30140906 |
| rs57891859 | TMEM163 | 31701892 | Nalls MA | 2:134707046 |
| rs1474055 | STK39 | 31701892 | Nalls MA | 2:168253884 |
| rs73038319 | TBC1D5 | 31701892 | Nalls MA | 3:18320267 |
| rs6808178 | RBMS3 | 31701892 | Nalls MA | 3:28664199 |
| rs12497850 | IP6K2 | 31701892 | Nalls MA | 3:48711556 |
| rs55961674 | KPNA1 | 31701892 | Nalls MA | 3:122478045 |
| rs11707416 | MED12L | 31701892 | Nalls MA | 3:151391177 |
| rs620513 | RN7SL474P | 31701892 | Nalls MA | 8:16840084 |
| rs10513789 | MCCC1 | 31701892 | Nalls MA | 3:183042285 |
| rs873786 | GAK | 31701892 | Nalls MA | 4:931588 |
| rs34311866 | TMEM175 | 31701892 | Nalls MA | 4:958159 |
| rs4698412 | BST1 | 31701892 | Nalls MA | 4:15735725 |
| rs34025766 | LCORL | 31701892 | Nalls MA | 4:17967188 |
| rs114138760 | PMVK | 31701892 | Nalls MA | 1:154925709 |
| rs4101061 | FAM47E, SCARB2 | 31701892 | Nalls MA | 4:76226816 |
| rs76763715 | GBA | 31701892 | Nalls MA | 1:155235843 |
| rs6658353 | RNU6-481P, FCGR2A | 31701892 | Nalls MA | 1:161499264 |
| rs11578699 | VAMP4, EEF1AKNMT | 31701892 | Nalls MA | 1:171750629 |
| rs823118 | RAB29, NUCKS1 | 31701892 | Nalls MA | 1:205754444 |
| rs11557080 | RAB29 | 31701892 | Nalls MA | 1:205768611 |
| rs4653767 | ITPKB | 31701892 | Nalls MA | 1:226728377 |
| rs10797576 | SIPA1L2 | 31701892 | Nalls MA | 1:232528865 |
| rs76116224 | KCNS3 | 31701892 | Nalls MA | 2:17966582 |
| rs2042477 | KCNIP3 | 31701892 | Nalls MA | 2:95335195 |
| rs11683001 | MAP4K4 | 31701892 | Nalls MA | 2:101780501 |
| rs112485576 | HLA-DQA1, HLA-DRB1 | 31701892 | Nalls MA | 6:32610995 |
| rs997368 | LINC02527 | 31701892 | Nalls MA | 6:111922088 |
| rs75859381 | LINC00326, HMGB1P13 | 31701892 | Nalls MA | 6:132889222 |
| rs199351 | GPNMB | 31701892 | Nalls MA | 7:23260430 |
| rs76949143 | SAPCD2P3, GS1-124K5.4 | 31701892 | Nalls MA | 7:66544864 |
| rs1293298 | CTSB | 31701892 | Nalls MA | 8:11854934 |
| rs1450522 | SPTSSB | 31701892 | Nalls MA | 3:161359842 |
| rs2280104 | BIN3 | 31701892 | Nalls MA | 8:22668467 |
| rs2086641 | CYRIB | 31701892 | Nalls MA | 8:129889663 |
| rs13294100 | SH3GL2 | 31701892 | Nalls MA | 9:17579692 |
| rs10756907 | SH3GL2 | 31701892 | Nalls MA | 9:17727067 |
| rs6476434 | UBAP2 | 31701892 | Nalls MA | 9:34046393 |
| rs943437 | LINC02527 | 32201043 | Smeland OB | 6:111921050 |
| rs34372695 | LAMTOR2, RAB25 | 32201043 | Smeland OB | 1:156060246 |
| rs1293298 | CTSB | 32201043 | Smeland OB | 8:11854934 |

| | | | | |
|---|---|---|---|---|
| rs11203810 | RN7SL474P | 32201043 | Smeland OB | 8:16854852 |
| rs4653767 | ITPKB | 32201043 | Smeland OB | 1:226728377 |
| rs10797576 | SIPA1L2 | 32201043 | Smeland OB | 1:232528865 |
| rs12467316 | LINC01127, IL1R2 | 32201043 | Smeland OB | 2:101988187 |
| rs10906923 | ITGA8 | 32201043 | Smeland OB | 10:15527599 |
| rs4667572 | STK39 | 32201043 | Smeland OB | 2:168299490 |
| rs3793947 | DLG2 | 32201043 | Smeland OB | 11:83833429 |
| rs13078687 | TBC1D5 | 32201043 | Smeland OB | 3:18181950 |
| rs77669894 | BICD1, RESF1 | 32201043 | Smeland OB | 12:32089513 |
| rs28370664 | SLC2A13 | 32201043 | Smeland OB | 12:40019503 |
| rs1450522 | SPTSSB | 32201043 | Smeland OB | 3:161359842 |
| rs140278703 | MCCC1 | 32201043 | Smeland OB | 3:183034255 |
| rs34884217 | TMEM175 | 32201043 | Smeland OB | 4:950422 |
| rs12651314 | CD38, BST1 | 32201043 | Smeland OB | 4:15751947 |
| rs3889108 | LINC02349 | 32201043 | Smeland OB | 15:61712597 |
| rs13107325 | SLC39A8 | 32201043 | Smeland OB | 4:102267552 |
| rs17483653 | CAMK2D | 32201043 | Smeland OB | 4:113422713 |
| rs1867598 | ELOVL7 | 32201043 | Smeland OB | 5:60842132 |
| rs6875262 | LINC02240, ZNF608 | 32201043 | Smeland OB | 5:124774580 |
| rs2647062 | HLA-DQA1, HLA-DRB1 | 32201043 | Smeland OB | 6:32602640 |
| rs302714 | RERE, RERE-AS1 | 32201043 | Smeland OB | 1:8426071 |
| rs28624974 | KLHL7 | 32201043 | Smeland OB | 7:23149317 |
| rs4915466 | MROH3P | 32201043 | Smeland OB | 1:200936379 |
| rs3805 | NUCKS1 | 32201043 | Smeland OB | 1:205714227 |
| rs4921739 | ZDHHC2 | 32201043 | Smeland OB | 8:17172148 |
| rs13294100 | SH3GL2 | 32201043 | Smeland OB | 9:17579692 |
| rs2278973 | CACNA1B | 32201043 | Smeland OB | 9:138121810 |
| rs6430538 | CCNT2, AS1 | 32201043 | Smeland OB | 2:134782397 |
| rs144814361 | BAG3 | 32201043 | Smeland OB | 10:119651405 |
| rs113732002 | NGEF | 32201043 | Smeland OB | 2:232982170 |
| rs3802920 | IGSF9B | 32201043 | Smeland OB | 11:133917106 |
| rs55740225 | PARP9 | 32201043 | Smeland OB | 3:122549781 |
| rs138895122 | ASS1P14 | 32201043 | Smeland OB | 12:33075690 |
| rs2276765 | MED12L | 32201043 | Smeland OB | 3:151413434 |
| rs144029582 | MTND1P24, LINC02400 | 32201043 | Smeland OB | 12:41754811 |
| rs11060112 | CCDC62 | 32201043 | Smeland OB | 12:122807956 |
| rs11158026 | GCH1 | 32201043 | Smeland OB | 14:54882151 |
| rs8005172 | GPR65 | 32201043 | Smeland OB | 14:88006268 |
| rs12643261 | FAM47E-STBD1, FAM47E | 32201043 | Smeland OB | 4:76255695 |
| rs1027647 | TPM1, LACTB | 32201043 | Smeland OB | 15:63082626 |
| rs140820592 | STX1B | 32201043 | Smeland OB | 16:31010559 |
| rs113564729 | KANSL1 | 32201043 | Smeland OB | 17:46101729 |
| rs148534175 | MED13, Y_RNA | 32201043 | Smeland OB | 17:62120839 |
| rs12456492 | RIT2 | 32201043 | Smeland OB | 18:43093415 |
| rs8087969 | SRSF10P1, SMAD4 | 32201043 | Smeland OB | 18:51157219 |
| rs6060983 | MYLK2 | 32201043 | Smeland OB | 20:31833121 |
| rs11701722 | DYRK1A | 32201043 | Smeland OB | 21:37398228 |
| rs56379273 | TRPM2 | 32201043 | Smeland OB | 21:44406603 |
| rs2979160 | PRAG1 | 32201043 | Smeland OB | 8:8450156 |
| rs9468195 | LINC01012 | 32201043 | Smeland OB | 6:27708530 |
| rs6826785 | SNCA | 32310270 | Foo JN | 4:89761323 |
| rs6679073 | RAB29, SLC41A1 | 32310270 | Foo JN | 1:205787356 |
| rs2292056 | MCCC1, MCCC1-AS1 | 32310270 | Foo JN | 3:183017423 |
| rs16846351 | ITPKB, ITPKB-IT1 | 32310270 | Foo JN | 1:226659011 |
| rs12278023 | DLG2 | 32310270 | Foo JN | 11:83799074 |
| rs1887316 | FYN | 32310270 | Foo JN | 6:111830249 |
| rs4130047 | RIT2 | 32310270 | Foo JN | 18:43098270 |
| rs6449168 | BST1 | 32310270 | Foo JN | 4:15726090 |
| rs34311866 | TMEM175 | 32310270 | Foo JN | 4:958159 |
| rs246814 | SV2C | 32310270 | Foo JN | 5:76303383 |
| rs356182 | SNCA | 31660654 | Bandres-Ciga S | 4:89704960 |
| rs190807041 | MUC19, LRRK2 | 31660654 | Bandres-Ciga S | 12:40379882 |
| rs113434679 | KANSL1 | 31660654 | Bandres-Ciga S | 17:46049399 |

| | | | | |
|---|---|---|---|---|
| rs9275152 | MTCO3P1, HLA-DQB1 | 31660654 | Bandres-Ciga S | 6:32684419 |
| rs62053943 | LINC02210, CRHR1 | 31660654 | Bandres-Ciga S | 17:45666837 |
| rs34311866 | TMEM175 | 31660654 | Bandres-Ciga S | 4:958159 |
| rs11150601 | SETD1A | 31660654 | Bandres-Ciga S | 16:30966478 |
| rs144755950 | GXYLT1 | 31701892 | Nalls MA | 12:42141601 |
| rs17686238 | RNA5SP443, ARHGAP27 | 31701892 | Nalls MA | 17:45339907 |
| rs9912362 | LINC02210, CRHR1 | 31701892 | Nalls MA | 17:45706862 |
| rs7225002 | KANSL1 | 31701892 | Nalls MA | 17:46111701 |
| rs199453 | NSF | 31701892 | Nalls MA | 17:46723580 |
| rs2295545 | DDRGK1, LZTS3 | 31701892 | Nalls MA | 20:3184040 |
| rs356219 | SNCA | 31755958 | Blauwendraat C | 4:89716450 |
| rs1293298 | CTSB | 31755958 | Blauwendraat C | 8:11854934 |
| rs6599388 | TMEM175 | 21292315 | Nalls MA | 4:945299 |
| rs11724635 | BST1 | 21292315 | Nalls MA | 4:15735478 |
| rs1491942 | LRRK2 | 21292315 | Nalls MA | 12:40227006 |
| rs34372695 | LAMTOR2, RAB25 | 21292315 | Nalls MA | 1:156060246 |
| rs6710823 | CCNT2, AS1 | 21292315 | Nalls MA | 2:134834811 |
| rs11711441 | MCCC1 | 21292315 | Nalls MA | 3:183103487 |
| rs356219 | SNCA | 21292315 | Nalls MA | 4:89716450 |
| rs2102808 | STK39 | 21292315 | Nalls MA | 2:168260515 |
| rs12817488 | CCDC62 | 21292315 | Nalls MA | 12:122811747 |
| rs2942168 | LINC02210, LINC02210-CRHR1 | 21292315 | Nalls MA | 17:45637484 |
| rs1223271 | ISM1, TASP1 | 19915575 | Simón-Sánchez J | 20:13316265 |
| rs6812193 | FAM47E-STBD1, FAM47E | 19915575 | Simón-Sánchez J | 4:76277833 |
| rs7077361 | ITGA8 | 19915575 | Simón-Sánchez J | 10:15519544 |
| rs2736990 | SNCA | 19915575 | Simón-Sánchez J | 4:89757390 |
| rs199533 | NSF | 19915575 | Simón-Sánchez J | 17:46751565 |
| rs393152 | LINC02210-CRHR1, LINC02210 | 19915575 | Simón-Sánchez J | 17:45641777 |
| rs823128 | NUCKS1 | 19915575 | Simón-Sánchez J | 1:205744250 |
| rs17115100 | CYP17A1, WBP1L | 19915575 | Simón-Sánchez J | 10:102831636 |
| rs6532197 | MMRN1, SNCA-AS1 | 19915575 | Simón-Sánchez J | 4:89876150 |
| rs12431733 | LINC02331 | 19915575 | Simón-Sánchez J | 14:53824112 |
| rs1480597 | | 17052657 | Fung HC | 10:44665661 |
| rs2242330 | STAP1 | 17052657 | Fung HC | 4:67581531 |
| rs10501570 | DLG2 | 17052657 | Fung HC | 11:84706803 |
| rs7702187 | SEMA5A | 16252231 | Maraganore DM | 5:9332169 |
| rs1564282 | GAK | 18985386 | Pankratz N | 4:858525 |
| rs11012 | PLEKHM1 | 20070850 | Edwards TL | 17:45436075 |
| rs12063142 | TAS1R2, PAX7 | 20070850 | Edwards TL | 1:18813023 |
| rs4837628 | BRINP1 | 20070850 | Edwards TL | 9:119297431 |
| rs11248060 | DGKQ | 20070850 | Edwards TL | 4:970571 |
| rs10464059 | RNU6-525P, GFPT2 | 20070850 | Edwards TL | 5:180367208 |
| rs2736990 | SNCA | 20070850 | Edwards TL | 4:89757390 |
| rs1442190 | CNTN1 | 24842889 | Vacic V | 12:40971838 |
| rs17577094 | KANSL1 | 24842889 | Vacic V | 17:46110126 |
| rs1630500 | KCNN3, PMVK | 24842889 | Vacic V | 1:154882579 |
| rs823114 | RAB29, NUCKS1 | 24842889 | Vacic V | 1:205750404 |
| rs11158026 | GCH1 | 25064009 | Nalls MA | 14:54882151 |
| rs1555399 | TMEM229B | 25064009 | Nalls MA | 14:67517653 |
| rs2414739 | LINC02349 | 25064009 | Nalls MA | 15:61701935 |
| rs14235 | BCKDK | 25064009 | Nalls MA | 16:31110472 |
| rs17649553 | MAPT | 25064009 | Nalls MA | 17:45917282 |
| rs12456492 | RIT2 | 25064009 | Nalls MA | 18:43093415 |
| rs62120679 | TMPRSS9 | 25064009 | Nalls MA | 19:2363321 |
| rs8118008 | DDRGK1, LZTS3 | 25064009 | Nalls MA | 20:3187520 |
| rs591323 | RN7SL474P | 25064009 | Nalls MA | 8:16839582 |
| rs823118 | RAB29, NUCKS1 | 25064009 | Nalls MA | 1:205754444 |
| rs10797576 | SIPA1L2 | 25064009 | Nalls MA | 1:232528865 |
| rs6430538 | CCNT2, AS1 | 25064009 | Nalls MA | 2:134782397 |
| rs1474055 | STK39 | 25064009 | Nalls MA | 2:168253884 |
| rs12637471 | MCCC1 | 25064009 | Nalls MA | 3:183044649 |

| | | | | |
|---|---|---|---|---|
| rs34311866 | TMEM175 | 25064009 | Nalls MA | 4:958159 |
| rs11724635 | BST1 | 25064009 | Nalls MA | 4:15735478 |
| rs6812193 | FAM47E-STBD1, FAM47E | 25064009 | Nalls MA | 4:76277833 |
| rs356182 | SNCA | 25064009 | Nalls MA | 4:89704960 |
| rs9275326 | HLA-DQB1, MTCO3P1 | 25064009 | Nalls MA | 6:32698883 |
| rs199347 | GPNMB | 25064009 | Nalls MA | 7:23254127 |
| rs117896735 | INPP5F | 25064009 | Nalls MA | 10:119776815 |
| rs3793947 | DLG2 | 25064009 | Nalls MA | 11:83833429 |
| rs329648 | IGSF9B, SPATA19 | 25064009 | Nalls MA | 11:133895472 |
| rs76904798 | LRRK2-DT, LRRK2 | 25064009 | Nalls MA | 12:40220632 |
| rs3773384 | CHL1, CHL1-AS1 | 29724592 | Pottier C | 3:374758 |
| rs34656641 | SP1 | 29724592 | Pottier C | 12:53403452 |
| rs17833740 | LINC01500 | 29724592 | Pottier C | 14:59154251 |
| rs356182 | SNCA | 34227697 | Loesch DP | 4:89704960 |
| rs34311866 | TMEM175 | 34064523 | Rodrigo LM | 4:958159 |
| rs983361 | SNCA,AS1 | 34064523 | Rodrigo LM | 4:89840793 |
| rs3806789 | SNCA,AS1 | 34064523 | Rodrigo LM | 4:89838405 |
| rs74125084 | ING1 | 34064523 | Rodrigo LM | 13:110720333 |
| rs11653889 | ITGAE, HASPIN | 34064523 | Rodrigo LM | 17:3724162 |
| rs116504637 | CYP4Z1 | 33987465 | Alfradique-Dunham I | 1:47092261 |
| rs11949046 | GABRG2 | 33987465 | Alfradique-Dunham I | 5:162251793 |
| rs78926797 | TMC5 | 33987465 | Alfradique-Dunham I | 16:19491341 |
| rs6932127 | NKAIN2, RNF217-AS1 | 33987465 | Alfradique-Dunham I | 6:124786353 |
| rs55971529 | FANCF, LINC01495 | 33987465 | Alfradique-Dunham I | 11:22561522 |
| rs988295487 | RNU6-161P, TPI1P1 | 33987465 | Alfradique-Dunham I | 1:76725240 |
| rs13330839 | RN7SKP190 | 33987465 | Alfradique-Dunham I | 16:82246021 |
| rs356220 | SNCA | 21084426 | Saad M | 4:89720189 |
| rs4698412 | BST1 | 21084426 | Saad M | 4:15735725 |
| rs4964469 | POLR3B, RFX4 | 21084426 | Saad M | 12:106556209 |
| rs6812193 | FAM47E-STBD1, FAM47E | 21738487 | Do CB | 4:76277833 |
| rs356220 | SNCA | 21738487 | Do CB | 4:89720189 |
| rs12185268 | SPPL2C, MAPT-AS1 | 21738487 | Do CB | 17:45846317 |
| rs10513789 | MCCC1 | 21738487 | Do CB | 3:183042285 |
| rs6599389 | TMEM175 | 21738487 | Do CB | 4:945325 |
| rs4130047 | RIT2 | 21738487 | Do CB | 18:43098270 |
| rs11868035 | SREBF1 | 21738487 | Do CB | 17:17811787 |
| rs34637584 | LRRK2 | 21738487 | Do CB | 12:40340400 |
| rs823156 | SLC41A1 | 21738487 | Do CB | 1:205795512 |
| rs2823357 | RNU6-1326P, CYCSP42 | 21738487 | Do CB | 21:15542586 |
| rs10958605 | TCIM | 22658654 | Chung SJ | 8:40196086 |
| rs959573 | LMNB1, MARCHF3 | 22658654 | Chung SJ | 5:126846170 |
| rs6482992 | CLRN3 | 22658654 | Chung SJ | 10:127880410 |
| rs17000647 | ODAPH | 22658654 | Chung SJ | 4:75556520 |
| rs3027247 | CTC1 | 22658654 | Chung SJ | 17:8227549 |
| rs2839398 | PRDM15 | 22451204 | Pankratz N | 21:41866243 |
| rs356220 | SNCA | 22451204 | Pankratz N | 4:89720189 |
| rs9917256 | STK39 | 22451204 | Pankratz N | 2:168286525 |
| rs4698412 | BST1 | 22451204 | Pankratz N | 4:15735725 |
| rs2275336 | CNKSR3 | 22451204 | Pankratz N | 6:154405854 |
| rs2395163 | TSBP1-AS1, HLA-DRA | 22451204 | Pankratz N | 6:32420032 |
| rs12726330 | SLC50A1 | 22451204 | Pankratz N | 1:155135691 |
| rs6430538 | CCNT2, AS1 | 22451204 | Pankratz N | 2:134782397 |
| rs11248060 | DGKQ | 22451204 | Pankratz N | 4:970571 |
| rs1296028 | CTSB, FDFT1 | 22451204 | Pankratz N | 8:11841238 |
| rs1536076 | SH3GL2 | 22451204 | Pankratz N | 9:17731923 |
| rs11026412 | ANO5 | 22451204 | Pankratz N | 11:22092547 |
| rs10877840 | SLC2A13 | 22451204 | Pankratz N | 12:39959194 |
| rs10519131 | LINC02349 | 22451204 | Pankratz N | 15:61708933 |
| rs11865038 | PRSS53, ZNF646 | 22451204 | Pankratz N | 16:31083850 |
| rs199515 | WNT3 | 22451204 | Pankratz N | 17:46779275 |
| rs12456492 | RIT2 | 22451204 | Pankratz N | 18:43093415 |
| rs2338971 | LINC01307 | 24511991 | Hill-Burns EM | 1:101414449 |
| rs356220 | SNCA | 24511991 | Hill-Burns EM | 4:89720189 |

| | | | | |
|---|---|---|---|---|
| rs356220 | SNCA | 24511991 | Hill-Burns EM | 4:89720189 |
| rs3129882 | HLA, DRA | 24511991 | Hill-Burns EM | 6:32441753 |
| rs3129882 | HLA, DRA | 24511991 | Hill-Burns EM | 6:32441753 |
| rs199498 | WNT3 | 24511991 | Hill-Burns EM | 17:46788237 |
| rs199498 | WNT3 | 24511991 | Hill-Burns EM | 17:46788237 |
| rs2338971 | LINC01307 | 24511991 | Hill-Burns EM | 1:101414449 |
| rs183211 | NSF | 21812969 | Liu X | 17:46710944 |
| rs10121009 | UNC13B | 21812969 | Liu X | 9:35269822 |
| rs415430 | WNT3 | 21812969 | Liu X | 17:46781778 |
| rs7118648 | CCDC82 | 23793441 | Davis MF | 11:96375264 |
| rs3935740 | TMC3-AS1, TMC3 | 23793441 | Davis MF | 15:81334384 |
| rs17497526 | COL13A1 | 23793441 | Davis MF | 10:69820364 |
| rs11060180 | CCDC62 | 25064009 | Nalls MA | 12:122819039 |
| rs2270968 | MCCC1 | 28011712 | Foo JN | 3:183037421 |
| rs8180209 | SNCA | 28011712 | Foo JN | 4:89723303 |
| rs1384236 | SLC2A13 | 28011712 | Foo JN | 12:40064582 |
| rs7479949 | DLG2 | 28011712 | Foo JN | 11:83792361 |
| rs11931074 | SNCA | 19915576 | Satake W | 4:89718364 |
| rs4538475 | BST1 | 19915576 | Satake W | 4:15736314 |
| rs1994090 | SLC2A13 | 19915576 | Satake W | 12:40034759 |
| rs947211 | SLC41A1, RAB29 | 19915576 | Satake W | 1:205783537 |
| rs356220 | SNCA | 20711177 | Hamza TH | 4:89720189 |
| rs11248051 | GAK | 20711177 | Hamza TH | 4:864544 |
| rs199533 | NSF | 20711177 | Hamza TH | 17:46751565 |
| rs3129882 | HLA, DRA | 20711177 | Hamza TH | 6:32441753 |
| rs34778348 | LRRK2 | 22438815 | Lill CM | 12:40363526 |
| rs1491942 | LRRK2 | 22438815 | Lill CM | 12:40227006 |
| rs11248060 | DGKQ | 22438815 | Lill CM | 4:970571 |
| rs356219 | SNCA | 22438815 | Lill CM | 4:89716450 |
| rs6532194 | MMRN1, SNCA-AS1 | 22438815 | Lill CM | 4:89859751 |
| rs8070723 | MAPT | 21044948 | Spencer CC | 17:46003698 |
| rs356220 | SNCA | 21044948 | Spencer CC | 4:89720189 |
| rs35643925 | SEMA4A, SLC25A44 | 34594039 | Sakaue S | 1:156185069 |
| rs356220 | SNCA | 34594039 | Sakaue S | 4:89720189 |
| rs344650 | LHFPL2 | 27402877 | Hill-Burns EM | 5:78564785 |
| rs74335301 | TRPS1 | 27402877 | Hill-Burns EM | 8:115626410 |
| rs79503702 | KLHDC1 | 27402877 | Hill-Burns EM | 14:49739452 |
| rs117267308 | TPM1 | 27402877 | Hill-Burns EM | 15:63054177 |
| rs356203 | SNCA | 31660654 | Bandres-Ciga S | 4:89744890 |
| rs9356013 | PRKN | 31660654 | Bandres-Ciga S | 6:162403694 |
| rs67383717 | LINC00476 | 27545685 | Biernacka JM | 9:95864266 |
| rs10253857 | ZNF789 | 33111402 | Tan MMX | 7:17673826 |
| rs13424530 | SULT1C2 | 33111402 | Tan MMX | 2:108292945 |
| rs4802739 | GPR32P1, GPR32 | 33111402 | Tan MMX | 19:50760039 |
| rs17554587 | SQOR | 33111402 | Tan MMX | 15:45744252 |
| rs17367669 | ZNF474, AS1 | 33111402 | Tan MMX | 5:122191027 |
| rs5870994 | ZNF474, AS1 | 33111402 | Tan MMX | 5:122193659 |
| rs7870456 | PTPRD | 33111402 | Tan MMX | 9:8454921 |
| rs72767442 | MCTP2 | 33111402 | Tan MMX | 15:94320087 |
| rs6741991 | KLHL29 | 33111402 | Tan MMX | 2:23493673 |
| rs35950207 | RNU6-121P, AQP10 | 33111402 | Tan MMX | 1:154319482 |
| rs74709761 | ANO2 | 33111402 | Tan MMX | 12:5829410 |
| rs12813102 | GPR19 | 33111402 | Tan MMX | 12:12677103 |
| rs4128840 | THSD4 | 33111402 | Tan MMX | 15:71520619 |
| rs429358 | APOE | 33111402 | Tan MMX | 19:44908684 |
| rs143371462 | SLCO1B3 | 33111402 | Tan MMX | 12:20812884 |
| rs113730632 | NR1D2 | 33111402 | Tan MMX | 3:23951314 |
| rs34105455 | DENND2B | 33111402 | Tan MMX | 11:8882396 |
| rs2956605 | CRISPLD1 | 33111402 | Tan MMX | 8:74970819 |
| rs5755468 | LINC02885 | 33111402 | Tan MMX | 22:34970241 |
| rs148603475 | NTRK2 | 33111402 | Tan MMX | 9:84627637 |
| rs1124933 | Y_RNA, RNU7-137P | 33111402 | Tan MMX | 20:18097908 |
| rs6488987 | AACS | 33111402 | Tan MMX | 12:125083207 |

| | | | | |
|---|---|---|---|---|
| rs17092224 | ASS1P13, DRAP1 | 33111402 | Tan MMX | 11:107127349 |
| rs429358 | APOE | 33111402 | Tan MMX | 19:44908684 |
| rs224750 | BCCIP, LINC02629 | 33111402 | Tan MMX | 10:33942102 |
| rs11634227 | MCTP2 | 33111402 | Tan MMX | 15:94318611 |
| rs79987229 | FAM184A | 33111402 | Tan MMX | 6:119112570 |
| rs62343939 | LINC02114 | 33111402 | Tan MMX | 5:4699328 |
| rs73656147 | PLPPR1 | 30338293 | Wallen ZD | 9:101115169 |
| rs9783733 | PWRN4 | 33884653 | Li C | 15:23923803 |
| rs3775458 | SNCA | 33884653 | Li C | 4:89818097 |
| rs356203 | SNCA | 30957308 | Blauwendraat C | 4:89744890 |
| rs983361 | SNCA, AS1 | 30957308 | Blauwendraat C | 4:89840793 |
| rs34311866 | TMEM175 | 30957308 | Blauwendraat C | 4:958159 |
| rs429358 | APOE | 30957308 | Blauwendraat C | 19:44908684 |
| rs1941184 | DSG3 | 19772629 | Latourelle JC | 18:31458160 |
| rs17565841 | OCA2 | 19772629 | Latourelle JC | 15:27752101 |
| rs10918270 | ATF6 | 19772629 | Latourelle JC | 1:161945711 |
| rs10767971 | PRRG4, QSER1 | 19772629 | Latourelle JC | 11:32874118 |
| rs7577851 | AAK1 | 19772629 | Latourelle JC | 2:69496578 |
| rs997277 | COL3A1, DIRC1 | 25663231 | Beecham GW | 2:188886137 |
| rs10788972 | TCEANC2 | 25663231 | Beecham GW | 1:54106570 |
| rs78736162 | MX2 | 25663231 | Beecham GW | 21:41368561 |
| rs6465122 | ZP3 | 25663231 | Beecham GW | 7:76411547 |
| rs12921479 | TRAPPC2L, PABPN1L | 25663231 | Beecham GW | 16:88862799 |
| rs141863958 | KCNIP4 | 25663231 | Beecham GW | 4:21291814 |
| rs816535 | MIR663AHG | 25663231 | Beecham GW | 20:26275360 |
| rs7984966 | HTR2A-AS1, HTR2A | 25663231 | Beecham GW | 13:46855311 |

**Table 4:** The list of genes used in spatial transcriptomics.

| N | Gene | N | Gene | N | Gene | N | Gene |
|---|---|---|---|---|---|---|---|
| 1 | SNAP25 | 25 | PDGFRA | 49 | PMEPA1 | 73 | VAV3 |
| 2 | SYT1 | 26 | DSCAM | 50 | GAD2 | 74 | DPP10 |
| 3 | SLC1A3 | 27 | OLIG1 | 51 | GAD1 | 75 | CD44 |
| 4 | SLC1A2 | 28 | CD2 | 52 | SEZ6 | 76 | SLC1A4 |
| 5 | AQP4 | 29 | THEMIS | 53 | GRIN2B | 77 | LAMA2 |
| 6 | PSD2 | 30 | SLC6A3 | 54 | CADPS2 | 78 | MARCKSL1 |
| 7 | SLC24A4 | 31 | TH | 55 | KLHL1 | 79 | TMEM163 |
| 8 | BCL2 | 32 | CKB | 56 | PRKG1 | 80 | P2RY12 |
| 9 | ITPKB | 33 | UCHL1 | 57 | PRKCA | 81 | CD163 |
| 10 | LRRK2 | 34 | ACTG1 | 58 | SGCD | 82 | C1QC |
| 11 | ITGAM | 35 | SCG2 | 59 | DAB1 | 83 | TRPM3 |
| 12 | DNAJC5 | 36 | ALDH1A1 | 60 | SOX6 | 84 | OPALIN |
| 13 | BHLHE41 | 37 | ATP5F1A | 61 | SDK1 | 85 | ACSBG1 |
| 14 | MOG | 38 | ATP13A2 | 62 | PDE4D | 86 | NFIB |
| 15 | ST18 | 39 | CBLN1 | 63 | GALNTL6 | 88 | CDH13 |
| 16 | COBL | 40 | PPP2R1A | 64 | UNC13C | 89 | CPED1 |
| 17 | PLCL1 | 41 | PGK1 | 65 | SCD | 90 | LGR6 |
| 18 | ELOVL7 | 42 | PINK1 | 66 | CTNNA3 | 91 | IP6K2 |
| 19 | FLT1 | 43 | VAT1 | 67 | SLC44A1 | 92 | BAG6 |

| | | | | | | | |
|---|---|---|---|---|---|---|---|
| 20 | PDGFRB | 44 | SNCA | 68 | GAB1 | 93 | SLC32A1 |
| 21 | COL12A1 | 45 | CLCN3 | 69 | ROBO1 | 94 | SYT2 |
| 22 | COL18A1 | 46 | VDAC1 | 70 | GABRA1 | 95 | OTX2 |
| 23 | CLDN5 | 47 | TIAM1 | 71 | GRIK1 | 96 | TPH1 |
| 24 | VCAN | 48 | GRID2 | 72 | LRRC4C | | |

**Table 5:** Top 30 markers of neuronal subtypes, states of astrocytes, states of microglia, states of oligodendrocytes.

| P-value (Wilcoxon Rank Sum test) | Ln fold change of the average expression | Adjusted p-value (Bonferroni Correction) | Cluster | Gene |
|---|---|---|---|---|
| 0 | 2.10063609927135 | 0 | neurons0 | ALDH1A1 |
| 0 | 1.60144327750264 | 0 | neurons0 | SLC18A2 |
| 0 | 1.57794687840324 | 0 | neurons0 | PEG10 |
| 0 | 1.47944361000439 | 0 | neurons0 | HSP90AA1 |
| 0 | 1.44837015036105 | 0 | neurons0 | TUBB2A |
| 0 | 1.38451879928164 | 0 | neurons0 | UCHL1 |
| 0 | 1.3774166629163 | 0 | neurons0 | SCG2 |
| 0 | 1.34139407347054 | 0 | neurons0 | TUBA1A |
| 0 | 1.33598988121485 | 0 | neurons0 | HSP90AB1 |
| 0 | 1.30183100825467 | 0 | neurons0 | SLC6A3 |
| 0 | 1.29657962697642 | 0 | neurons0 | ATP6V1B2 |
| 0 | 1.23020232006313 | 0 | neurons0 | TUBA1B |
| 0 | 1.22914232778746 | 0 | neurons0 | GAPDH |
| 0 | 1.22635068728498 | 0 | neurons0 | NEFL |
| 0 | 1.21584996972611 | 0 | neurons0 | TH |
| 0 | 1.21272269527338 | 0 | neurons0 | ACTG1 |
| 0 | 1.21189882315672 | 0 | neurons0 | RTN4 |
| 0 | 1.11437629344107 | 0 | neurons0 | SERINC1 |
| 0 | 1.09099814419483 | 0 | neurons0 | HSPA8 |
| 0 | 1.06112915238591 | 0 | neurons0 | UBB |
| 0 | 1.05565401111523 | 0 | neurons0 | PLD3 |
| 0 | 1.05248960136111 | 0 | neurons0 | SARAF |
| 0 | 1.04936675031283 | 0 | neurons0 | YWHAG |
| 0 | 1.03226952731248 | 0 | neurons0 | TERF2IP |
| 0 | 1.02047233645528 | 0 | neurons0 | PGRMC1 |
| 0 | 1.01620260345566 | 0 | neurons0 | CHGB |
| 0 | 1.01539301629168 | 0 | neurons0 | MAP1B |
| 0 | 1.01317407397453 | 0 | neurons0 | YWHAB |
| 0 | 1.01142475039306 | 0 | neurons0 | STMN2 |

| | | | | |
|---|---|---|---|---|
| 0 | 0.993457618936411 | 0 | neurons0 | ATP1B1 |
| 0 | 2.26026543523279 | 0 | neurons1 | GALNTL6 |
| 0 | 2.23757360877458 | 0 | neurons1 | PDE4D |
| 0 | 2.13133580970016 | 0 | neurons1 | PRKG1 |
| 0 | 2.02304530608943 | 0 | neurons1 | DAB1 |
| 0 | 1.99140631647865 | 0 | neurons1 | SGCD |
| 0 | 1.87448629273154 | 0 | neurons1 | IQCJ-SCHIP1 |
| 0 | 1.86035952050796 | 0 | neurons1 | CSMD1 |
| 0 | 1.832974873736 | 0 | neurons1 | PCDH15 |
| 0 | 1.7258492317283 | 0 | neurons1 | NEAT1 |
| 0 | 1.62831496671539 | 0 | neurons1 | AC073050.1 |
| 0 | 1.53542448850782 | 0 | neurons1 | SLC44A5 |
| 0 | 1.52830254162751 | 0 | neurons1 | RGS6 |
| 1.64795931348539e-302 | 1.61598079746433 | 1.0245363051923e-298 | neurons1 | DCC |
| 6.63894500728683e-292 | 1.47157175532948 | 4.12743211103022e-288 | neurons1 | ADARB2 |
| 6.95904689356131e-291 | 1.55119214234239 | 4.32643945372707e-287 | neurons1 | PDE4B |
| 1.02924393197418e-290 | 1.77831966381378 | 6.39880952508345e-287 | neurons1 | LRP1B |
| 3.07259125273642e-274 | 1.41095626467983 | 1.91022998182623e-270 | neurons1 | PLEKHA5 |
| 2.24875967120787e-256 | 1.54365367347044 | 1.39805388758993e-252 | neurons1 | MACROD2 |
| 1.68873298829065e-255 | 1.38534463873227 | 1.04988529882029e-251 | neurons1 | FAM155A |
| 5.94516422589769e-255 | 1.62299571176039 | 3.6961085992406e-251 | neurons1 | CNTNAP2 |
| 7.03501115993707e-250 | 1.42143193252878 | 4.37366643813287e-246 | neurons1 | FRMD4A |
| 1.45655759838074e-248 | 1.33895788339141 | 9.05541858913307e-245 | neurons1 | NRXN3 |
| 1.66729755971344e-245 | 1.6148919576547 | 1.03655889287385e-241 | neurons1 | ERBB4 |
| 2.14390065508919e-243 | 1.35352201615133 | 1.33286303726895e-239 | neurons1 | AL589740.1 |
| 5.97027264793728e-242 | 1.58769809062867 | 3.7117185052226e-238 | neurons1 | FGF12 |
| 2.0745802820256e-239 | 1.7877842209946 | 1.28976656133532e-235 | neurons1 | KLHL1 |
| 2.02227130535735e-234 | 1.53452182634097 | 1.25724607054067e-230 | neurons1 | ROBO2 |
| 5.38396939185751e-227 | 1.43061475079612 | 3.34721377091781e-223 | neurons1 | RALYL |
| 6.35333003953782e-226 | 1.50025504712092 | 3.94986528558066e-222 | neurons1 | KCNQ3 |
| 3.66465583442554e-198 | 1.43306249072775 | 2.27831653226236e-194 | neurons1 | SEMA6D |
| 0 | 2.32876902733058 | 0 | neurons2 | RBFOX1 |
| 0 | 1.51912577791838 | 0 | neurons2 | FGF14 |
| 0 | 1.38698882989493 | 0 | neurons2 | RYR2 |
| 0 | 1.27928140939911 | 0 | neurons2 | PCDH11X |
| 1.84108711848972e-288 | 1.84620351846826 | 1.14460386156506e-284 | neurons2 | GRID2 |
| 4.21985226983994e-284 | 1.47800045519075 | 2.62348215615949e-280 | neurons2 | CDH18 |
| 2.88153998244763e-270 | 1.38115196742105 | 1.79145340708769e-266 | neurons2 | KIAA1217 |
| 6.41710925202459e-264 | 1.19644594207074 | 3.98951682198369e-260 | neurons2 | HTR2C |
| 1.68582439115773e-262 | 1.44185867699179 | 1.04807702398276e-258 | neurons2 | OPCML |
| 2.18099489386258e-252 | 1.22192170639522 | 1.35592452551437e-248 | neurons2 | GRM7 |

| | | | | |
|---|---|---|---|---|
| 3.09414429476839e-249 | 1.46507583172158 | 1.92362950805751e-245 | neurons2 | GRIK2 |
| 2.15287357608456e-235 | 1.66449391931306 | 1.33844150225177e-231 | neurons2 | ROBO1 |
| 8.11830259533803e-226 | 1.29519998108133 | 5.04714872352165e-222 | neurons2 | HS3ST4 |
| 1.3992469138617e-221 | 1.42691660004632 | 8.69911806347819e-218 | neurons2 | SGCZ |
| 1.99313139756788e-219 | 1.13829979794793 | 1.23912978986795e-215 | neurons2 | NAV3 |
| 6.78876196777036e-206 | 1.08710334281746 | 4.22057331536283e-202 | neurons2 | AGBL4 |
| 3.9846112229798e-201 | 1.22401388867605 | 2.47723279732654e-197 | neurons2 | LHFPL3 |
| 5.75198282305991e-190 | 1.41705455748908 | 3.57600772109634e-186 | neurons2 | NRG3 |
| 1.75207279234999e-189 | 1.35964198350581 | 1.08926365500399e-185 | neurons2 | TENM2 |
| 9.10140034353962e-186 | 1.06147892741632 | 5.65834059357858e-182 | neurons2 | STXBP5L |
| 5.0250079579868e-182 | 1.43607895544552 | 3.12404744748039e-178 | neurons2 | PTPRD |
| 1.77908187349488e-175 | 1.29276083992591 | 1.10605520075177e-171 | neurons2 | TRPM3 |
| 3.28866483688427e-175 | 1.13680374828962 | 2.04456292909095e-171 | neurons2 | GRIP1 |
| 6.60701756497971e-155 | 1.08219899347394 | 4.10758282014789e-151 | neurons2 | NCAM2 |
| 3.11945024733657e-153 | 1.1648385739194 | 1.93936221876914e-149 | neurons2 | CNTN5 |
| 4.10393806099349e-153 | 1.12865542925183 | 2.55141829251965e-149 | neurons2 | KAZN |
| 5.53025687074696e-148 | 1.23918737122834 | 3.43816069654339e-144 | neurons2 | NEGR1 |
| 2.8212703985172e-144 | 1.04317127346159 | 1.75398380675814e-140 | neurons2 | DLG2 |
| 4.00005630661181e-140 | 1.2145285812257 | 2.48683500582056e-136 | neurons2 | NRG1 |
| 4.07744904104815e-94 | 1.15746128076849 | 2.53495006881964e-90 | neurons2 | DPP10 |
| 0 | 1.42461343949247 | 0 | neurons3 | GAD1 |
| 0 | 1.35812118771256 | 0 | neurons3 | GAD2 |
| 2.91805572228008e-267 | 2.05602778317478 | 1.81415524254153e-263 | neurons3 | GABRA1 |
| 2.34748322103399e-209 | 1.5463300485525 | 1.45943031851683e-205 | neurons3 | GABRB2 |
| 5.72080528615052e-183 | 1.06335458334122 | 3.55662464639978e-179 | neurons3 | SCN1A |
| 5.99347025460838e-179 | 0.942243604780411 | 3.72614045729003e-175 | neurons3 | CALB2 |
| 1.09599061379104e-141 | 0.85069115294771 | 6.81377364593891e-138 | neurons3 | PCP4 |
| 6.60185285133834e-141 | 0.741602656147952 | 4.10437191767704e-137 | neurons3 | KCNC1 |
| 5.71127505347971e-89 | 0.577896600204892 | 3.55069970074833e-85 | neurons3 | HTR2C |
| 1.40887342222317e-86 | 1.14874993145978 | 8.75896606596145e-83 | neurons3 | SPARCL1 |
| 1.36870262792385e-85 | 0.818140438582976 | 8.50922423780256e-82 | neurons3 | SLC24A2 |
| 1.43631248903225e-80 | 0.877592972621254 | 8.9295547443135e-77 | neurons3 | VGF |
| 3.91754494972104e-80 | 0.753339935022328 | 2.43553769524157e-76 | neurons3 | GNAS |
| 4.35443958407382e-72 | 0.755645315565967 | 2.70715508941869e-68 | neurons3 | GABRG2 |
| 2.44665947229797e-69 | 0.698718225609401 | 1.52108819392765e-65 | neurons3 | CHGA |
| 2.49761645813101e-65 | 0.743877392385975 | 1.55276815202005e-61 | neurons3 | OXR1 |
| 2.89933519458964e-65 | 0.696613757921842 | 1.80251669047638e-61 | neurons3 | ATP1B1 |
| 1.1442707153134e-64 | 0.926812767877073 | 7.11393103710341e-61 | neurons3 | NEFM |
| 1.4295795054254e-63 | 0.695048896592796 | 8.88769578522974e-60 | neurons3 | CKB |
| 2.96067357476482e-63 | 0.668847402669786 | 1.84065076143129e-59 | neurons3 | SLC12A5 |
| 8.73071810393379e-61 | 0.66274913953167 | 5.42788744521563e-57 | neurons3 | SLC22A17 |

| | | | | |
|---|---|---|---|---|
| 7.16663609193379e-59 | 0.61721248180056 | 4.45549765835524e-55 | neurons3 | SORL1 |
| 3.70539213553163e-58 | 0.733082777479108 | 2.30364229066002e-54 | neurons3 | INA |
| 5.90731715588274e-55 | 0.850142882787111 | 3.6725790758123e-51 | neurons3 | SPP1 |
| 3.78933676955289e-53 | 0.610886971165792 | 2.35583066963103e-49 | neurons3 | SV2A |
| 5.45891689938521e-52 | 0.632481949985293 | 3.39380863634779e-48 | neurons3 | SEZ6L2 |
| 1.60827166547003e-49 | 0.604572597002252 | 9.99862494422718e-46 | neurons3 | CLU |
| 1.08280156229739e-44 | 0.587779758848618 | 6.73177731280284e-41 | neurons3 | DNM1 |
| 4.91764499878345e-43 | 0.77212449976066 | 3.05729989574367e-39 | neurons3 | NEFH |
| 1.62457879398385e-29 | 0.597362808162695 | 1.01000063621976e-25 | neurons3 | SLC6A1 |
| 3.46261918333643e-181 | 1.15413686197213 | 2.15271034628026e-177 | neurons4 | ZEB2 |
| 1.29595455289171e-153 | 1.65386808847459 | 8.05694945532775e-150 | neurons4 | ST18 |
| 5.45457223602582e-140 | 1.36307950966105 | 3.39110755913725e-136 | neurons4 | NCKAP5 |
| 2.7019512600947e-130 | 1.15127792352704 | 1.67980309840088e-126 | neurons4 | TMEM144 |
| 8.10457246580761e-116 | 0.978295944789811 | 5.03861270199259e-112 | neurons4 | GAB1 |
| 2.58329143262164e-115 | 1.61342301846766 | 1.60603228366088e-111 | neurons4 | CTNNA3 |
| 1.0133134993919e-106 | 1.23048542124529 | 6.29977002571943e-103 | neurons4 | MAN2A1 |
| 3.20048631960231e-95 | 1.57279831234984 | 1.98974234489676e-91 | neurons4 | QKI |
| 5.95670221310257e-95 | 1.09750743269127 | 3.70328176588587e-91 | neurons4 | CDH20 |
| 7.53418733114385e-95 | 1.27271786620703 | 4.68400426377213e-91 | neurons4 | PIP4K2A |
| 3.83755650882977e-94 | 1.1087897017111 | 2.38580888153947e-90 | neurons4 | PHLPP1 |
| 8.13665116888126e-89 | 1.51654813795091 | 5.05855603169348e-85 | neurons4 | SLC44A1 |
| 3.20977838933928e-79 | 1.20432335654151 | 1.99551922465223e-75 | neurons4 | FMNL2 |
| 8.03572124810935e-77 | 1.22342829853564 | 4.99580789994958e-73 | neurons4 | PLCL1 |
| 9.71289911956621e-76 | 1.09284898835215 | 6.03850938263432e-72 | neurons4 | ERBIN |
| 1.15129882858448e-71 | 1.68229279358586 | 7.15762481730969e-68 | neurons4 | PCDH9 |
| 1.79244893086772e-71 | 1.04302564546601 | 1.11436550032046e-67 | neurons4 | MAP7 |
| 3.61304887109e-71 | 0.976201801372753 | 2.24623248315665e-67 | neurons4 | TF |
| 4.67918332830259e-69 | 1.2887158570258 | 2.90904827520572e-65 | neurons4 | RNF220 |
| 9.87185003187467e-66 | 1.13141889156309 | 6.13732916481648e-62 | neurons4 | ZBTB20 |
| 1.10459893894999e-64 | 1.35598256978423 | 6.86729160345208e-61 | neurons4 | MBP |
| 1.70658357950678e-63 | 1.10347708578563 | 1.06098301137937e-59 | neurons4 | TMTC2 |
| 2.68609664699896e-59 | 1.15132015953583 | 1.66994628543925e-55 | neurons4 | GPM6B |
| 2.3357999822074e-55 | 1.06269481577251 | 1.45216684893834e-51 | neurons4 | PDE4B |
| 3.89201468921598e-55 | 1.00969346671909 | 2.41966553228558e-51 | neurons4 | WWOX |
| 7.12737804741638e-55 | 1.11797858253422 | 4.43109093207876e-51 | neurons4 | ELMO1 |
| 1.95250952243219e-52 | 1.01907718089255 | 1.2138751700961e-48 | neurons4 | NPAS3 |
| 1.11234542462499e-51 | 1.05495791564626 | 6.91545150489355e-48 | neurons4 | IL1RAPL1 |
| 4.1557361646918e-46 | 1.41860341910475 | 2.58362117358889e-42 | neurons4 | PLP1 |
| 2.62481508665149e-40 | 1.05969941978134 | 1.63184753937123e-36 | neurons4 | PPP2R2B |
| 0 | 3.14207423334686 | 0 | neurons5 | GRIK1 |
| 1.45500548055413e-294 | 1.49081149797796 | 9.04576907260505e-291 | neurons5 | PDZRN3 |

| | | | | |
|---|---|---|---|---|
| 1.96901370649845e-251 | 1.81552886769453 | 1.22413582133009e-247 | neurons5 | SYN3 |
| 2.84110029632872e-246 | 1.65851261210126 | 1.76631205422757e-242 | neurons5 | ZFPM2 |
| 5.48391495757311e-233 | 1.7780184116792 | 3.4093499291232e-229 | neurons5 | SYNPR |
| 1.83315382446706e-226 | 1.56719445356647 | 1.13967173267117e-222 | neurons5 | UNC5D |
| 4.60959664812414e-211 | 1.42806512231473 | 2.86578623613878e-207 | neurons5 | CDH13 |
| 2.96731327894203e-198 | 1.53676975846182 | 1.84477866551826e-194 | neurons5 | GALNT13 |
| 1.46819112876634e-186 | 1.38820602507148 | 9.12774424754035e-183 | neurons5 | KCTD8 |
| 6.01871817819884e-172 | 1.37435355327324 | 3.74183709138622e-168 | neurons5 | TMEM108 |
| 2.40850969655412e-170 | 1.40805967311009 | 1.4973704783477e-166 | neurons5 | SYN2 |
| 6.59470620357939e-169 | 1.70416454816429 | 4.0999288467653e-165 | neurons5 | OPCML |
| 2.33740207383915e-161 | 1.49685959167321 | 1.4531628693058e-157 | neurons5 | SGCZ |
| 1.34391318326197e-159 | 1.51782544291698 | 8.35510826033969e-156 | neurons5 | CACNA2D3 |
| 1.74072492615914e-150 | 2.25426356899565 | 1.08220868659314e-146 | neurons5 | DLGAP1 |
| 8.49135081929934e-148 | 1.44453600308526 | 5.2790728043584e-144 | neurons5 | FAT3 |
| 1.03080093437452e-146 | 1.38153378086056 | 6.40848940900636e-143 | neurons5 | CACNA1C |
| 1.32286310299391e-140 | 1.70208004114242 | 8.22423991131314e-137 | neurons5 | FGF14 |
| 4.31571838076487e-138 | 1.71102020879048 | 2.68308211732152e-134 | neurons5 | MDGA2 |
| 1.68250033105532e-134 | 1.4651240786243 | 1.04601045581709e-130 | neurons5 | CNTNAP5 |
| 5.02705834170876e-126 | 1.8166651043687 | 3.12532217104033e-122 | neurons5 | AUTS2 |
| 1.45113377677852e-125 | 1.98225288212767 | 9.02169869023207e-122 | neurons5 | KCND2 |
| 6.76136084901503e-111 | 1.49324388256804 | 4.20353803983264e-107 | neurons5 | DSCAM |
| 8.86043028422437e-111 | 1.52979539604377 | 5.50852950770229e-107 | neurons5 | RORA |
| 2.51148301887076e-108 | 2.55573141276963 | 1.56138899283195e-104 | neurons5 | KCNIP4 |
| 9.38940060536467e-107 | 1.46132121283393 | 5.83739035635521e-103 | neurons5 | LRRC4C |
| 5.95024257847732e-104 | 1.50637097576337 | 3.69926581103935e-100 | neurons5 | NRG3 |
| 1.22344582315042e-100 | 1.43052935642731 | 7.60616268252614e-97 | neurons5 | PLCB1 |
| 1.34988497051206e-91 | 1.44893208394242 | 8.39223486167346e-88 | neurons5 | DLG2 |
| 6.14094040759125e-84 | 1.38365062423787 | 3.81782265139948e-80 | neurons5 | LSAMP |
| 0 | 0.380988674667607 | 0 | astrocytes0 | GPC5 |
| 0 | 0.348751205649349 | 0 | astrocytes0 | NRXN1 |
| 3.92593144994093e-250 | 0.271266341456658 | 2.96721898986536e-246 | astrocytes0 | ADGRB3 |
| 2.40050323711436e-231 | 0.258561906371582 | 1.81430034661103e-227 | astrocytes0 | CADM2 |
| 1.63921694849169e-208 | 0.343808839473779 | 1.23892016967002e-204 | astrocytes0 | KCND2 |
| 3.63698656631333e-199 | 0.324775799471213 | 2.74883444681962e-195 | astrocytes0 | SLC6A11 |
| 1.1090399532462e-197 | 0.253049832332989 | 8.38212396663482e-194 | astrocytes0 | GPM6A |
| 8.87619926539914e-192 | 0.339539294523512 | 6.70863140478867e-188 | astrocytes0 | LRRC4C |
| 7.76276450014668e-191 | 0.612121771792742 | 5.86709740921086e-187 | astrocytes0 | XIST |
| 4.26154517467158e-172 | 0.322423386908026 | 3.22087584301678e-168 | astrocytes0 | FAM155A |
| 2.03842370329306e-151 | 0.333725896082354 | 1.54064063494889e-147 | astrocytes0 | SLC1A2 |
| 6.7674616393802e-146 | 0.299418177032957 | 5.11484750704356e-142 | astrocytes0 | CABLES1 |
| 5.21030107944326e-144 | 0.275744400175518 | 3.93794555584321e-140 | astrocytes0 | RGS7 |

| | | | | |
|---|---|---|---|---|
| 4.12325535682796e-137 | 0.535370881495168 | 3.11635639869057e-133 | astrocytes0 | LINC00499 |
| 3.50634997670215e-136 | 0.276328008075393 | 2.65009931239149e-132 | astrocytes0 | RGS6 |
| 4.20428205117228e-120 | 0.265148368944663 | 3.17759637427601e-116 | astrocytes0 | CACNB2 |
| 1.13224269231107e-110 | 0.267278971940772 | 8.55749026848705e-107 | astrocytes0 | GUCY1A2 |
| 1.69623610861772e-101 | 0.259030667707301 | 1.28201525089327e-97 | astrocytes0 | PTCH1 |
| 9.73684543767895e-84 | 0.354633696912948 | 7.35910778179775e-80 | astrocytes0 | VAV3 |
| 2.94321308604584e-77 | 0.390579510702723 | 2.22448045043344e-73 | astrocytes0 | GRM5 |
| 3.22061466535846e-77 | 0.260416900434995 | 2.43414056407792e-73 | astrocytes0 | TMEM108 |
| 7.94108968618914e-71 | 0.326297401584594 | 6.00187558482175e-67 | astrocytes0 | AC012593.1 |
| 4.13546110316199e-70 | 0.34190593506946 | 3.12558150176983e-66 | astrocytes0 | SGCZ |
| 9.39579626041345e-63 | 0.3311235496172 | 7.10134281362048e-59 | astrocytes0 | RIMS1 |
| 6.42251293109106e-59 | 0.250342803969962 | 4.85413527331863e-55 | astrocytes0 | RALYL |
| 2.51136673698412e-54 | 0.287680749228085 | 1.8980909798126e-50 | astrocytes0 | CCDC129 |
| 1.41733104074867e-43 | 0.270967421200149 | 1.07121880059784e-39 | astrocytes0 | ZNF98 |
| 0 | 2.3199406575042 | 0 | astrocytes1 | DPP10 |
| 0 | 0.968116655742867 | 0 | astrocytes1 | ADAMTSL3 |
| 0 | 0.915861446132331 | 0 | astrocytes1 | STXBP5L |
| 0 | 0.88553750563614 | 0 | astrocytes1 | C8orf34 |
| 0 | 0.875121970644028 | 0 | astrocytes1 | CPAMD8 |
| 0 | 0.861741399480078 | 0 | astrocytes1 | DCLK1 |
| 0 | 0.82616976255629 | 0 | astrocytes1 | CCDC85A |
| 0 | 0.698805560624623 | 0 | astrocytes1 | CD44 |
| 0 | 0.656602345727454 | 0 | astrocytes1 | GALNT15 |
| 0 | 0.654650982113482 | 0 | astrocytes1 | MAN1C1 |
| 0 | 0.637620465402837 | 0 | astrocytes1 | LINC00609 |
| 0 | 0.604373978611923 | 0 | astrocytes1 | PRKG1 |
| 0 | 0.561358761912141 | 0 | astrocytes1 | ABI3BP |
| 0 | 0.531254406695393 | 0 | astrocytes1 | SLC38A1 |
| 0 | 0.499233961178096 | 0 | astrocytes1 | CERS6 |
| 0 | 0.429521317969376 | 0 | astrocytes1 | LRAT |
| 1.40287238447167e-285 | 0.458784957698346 | 1.06029094818369e-281 | astrocytes1 | NFASC |
| 1.71980967571319e-278 | 0.690833547752815 | 1.29983215290403e-274 | astrocytes1 | MIR4300HG |
| 2.68826594041679e-260 | 0.589305380324721 | 2.03179139776701e-256 | astrocytes1 | TTN |
| 6.25234329519639e-237 | 0.470700101689373 | 4.72552106250943e-233 | astrocytes1 | ST6GAL1 |
| 2.20906379425703e-220 | 0.457475204791557 | 1.66961041569947e-216 | astrocytes1 | COL21A1 |
| 5.52424000202281e-207 | 0.500691473936896 | 4.17522059352884e-203 | astrocytes1 | HS6ST3 |
| 3.3630298011393e-196 | 0.514336970545819 | 2.54177792370109e-192 | astrocytes1 | KAZN |
| 3.73991617786544e-196 | 0.467148561700296 | 2.8266286472307e-192 | astrocytes1 | CALN1 |
| 4.97718791324144e-193 | 0.428628617946573 | 3.76175862482788e-189 | astrocytes1 | ARHGEF3 |
| 1.08465130809787e-151 | 0.522019013307014 | 8.19779458660368e-148 | astrocytes1 | MAOB |
| 5.82794783910644e-108 | 0.421828917233308 | 4.40476297679665e-104 | astrocytes1 | CADPS |

| | | | | |
|---|---|---|---|---|
| 2.31696650386618e-91 | 0.440842827048896 | 1.75116328362206e-87 | astrocytes1 | LINC01094 |
| 1.14271683682318e-77 | 0.616324723100614 | 8.63665385270957e-74 | astrocytes1 | ROBO2 |
| 8.45298023003257e-06 | 0.455493392088262 | 0.0638876245785861 | astrocytes1 | NRXN3 |
| 0 | 0.899904335578988 | 0 | astrocytes2 | SYT1 |
| 0 | 0.806861983559021 | 0 | astrocytes2 | UCHL1 |
| 0 | 0.742841213613937 | 0 | astrocytes2 | ID2 |
| 0 | 0.627297433482444 | 0 | astrocytes2 | PEG10 |
| 0 | 0.618365189191557 | 0 | astrocytes2 | ATP1B1 |
| 0 | 0.589373007922373 | 0 | astrocytes2 | TUBA1B |
| 0 | 0.584647275762757 | 0 | astrocytes2 | UBC |
| 0 | 0.58123514254972 | 0 | astrocytes2 | GAPDH |
| 0 | 0.553851728375942 | 0 | astrocytes2 | TUBB2A |
| 0 | 0.538632930345017 | 0 | astrocytes2 | STMN2 |
| 0 | 0.538492148125031 | 0 | astrocytes2 | UBB |
| 0 | 0.5253633759703 | 0 | astrocytes2 | FRMD4A |
| 0 | 0.516285057131522 | 0 | astrocytes2 | SNAP25 |
| 0 | 0.498221950862433 | 0 | astrocytes2 | NEFL |
| 0 | 0.495352680281847 | 0 | astrocytes2 | ACTG1 |
| 0 | 0.486575089116689 | 0 | astrocytes2 | C1orf61 |
| 0 | 0.458141017311943 | 0 | astrocytes2 | SLC18A2 |
| 0 | 0.457929586972579 | 0 | astrocytes2 | NEFM |
| 0 | 0.457325511101989 | 0 | astrocytes2 | TUBA1A |
| 0 | 0.443580273518333 | 0 | astrocytes2 | JUNB |
| 0 | 0.434176396079583 | 0 | astrocytes2 | SCG2 |
| 2.22679859624712e-307 | 0.495934700641848 | 1.68301437904357e-303 | astrocytes2 | HSP90AA1 |
| 3.13459951788259e-291 | 0.50820502024872 | 2.36913031561566e-287 | astrocytes2 | HSP90AB1 |
| 5.47126447453615e-291 | 0.472452378896464 | 4.13518168985443e-287 | astrocytes2 | JUN |
| 3.88360283989055e-265 | 0.494192511150853 | 2.93522702638928e-261 | astrocytes2 | MAP1B |
| 1.55774019173062e-237 | 0.518982339268283 | 1.17734003691e-233 | astrocytes2 | HES1 |
| 3.76566288862468e-222 | 0.59396735229554 | 2.84608801122253e-218 | astrocytes2 | TIMP3 |
| 2.17032948382811e-216 | 0.533141418124452 | 1.64033502387729e-212 | astrocytes2 | AC009975.1 |
| 1.79137641177093e-208 | 0.446560788118238 | 1.35392229201647e-204 | astrocytes2 | ZFP36L2 |
| 2.34399520376852e-205 | 0.443572858594325 | 1.77159157500824e-201 | astrocytes2 | MMD2 |
| 0 | 0.773794054687926 | 0 | astrocytes3 | NEAT1 |
| 0 | 0.738860082228327 | 0 | astrocytes3 | RANBP3L |
| 0 | 0.557479488505007 | 0 | astrocytes3 | SLC39A11 |
| 0 | 0.46879981818323 | 0 | astrocytes3 | AC016831.7 |
| 0 | 0.42105698397376 | 0 | astrocytes3 | PARD3 |
| 1.57491875172592e-296 | 0.4635752495691 | 1.19032359255445e-292 | astrocytes3 | AL355974.2 |
| 9.38539680179327e-294 | 0.371296816995558 | 7.09348290279536e-290 | astrocytes3 | MRPS6 |
| 5.46421404487506e-291 | 0.455189319768446 | 4.12985297511657e-287 | astrocytes3 | FKBP5 |

| | | | | |
|---|---|---|---|---|
| 2.56198730387146e-258 | 0.519141910508683 | 1.93635000426605e-254 | astrocytes3 | TPD52L1 |
| 7.19498411815436e-233 | 0.385101794087768 | 5.43796899650107e-229 | astrocytes3 | LHFPL6 |
| 1.5607393342774e-219 | 0.467891862192126 | 1.17960678884686e-215 | astrocytes3 | OGFRL1 |
| 2.93554768532065e-218 | 0.388733043868121 | 2.21868694056535e-214 | astrocytes3 | HSPH1 |
| 5.59420379061431e-211 | 0.418799371804219 | 4.2280992249463e-207 | astrocytes3 | IFRD1 |
| 5.12363432440341e-192 | 0.33478561844148 | 3.8724428223841e-188 | astrocytes3 | AC092691.1 |
| 9.28212290493471e-184 | 0.430217733919659 | 7.01542849154967e-180 | astrocytes3 | SLC7A11 |
| 1.25211883558218e-174 | 0.337427942776556 | 9.46351415933008e-171 | astrocytes3 | TTTY14 |
| 3.79473846739292e-160 | 0.340230969755419 | 2.86806333365557e-156 | astrocytes3 | MAPK4 |
| 2.9518506091515e-156 | 0.344410811105553 | 2.2310086903967e-152 | astrocytes3 | BOC |
| 1.07992723929296e-154 | 0.349109375032278 | 8.16209007457617e-151 | astrocytes3 | BAG3 |
| 5.98581949701172e-151 | 0.349560823930651 | 4.52408237584146e-147 | astrocytes3 | PTGES3 |
| 5.31326867691778e-145 | 0.324418042828375 | 4.01576846601446e-141 | astrocytes3 | SESN1 |
| 1.71040199523348e-143 | 0.328361253829471 | 1.29272182799746e-139 | astrocytes3 | SYBU |
| 4.05618204901756e-143 | 0.550677702601665 | 3.06566239264747e-139 | astrocytes3 | CHI3L1 |
| 4.73904289077599e-143 | 0.338349283493608 | 3.5817686168485e-139 | astrocytes3 | SAMD4A |
| 1.54740280534227e-133 | 0.349527260143272 | 1.16952704027768e-129 | astrocytes3 | CHST11 |
| 9.08408160510132e-106 | 0.357671820309176 | 6.86574887713558e-102 | astrocytes3 | CACNA2D3 |
| 2.31265399456652e-81 | 0.368081276982881 | 1.74790388909338e-77 | astrocytes3 | ZNF98 |
| 3.87630590090989e-70 | 0.357453978203457 | 2.92971199990769e-66 | astrocytes3 | HMGB1 |
| 6.76283590606626e-62 | 0.376096636567075 | 5.11135137780488e-58 | astrocytes3 | HSPB1 |
| 7.55346902852928e-30 | 0.334543780281671 | 5.70891189176243e-26 | astrocytes3 | LAMA2 |
| 8.85755211059754e-110 | 0.346497248793733 | 6.69453788518962e-106 | astrocytes4 | CHI3L1 |
| 1.59087038198262e-100 | 0.450490441689186 | 1.20237983470246e-96 | astrocytes4 | CLU |
| 1.37932720097997e-96 | 0.451316904177429 | 1.04249549850066e-92 | astrocytes4 | MT2A |
| 1.0999961954746e-59 | 0.275566848747627 | 8.31377124539705e-56 | astrocytes4 | RASGEF1B |
| 3.77173664619922e-59 | 0.360072974925825 | 2.85067855719737e-55 | astrocytes4 | APOE |
| 6.18089852628764e-53 | 0.275275238978834 | 4.6715231061682e-49 | astrocytes4 | NMT1 |
| 1.29763011712598e-50 | 0.261697800104332 | 9.80748842523813e-47 | astrocytes4 | SRRM2 |
| 1.63202140283388e-48 | 0.287848577210354 | 1.23348177626184e-44 | astrocytes4 | ANGPTL4 |
| 1.28730207496123e-44 | 0.255550894205184 | 9.72942908255696e-41 | astrocytes4 | GFAP |
| 4.55983563130862e-34 | 0.295320416681688 | 3.44632377014305e-30 | astrocytes4 | MT1E |
| 1.26830481287683e-29 | 0.295350291952434 | 9.58584777572311e-26 | astrocytes4 | CST3 |
| 3.59139581602241e-19 | 0.319081550329269 | 2.71437695774974e-15 | astrocytes4 | PLCG2 |
| 1.61197325018716e-16 | 0.38429592606328 | 1.21832938249146e-12 | astrocytes4 | LINGO1 |
| 4.39308599073566e-10 | 0.274990615216682 | 3.32029439179801e-06 | astrocytes4 | MT3 |
| 4.28740643098257e-08 | 0.30626173817114 | 0.000324042178053663 | astrocytes4 | SLC26A3 |
| 0 | 2.55509301089466 | 0 | astrocytes5 | IL1RAPL1 |
| 0 | 2.23539620015456 | 0 | astrocytes5 | ST18 |
| 0 | 2.02614865667253 | 0 | astrocytes5 | SLC24A2 |
| 0 | 2.01549231190751 | 0 | astrocytes5 | ELMO1 |

| | | | | |
|---|---|---|---|---|
| 0 | 2.013517669834 | 0 | astrocytes5 | CTNNA3 |
| 0 | 1.97055117348543 | 0 | astrocytes5 | MBP |
| 0 | 1.96065399819442 | 0 | astrocytes5 | EDIL3 |
| 0 | 1.91189221008597 | 0 | astrocytes5 | PEX5L |
| 0 | 1.64974591350324 | 0 | astrocytes5 | SPOCK3 |
| 0 | 1.58991506485247 | 0 | astrocytes5 | TTLL7 |
| 0 | 1.55663913624623 | 0 | astrocytes5 | ANK3 |
| 0 | 1.53341702916794 | 0 | astrocytes5 | PRUNE2 |
| 0 | 1.44949892854479 | 0 | astrocytes5 | FRMD4B |
| 0 | 1.38421950790868 | 0 | astrocytes5 | MARCH1 |
| 0 | 1.31874329573684 | 0 | astrocytes5 | SHTN1 |
| 0 | 1.28116073790116 | 0 | astrocytes5 | CLMN |
| 1.67600088901714e-306 | 1.74355449195932 | 1.26672147191915e-302 | astrocytes5 | PLCL1 |
| 2.84068153604174e-301 | 1.40095959832005 | 2.14698710820004e-297 | astrocytes5 | NLGN1 |
| 4.44025347061776e-273 | 1.7446960854122 | 3.3559435730929e-269 | astrocytes5 | RNF220 |
| 2.29195598677019e-266 | 1.30396396189062 | 1.73226033480091e-262 | astrocytes5 | UNC5C |
| 3.06023497830482e-239 | 1.55559268181398 | 2.31292559660278e-235 | astrocytes5 | PTPRD |
| 1.39515043502229e-238 | 1.36334743354175 | 1.05445469878984e-234 | astrocytes5 | DNM3 |
| 2.9757062941082e-234 | 1.33494211208821 | 2.24903881708698e-230 | astrocytes5 | QDPR |
| 4.52145384855151e-216 | 1.60288139531514 | 3.41731481873523e-212 | astrocytes5 | MAN2A1 |
| 2.91323412633862e-215 | 1.5168084000605 | 2.20182235268673e-211 | astrocytes5 | PIP4K2A |
| 4.70397257135086e-208 | 2.20831049240501 | 3.55526246942698e-204 | astrocytes5 | PLP1 |
| 2.15259917358041e-173 | 1.28707324891934 | 1.62693445539207e-169 | astrocytes5 | SLC44A1 |
| 8.32953217592907e-172 | 1.18683023701333 | 6.29546041856719e-168 | astrocytes5 | FRYL |
| 2.42986596419671e-171 | 1.45947330972408 | 1.83649269573987e-167 | astrocytes5 | CNTNAP2 |
| 2.40371725628783e-159 | 1.20464455232418 | 1.81672950230234e-155 | astrocytes5 | SPP1 |
| 2.65134492612137e-176 | 0.2574796389643 | 1.69977723213641e-172 | microglia0 | APOE |
| 6.16840330106818e-127 | 0.345305698114125 | 3.95456335631481e-123 | microglia0 | SPP1 |
| 2.32921156972893e-94 | 0.263855652638987 | 1.49325753735322e-90 | microglia0 | FCGR3A |
| 2.02595172593335e-75 | 0.258309172064581 | 1.29883765149587e-71 | microglia0 | VSIG4 |
| 1.217114831806e-67 | 0.255593305579039 | 7.80292318670829e-64 | microglia0 | NCK2 |
| 4.61607641830674e-22 | 0.352543592170951 | 2.95936659177645e-18 | microglia0 | TMEM163 |
| 0 | 0.95922193867867 | 0 | microglia1 | SYT1 |
| 0 | 0.94695574890768 | 0 | microglia1 | MAP1B |
| 0 | 0.855095191424653 | 0 | microglia1 | ALDH1A1 |
| 0 | 0.79982089250456 | 0 | microglia1 | UCHL1 |
| 0 | 0.63187911376326 | 0 | microglia1 | PEG10 |
| 0 | 0.5989551251282 | 0 | microglia1 | SNAP25 |
| 0 | 0.505401108466269 | 0 | microglia1 | STMN2 |
| 0 | 0.499815241692593 | 0 | microglia1 | TUBB2A |
| 3.27144759335858e-307 | 0.49485880019924 | 2.09732505210219e-303 | microglia1 | SLC18A2 |

| | | | | |
|---|---|---|---|---|
| 6.35438103673538e-297 | 0.624295487465996 | 4.07379368265126e-293 | microglia1 | ATP1B1 |
| 3.51030565393252e-284 | 0.505312749878735 | 2.25045695473614e-280 | microglia1 | NEFM |
| 3.91976288472e-267 | 0.502342639814799 | 2.51295998539399e-263 | microglia1 | NEFL |
| 2.45038659108309e-233 | 0.45474599555428 | 1.57094284354337e-229 | microglia1 | SCG2 |
| 8.64724760379737e-229 | 0.463960021705735 | 5.54375043879449e-225 | microglia1 | TUBA1A |
| 2.91627973238987e-196 | 0.411638889946608 | 1.86962693643514e-192 | microglia1 | PEBP1 |
| 3.08556808039281e-191 | 0.511043550539774 | 1.97815769633983e-187 | microglia1 | RASGEF1C |
| 7.08364949729293e-189 | 0.431130018903734 | 4.5413276927145e-185 | microglia1 | AP003481.1 |
| 1.15580726011401e-183 | 0.660731045648907 | 7.40988034459095e-180 | microglia1 | CCDC26 |
| 3.80907199104908e-179 | 0.550972786841928 | 2.44199605346157e-175 | microglia1 | HSP90AA1 |
| 1.34828472365079e-174 | 0.432876411723993 | 8.6438533633252e-171 | microglia1 | SYNDIG1 |
| 5.63697863044856e-149 | 0.48076683034522 | 3.61386699998057e-145 | microglia1 | OLR1 |
| 4.04178179540347e-147 | 0.409697114843534 | 2.59118630903316e-143 | microglia1 | ZFP36L1 |
| 5.36084565880777e-142 | 0.401337100220235 | 3.43683815186166e-138 | microglia1 | FOS |
| 1.06135528686984e-114 | 0.419164842061733 | 6.80434874412254e-111 | microglia1 | CX3CR1 |
| 5.47651837616115e-94 | 0.547819325706227 | 3.51099593095691e-90 | microglia1 | SLC2A3 |
| 1.74545234763208e-85 | 0.438711800885847 | 1.11900950006692e-81 | microglia1 | XACT |
| 8.81287423767925e-56 | 0.510619287894374 | 5.64993367377617e-52 | microglia1 | HIST1H2AC |
| 3.94655731289862e-48 | 0.435898731551206 | 2.5301378932993e-44 | microglia1 | SRGN |
| 6.57235267308951e-44 | 0.403887679059111 | 4.21353529871768e-40 | microglia1 | RTTN |
| 3.02352015743043e-37 | 0.51280052048403 | 1.93837877292865e-33 | microglia1 | GRID2 |
| 2.30284202614437e-158 | 0.34549365857251 | 1.47635202296116e-154 | microglia2 | FRMD4A |
| 3.10429653136928e-156 | 0.367197954225265 | 1.99016450626084e-152 | microglia2 | ARHGAP15 |
| 6.83323929164981e-150 | 0.324879233408878 | 4.38078970987669e-146 | microglia2 | DOCK8 |
| 9.53532242052666e-143 | 0.315643627055013 | 6.11309520379964e-139 | microglia2 | CELF2 |
| 8.37466867673913e-140 | 0.469526305177982 | 5.36900008865746e-136 | microglia2 | HS3ST4 |
| 3.83427851742181e-116 | 0.388682000255027 | 2.45815595751912e-112 | microglia2 | JAZF1 |
| 9.47905832280907e-112 | 0.308267948538267 | 6.0770242907529e-108 | microglia2 | SSH2 |
| 4.13376308034961e-109 | 0.448111166265465 | 2.65015551081214e-105 | microglia2 | AC244021.1 |
| 1.08551536151055e-102 | 0.438517453277405 | 6.95923898264415e-99 | microglia2 | KCNIP1 |
| 2.90706073251146e-101 | 0.352378876416758 | 1.8637166356131e-97 | microglia2 | CPED1 |
| 3.14189079383148e-92 | 0.392804024976114 | 2.01426618792536e-88 | microglia2 | SYNDIG1 |
| 9.11059713479341e-82 | 0.28848674878056 | 5.84080382311605e-78 | microglia2 | MAML2 |
| 5.69274917173999e-81 | 0.392646984840841 | 3.64962149400251e-77 | microglia2 | ST6GALNAC3 |
| 4.96382588015824e-73 | 0.345815695369694 | 3.18230877176945e-69 | microglia2 | CSGALNACT1 |
| 4.24175895563064e-71 | 0.383935062080751 | 2.71939166645481e-67 | microglia2 | OXR1 |
| 6.07432877749601e-70 | 0.314548527791116 | 3.89425217925269e-66 | microglia2 | IGSF21 |
| 1.87428902635404e-68 | 0.310305349327344 | 1.20160669479558e-64 | microglia2 | LINC02232 |
| 7.562720414784e-66 | 0.299494402097011 | 4.84846005791802e-62 | microglia2 | RUNX1 |
| 1.08312776740464e-62 | 0.333853117882813 | 6.94393211683117e-59 | microglia2 | AC074327.1 |
| 1.54250050522838e-61 | 0.335644689520632 | 9.88897073901917e-58 | microglia2 | FP236383.1 |

| | | | | |
|---|---|---|---|---|
| 1.91304738277684e-61 | 0.373255717301849 | 1.22645467709823e-57 | microglia2 | KHDRBS3 |
| 4.97403337804716e-56 | 0.312602560870122 | 3.18885279866604e-52 | microglia2 | P2RY12 |
| 9.96356073943069e-56 | 0.290351532092622 | 6.38763879004901e-52 | microglia2 | CCND3 |
| 2.53158447768964e-52 | 0.355735579001186 | 1.62299880864683e-48 | microglia2 | FOXP2 |
| 4.46591196274652e-52 | 0.357455140912786 | 2.86309615931679e-48 | microglia2 | FP671120.1 |
| 1.31709464136787e-48 | 0.302198177736203 | 8.44389374580945e-45 | microglia2 | RASGEF1C |
| 8.8293031559213e-46 | 0.455821124095272 | 5.66046625326114e-42 | microglia2 | AC008691.1 |
| 1.74315415308797e-34 | 0.32114529347271 | 1.1175361275447e-30 | microglia2 | SDK1 |
| 1.75070762046287e-27 | 0.296555751632884 | 1.12237865547875e-23 | microglia2 | XIST |
| 2.8651420861946e-17 | 0.375167436211262 | 1.83684259145935e-13 | microglia2 | XACT |
| 0 | 1.2574624834321 | 0 | microglia3 | DIRC3 |
| 0 | 0.934893852040158 | 0 | microglia3 | IQGAP2 |
| 1.13614053614508e-302 | 1.06585748329646 | 7.28379697721492e-299 | microglia3 | STARD13 |
| 1.42197707949018e-301 | 0.910987249691888 | 9.116295070439e-298 | microglia3 | CD163 |
| 8.25592874227553e-290 | 0.931784750472173 | 5.29287591667284e-286 | microglia3 | GPNMB |
| 7.84188897546219e-282 | 1.12008312652162 | 5.02743502216881e-278 | microglia3 | CPM |
| 2.41288155173561e-266 | 0.897727618637195 | 1.5468983628177e-262 | microglia3 | FMN1 |
| 2.98648331506782e-257 | 0.551106740522516 | 1.91463445328998e-253 | microglia3 | SH3PXD2A |
| 2.01290780189572e-249 | 0.599684847440469 | 1.29047519179535e-245 | microglia3 | SLC38A6 |
| 5.2097738623412e-249 | 0.930577776095368 | 3.33998602314694e-245 | microglia3 | RGL1 |
| 1.08085923863714e-235 | 0.594501067441923 | 6.92938857890273e-232 | microglia3 | MCTP1 |
| 1.02739737684784e-229 | 0.681725375077042 | 6.58664458297151e-226 | microglia3 | ATG7 |
| 7.25318183271446e-212 | 0.746071919608708 | 4.65001487295324e-208 | microglia3 | MITF |
| 1.98053891147611e-194 | 0.727678447728046 | 1.26972349614733e-190 | microglia3 | KCNMA1 |
| 6.64322771676677e-193 | 0.59285162659252 | 4.25897328921917e-189 | microglia3 | RBM47 |
| 9.75070848945827e-181 | 0.637652307508836 | 6.25117921259169e-177 | microglia3 | DPYD |
| 6.46230656167923e-169 | 0.629533697802061 | 4.14298473669255e-165 | microglia3 | ABCA1 |
| 1.92898697009103e-152 | 0.626196172597963 | 1.23667354652536e-148 | microglia3 | FRMD4B |
| 4.83983867634672e-150 | 0.629159119989005 | 3.10282057540588e-146 | microglia3 | ZNF804A |
| 3.49507914137852e-146 | 0.862533025993687 | 2.24069523753777e-142 | microglia3 | NHSL1 |
| 8.80700102362758e-145 | 0.628403529701564 | 5.64616835624764e-141 | microglia3 | MERTK |
| 1.2443513499909e-144 | 0.707011570261762 | 7.97753650479167e-141 | microglia3 | TPRG1 |
| 7.90250954919383e-128 | 0.646955294121023 | 5.06629887198816e-124 | microglia3 | PRKCE |
| 2.85246273669291e-126 | 0.658643116455521 | 1.82871386049382e-122 | microglia3 | SAMD4A |
| 8.34435690642751e-117 | 0.601264881503006 | 5.34956721271067e-113 | microglia3 | LGMN |
| 1.72808682579235e-114 | 0.529334108549776 | 1.10787646401547e-110 | microglia3 | NPL |
| 1.76280055839529e-112 | 0.712883258691975 | 1.13013143798722e-108 | microglia3 | MTSS1 |
| 3.17617120490532e-86 | 0.573138768848355 | 2.0362433594648e-82 | microglia3 | MYO1E |
| 5.39309464117672e-76 | 0.599898898204799 | 3.4575129744584e-72 | microglia3 | SELENOP |
| 3.48864559700518e-54 | 0.534598979845195 | 2.23657069224002e-50 | microglia3 | NRP1 |
| 0 | 1.69901468370322 | 0 | microglia4 | FTL |

| | | | | |
|---|---|---|---|---|
| 0 | 1.48524623705572 | 0 | microglia4 | FTH1 |
| 9.11963200594129e-307 | 1.10972069947656 | 5.84659607900896e-303 | microglia4 | RPL13 |
| 1.14867244824368e-302 | 1.15750504814553 | 7.36413906578811e-299 | microglia4 | TMSB4X |
| 2.1651140021162e-295 | 1.31389173159875 | 1.38805458675669e-291 | microglia4 | RPLP1 |
| 2.61313917262018e-284 | 1.04799290483902 | 1.6752835235668e-280 | microglia4 | RPS19 |
| 2.2516390018171e-273 | 1.27473360859807 | 1.44352576406494e-269 | microglia4 | RPL10 |
| 2.93630436957267e-272 | 1.14626565521415 | 1.88246473133304e-268 | microglia4 | RPLP2 |
| 3.83958561887136e-263 | 1.06885294179165 | 2.46155834025843e-259 | microglia4 | RPS15 |
| 1.10236285229072e-248 | 1.1358749863213 | 7.06724824603579e-245 | microglia4 | TMSB10 |
| 4.96439359571287e-248 | 1.11816257442948 | 3.18267273421152e-244 | microglia4 | RPL11 |
| 9.07713425765111e-247 | 1.12420384749698 | 5.81935077258013e-243 | microglia4 | RPL32 |
| 6.14455438221123e-243 | 1.10789426009598 | 3.93927381443562e-239 | microglia4 | RPL19 |
| 2.79660082388919e-236 | 1.2953025802338 | 1.79290078819536e-232 | microglia4 | APOE |
| 7.05358717172878e-233 | 1.0759776318614 | 4.52205473579532e-229 | microglia4 | RPL41 |
| 3.32063519487461e-227 | 0.967894366008239 | 2.12885922343411e-223 | microglia4 | C1QB |
| 8.96455352165037e-226 | 0.935480725301219 | 5.74717526273005e-222 | microglia4 | RPS18 |
| 2.44199817233524e-220 | 1.02098344124661 | 1.56556502828412e-216 | microglia4 | RPL28 |
| 5.73421872395921e-220 | 1.02344301904536 | 3.67620762393025e-216 | microglia4 | TPT1 |
| 1.1399015074234e-214 | 1.00796065401126 | 7.30790856409145e-211 | microglia4 | PLEKHA7 |
| 1.65957490590804e-206 | 0.92856738256143 | 1.06395347217764e-202 | microglia4 | RPS14 |
| 3.88146673377083e-194 | 0.92757302906049 | 2.48840832302048e-190 | microglia4 | RPS3 |
| 1.40097554607986e-182 | 0.904955710139062 | 8.98165422591797e-179 | microglia4 | RPS11 |
| 3.98563289585159e-180 | 0.912490790886114 | 2.55518924953046e-176 | microglia4 | RPS4X |
| 7.99371821166153e-170 | 0.899125568031421 | 5.12477274549621e-166 | microglia4 | C1QA |
| 2.86153295937612e-165 | 0.977181990324365 | 1.83452878025603e-161 | microglia4 | APOC1 |
| 3.48475372350117e-162 | 0.928863133990254 | 2.2340756121366e-158 | microglia4 | RPS24 |
| 5.26164647674179e-155 | 0.901935341944145 | 3.37324155623916e-151 | microglia4 | RPS23 |
| 1.38220926209179e-135 | 0.965673612396673 | 8.86134357927047e-132 | microglia4 | CD14 |
| 5.40303802972109e-115 | 0.944247676281764 | 3.46388768085419e-111 | microglia4 | SPP1 |
| 0 | 1.97501153675965 | 0 | microglia5 | MAGI2 |
| 0 | 1.94448104053882 | 0 | microglia5 | NCAM2 |
| 0 | 1.82477362781068 | 0 | microglia5 | PEX5L |
| 0 | 1.7969377051424 | 0 | microglia5 | SLC24A2 |
| 0 | 1.71178947770203 | 0 | microglia5 | PTPRD |
| 0 | 1.62841955362057 | 0 | microglia5 | MAP7 |
| 0 | 1.62392835086112 | 0 | microglia5 | NTM |
| 0 | 1.61798286305082 | 0 | microglia5 | FRMD5 |
| 0 | 1.50151588663378 | 0 | microglia5 | TTLL7 |
| 0 | 1.47048073234021 | 0 | microglia5 | PRUNE2 |
| 8.37265233708275e-308 | 2.01411179927447 | 5.36770741330375e-304 | microglia5 | PPP2R2B |
| 1.56396569551456e-291 | 1.76375099880026 | 1.00265840739438e-287 | microglia5 | EDIL3 |

| | | | | |
|---|---|---|---|---|
| 3.46873294460673e-288 | 1.925841512512 | 2.22380469078737e-284 | microglia5 | ST18 |
| 1.98138243438494e-276 | 1.53177265392732 | 1.27026427868419e-272 | microglia5 | TMTC2 |
| 5.01771028534467e-271 | 2.7905810571972 | 3.21685406393447e-267 | microglia5 | PCDH9 |
| 1.09924918312325e-256 | 1.60102309141699 | 7.04728651300316e-253 | microglia5 | GPM6B |
| 1.01131432156052e-253 | 1.63272558840468 | 6.48353611552452e-250 | microglia5 | ERBB4 |
| 5.55732939363342e-252 | 1.68780615928211 | 3.56280387425839e-248 | microglia5 | RNF220 |
| 5.26382984345613e-245 | 2.34235030727667 | 3.37464131263972e-241 | microglia5 | IL1RAPL1 |
| 1.03654624113921e-237 | 2.06252074289651 | 6.64529795194347e-234 | microglia5 | CADM2 |
| 3.42128505944128e-230 | 1.46726174245535 | 2.19338585160781e-226 | microglia5 | SPOCK3 |
| 1.65308662506076e-225 | 1.81913579797583 | 1.05979383532645e-221 | microglia5 | CTNNA3 |
| 8.80883911365279e-190 | 1.55803450374151 | 5.6473467557628e-186 | microglia5 | NPAS3 |
| 4.06120984019199e-188 | 1.51401973151388 | 2.60364162854708e-184 | microglia5 | DLG2 |
| 1.62868099020475e-187 | 1.7162636790655 | 1.04414738282027e-183 | microglia5 | SLC44A1 |
| 7.96679075951513e-169 | 1.63475981258851 | 5.10750955592515e-165 | microglia5 | LRP1B |
| 3.25851249044077e-150 | 1.68688677297463 | 2.08903235762158e-146 | microglia5 | LSAMP |
| 2.47155264897257e-108 | 1.99989022332106 | 1.58451240325631e-104 | microglia5 | PLP1 |
| 5.38295434239284e-29 | 1.47306813783036 | 3.45101202890805e-25 | microglia5 | TRPM3 |
| 2.0221086945175e-27 | 1.46709870166279 | 1.29637388405517e-23 | microglia5 | NRG3 |
| 0 | 0.529862402642509 | 0 | oligos0 | SLC38A2 |
| 0 | 0.426510916406258 | 0 | oligos0 | QDPR |
| 0 | 0.415642399296714 | 0 | oligos0 | LAMP2 |
| 0 | 0.401493669075189 | 0 | oligos0 | MID1IP1 |
| 0 | 0.396763035785912 | 0 | oligos0 | AMER2 |
| 0 | 0.39409131747268 | 0 | oligos0 | SUN2 |
| 0 | 0.390283763858314 | 0 | oligos0 | ERBIN |
| 0 | 0.387309441485747 | 0 | oligos0 | PLXDC2 |
| 0 | 0.387170972640179 | 0 | oligos0 | NDRG2 |
| 0 | 0.385663811726068 | 0 | oligos0 | NFIB |
| 0 | 0.356643623292852 | 0 | oligos0 | RGCC |
| 0 | 0.354022909605429 | 0 | oligos0 | GPRC5B |
| 0 | 0.352049563524781 | 0 | oligos0 | KCNMB4 |
| 0 | 0.344539809185015 | 0 | oligos0 | FKBP5 |
| 0 | 0.339152115972386 | 0 | oligos0 | SPOCK1 |
| 0 | 0.335618152929478 | 0 | oligos0 | LRP4 |
| 0 | 0.333686760291406 | 0 | oligos0 | CCP110 |
| 0 | 0.332559476449247 | 0 | oligos0 | SLC6A15 |
| 0 | 0.313856192266016 | 0 | oligos0 | SGK1 |
| 0 | 0.303473232016364 | 0 | oligos0 | KIAA0930 |
| 0 | 0.29208708708815 | 0 | oligos0 | ERBB4 |
| 0 | 0.290306999540693 | 0 | oligos0 | CLMN |
| 0 | 0.278498907314549 | 0 | oligos0 | FCHSD2 |

| | | | | |
|---|---|---|---|---|
| 0 | 0.268901838543256 | 0 | oligos0 | NDRG1 |
| 3.09139557279996e-305 | 0.337141950759615 | 2.22054943657071e-301 | oligos0 | PHYHIPL |
| 1.91565350573282e-286 | 0.281207477523855 | 1.37601391316788e-282 | oligos0 | ZBTB16 |
| 2.99578880239095e-281 | 0.280303682365653 | 2.15187509675742e-277 | oligos0 | LINC00844 |
| 1.59249579131009e-260 | 0.276362689773426 | 1.14388972689804e-256 | oligos0 | CPQ |
| 1.82414608927276e-246 | 0.287474516490403 | 1.31028413592463e-242 | oligos0 | PPA1 |
| 2.90595806237855e-158 | 0.276189685816323 | 2.08734967620651e-154 | oligos0 | ZNF365 |
| 0 | 1.74541257368679 | 0 | oligos1 | RBFOX1 |
| 0 | 0.84197955068278 | 0 | oligos1 | MACROD2 |
| 0 | 0.805276247827256 | 0 | oligos1 | AFF3 |
| 0 | 0.800017900775406 | 0 | oligos1 | RASGRF1 |
| 0 | 0.78339147415064 | 0 | oligos1 | FSTL5 |
| 0 | 0.747498254647312 | 0 | oligos1 | MSRA |
| 0 | 0.734072249036134 | 0 | oligos1 | COL18A1 |
| 0 | 0.713713445144488 | 0 | oligos1 | LINC00609 |
| 0 | 0.697927601807058 | 0 | oligos1 | NRXN1 |
| 0 | 0.604657920990067 | 0 | oligos1 | KCTD8 |
| 0 | 0.603072876759428 | 0 | oligos1 | SLC35F3 |
| 0 | 0.603004242710548 | 0 | oligos1 | ACTN2 |
| 0 | 0.599232458389796 | 0 | oligos1 | ABCA6 |
| 0 | 0.585443672462977 | 0 | oligos1 | TMEM178A |
| 0 | 0.562637938938578 | 0 | oligos1 | SCCPDH |
| 0 | 0.547663149756712 | 0 | oligos1 | SNX24 |
| 0 | 0.545947094322841 | 0 | oligos1 | FBXO32 |
| 0 | 0.54190189586122 | 0 | oligos1 | CALN1 |
| 0 | 0.538623223083935 | 0 | oligos1 | SPP1 |
| 0 | 0.533505583943035 | 0 | oligos1 | SEMA5A |
| 0 | 0.521234399139847 | 0 | oligos1 | SELENOP |
| 0 | 0.517238115757488 | 0 | oligos1 | INPP5F |
| 0 | 0.508009718988598 | 0 | oligos1 | CCSER1 |
| 0 | 0.504836636448284 | 0 | oligos1 | ZDHHC14 |
| 0 | 0.502505063848542 | 0 | oligos1 | DHCR24 |
| 0 | 0.502041669300939 | 0 | oligos1 | PCLO |
| 0 | 0.499545260985086 | 0 | oligos1 | PLEKHG1 |
| 0 | 0.49752359241893 | 0 | oligos1 | FGD4 |
| 0 | 0.493917438884557 | 0 | oligos1 | NEAT1 |
| 1.57529797121293e-276 | 0.624176068958905 | 1.13153653272225e-272 | oligos1 | LINC01505 |
| 0 | 1.00352577361403 | 0 | oligos2 | SYT1 |
| 0 | 0.866601907649417 | 0 | oligos2 | ALDH1A1 |
| 0 | 0.610281615223405 | 0 | oligos2 | UCHL1 |
| 0 | 0.591123630720703 | 0 | oligos2 | SNAP25 |

| | | | | |
|---|---|---|---|---|
| 0 | 0.568383488143585 | 0 | oligos2 | PEG10 |
| 0 | 0.527279226400602 | 0 | oligos2 | STMN2 |
| 0 | 0.517130266235044 | 0 | oligos2 | NEFL |
| 0 | 0.49210726060478 | 0 | oligos2 | SLC18A2 |
| 0 | 0.481274368728525 | 0 | oligos2 | MAP1B |
| 0 | 0.475091356559261 | 0 | oligos2 | TUBB2A |
| 0 | 0.471544174057946 | 0 | oligos2 | SCG2 |
| 0 | 0.46692928894861 | 0 | oligos2 | GALNTL6 |
| 0 | 0.466876426210382 | 0 | oligos2 | OPALIN |
| 0 | 0.460473339362734 | 0 | oligos2 | NEFM |
| 0 | 0.414842840235777 | 0 | oligos2 | SVEP1 |
| 0 | 0.399121255951545 | 0 | oligos2 | NRXN3 |
| 0 | 0.381832150598166 | 0 | oligos2 | YWHAG |
| 0 | 0.379556012060585 | 0 | oligos2 | TUBA1B |
| 0 | 0.37713438952383 | 0 | oligos2 | MT-CO1 |
| 0 | 0.364133151097982 | 0 | oligos2 | PPP2R2B |
| 0 | 0.359848995693135 | 0 | oligos2 | SLC6A3 |
| 0 | 0.3544110744126 | 0 | oligos2 | SV2C |
| 0 | 0.346487008210124 | 0 | oligos2 | THY1 |
| 0 | 0.338407281167295 | 0 | oligos2 | NDRG4 |
| 0 | 0.333122946097715 | 0 | oligos2 | CALY |
| 0 | 0.327880641697645 | 0 | oligos2 | CNTN1 |
| 1.83133180850407e-306 | 0.367212838907358 | 1.31544563804847e-302 | oligos2 | GNAS |
| 6.19382676549793e-305 | 0.413669160320624 | 4.44902575890214e-301 | oligos2 | KCNAB1 |
| 2.01548038315438e-295 | 0.340319904365576 | 1.44771955921979e-291 | oligos2 | NSF |
| 1.85015509696612e-141 | 0.322897140156815 | 1.32896640615077e-137 | oligos2 | PDE1A |
| 0 | 1.37693096328032 | 0 | oligos3 | CTNNA2 |
| 0 | 0.890507147033957 | 0 | oligos3 | LAMA2 |
| 0 | 0.801364140911627 | 0 | oligos3 | XIST |
| 0 | 0.6937638684925 | 0 | oligos3 | GRIN2A |
| 0 | 0.68068006845283 | 0 | oligos3 | KCNAB1 |
| 0 | 0.637243045520704 | 0 | oligos3 | FCHSD2 |
| 0 | 0.602186483812519 | 0 | oligos3 | ROR1 |
| 0 | 0.59500417139533 | 0 | oligos3 | LRRC7 |
| 0 | 0.57631609060962 | 0 | oligos3 | PALM2 |
| 0 | 0.576275924212079 | 0 | oligos3 | SVEP1 |
| 0 | 0.567241109964659 | 0 | oligos3 | CTNND2 |
| 0 | 0.557855805747709 | 0 | oligos3 | PLXDC2 |
| 0 | 0.547905821149055 | 0 | oligos3 | LINC01608 |
| 0 | 0.53710499382005 | 0 | oligos3 | ANK3 |
| 0 | 0.537055540816676 | 0 | oligos3 | AUTS2 |

| | | | | |
|---|---|---|---|---|
| 0 | 0.531725713405106 | 0 | oligos3 | ASTN2 |
| 0 | 0.506790003820184 | 0 | oligos3 | ANO4 |
| 0 | 0.502823387168423 | 0 | oligos3 | PPFIA2 |
| 0 | 0.494595374834865 | 0 | oligos3 | KCNH8 |
| 0 | 0.493012382281819 | 0 | oligos3 | NLGN1 |
| 0 | 0.486803334888019 | 0 | oligos3 | OPALIN |
| 0 | 0.462493040026598 | 0 | oligos3 | PPP2R2B |
| 0 | 0.406998634351884 | 0 | oligos3 | PLCL2 |
| 0 | 0.397902525731621 | 0 | oligos3 | CREB5 |
| 0 | 0.393560251292445 | 0 | oligos3 | FRY |
| 0 | 0.385137551265938 | 0 | oligos3 | DYSF |
| 2.26132494078155e-304 | 0.432613918696476 | 1.62430970490472e-300 | oligos3 | NRXN3 |
| 1.11695588617096e-294 | 0.398310997686247 | 8.02309413036601e-291 | oligos3 | TIAM1 |
| 1.86969867441488e-231 | 0.558479337626419 | 1.34300455783221e-227 | oligos3 | DCC |
| 1.48522700122473e-102 | 0.388807917847227 | 1.06683855497972e-98 | oligos3 | PTPRM |
| 0 | 0.454916598997 | 0 | oligos4 | SLC5A11 |
| 0 | 0.430998986764344 | 0 | oligos4 | OTUD7A |
| 0 | 0.319420117498636 | 0 | oligos4 | PCDH9 |
| 1.12369449964832e-235 | 0.278772975973642 | 8.07149759097391e-232 | oligos4 | RNF220 |
| 5.97064268105594e-217 | 0.273020781040593 | 4.28871263780248e-213 | oligos4 | DOCK5 |
| 7.85599512653413e-215 | 0.372492531688983 | 5.64296129938947e-211 | oligos4 | HIP1 |
| 7.15172135283743e-202 | 0.495479600936655 | 5.13708144774313e-198 | oligos4 | FP236383.1 |
| 1.12260885993137e-172 | 0.256171201779147 | 8.06369944088705e-169 | oligos4 | DST |
| 6.0341042553593e-170 | 0.392301343279602 | 4.33429708662459e-166 | oligos4 | L3MBTL4 |
| 7.73357798591512e-166 | 0.337059431727063 | 5.55502906728283e-162 | oligos4 | KIRREL3 |
| 4.0268523008002e-162 | 0.277155447339414 | 2.89248800766478e-158 | oligos4 | DPYD |
| 1.80751981503957e-157 | 0.288186877222417 | 1.29834148314292e-153 | oligos4 | DSCAML1 |
| 5.22323967360136e-157 | 0.523745278548172 | 3.75185305754786e-153 | oligos4 | FP671120.1 |
| 1.32994670907237e-153 | 0.289326319921451 | 9.55300721126682e-150 | oligos4 | SAMD12 |
| 1.37608706208081e-140 | 0.384632061399834 | 9.88443336692644e-137 | oligos4 | AC026316.5 |
| 1.11528177414574e-130 | 0.259647350348251 | 8.01106898368888e-127 | oligos4 | DOCK3 |
| 1.43622739050163e-123 | 0.325967400261248 | 1.03164213459732e-119 | oligos4 | GREB1L |
| 1.58822703514353e-117 | 0.297940070833406 | 1.1408234793436e-113 | oligos4 | GNG7 |
| 4.98871815608922e-117 | 0.280190490479776 | 3.58339625151889e-113 | oligos4 | FHIT |
| 6.64180860646684e-110 | 0.271745556673348 | 4.77081112202513e-106 | oligos4 | GPI |
| 7.61332026811588e-110 | 0.263548571842852 | 5.46864794858764e-106 | oligos4 | SLC22A15 |
| 3.22590833421908e-104 | 0.257082762394883 | 2.31716995646956e-100 | oligos4 | ZHX2 |
| 1.73315768819687e-103 | 0.278802641199196 | 1.24492716743181e-99 | oligos4 | ANKRD18A |
| 2.11678878810634e-96 | 0.25848203174368 | 1.52048938649679e-92 | oligos4 | MT-ND4 |
| 8.83995168564095e-90 | 0.273210515511183 | 6.3497372957959e-86 | oligos4 | LINC00854 |
| 1.59824119154777e-89 | 0.264268174234701 | 1.14801664788877e-85 | oligos4 | EVA1C |

| | | | | |
|---|---|---|---|---|
| 4.04828882108007e-83 | 0.278978480716574 | 2.90788586018181e-79 | oligos4 | AC106869.1 |
| 8.8768947356477e-73 | 0.354691295822034 | 6.37627348861574e-69 | oligos4 | AC012593.1 |
| 2.9000277560138e-25 | 0.338815777331657 | 2.08308993714471e-21 | oligos4 | XIST |
| 7.25393432643058e-20 | 0.282217063952454 | 5.21050102667509e-16 | oligos4 | AC100801.1 |
| 0 | 1.49046314044338 | 0 | oligos5 | CRYAB |
| 0 | 1.45347348589548 | 0 | oligos5 | S100B |
| 0 | 1.4503309521293 | 0 | oligos5 | FTH1 |
| 0 | 1.38995738978924 | 0 | oligos5 | FTL |
| 0 | 1.07020776568975 | 0 | oligos5 | GAPDH |
| 0 | 1.05956648168814 | 0 | oligos5 | PPP1R14A |
| 0 | 0.959066507951811 | 0 | oligos5 | MARCKSL1 |
| 0 | 0.900760542184166 | 0 | oligos5 | NKX6-2 |
| 0 | 0.896696113250431 | 0 | oligos5 | FEZ1 |
| 0 | 0.857546386353958 | 0 | oligos5 | PLCG2 |
| 0 | 0.84646456109344 | 0 | oligos5 | RPL10 |
| 0 | 0.842755052145574 | 0 | oligos5 | SVIP |
| 0 | 0.840588142582199 | 0 | oligos5 | MAG |
| 0 | 0.835186737365357 | 0 | oligos5 | C4orf48 |
| 0 | 0.835135039211081 | 0 | oligos5 | RPL28 |
| 0 | 0.829477616471715 | 0 | oligos5 | RPS15 |
| 0 | 0.794164850431862 | 0 | oligos5 | MT3 |
| 0 | 0.790730014068484 | 0 | oligos5 | RPLP1 |
| 0 | 0.785262654835824 | 0 | oligos5 | TSC22D4 |
| 0 | 0.784742912876312 | 0 | oligos5 | STMN4 |
| 0 | 0.778915171966516 | 0 | oligos5 | ACTB |
| 0 | 0.774897055958038 | 0 | oligos5 | LGALS1 |
| 0 | 0.766884415630953 | 0 | oligos5 | S100A6 |
| 0 | 0.759409976586994 | 0 | oligos5 | FIS1 |
| 0 | 0.752512320768352 | 0 | oligos5 | CDKN1C |
| 0 | 0.727705574247061 | 0 | oligos5 | EIF1 |
| 0 | 0.709397656067712 | 0 | oligos5 | RPL13 |
| 0 | 0.698626673117015 | 0 | oligos5 | SERF2 |
| 0 | 0.690529306902338 | 0 | oligos5 | DYNC1I2 |
| 0 | 0.68662192637845 | 0 | oligos5 | APLP1 |

**Table 6:** Pathway (GO and KEGG) over-representation analysis on cell-type markers reported in **Table 5.**

| ID | Description | GeneRatio | BgRatio | pvalue | p.adjust | Count | subtype | groupName |
|---|---|---|---|---|---|---|---|---|
| GO:0006458 | 'de novo' protein folding | 9/616 | 41/18800 | 1E-05 | 0.00014 | 9 | neurons0 | Unfolded Protein Response |
| GO:0042743 | hydrogen peroxide metabolic process | 10/616 | 55/18800 | 1E-05 | 0.00025 | 10 | neurons0 | Oxidative stress |
| GO:0008637 | apoptotic mitochondrial changes | 14/616 | 107/18800 | 1E-05 | 0.00025 | 14 | neurons0 | Apoptosis |
| GO:0055076 | transition metal ion homeostasis | 16/616 | 139/18800 | 1E-05 | 0.00031 | 16 | neurons0 | Ion transport |
| GO:0034975 | protein folding in endoplasmic reticulum | 5/616 | 11/18800 | 1E-05 | 0.00034 | 5 | neurons0 | Unfolded Protein Response |
| GO:0048499 | synaptic vesicle membrane organization | 7/616 | 26/18800 | 1E-05 | 0.00034 | 7 | neurons0 | Vesicle |
| GO:0051881 | regulation of mitochondrial membrane potential | 11/616 | 71/18800 | 2E-05 | 0.00041 | 11 | neurons0 | Mitochondrial changes |
| GO:0055072 | iron ion homeostasis | 12/616 | 85/18800 | 2E-05 | 0.00045 | 12 | neurons0 | Ion transport |
| GO:0014046 | dopamine secretion | 8/616 | 37/18800 | 2E-05 | 0.00045 | 8 | neurons0 | Dopamine metabolism |
| GO:0014059 | regulation of dopamine secretion | 8/616 | 37/18800 | 2E-05 | 0.00045 | 8 | neurons0 | Dopamine metabolism |
| GO:0051084 | 'de novo' post-translational protein folding | 8/616 | 37/18800 | 2E-05 | 0.00045 | 8 | neurons0 | Unfolded Protein Response |
| GO:0042776 | proton motive force-driven mitochondrial ATP synthesis | 6/616 | 19/18800 | 2E-05 | 0.00048 | 6 | neurons0 | Mitochondrial changes |
| GO:0099173 | postsynapse organization | 17/616 | 163/18800 | 3E-05 | 0.00054 | 17 | neurons0 | Synapse assembly |
| GO:0042053 | regulation of dopamine metabolic process | 6/616 | 20/18800 | 3E-05 | 0.00064 | 6 | neurons0 | Dopamine metabolism |
| GO:0017158 | regulation of calcium ion-dependent exocytosis | 8/616 | 40/18800 | 4E-05 | 0.00077 | 8 | neurons0 | Calcium transport |
| GO:0017156 | calcium-ion regulated exocytosis | 10/616 | 65/18800 | 5E-05 | 0.00089 | 10 | neurons0 | Calcium transport |
| GO:0042417 | dopamine metabolic process | 8/616 | 42/18800 | 6E-05 | 0.00104 | 8 | neurons0 | Dopamine metabolism |
| GO:0031338 | regulation of vesicle fusion | 6/616 | 22/18800 | 6E-05 | 0.00105 | 6 | neurons0 | Vesicle |
| GO:0051085 | chaperone cofactor-dependent protein refolding | 7/616 | 32/18800 | 6E-05 | 0.00117 | 7 | neurons0 | Unfolded Protein Response |
| GO:2000300 | regulation of synaptic vesicle exocytosis | 9/616 | 56/18800 | 8E-05 | 0.0014 | 9 | neurons0 | Vesicle |
| GO:0007416 | synapse assembly | 17/616 | 180/18800 | 9E-05 | 0.00159 | 17 | neurons0 | Synapse assembly |
| GO:0047496 | vesicle transport along microtubule | 8/616 | 45/18800 | 9E-05 | 0.0016 | 8 | neurons0 | Vesicle |
| GO:0099174 | regulation of presynapse organization | 7/616 | 34/18800 | 1E-04 | 0.00164 | 7 | neurons0 | Synapse assembly |
| GO:1905606 | regulation of presynapse assembly | 7/616 | 34/18800 | 1E-04 | 0.00164 | 7 | neurons0 | Synapse assembly |
| GO:0032469 | endoplasmic reticulum calcium ion homeostasis | 6/616 | 25/18800 | 0.00013 | 0.00206 | 6 | neurons0 | Calcium transport |

| | | | | | | | | |
|---|---|---|---|---|---|---|---|---|
| GO:2001243 | negative regulation of intrinsic apoptotic signaling pathway | 12/616 | 102/18800 | 0.00013 | 0.00206 | 12 | neurons0 | Apoptosis |
| GO:0016239 | positive regulation of macroautophagy | 10/616 | 73/18800 | 0.00013 | 0.00208 | 10 | neurons0 | Autophagy |
| GO:1903146 | regulation of autophagy of mitochondrion | 7/616 | 36/18800 | 0.00014 | 0.00229 | 7 | neurons0 | Autophagy |
| GO:1903578 | regulation of ATP metabolic process | 11/616 | 89/18800 | 0.00015 | 0.00242 | 11 | neurons0 | Energy production |
| GO:0045956 | positive regulation of calcium ion-dependent exocytosis | 5/616 | 17/18800 | 0.00017 | 0.00255 | 5 | neurons0 | Calcium transport |
| GO:0048791 | calcium ion-regulated exocytosis of neurotransmitter | 5/616 | 17/18800 | 0.00017 | 0.00255 | 5 | neurons0 | Calcium transport |
| GO:0099054 | presynapse assembly | 8/616 | 50/18800 | 2E-04 | 3 | 8 | neurons0 | Synapse assembly |
| GO:2001234 | negative regulation of apoptotic signaling pathway | 19/616 | 230/18800 | 0.00022 | 0.00318 | 19 | neurons0 | Apoptosis |
| GO:0006979 | response to oxidative stress | 29/616 | 433/18800 | 0.00024 | 0.0034 | 29 | neurons0 | Oxidative stress |
| GO:0097479 | synaptic vesicle localization | 8/616 | 52/18800 | 0.00027 | 0.00379 | 8 | neurons0 | Vesicle |
| GO:0099172 | presynapse organization | 8/616 | 52/18800 | 0.00027 | 0.00379 | 8 | neurons0 | Synapse assembly |
| GO:0048489 | synaptic vesicle transport | 7/616 | 40/18800 | 0.00028 | 0.00397 | 7 | neurons0 | Vesicle |
| GO:0098780 | response to mitochondrial depolarisation | 5/616 | 19/18800 | 0.00029 | 0.0041 | 5 | neurons0 | Mitochondrial changes |
| GO:0006122 | mitochondrial electron transport, ubiquinol to cytochrome c | 4/616 | 11/18800 | 0.00031 | 0.00424 | 4 | neurons0 | Mitochondrial changes |
| GO:0042744 | hydrogen peroxide catabolic process | 6/616 | 30/18800 | 0.00037 | 0.00486 | 6 | neurons0 | Oxidative stress |
| GO:0051402 | neuron apoptotic process | 19/616 | 241/18800 | 0.00039 | 0.00515 | 19 | neurons0 | Apoptosis |
| GO:0098869 | cellular oxidant detoxification | 11/616 | 100/18800 | 0.00043 | 0.00562 | 11 | neurons0 | Oxidative stress |
| GO:0016188 | synaptic vesicle maturation | 4/616 | 12/18800 | 0.00046 | 0.00577 | 4 | neurons0 | Vesicle |
| GO:0043467 | regulation of generation of precursor metabolites and energy | 13/616 | 134/18800 | 0.00046 | 0.00584 | 13 | neurons0 | Energy production |
| GO:0031629 | synaptic vesicle fusion to presynaptic active zone membrane | 5/616 | 21/18800 | 0.00049 | 0.00609 | 5 | neurons0 | Vesicle |
| GO:0099518 | vesicle cytoskeletal trafficking | 9/616 | 71/18800 | 5E-04 | 0.0062 | 9 | neurons0 | Vesicle |
| GO:1902235 | regulation of endoplasmic reticulum stress-induced intrinsic apoptotic signaling pathway | 6/616 | 32/18800 | 0.00053 | 0.0065 | 6 | neurons0 | Apoptosis |
| GO:0099500 | vesicle fusion to plasma membrane | 5/616 | 22/18800 | 0.00062 | 0.00732 | 5 | neurons0 | Vesicle |
| GO:1903599 | positive regulation of autophagy of mitochondrion | 4/616 | 13/18800 | 0.00064 | 0.00749 | 4 | neurons0 | Autophagy |
| GO:1904925 | positive regulation of autophagy of mitochondrion in response to mitochondrial depolarization | 4/616 | 13/18800 | 0.00064 | 0.00749 | 4 | neurons0 | Mitochondrial changes |

| | | | | | | | | |
|---|---|---|---|---|---|---|---|---|
| GO:0060560 | developmental growth involved in morphogenesis | 18/616 | 234/18800 | 0.00073 | 0.00842 | 18 | neurons0 | Morphogenesis |
| GO:0097345 | mitochondrial outer membrane permeabilization | 6/616 | 35/18800 | 0.00087 | 0.00961 | 6 | neurons0 | Mitochondrial changes |
| GO:1904923 | regulation of autophagy of mitochondrion in response to mitochondrial depolarization | 4/616 | 14/18800 | 0.00088 | 0.00961 | 4 | neurons0 | Mitochondrial changes |
| GO:0051560 | mitochondrial calcium ion homeostasis | 5/616 | 25/18800 | 0.00115 | 0.01211 | 5 | neurons0 | Mitochondrial changes |
| GO:0043281 | regulation of cysteine-type endopeptidase activity involved in apoptotic process | 16/616 | 204/18800 | 0.00116 | 0.01211 | 16 | neurons0 | Apoptosis |
| GO:0034620 | cellular response to unfolded protein | 10/616 | 98/18800 | 0.00139 | 0.01421 | 10 | neurons0 | Unfolded Protein Response |
| GO:0035418 | protein localization to synapse | 8/616 | 67/18800 | 0.00151 | 0.01516 | 8 | neurons0 | Synapse assembly |
| GO:2001235 | positive regulation of apoptotic signaling pathway | 12/616 | 134/18800 | 0.00153 | 0.01529 | 12 | neurons0 | Apoptosis |
| GO:0001963 | synaptic transmission, dopaminergic | 5/616 | 27/18800 | 0.00165 | 0.01614 | 5 | neurons0 | Dopamine metabolism |
| GO:0010508 | positive regulation of autophagy | 12/616 | 136/18800 | 0.00174 | 0.01688 | 12 | neurons0 | Autophagy |
| GO:0007007 | inner mitochondrial membrane organization | 6/616 | 40/18800 | 0.00179 | 0.01736 | 6 | neurons0 | Mitochondrial changes |
| GO:0007212 | dopamine receptor signaling pathway | 6/616 | 40/18800 | 0.00179 | 0.01736 | 6 | neurons0 | Dopamine metabolism |
| GO:0099175 | regulation of postsynapse organization | 9/616 | 86/18800 | 2 | 0.0191 | 9 | neurons0 | Synapse assembly |
| GO:0051963 | regulation of synapse assembly | 10/616 | 103/18800 | 0.00203 | 0.01917 | 10 | neurons0 | Synapse assembly |
| GO:1902110 | positive regulation of mitochondrial membrane permeability involved in apoptotic process | 6/616 | 41/18800 | 0.00204 | 0.01921 | 6 | neurons0 | Apoptosis |
| GO:0098876 | vesicle-mediated transport to the plasma membrane | 12/616 | 139/18800 | 0.00209 | 0.01958 | 12 | neurons0 | Vesicle |
| GO:0097120 | receptor localization to synapse | 7/616 | 56/18800 | 0.00226 | 0.02102 | 7 | neurons0 | Synapse assembly |
| GO:1902175 | regulation of oxidative stress-induced intrinsic apoptotic signaling pathway | 5/616 | 29/18800 | 0.0023 | 0.02135 | 5 | neurons0 | Apoptosis |
| GO:0006906 | vesicle fusion | 10/616 | 105/18800 | 0.00234 | 0.02167 | 10 | neurons0 | Vesicle |
| GO:1902686 | mitochondrial outer membrane permeabilization involved in programmed cell death | 6/616 | 43/18800 | 0.00262 | 0.0238 | 6 | neurons0 | Mitochondrial changes |
| GO:0090151 | establishment of protein localization to mitochondrial membrane | 5/616 | 30/18800 | 0.00268 | 0.02414 | 5 | neurons0 | Mitochondrial changes |
| GO:0034599 | cellular response to oxidative stress | 19/616 | 284/18800 | 0.00271 | 0.02423 | 19 | neurons0 | Oxidative stress |
| GO:0006839 | mitochondrial transport | 14/616 | 182/18800 | 0.00275 | 0.02453 | 14 | neurons0 | Mitochondrial changes |

| | | | | | | | | |
|---|---|---|---|---|---|---|---|---|
| GO:0006695 | cholesterol biosynthetic process | 7/616 | 58/18800 | 0.00277 | 0.02457 | 7 | neurons0 | Cholesterol metabolism |
| GO:0043524 | negative regulation of neuron apoptotic process | 12/616 | 145/18800 | 0.00297 | 0.02603 | 12 | neurons0 | Apoptosis |
| GO:0008631 | intrinsic apoptotic signaling pathway in response to oxidative stress | 6/616 | 45/18800 | 0.00332 | 0.02838 | 6 | neurons0 | Apoptosis |
| GO:0055074 | calcium ion homeostasis | 27/616 | 468/18800 | 0.00336 | 0.02849 | 27 | neurons0 | Calcium transport |
| GO:0061912 | selective autophagy | 8/616 | 76/18800 | 0.00338 | 0.02855 | 8 | neurons0 | Autophagy |
| GO:0043523 | regulation of neuron apoptotic process | 15/616 | 207/18800 | 0.00349 | 0.02912 | 15 | neurons0 | Apoptosis |
| GO:0043653 | mitochondrial fragmentation involved in apoptotic process | 3/616 | 10/18800 | 0.00354 | 0.02912 | 3 | neurons0 | Apoptosis |
| GO:1990034 | calcium ion export across plasma membrane | 3/616 | 10/18800 | 0.00354 | 0.02912 | 3 | neurons0 | Calcium transport |
| GO:0010310 | regulation of hydrogen peroxide metabolic process | 4/616 | 20/18800 | 0.00364 | 0.02951 | 4 | neurons0 | Oxidative stress |
| GO:1902236 | negative regulation of endoplasmic reticulum stress-induced intrinsic apoptotic signaling pathway | 4/616 | 20/18800 | 0.00364 | 0.02951 | 4 | neurons0 | Apoptosis |
| GO:0043154 | negative regulation of cysteine-type endopeptidase activity involved in apoptotic process | 8/616 | 77/18800 | 0.00366 | 0.0296 | 8 | neurons0 | Apoptosis |
| GO:0035794 | positive regulation of mitochondrial membrane permeability | 6/616 | 46/18800 | 0.00371 | 0.0296 | 6 | neurons0 | Mitochondrial changes |
| GO:0036475 | neuron death in response to oxidative stress | 5/616 | 33/18800 | 0.00412 | 0.03244 | 5 | neurons0 | Oxidative stress |
| GO:0090141 | positive regulation of mitochondrial fission | 4/616 | 21/18800 | 0.00438 | 0.03434 | 4 | neurons0 | Mitochondrial changes |
| GO:0031346 | positive regulation of cell projection organization | 21/616 | 341/18800 | 0.00441 | 0.03445 | 21 | neurons0 | Cell projection organization |
| GO:0070059 | intrinsic apoptotic signaling pathway in response to endoplasmic reticulum stress | 7/616 | 63/18800 | 0.00443 | 0.0345 | 7 | neurons0 | Apoptosis |
| GO:1902108 | regulation of mitochondrial membrane permeability involved in apoptotic process | 6/616 | 48/18800 | 0.0046 | 0.03547 | 6 | neurons0 | Apoptosis |
| GO:0006874 | cellular calcium ion homeostasis | 26/616 | 456/18800 | 0.00462 | 0.03554 | 26 | neurons0 | Calcium transport |
| GO:0051561 | positive regulation of mitochondrial calcium ion concentration | 3/616 | 11/18800 | 0.00474 | 0.03604 | 3 | neurons0 | Mitochondrial changes |
| GO:0050770 | regulation of axonogenesis | 12/616 | 154/18800 | 0.00483 | 0.03651 | 12 | neurons0 | Axonogenesis |
| GO:0035967 | cellular response to topologically incorrect protein | 10/616 | 117/18800 | 0.00512 | 0.03835 | 10 | neurons0 | Unfolded Protein Response |
| GO:0062237 | protein localization to postsynapse | 5/616 | 36/18800 | 0.00604 | 0.04389 | 5 | neurons0 | Synapse assembly |
| GO:0008203 | cholesterol metabolic process | 11/616 | 139/18800 | 0.00609 | 0.04389 | 11 | neurons0 | Cholesterol metabolism |

| | | | | | | | | |
|---|---|---|---|---|---|---|---|---|
| GO:0140112 | extracellular vesicle biogenesis | 4/616 | 23/18800 | 0.00615 | 0.04389 | 4 | neurons0 | Vesicle |
| GO:0051583 | dopamine uptake involved in synaptic transmission | 3/616 | 12/18800 | 0.00617 | 0.04389 | 3 | neurons0 | Dopamine metabolism |
| GO:1903580 | positive regulation of ATP metabolic process | 5/616 | 37/18800 | 0.0068 | 0.04734 | 5 | neurons0 | Energy production |
| GO:0005513 | detection of calcium ion | 3/616 | 13/18800 | 0.00783 | 0.05343 | 3 | neurons0 | Calcium transport |
| GO:0043280 | positive regulation of cysteine-type endopeptidase activity involved in apoptotic process | 10/616 | 125/18800 | 0.00809 | 0.05489 | 10 | neurons0 | Apoptosis |
| GO:0060074 | synapse maturation | 4/616 | 25/18800 | 0.00835 | 0.05579 | 4 | neurons0 | Synapse assembly |
| GO:0000266 | mitochondrial fission | 5/616 | 40/18800 | 0.00948 | 0.06225 | 5 | neurons0 | Mitochondrial changes |
| GO:0010882 | regulation of cardiac muscle contraction by calcium ion signaling | 4/616 | 26/18800 | 0.00962 | 0.0628 | 4 | neurons0 | Muscle contraction |
| GO:0051204 | protein insertion into mitochondrial membrane | 4/616 | 26/18800 | 0.00962 | 0.0628 | 4 | neurons0 | Mitochondrial changes |
| GO:0090128 | regulation of synapse maturation | 3/616 | 14/18800 | 0.00973 | 0.0628 | 3 | neurons0 | Synapse assembly |
| GO:0048278 | vesicle docking | 6/616 | 57/18800 | 0.01063 | 0.06813 | 6 | neurons0 | Vesicle |
| GO:0006809 | nitric oxide biosynthetic process | 7/616 | 75/18800 | 0.01136 | 0.07182 | 7 | neurons0 | Oxidative stress |
| GO:0060997 | dendritic spine morphogenesis | 6/616 | 58/18800 | 0.01154 | 0.07273 | 6 | neurons0 | Morphogenesis |
| GO:0008053 | mitochondrial fusion | 4/616 | 28/18800 | 0.01251 | 0.07666 | 4 | neurons0 | Mitochondrial changes |
| GO:0006919 | activation of cysteine-type endopeptidase activity involved in apoptotic process | 7/616 | 77/18800 | 0.01302 | 0.07865 | 7 | neurons0 | Apoptosis |
| GO:0050772 | positive regulation of axonogenesis | 7/616 | 77/18800 | 0.01302 | 0.07865 | 7 | neurons0 | Axonogenesis |
| GO:0061001 | regulation of dendritic spine morphogenesis | 5/616 | 44/18800 | 0.01407 | 0.08411 | 5 | neurons0 | Morphogenesis |
| GO:1903203 | regulation of oxidative stress-induced neuron death | 4/616 | 29/18800 | 0.01415 | 0.08416 | 4 | neurons0 | Oxidative stress |
| GO:0006878 | cellular copper ion homeostasis | 3/616 | 16/18800 | 0.01426 | 0.08416 | 3 | neurons0 | Ion transport |
| GO:0045428 | regulation of nitric oxide biosynthetic process | 6/616 | 61/18800 | 0.01459 | 0.08554 | 6 | neurons0 | Oxidative stress |
| GO:2001244 | positive regulation of intrinsic apoptotic signaling pathway | 6/616 | 61/18800 | 0.01459 | 0.08554 | 6 | neurons0 | Apoptosis |
| GO:0010770 | positive regulation of cell morphogenesis involved in differentiation | 7/616 | 79/18800 | 0.01485 | 0.08683 | 7 | neurons0 | Morphogenesis |
| GO:0046902 | regulation of mitochondrial membrane permeability | 6/616 | 62/18800 | 0.01572 | 0.09153 | 6 | neurons0 | Mitochondrial changes |
| GO:0042063 | gliogenesis | 17/616 | 291/18800 | 0.01584 | 0.09207 | 17 | neurons0 | Gliogenesis |
| GO:0090140 | regulation of mitochondrial fission | 4/616 | 30/18800 | 0.01591 | 0.09236 | 4 | neurons0 | Mitochondrial changes |

| | | | | | | | | |
|---|---|---|---|---|---|---|---|---|
| GO:0048813 | dendrite morphogenesis | 10/616 | 139/18800 | 0.01625 | 0.09392 | 10 | neurons0 | Morphogenesis |
| GO:0046209 | nitric oxide metabolic process | 7/616 | 81/18800 | 0.01686 | 0.09593 | 7 | neurons0 | Oxidative stress |
| GO:0071732 | cellular response to nitric oxide | 3/616 | 17/18800 | 0.0169 | 0.09593 | 3 | neurons0 | Oxidative stress |
| GO:0090494 | dopamine uptake | 3/616 | 17/18800 | 0.0169 | 0.09593 | 3 | neurons0 | Dopamine metabolism |
| GO:0051965 | positive regulation of synapse assembly | 6/616 | 63/18800 | 0.01691 | 0.09593 | 6 | neurons0 | Synapse assembly |
| GO:0098685 | Schaffer collateral - CA1 synapse | 11/636 | 72/19594 | 2E-05 | 0.00013 | 11 | neurons0 | Synapse assembly |
| GO:0032279 | asymmetric synapse | 26/636 | 323/19594 | 2E-05 | 0.00014 | 26 | neurons0 | Synapse assembly |
| GO:0098563 | intrinsic component of synaptic vesicle membrane | 8/636 | 40/19594 | 4E-05 | 0.00023 | 8 | neurons0 | Vesicle |
| GO:0005750 | mitochondrial respiratory chain complex III | 5/636 | 14/19594 | 6E-05 | 0.00033 | 5 | neurons0 | Mitochondrial changes |
| GO:0030136 | clathrin-coated vesicle | 17/636 | 192/19594 | 0.00018 | 0.00096 | 17 | neurons0 | Vesicle |
| GO:0097470 | ribbon synapse | 4/636 | 10/19594 | 2E-04 | 0.00102 | 4 | neurons0 | Synapse assembly |
| GO:0030285 | integral component of synaptic vesicle membrane | 6/636 | 28/19594 | 0.00023 | 0.00116 | 6 | neurons0 | Vesicle |
| GO:0030120 | vesicle coat | 8/636 | 52/19594 | 0.00025 | 0.00124 | 8 | neurons0 | Vesicle |
| GO:0005798 | Golgi-associated vesicle | 10/636 | 84/19594 | 0.00038 | 0.0018 | 10 | neurons0 | Vesicle |
| GO:0060198 | clathrin-sculpted vesicle | 4/636 | 12/19594 | 0.00044 | 0.00202 | 4 | neurons0 | Vesicle |
| GO:0042645 | mitochondrial nucleoid | 7/636 | 44/19594 | 0.00049 | 0.00221 | 7 | neurons0 | Mitochondrial changes |
| GO:0030140 | trans-Golgi network transport vesicle | 6/636 | 32/19594 | 5E-04 | 0.00223 | 6 | neurons0 | Vesicle |
| GO:0032592 | integral component of mitochondrial membrane | 10/636 | 88/19594 | 0.00056 | 0.00247 | 10 | neurons0 | Mitochondrial changes |
| GO:0030666 | endocytic vesicle membrane | 16/636 | 194/19594 | 0.00061 | 0.00266 | 16 | neurons0 | Vesicle |
| GO:0030130 | clathrin coat of trans-Golgi network vesicle | 4/636 | 13/19594 | 0.00062 | 0.00268 | 4 | neurons0 | Vesicle |
| GO:0030662 | coated vesicle membrane | 15/636 | 176/19594 | 0.00064 | 0.00274 | 15 | neurons0 | Vesicle |
| GO:0098573 | intrinsic component of mitochondrial membrane | 10/636 | 90/19594 | 0.00067 | 0.00284 | 10 | neurons0 | Mitochondrial changes |
| GO:0045334 | clathrin-coated endocytic vesicle | 10/636 | 91/19594 | 0.00073 | 0.00308 | 10 | neurons0 | Vesicle |
| GO:0005759 | mitochondrial matrix | 29/636 | 473/19594 | 0.00085 | 0.00353 | 29 | neurons0 | Mitochondrial changes |
| GO:0031307 | integral component of mitochondrial outer membrane | 5/636 | 24/19594 | 9E-04 | 0.00373 | 5 | neurons0 | Mitochondrial changes |
| GO:0005751 | mitochondrial respiratory chain complex IV | 5/636 | 25/19594 | 0.0011 | 0.00441 | 5 | neurons0 | Mitochondrial changes |
| GO:0031306 | intrinsic component of mitochondrial outer membrane | 5/636 | 25/19594 | 0.0011 | 0.00441 | 5 | neurons0 | Mitochondrial changes |
| GO:0012510 | trans-Golgi network transport vesicle membrane | 4/636 | 15/19594 | 0.00113 | 0.00447 | 4 | neurons0 | Vesicle |

| | | | | | | | | |
|---|---|---|---|---|---|---|---|---|
| GO:0030135 | coated vesicle | 20/636 | 290/19594 | 0.00133 | 0.00516 | 20 | neurons0 | Vesicle |
| GO:0045335 | phagocytic vesicle | 12/636 | 138/19594 | 0.00182 | 0.00702 | 12 | neurons0 | Vesicle |
| GO:0030669 | clathrin-coated endocytic vesicle membrane | 8/636 | 72/19594 | 0.00227 | 0.00863 | 8 | neurons0 | Vesicle |
| GO:0005742 | mitochondrial outer membrane translocase complex | 4/636 | 21/19594 | 0.00424 | 0.01505 | 4 | neurons0 | Mitochondrial changes |
| GO:0000276 | mitochondrial proton-transporting ATP synthase complex, coupling factor F(o) | 3/636 | 11/19594 | 0.00462 | 0.01631 | 3 | neurons0 | Mitochondrial changes |
| GO:0005747 | mitochondrial respiratory chain complex I | 6/636 | 49/19594 | 0.00488 | 0.01691 | 6 | neurons0 | Mitochondrial changes |
| GO:0098799 | outer mitochondrial membrane protein complex | 4/636 | 22/19594 | 0.00505 | 0.0174 | 4 | neurons0 | Mitochondrial changes |
| GO:0005758 | mitochondrial intermembrane space | 8/636 | 83/19594 | 0.00549 | 0.01858 | 8 | neurons0 | Mitochondrial changes |
| GO:0060076 | excitatory synapse | 6/636 | 51/19594 | 0.00595 | 0.01969 | 6 | neurons0 | Synapse assembly |
| GO:0071682 | endocytic vesicle lumen | 4/636 | 23/19594 | 0.00596 | 0.01969 | 4 | neurons0 | Vesicle |
| GO:0030660 | Golgi-associated vesicle membrane | 6/636 | 52/19594 | 0.00655 | 0.02119 | 6 | neurons0 | Vesicle |
| GO:0098992 | neuronal dense core vesicle | 3/636 | 13/19594 | 0.00763 | 0.02444 | 3 | neurons0 | Vesicle |
| GO:0043209 | myelin sheath | 5/636 | 45/19594 | 0.01485 | 0.04419 | 5 | neurons0 | Enhanced myelination |
| GO:0060077 | inhibitory synapse | 3/636 | 19/19594 | 0.02239 | 0.06247 | 3 | neurons0 | Synapse assembly |
| GO:0099186 | structural constituent of postsynapse | 5/622 | 13/18410 | 4E-05 | 0.00083 | 5 | neurons0 | Synapse assembly |
| GO:0016209 | antioxidant activity | 11/622 | 85/18410 | 0.00013 | 0.00211 | 11 | neurons0 | Oxidative stress |
| GO:0015453 | oxidoreduction-driven active transmembrane transporter activity | 10/622 | 72/18410 | 0.00015 | 0.00224 | 10 | neurons0 | Oxidative stress |
| GO:0098918 | structural constituent of synapse | 5/622 | 19/18410 | 0.00034 | 0.00455 | 5 | neurons0 | Synapse assembly |
| GO:0000287 | magnesium ion binding | 18/622 | 222/18410 | 0.00056 | 0.00661 | 18 | neurons0 | Ion transport |
| GO:0016668 | oxidoreductase activity, acting on a sulfur group of donors, NAD(P) as acceptor | 4/622 | 13/18410 | 0.00072 | 0.00822 | 4 | neurons0 | Oxidative stress |
| GO:0008198 | ferrous iron binding | 5/622 | 26/18410 | 0.00158 | 0.01646 | 5 | neurons0 | Iron transport |
| GO:0043028 | cysteine-type endopeptidase regulator activity involved in apoptotic process | 6/622 | 40/18410 | 0.00209 | 0.02013 | 6 | neurons0 | Apoptosis |
| GO:0016684 | oxidoreductase activity, acting on peroxide as acceptor | 7/622 | 56/18410 | 0.00268 | 0.02387 | 7 | neurons0 | Oxidative stress |
| GO:0016667 | oxidoreductase activity, acting on a sulfur group of donors | 7/622 | 57/18410 | 0.00297 | 0.02588 | 7 | neurons0 | Oxidative stress |
| GO:0015662 | P-type ion transporter activity | 4/622 | 20/18410 | 0.00406 | 0.03071 | 4 | neurons0 | Ion transport |
| GO:0016903 | oxidoreductase activity, acting on the aldehyde or oxo group of donors | 6/622 | 46/18410 | 0.00431 | 0.03199 | 6 | neurons0 | Oxidative stress |

| | | | | | | | | |
|---|---|---|---|---|---|---|---|---|
| GO:0005507 | copper ion binding | 7/622 | 61/18410 | 0.00436 | 0.03207 | 7 | neurons0 | Ion transport |
| GO:0008199 | ferric iron binding | 3/622 | 11/18410 | 0.00517 | 0.0359 | 3 | neurons0 | Iron transport |
| GO:0043027 | cysteine-type endopeptidase inhibitor activity involved in apoptotic process | 4/622 | 22/18410 | 0.00581 | 0.03963 | 4 | neurons0 | Apoptosis |
| GO:0015485 | cholesterol binding | 6/622 | 50/18410 | 0.00653 | 0.04332 | 6 | neurons0 | Cholesterol metabolism |
| GO:0004601 | peroxidase activity | 6/622 | 52/18410 | 0.0079 | 0.05023 | 6 | neurons0 | Oxidative stress |
| GO:0016620 | oxidoreductase activity, acting on the aldehyde or oxo group of donors, NAD or NADP as acceptor | 5/622 | 38/18410 | 0.00865 | 0.05274 | 5 | neurons0 | Oxidative stress |
| GO:0048306 | calcium-dependent protein binding | 8/622 | 87/18410 | 0.00913 | 0.05501 | 8 | neurons0 | Calcium transport |
| GO:0016651 | oxidoreductase activity, acting on NAD(P)H | 8/622 | 88/18410 | 0.00976 | 0.05717 | 8 | neurons0 | Oxidative stress |
| GO:0016655 | oxidoreductase activity, acting on NAD(P)H, quinone or similar compound as acceptor | 6/622 | 57/18410 | 0.01223 | 0.06797 | 6 | neurons0 | Oxidative stress |
| hsa04260 | Cardiac muscle contraction | 13/358 | 87/8170 | 1E-04 | 9E-04 | 13 | neurons0 | Muscle contraction |
| hsa04728 | Dopaminergic synapse | 12/358 | 132/8170 | 0.01284 | 0.08102 | 12 | neurons0 | Dopamine metabolism |
| GO:0099172 | presynapse organization | 10/508 | 52/18800 | 0 | 1E-04 | 10 | neurons1 | Synapse assembly |
| GO:0035249 | synaptic transmission, glutamatergic | 13/508 | 94/18800 | 0 | 0.00013 | 13 | neurons1 | Glutamatergic synapse |
| GO:0099643 | signal release from synapse | 16/508 | 145/18800 | 0 | 0.00016 | 16 | neurons1 | Synapse assembly |
| GO:0031346 | positive regulation of cell projection organization | 26/508 | 341/18800 | 0 | 0.00017 | 26 | neurons1 | Cell projection organization |
| GO:0048041 | focal adhesion assembly | 12/508 | 83/18800 | 0 | 0.00017 | 12 | neurons1 | Cell adhesion |
| GO:0099175 | regulation of postsynapse organization | 12/508 | 86/18800 | 0 | 0.00025 | 12 | neurons1 | Synapse assembly |
| GO:0051965 | positive regulation of synapse assembly | 10/508 | 63/18800 | 1E-05 | 0.00047 | 10 | neurons1 | Synapse assembly |
| GO:0099003 | vesicle-mediated transport in synapse | 18/508 | 197/18800 | 1E-05 | 0.00047 | 18 | neurons1 | Vesicle |
| GO:0099504 | synaptic vesicle cycle | 17/508 | 183/18800 | 1E-05 | 0.00067 | 17 | neurons1 | Vesicle |
| GO:0060997 | dendritic spine morphogenesis | 9/508 | 58/18800 | 2E-05 | 0.0014 | 9 | neurons1 | Morphogenesis |
| GO:0001952 | regulation of cell-matrix adhesion | 13/508 | 123/18800 | 3E-05 | 0.00166 | 13 | neurons1 | Cell adhesion |
| GO:0002028 | regulation of sodium ion transport | 11/508 | 90/18800 | 3E-05 | 0.00174 | 11 | neurons1 | Ion transport |
| GO:0016079 | synaptic vesicle exocytosis | 12/508 | 107/18800 | 3E-05 | 0.0018 | 12 | neurons1 | Vesicle |
| GO:0032412 | regulation of ion transmembrane transporter activity | 20/508 | 263/18800 | 3E-05 | 0.00184 | 20 | neurons1 | Ion transport |
| GO:0050772 | positive regulation of axonogenesis | 10/508 | 77/18800 | 4E-05 | 0.00217 | 10 | neurons1 | Axonogenesis |

| | | | | | | | | |
|---|---|---|---|---|---|---|---|---|
| GO:0099560 | synaptic membrane adhesion | 6/508 | 25/18800 | 4E-05 | 0.00219 | 6 | neurons1 | Cell adhesion |
| GO:0050771 | negative regulation of axonogenesis | 9/508 | 64/18800 | 5E-05 | 0.00263 | 9 | neurons1 | Axonogenesis |
| GO:0021952 | central nervous system projection neuron axonogenesis | 6/508 | 26/18800 | 5E-05 | 0.00271 | 6 | neurons1 | Axonogenesis |
| GO:0090630 | activation of GTPase activity | 12/508 | 114/18800 | 6E-05 | 0.00279 | 12 | neurons1 | GTPase activity |
| GO:0007158 | neuron cell-cell adhesion | 5/508 | 17/18800 | 7E-05 | 0.00301 | 5 | neurons1 | Cell adhesion |
| GO:0007160 | cell-matrix adhesion | 18/508 | 235/18800 | 8E-05 | 0.0033 | 18 | neurons1 | Cell adhesion |
| GO:0099566 | regulation of postsynaptic cytosolic calcium ion concentration | 4/508 | 10/18800 | 1E-04 | 0.00418 | 4 | neurons1 | Calcium transport |
| GO:0071805 | potassium ion transmembrane transport | 17/508 | 219/18800 | 1E-04 | 0.00421 | 17 | neurons1 | Ion transport |
| GO:0097120 | receptor localization to synapse | 8/508 | 56/18800 | 0.00012 | 0.00502 | 8 | neurons1 | Synapse assembly |
| GO:0061001 | regulation of dendritic spine morphogenesis | 7/508 | 44/18800 | 0.00016 | 0.00626 | 7 | neurons1 | Morphogenesis |
| GO:0031345 | negative regulation of cell projection organization | 15/508 | 188/18800 | 0.00019 | 0.00704 | 15 | neurons1 | Cell projection organization |
| GO:0099558 | maintenance of synapse structure | 5/508 | 21/18800 | 2E-04 | 0.00743 | 5 | neurons1 | Synapse assembly |
| GO:0001738 | morphogenesis of a polarized epithelium | 10/508 | 94/18800 | 0.00023 | 0.00811 | 10 | neurons1 | Morphogenesis |
| GO:0022604 | regulation of cell morphogenesis | 20/508 | 305/18800 | 0.00025 | 0.00872 | 20 | neurons1 | Morphogenesis |
| GO:0099174 | regulation of presynapse organization | 6/508 | 34/18800 | 0.00027 | 0.0089 | 6 | neurons1 | Synapse assembly |
| GO:1905606 | regulation of presynapse assembly | 6/508 | 34/18800 | 0.00027 | 0.0089 | 6 | neurons1 | Synapse assembly |
| GO:0021955 | central nervous system neuron axonogenesis | 6/508 | 35/18800 | 0.00031 | 0.01014 | 6 | neurons1 | Axonogenesis |
| GO:0048814 | regulation of dendrite morphogenesis | 8/508 | 64/18800 | 0.00032 | 0.01014 | 8 | neurons1 | Morphogenesis |
| GO:0006813 | potassium ion transport | 17/508 | 243/18800 | 0.00035 | 0.01105 | 17 | neurons1 | Ion transport |
| GO:0010959 | regulation of metal ion transport | 23/508 | 403/18800 | 0.00066 | 0.01781 | 23 | neurons1 | Ion transport |
| GO:0007156 | homophilic cell adhesion via plasma membrane adhesion molecules | 13/508 | 168/18800 | 0.00066 | 0.01781 | 13 | neurons1 | Cell adhesion |
| GO:0010810 | regulation of cell-substrate adhesion | 15/508 | 217/18800 | 0.00086 | 0.02188 | 15 | neurons1 | Cell adhesion |
| GO:2000649 | regulation of sodium ion transmembrane transporter activity | 7/508 | 58/18800 | 0.00092 | 0.02288 | 7 | neurons1 | Ion transport |
| GO:0031589 | cell-substrate adhesion | 21/508 | 364/18800 | 0.00097 | 0.02376 | 21 | neurons1 | Cell adhesion |
| GO:0097553 | calcium ion transmembrane import into cytosol | 12/508 | 154/18800 | 0.00099 | 0.02397 | 12 | neurons1 | Calcium transport |
| GO:0006936 | muscle contraction | 20/508 | 349/18800 | 0.00138 | 0.03121 | 20 | neurons1 | Muscle contraction |

| | | | | | | | | |
|---|---|---|---|---|---|---|---|---|
| GO:0098703 | calcium ion import across plasma membrane | 5/508 | 32/18800 | 0.00156 | 0.03488 | 5 | neurons1 | Calcium transport |
| GO:1902656 | calcium ion import into cytosol | 5/508 | 33/18800 | 0.00179 | 0.03869 | 5 | neurons1 | Calcium transport |
| GO:0048755 | branching morphogenesis of a nerve | 3/508 | 10/18800 | 0.00204 | 0.04293 | 3 | neurons1 | Morphogenesis |
| GO:0035418 | protein localization to synapse | 7/508 | 67/18800 | 0.00216 | 0.04431 | 7 | neurons1 | Synapse assembly |
| GO:1902305 | regulation of sodium ion transmembrane transport | 7/508 | 67/18800 | 0.00216 | 0.04431 | 7 | neurons1 | Ion transport |
| GO:0070588 | calcium ion transmembrane transport | 18/508 | 314/18800 | 0.00233 | 0.04617 | 18 | neurons1 | Calcium transport |
| GO:0060402 | calcium ion transport into cytosol | 12/508 | 171/18800 | 0.00242 | 0.04731 | 12 | neurons1 | Calcium transport |
| GO:0045161 | neuronal ion channel clustering | 3/508 | 11/18800 | 0.00275 | 0.05195 | 3 | neurons1 | Ion transport |
| GO:0051966 | regulation of synaptic transmission, glutamatergic | 7/508 | 70/18800 | 0.00278 | 0.05209 | 7 | neurons1 | Glutamatergic synapse |
| GO:0006941 | striated muscle contraction | 12/508 | 178/18800 | 0.00336 | 0.05972 | 12 | neurons1 | Muscle contraction |
| GO:1901379 | regulation of potassium ion transmembrane transport | 8/508 | 95/18800 | 0.00419 | 0.07013 | 8 | neurons1 | Ion transport |
| GO:1904861 | excitatory synapse assembly | 4/508 | 25/18800 | 0.00425 | 0.07013 | 4 | neurons1 | Synapse assembly |
| GO:0001953 | negative regulation of cell-matrix adhesion | 5/508 | 40/18800 | 0.00426 | 0.07013 | 5 | neurons1 | Cell adhesion |
| GO:0060048 | cardiac muscle contraction | 10/508 | 138/18800 | 0.00431 | 0.07024 | 10 | neurons1 | Muscle contraction |
| GO:1900242 | regulation of synaptic vesicle endocytosis | 3/508 | 13/18800 | 0.00458 | 0.07338 | 3 | neurons1 | Vesicle |
| GO:0072028 | nephron morphogenesis | 7/508 | 77/18800 | 0.00476 | 0.07484 | 7 | neurons1 | Morphogenesis |
| GO:0060401 | cytosolic calcium ion transport | 12/508 | 190/18800 | 0.00564 | 0.08523 | 12 | neurons1 | Calcium transport |
| GO:0048512 | circadian behavior | 5/508 | 43/18800 | 0.00583 | 0.08665 | 5 | neurons1 | Circadian rhythm |
| GO:2000650 | negative regulation of sodium ion transmembrane transporter activity | 3/508 | 15/18800 | 7 | 0.09783 | 3 | neurons1 | Ion transport |
| GO:0005891 | voltage-gated calcium channel complex | 8/524 | 44/19594 | 2E-05 | 0.00021 | 8 | neurons1 | Calcium transport |
| GO:0098685 | Schaffer collateral - CA1 synapse | 9/524 | 72/19594 | 0.00012 | 0.00123 | 9 | neurons1 | Synapse assembly |
| GO:0060076 | excitatory synapse | 7/524 | 51/19594 | 0.00039 | 0.00349 | 7 | neurons1 | Synapse assembly |
| GO:0098688 | parallel fiber to Purkinje cell synapse | 3/524 | 15/19594 | 0.00681 | 0.0439 | 3 | neurons1 | Synapse assembly |
| GO:0098686 | hippocampal mossy fiber to CA3 synapse | 4/524 | 34/19594 | 0.01243 | 0.06912 | 4 | neurons1 | Synapse assembly |
| GO:0022824 | transmitter-gated ion channel activity | 10/507 | 60/18410 | 1E-05 | 0.00021 | 10 | neurons1 | Ion transport |
| GO:0008331 | high voltage-gated calcium channel activity | 5/507 | 11/18410 | 1E-05 | 0.00024 | 5 | neurons1 | Calcium transport |

| | | | | | | | | |
|---|---|---|---|---|---|---|---|---|
| GO:0015276 | ligand-gated ion channel activity | 15/507 | 145/18410 | 1E-05 | 0.00035 | 15 | neurons1 | Ion transport |
| GO:0005244 | voltage-gated ion channel activity | 18/507 | 201/18410 | 1E-05 | 0.00035 | 18 | neurons1 | Ion transport |
| GO:0005230 | extracellular ligand-gated ion channel activity | 10/507 | 73/18410 | 3E-05 | 0.00084 | 10 | neurons1 | Ion transport |
| GO:0031267 | small GTPase binding | 20/507 | 267/18410 | 5E-05 | 0.00145 | 20 | neurons1 | GTPase activity |
| GO:0051020 | GTPase binding | 20/507 | 298/18410 | 0.00024 | 0.00473 | 20 | neurons1 | GTPase activity |
| GO:1904315 | transmitter-gated ion channel activity involved in regulation of postsynaptic membrane potential | 7/507 | 46/18410 | 0.00024 | 0.00473 | 7 | neurons1 | Ion transport |
| GO:0015085 | calcium ion transmembrane transporter activity | 12/507 | 135/18410 | 0.00036 | 0.00607 | 12 | neurons1 | Calcium transport |
| GO:0046873 | metal ion transmembrane transporter activity | 25/507 | 428/18410 | 0.00036 | 0.00607 | 25 | neurons1 | Ion transport |
| GO:0005262 | calcium channel activity | 11/507 | 119/18410 | 0.00045 | 0.00729 | 11 | neurons1 | Calcium transport |
| GO:0098632 | cell-cell adhesion mediator activity | 7/507 | 54/18410 | 0.00067 | 0.01027 | 7 | neurons1 | Cell adhesion |
| GO:0015079 | potassium ion transmembrane transporter activity | 12/507 | 154/18410 | 0.00117 | 0.01634 | 12 | neurons1 | Ion transport |
| GO:0098631 | cell adhesion mediator activity | 7/507 | 64/18410 | 0.00185 | 0.02407 | 7 | neurons1 | Cell adhesion |
| GO:0005245 | voltage-gated calcium channel activity | 5/507 | 45/18410 | 0.00766 | 0.06832 | 5 | neurons1 | Calcium transport |
| hsa04726 | Serotonergic synapse | 13/214 | 115/8170 | 1E-05 | 0.00022 | 13 | neurons1 | Synapse assembly |
| hsa04725 | Cholinergic synapse | 12/214 | 113/8170 | 4E-05 | 0.00064 | 12 | neurons1 | Synapse assembly |
| hsa04270 | Vascular smooth muscle contraction | 13/214 | 134/8170 | 5E-05 | 0.00072 | 13 | neurons1 | Muscle contraction |
| hsa04020 | Calcium signaling pathway | 18/214 | 240/8170 | 5E-05 | 0.00081 | 18 | neurons1 | Calcium transport |
| hsa04727 | GABAergic synapse | 9/214 | 89/8170 | 0.00051 | 0.00423 | 9 | neurons1 | Synapse assembly |
| hsa04670 | Leukocyte transendothelial migration | 8/214 | 114/8170 | 0.00995 | 0.04786 | 8 | neurons1 | Leukocytes |
| hsa04961 | Endocrine and other factor-regulated calcium reabsorption | 5/214 | 53/8170 | 0.01217 | 0.0534 | 5 | neurons1 | Calcium transport |
| hsa04510 | Focal adhesion | 11/214 | 201/8170 | 0.01645 | 0.0623 | 11 | neurons1 | Cell adhesion |
| GO:0043547 | positive regulation of GTPase activity | 28/713 | 272/18800 | 0 | 0.00017 | 28 | neurons2 | GTPase activity |
| GO:0099560 | synaptic membrane adhesion | 8/713 | 25/18800 | 0 | 0.00023 | 8 | neurons2 | Cell adhesion |
| GO:0007157 | heterophilic cell-cell adhesion via plasma membrane cell adhesion molecules | 11/713 | 51/18800 | 0 | 0.00023 | 11 | neurons2 | Cell adhesion |
| GO:0051965 | positive regulation of synapse assembly | 12/713 | 63/18800 | 0 | 0.00031 | 12 | neurons2 | Synapse assembly |

| | | | | | | | | |
|---|---|---|---|---|---|---|---|---|
| GO:0099175 | regulation of postsynapse organization | 14/713 | 86/18800 | 0 | 0.00032 | 14 | neurons2 | Synapse assembly |
| GO:0050772 | positive regulation of axonogenesis | 13/713 | 77/18800 | 1E-05 | 0.00046 | 13 | neurons2 | Axonogenesis |
| GO:0097120 | receptor localization to synapse | 11/713 | 56/18800 | 1E-05 | 0.00052 | 11 | neurons2 | Synapse assembly |
| GO:0031589 | cell-substrate adhesion | 32/713 | 364/18800 | 1E-05 | 0.00072 | 32 | neurons2 | Cell adhesion |
| GO:0070588 | calcium ion transmembrane transport | 29/713 | 314/18800 | 1E-05 | 0.00072 | 29 | neurons2 | Calcium transport |
| GO:0051966 | regulation of synaptic transmission, glutamatergic | 12/713 | 70/18800 | 1E-05 | 0.00082 | 12 | neurons2 | Glutamatergic synapse |
| GO:0048041 | focal adhesion assembly | 13/713 | 83/18800 | 1E-05 | 0.00094 | 13 | neurons2 | Cell adhesion |
| GO:0048814 | regulation of dendrite morphogenesis | 11/713 | 64/18800 | 3E-05 | 0.00151 | 11 | neurons2 | Morphogenesis |
| GO:0006816 | calcium ion transport | 34/713 | 424/18800 | 4E-05 | 0.00194 | 34 | neurons2 | Calcium transport |
| GO:0021952 | central nervous system projection neuron axonogenesis | 7/713 | 26/18800 | 4E-05 | 0.00206 | 7 | neurons2 | Axonogenesis |
| GO:0001952 | regulation of cell-matrix adhesion | 15/713 | 123/18800 | 7E-05 | 0.00301 | 15 | neurons2 | Cell adhesion |
| GO:0032412 | regulation of ion transmembrane transporter activity | 24/713 | 263/18800 | 7E-05 | 0.00312 | 24 | neurons2 | Ion transport |
| GO:0097553 | calcium ion transmembrane import into cytosol | 17/713 | 154/18800 | 8E-05 | 0.00345 | 17 | neurons2 | Calcium transport |
| GO:0010810 | regulation of cell-substrate adhesion | 21/713 | 217/18800 | 8E-05 | 0.00358 | 21 | neurons2 | Cell adhesion |
| GO:0031345 | negative regulation of cell projection organization | 19/713 | 188/18800 | 1E-04 | 0.00416 | 19 | neurons2 | Cell projection organization |
| GO:0022604 | regulation of cell morphogenesis | 26/713 | 305/18800 | 0.00011 | 0.00423 | 26 | neurons2 | Morphogenesis |
| GO:0060401 | cytosolic calcium ion transport | 19/713 | 190/18800 | 0.00012 | 0.00449 | 19 | neurons2 | Calcium transport |
| GO:0051893 | regulation of focal adhesion assembly | 10/713 | 63/18800 | 0.00012 | 0.00454 | 10 | neurons2 | Cell adhesion |
| GO:0099643 | signal release from synapse | 16/713 | 145/18800 | 0.00013 | 0.00462 | 16 | neurons2 | Synapse assembly |
| GO:1990806 | ligand-gated ion channel signaling pathway | 6/713 | 22/18800 | 0.00013 | 0.00464 | 6 | neurons2 | Ion transport |
| GO:0098703 | calcium ion import across plasma membrane | 7/713 | 32/18800 | 0.00016 | 0.00542 | 7 | neurons2 | Calcium transport |
| GO:0070509 | calcium ion import | 12/713 | 91/18800 | 0.00016 | 0.00542 | 12 | neurons2 | Calcium transport |
| GO:1902656 | calcium ion import into cytosol | 7/713 | 33/18800 | 2E-04 | 0.00635 | 7 | neurons2 | Calcium transport |
| GO:0035418 | protein localization to synapse | 10/713 | 67/18800 | 2E-04 | 0.00647 | 10 | neurons2 | Synapse assembly |
| GO:0055074 | calcium ion homeostasis | 34/713 | 468/18800 | 0.00024 | 0.00731 | 34 | neurons2 | Calcium transport |
| GO:1901379 | regulation of potassium ion transmembrane transport | 12/713 | 95/18800 | 0.00025 | 0.00743 | 12 | neurons2 | Ion transport |

| | | | | | | | | |
|---|---|---|---|---|---|---|---|---|
| GO:0007160 | cell-matrix adhesion | 21/713 | 235/18800 | 0.00025 | 0.00761 | 21 | neurons2 | Cell adhesion |
| GO:0019722 | calcium-mediated signaling | 19/713 | 202/18800 | 0.00026 | 0.00766 | 19 | neurons2 | Calcium transport |
| GO:0071805 | potassium ion transmembrane transport | 20/713 | 219/18800 | 0.00027 | 0.00779 | 20 | neurons2 | Ion transport |
| GO:0060402 | calcium ion transport into cytosol | 17/713 | 171/18800 | 0.00028 | 0.00809 | 17 | neurons2 | Calcium transport |
| GO:0021955 | central nervous system neuron axonogenesis | 7/713 | 35/18800 | 0.00029 | 0.00835 | 7 | neurons2 | Axonogenesis |
| GO:0006874 | cellular calcium ion homeostasis | 33/713 | 456/18800 | 0.00031 | 0.00881 | 33 | neurons2 | Calcium transport |
| GO:0099566 | regulation of postsynaptic cytosolic calcium ion concentration | 4/713 | 10/18800 | 0.00036 | 0.00975 | 4 | neurons2 | Calcium transport |
| GO:0010959 | regulation of metal ion transport | 30/713 | 403/18800 | 0.00036 | 0.00979 | 30 | neurons2 | Ion transport |
| GO:0090630 | activation of GTPase activity | 13/713 | 114/18800 | 0.00038 | 0.01033 | 13 | neurons2 | GTPase activity |
| GO:0051480 | regulation of cytosolic calcium ion concentration | 27/713 | 356/18800 | 0.00053 | 0.01337 | 27 | neurons2 | Calcium transport |
| GO:0002028 | regulation of sodium ion transport | 11/713 | 90/18800 | 0.00059 | 0.01455 | 11 | neurons2 | Ion transport |
| GO:0099504 | synaptic vesicle cycle | 17/713 | 183/18800 | 0.00062 | 0.01507 | 17 | neurons2 | Vesicle |
| GO:0051932 | synaptic transmission, GABAergic | 8/713 | 51/18800 | 0.00062 | 0.01507 | 8 | neurons2 | GABAergic synapse |
| GO:0060560 | developmental growth involved in morphogenesis | 20/713 | 234/18800 | 0.00063 | 0.01515 | 20 | neurons2 | Morphogenesis |
| GO:0007162 | negative regulation of cell adhesion | 24/713 | 305/18800 | 0.00063 | 0.01515 | 24 | neurons2 | Cell adhesion |
| GO:0032414 | positive regulation of ion transmembrane transporter activity | 12/713 | 106/18800 | 0.00068 | 0.01597 | 12 | neurons2 | Ion transport |
| GO:0010812 | negative regulation of cell-substrate adhesion | 9/713 | 66/18800 | 0.00083 | 0.01852 | 9 | neurons2 | Cell adhesion |
| GO:0043266 | regulation of potassium ion transport | 12/713 | 110/18800 | 0.00095 | 0.02076 | 12 | neurons2 | Ion transport |
| GO:0006813 | potassium ion transport | 20/713 | 243/18800 | 1 | 0.0216 | 20 | neurons2 | Ion transport |
| GO:0051592 | response to calcium ion | 14/713 | 144/18800 | 0.00117 | 0.02385 | 14 | neurons2 | Calcium transport |
| GO:0034767 | positive regulation of ion transmembrane transport | 15/713 | 163/18800 | 0.00139 | 0.02737 | 15 | neurons2 | Ion transport |
| GO:0099003 | vesicle-mediated transport in synapse | 17/713 | 197/18800 | 0.0014 | 0.02751 | 17 | neurons2 | Vesicle |
| GO:2000649 | regulation of sodium ion transmembrane transporter activity | 8/713 | 58/18800 | 0.00148 | 0.02883 | 8 | neurons2 | Ion transport |
| GO:0007204 | positive regulation of cytosolic calcium ion concentration | 24/713 | 325/18800 | 0.00151 | 0.0292 | 24 | neurons2 | Calcium transport |
| GO:0097091 | synaptic vesicle clustering | 4/713 | 14/18800 | 0.00151 | 0.0292 | 4 | neurons2 | Vesicle |
| GO:0032228 | regulation of synaptic transmission, GABAergic | 6/713 | 34/18800 | 0.00158 | 0.02939 | 6 | neurons2 | GABAergic synapse |

| | | | | | | | | |
|---|---|---|---|---|---|---|---|---|
| GO:0099174 | regulation of presynapse organization | 6/713 | 34/18800 | 0.00158 | 0.02939 | 6 | neurons2 | Synapse assembly |
| GO:1905606 | regulation of presynapse assembly | 6/713 | 34/18800 | 0.00158 | 0.02939 | 6 | neurons2 | Synapse assembly |
| GO:0048854 | brain morphogenesis | 6/713 | 36/18800 | 0.00215 | 0.03715 | 6 | neurons2 | Morphogenesis |
| GO:0003148 | outflow tract septum morphogenesis | 5/713 | 25/18800 | 0.00219 | 0.03722 | 5 | neurons2 | Morphogenesis |
| GO:1904861 | excitatory synapse assembly | 5/713 | 25/18800 | 0.00219 | 0.03722 | 5 | neurons2 | Synapse assembly |
| GO:0034446 | substrate adhesion-dependent cell spreading | 11/713 | 109/18800 | 0.00285 | 0.04523 | 11 | neurons2 | Cell adhesion |
| GO:0035725 | sodium ion transmembrane transport | 15/713 | 177/18800 | 0.0031 | 0.04845 | 15 | neurons2 | Ion transport |
| GO:0006814 | sodium ion transport | 19/713 | 249/18800 | 0.00313 | 0.04845 | 19 | neurons2 | Ion transport |
| GO:0006941 | striated muscle contraction | 15/713 | 178/18800 | 0.00327 | 0.0493 | 15 | neurons2 | Muscle contraction |
| GO:0001953 | negative regulation of cell-matrix adhesion | 6/713 | 40/18800 | 0.00372 | 0.05433 | 6 | neurons2 | Cell adhesion |
| GO:1902305 | regulation of sodium ion transmembrane transport | 8/713 | 67/18800 | 0.00373 | 0.05433 | 8 | neurons2 | Ion transport |
| GO:0060412 | ventricular septum morphogenesis | 6/713 | 42/18800 | 0.00477 | 0.06743 | 6 | neurons2 | Morphogenesis |
| GO:2000300 | regulation of synaptic vesicle exocytosis | 7/713 | 56/18800 | 0.00506 | 0.06916 | 7 | neurons2 | Vesicle |
| GO:0016339 | calcium-dependent cell-cell adhesion via plasma membrane cell adhesion molecules | 6/713 | 43/18800 | 0.00537 | 0.07139 | 6 | neurons2 | Cell adhesion |
| GO:0021575 | hindbrain morphogenesis | 6/713 | 43/18800 | 0.00537 | 0.07139 | 6 | neurons2 | Morphogenesis |
| GO:0051058 | negative regulation of small GTPase mediated signal transduction | 7/713 | 57/18800 | 0.00558 | 0.07394 | 7 | neurons2 | GTPase activity |
| GO:1901381 | positive regulation of potassium ion transmembrane transport | 6/713 | 44/18800 | 0.00602 | 0.07835 | 6 | neurons2 | Ion transport |
| GO:0060997 | dendritic spine morphogenesis | 7/713 | 58/18800 | 0.00614 | 0.07917 | 7 | neurons2 | Morphogenesis |
| GO:0045161 | neuronal ion channel clustering | 3/713 | 11/18800 | 0.00713 | 0.0872 | 3 | neurons2 | Ion transport |
| GO:0099558 | maintenance of synapse structure | 4/713 | 21/18800 | 0.00734 | 0.08813 | 4 | neurons2 | Synapse assembly |
| GO:0043270 | positive regulation of ion transport | 19/713 | 273/18800 | 0.00829 | 0.09468 | 19 | neurons2 | Ion transport |
| GO:0050850 | positive regulation of calcium-mediated signaling | 5/713 | 34/18800 | 0.00865 | 0.09737 | 5 | neurons2 | Calcium transport |
| GO:0098686 | hippocampal mossy fiber to CA3 synapse | 8/739 | 34/19594 | 3E-05 | 0.00032 | 8 | neurons2 | Synapse assembly |
| GO:0005891 | voltage-gated calcium channel complex | 9/739 | 44/19594 | 3E-05 | 0.00034 | 9 | neurons2 | Calcium transport |
| GO:1990454 | L-type voltage-gated calcium channel complex | 4/739 | 11/19594 | 0.00054 | 0.00423 | 4 | neurons2 | Calcium transport |

| | | | | | | | | |
|---|---|---|---|---|---|---|---|---|
| GO:0060077 | inhibitory synapse | 5/739 | 19/19594 | 0.00056 | 0.00438 | 5 | neurons2 | Synapse assembly |
| GO:0032280 | symmetric synapse | 3/739 | 10/19594 | 0.00526 | 0.03038 | 3 | neurons2 | Synapse assembly |
| GO:0098688 | parallel fiber to Purkinje cell synapse | 3/739 | 15/19594 | 0.01732 | 0.08642 | 3 | neurons2 | Synapse assembly |
| GO:0022824 | transmitter-gated ion channel activity | 11/708 | 60/18410 | 2E-05 | 0.00056 | 11 | neurons2 | Ion transport |
| GO:0046873 | metal ion transmembrane transporter activity | 35/708 | 428/18410 | 2E-05 | 0.00079 | 35 | neurons2 | Ion transport |
| GO:0008331 | high voltage-gated calcium channel activity | 5/708 | 11/18410 | 3E-05 | 0.00096 | 5 | neurons2 | Calcium transport |
| GO:0005244 | voltage-gated ion channel activity | 21/708 | 201/18410 | 3E-05 | 0.00096 | 21 | neurons2 | Ion transport |
| GO:0005230 | extracellular ligand-gated ion channel activity | 11/708 | 73/18410 | 1E-04 | 0.00246 | 11 | neurons2 | Ion transport |
| GO:0005245 | voltage-gated calcium channel activity | 8/708 | 45/18410 | 0.00028 | 0.00636 | 8 | neurons2 | Calcium transport |
| GO:0015276 | ligand-gated ion channel activity | 15/708 | 145/18410 | 0.00048 | 0.00907 | 15 | neurons2 | Ion transport |
| GO:0098632 | cell-cell adhesion mediator activity | 8/708 | 54/18410 | 1 | 0.01587 | 8 | neurons2 | Cell adhesion |
| GO:0098631 | cell adhesion mediator activity | 8/708 | 64/18410 | 0.00304 | 0.03737 | 8 | neurons2 | Cell adhesion |
| GO:0098918 | structural constituent of synapse | 4/708 | 19/18410 | 0.0053 | 0.05748 | 4 | neurons2 | Synapse assembly |
| GO:0005432 | calcium:sodium antiporter activity | 3/708 | 10/18410 | 0.00555 | 0.05748 | 3 | neurons2 | Calcium transport |
| GO:0015368 | calcium:cation antiporter activity | 3/708 | 11/18410 | 0.00741 | 0.0672 | 3 | neurons2 | Calcium transport |
| GO:1904315 | transmitter-gated ion channel activity involved in regulation of postsynaptic membrane potential | 6/708 | 46/18410 | 8 | 0.07078 | 6 | neurons2 | Ion transport |
| GO:0022851 | GABA-gated chloride ion channel activity | 3/708 | 13/18410 | 0.01214 | 0.09165 | 3 | neurons2 | Ion transport |
| hsa04727 | GABAergic synapse | 13/267 | 89/8170 | 1E-05 | 0.00011 | 13 | neurons2 | Synapse assembly |
| hsa04020 | Calcium signaling pathway | 22/267 | 240/8170 | 1E-05 | 0.00015 | 22 | neurons2 | Calcium transport |
| hsa04725 | Cholinergic synapse | 14/267 | 113/8170 | 2E-05 | 0.00021 | 14 | neurons2 | Synapse assembly |
| hsa04726 | Serotonergic synapse | 13/267 | 115/8170 | 9E-05 | 0.00079 | 13 | neurons2 | Synapse assembly |
| hsa04514 | Cell adhesion molecules | 15/267 | 157/8170 | 0.00018 | 0.00125 | 15 | neurons2 | Cell adhesion |
| hsa04270 | Vascular smooth muscle contraction | 13/267 | 134/8170 | 0.00042 | 0.00255 | 13 | neurons2 | Muscle contraction |
| hsa04260 | Cardiac muscle contraction | 8/267 | 87/8170 | 0.00736 | 0.0318 | 8 | neurons2 | Muscle contraction |
| hsa04660 | T cell receptor signaling pathway | 8/267 | 104/8170 | 0.02024 | 0.07168 | 8 | neurons2 | T cell activity |

| | | | | | | | | |
|---|---|---|---|---|---|---|---|---|
| hsa04961 | Endocrine and other factor-regulated calcium reabsorption | 5/267 | 53/8170 | 0.02875 | 0.0894 | 5 | neurons2 | Calcium transport |
| hsa04510 | Focal adhesion | 12/267 | 201/8170 | 0.03188 | 0.09677 | 12 | neurons2 | Cell adhesion |
| GO:0043270 | positive regulation of ion transport | 12/151 | 273/18800 | 0 | 0.00016 | 12 | neurons3 | Ion transport |
| GO:0006874 | cellular calcium ion homeostasis | 15/151 | 456/18800 | 0 | 0.00027 | 15 | neurons3 | Calcium transport |
| GO:0006813 | potassium ion transport | 11/151 | 243/18800 | 0 | 0.00027 | 11 | neurons3 | Ion transport |
| GO:0010959 | regulation of metal ion transport | 14/151 | 403/18800 | 0 | 0.00028 | 14 | neurons3 | Ion transport |
| GO:0099587 | inorganic ion import across plasma membrane | 8/151 | 118/18800 | 0 | 0.00028 | 8 | neurons3 | Ion transport |
| GO:0099509 | regulation of presynaptic cytosolic calcium ion concentration | 4/151 | 15/18800 | 1E-05 | 0.00028 | 4 | neurons3 | Calcium transport |
| GO:0097120 | receptor localization to synapse | 6/151 | 56/18800 | 1E-05 | 3E-04 | 6 | neurons3 | Synapse assembly |
| GO:0055074 | calcium ion homeostasis | 15/151 | 468/18800 | 1E-05 | 3E-04 | 15 | neurons3 | Calcium transport |
| GO:1904862 | inhibitory synapse assembly | 4/151 | 16/18800 | 1E-05 | 0.00032 | 4 | neurons3 | Synapse assembly |
| GO:0034765 | regulation of ion transmembrane transport | 15/151 | 476/18800 | 1E-05 | 0.00032 | 15 | neurons3 | Ion transport |
| GO:0006816 | calcium ion transport | 14/151 | 424/18800 | 1E-05 | 0.00035 | 14 | neurons3 | Calcium transport |
| GO:0048488 | synaptic vesicle endocytosis | 6/151 | 61/18800 | 1E-05 | 0.00037 | 6 | neurons3 | Vesicle |
| GO:0062237 | protein localization to postsynapse | 5/151 | 36/18800 | 1E-05 | 0.00037 | 5 | neurons3 | Synapse assembly |
| GO:0007409 | axonogenesis | 14/151 | 430/18800 | 1E-05 | 0.00038 | 14 | neurons3 | Axonogenesis |
| GO:0071805 | potassium ion transmembrane transport | 10/151 | 219/18800 | 1E-05 | 0.00039 | 10 | neurons3 | Ion transport |
| GO:0051402 | neuron apoptotic process | 10/151 | 241/18800 | 3E-05 | 0.00082 | 10 | neurons3 | Apoptosis |
| GO:0036465 | synaptic vesicle recycling | 6/151 | 73/18800 | 3E-05 | 0.00083 | 6 | neurons3 | Vesicle |
| GO:0051480 | regulation of cytosolic calcium ion concentration | 12/151 | 356/18800 | 3E-05 | 0.00096 | 12 | neurons3 | Calcium transport |
| GO:0097553 | calcium ion transmembrane import into cytosol | 8/151 | 154/18800 | 3E-05 | 0.00105 | 8 | neurons3 | Calcium transport |
| GO:0070839 | metal ion export | 5/151 | 49/18800 | 4E-05 | 0.0013 | 5 | neurons3 | Ion transport |
| GO:0010882 | regulation of cardiac muscle contraction by calcium ion signaling | 4/151 | 26/18800 | 5E-05 | 0.00145 | 4 | neurons3 | Muscle contraction |
| GO:0099173 | postsynapse organization | 8/151 | 163/18800 | 5E-05 | 0.00145 | 8 | neurons3 | Synapse assembly |
| GO:0051932 | synaptic transmission, GABAergic | 5/151 | 51/18800 | 5E-05 | 0.0015 | 5 | neurons3 | GABAergic synapse |
| GO:1990034 | calcium ion export across plasma membrane | 3/151 | 10/18800 | 6E-05 | 0.00159 | 3 | neurons3 | Calcium transport |
| GO:0007263 | nitric oxide mediated signal transduction | 4/151 | 27/18800 | 6E-05 | 0.00161 | 4 | neurons3 | Oxidative stress |

| | | | | | | | | |
|---|---|---|---|---|---|---|---|---|
| GO:0060402 | calcium ion transport into cytosol | 8/151 | 171/18800 | 7E-05 | 0.00192 | 8 | neurons3 | Calcium transport |
| GO:0060048 | cardiac muscle contraction | 7/151 | 138/18800 | 0.00013 | 0.00297 | 7 | neurons3 | Muscle contraction |
| GO:0043524 | negative regulation of neuron apoptotic process | 7/151 | 145/18800 | 0.00017 | 0.00357 | 7 | neurons3 | Apoptosis |
| GO:0099643 | signal release from synapse | 7/151 | 145/18800 | 0.00017 | 0.00357 | 7 | neurons3 | Synapse assembly |
| GO:0016079 | synaptic vesicle exocytosis | 6/151 | 107/18800 | 0.00023 | 0.00429 | 6 | neurons3 | Vesicle |
| GO:0019722 | calcium-mediated signaling | 8/151 | 202/18800 | 0.00023 | 0.00437 | 8 | neurons3 | Calcium transport |
| GO:0061684 | chaperone-mediated autophagy | 3/151 | 16/18800 | 0.00026 | 0.00469 | 3 | neurons3 | Autophagy |
| GO:0043523 | regulation of neuron apoptotic process | 8/151 | 207/18800 | 0.00027 | 0.00479 | 8 | neurons3 | Apoptosis |
| GO:0032412 | regulation of ion transmembrane transporter activity | 9/151 | 263/18800 | 0.00028 | 0.00483 | 9 | neurons3 | Ion transport |
| GO:0006458 | 'de novo' protein folding | 4/151 | 41/18800 | 0.00032 | 0.00533 | 4 | neurons3 | Unfolded Protein Response |
| GO:0046034 | ATP metabolic process | 9/151 | 273/18800 | 0.00037 | 0.00586 | 9 | neurons3 | Energy production |
| GO:0055117 | regulation of cardiac muscle contraction | 5/151 | 77/18800 | 0.00039 | 0.00613 | 5 | neurons3 | Muscle contraction |
| GO:0051928 | positive regulation of calcium ion transport | 6/151 | 119/18800 | 4E-04 | 0.00628 | 6 | neurons3 | Calcium transport |
| GO:0006937 | regulation of muscle contraction | 7/151 | 170/18800 | 0.00045 | 0.00688 | 7 | neurons3 | Muscle contraction |
| GO:0006936 | muscle contraction | 10/151 | 349/18800 | 0.00053 | 0.00791 | 10 | neurons3 | Muscle contraction |
| GO:0006941 | striated muscle contraction | 7/151 | 178/18800 | 6E-04 | 0.00859 | 7 | neurons3 | Muscle contraction |
| GO:0002028 | regulation of sodium ion transport | 5/151 | 90/18800 | 0.00079 | 0.01065 | 5 | neurons3 | Ion transport |
| GO:0060074 | synapse maturation | 3/151 | 25/18800 | 0.00103 | 0.01293 | 3 | neurons3 | Synapse assembly |
| GO:0006942 | regulation of striated muscle contraction | 5/151 | 96/18800 | 0.00106 | 0.01312 | 5 | neurons3 | Muscle contraction |
| GO:0051592 | response to calcium ion | 6/151 | 144/18800 | 0.00109 | 0.01334 | 6 | neurons3 | Calcium transport |
| GO:0048499 | synaptic vesicle membrane organization | 3/151 | 26/18800 | 0.00115 | 0.01383 | 3 | neurons3 | Vesicle |
| GO:1901018 | positive regulation of potassium ion transmembrane transporter activity | 3/151 | 27/18800 | 0.00129 | 0.01482 | 3 | neurons3 | Ion transport |
| GO:0010522 | regulation of calcium ion transport into cytosol | 5/151 | 103/18800 | 0.00146 | 0.01621 | 5 | neurons3 | Calcium transport |
| GO:0050770 | regulation of axonogenesis | 6/151 | 154/18800 | 0.00154 | 0.01691 | 6 | neurons3 | Axonogenesis |
| GO:0006906 | vesicle fusion | 5/151 | 105/18800 | 0.00159 | 0.01725 | 5 | neurons3 | Vesicle |
| GO:0006457 | protein folding | 7/151 | 212/18800 | 0.00165 | 0.0177 | 7 | neurons3 | Unfolded Protein Response |

| | | | | | | | | |
|---|---|---|---|---|---|---|---|---|
| GO:0032414 | positive regulation of ion transmembrane transporter activity | 5/151 | 106/18800 | 0.00165 | 0.0177 | 5 | neurons3 | Ion transport |
| GO:0035966 | response to topologically incorrect protein | 6/151 | 160/18800 | 0.00187 | 0.01951 | 6 | neurons3 | Unfolded Protein Response |
| GO:0034767 | positive regulation of ion transmembrane transport | 6/151 | 163/18800 | 0.00205 | 0.02064 | 6 | neurons3 | Ion transport |
| GO:1902305 | regulation of sodium ion transmembrane transport | 4/151 | 67/18800 | 0.00207 | 0.02074 | 4 | neurons3 | Ion transport |
| GO:0051085 | chaperone cofactor-dependent protein refolding | 3/151 | 32/18800 | 0.00212 | 0.02077 | 3 | neurons3 | Unfolded Protein Response |
| GO:0098703 | calcium ion import across plasma membrane | 3/151 | 32/18800 | 0.00212 | 0.02077 | 3 | neurons3 | Calcium transport |
| GO:0010765 | positive regulation of sodium ion transport | 3/151 | 33/18800 | 0.00232 | 0.02202 | 3 | neurons3 | Ion transport |
| GO:1902656 | calcium ion import into cytosol | 3/151 | 33/18800 | 0.00232 | 0.02202 | 3 | neurons3 | Calcium transport |
| GO:0061077 | chaperone-mediated protein folding | 4/151 | 70/18800 | 0.00243 | 0.02279 | 4 | neurons3 | Unfolded Protein Response |
| GO:1904427 | positive regulation of calcium ion transmembrane transport | 4/151 | 71/18800 | 0.00256 | 0.02356 | 4 | neurons3 | Calcium transport |
| GO:0010749 | regulation of nitric oxide mediated signal transduction | 2/151 | 10/18800 | 0.00276 | 0.02445 | 2 | neurons3 | Oxidative stress |
| GO:1905383 | protein localization to presynapse | 2/151 | 10/18800 | 0.00276 | 0.02445 | 2 | neurons3 | Synapse assembly |
| GO:0050848 | regulation of calcium-mediated signaling | 4/151 | 75/18800 | 0.00312 | 0.02715 | 4 | neurons3 | Calcium transport |
| GO:0014046 | dopamine secretion | 3/151 | 37/18800 | 0.00323 | 0.0276 | 3 | neurons3 | Dopamine metabolism |
| GO:0014059 | regulation of dopamine secretion | 3/151 | 37/18800 | 0.00323 | 0.0276 | 3 | neurons3 | Dopamine metabolism |
| GO:0051084 | 'de novo' post-translational protein folding | 3/151 | 37/18800 | 0.00323 | 0.0276 | 3 | neurons3 | Unfolded Protein Response |
| GO:1901021 | positive regulation of calcium ion transmembrane transporter activity | 3/151 | 37/18800 | 0.00323 | 0.0276 | 3 | neurons3 | Calcium transport |
| GO:0034975 | protein folding in endoplasmic reticulum | 2/151 | 11/18800 | 0.00336 | 0.02785 | 2 | neurons3 | Unfolded Protein Response |
| GO:0050772 | positive regulation of axonogenesis | 4/151 | 77/18800 | 0.00344 | 0.02828 | 4 | neurons3 | Axonogenesis |
| GO:0007212 | dopamine receptor signaling pathway | 3/151 | 40/18800 | 0.00404 | 0.03238 | 3 | neurons3 | Dopamine metabolism |
| GO:0031345 | negative regulation of cell projection organization | 6/151 | 188/18800 | 0.00416 | 0.03302 | 6 | neurons3 | Cell projection organization |
| GO:0051924 | regulation of calcium ion transport | 7/151 | 251/18800 | 0.00422 | 0.03339 | 7 | neurons3 | Calcium transport |
| GO:0070296 | sarcoplasmic reticulum calcium ion transport | 3/151 | 42/18800 | 0.00464 | 0.03591 | 3 | neurons3 | Calcium transport |
| GO:0005513 | detection of calcium ion | 2/151 | 13/18800 | 0.00472 | 0.03597 | 2 | neurons3 | Calcium transport |
| GO:0007204 | positive regulation of cytosolic calcium ion concentration | 8/151 | 325/18800 | 0.00481 | 0.03658 | 8 | neurons3 | Calcium transport |

| | | | | | | | | |
|---|---|---|---|---|---|---|---|---|
| GO:1901381 | positive regulation of potassium ion transmembrane transport | 3/151 | 44/18800 | 0.00529 | 0.03948 | 3 | neurons3 | Ion transport |
| GO:0006986 | response to unfolded protein | 5/151 | 139/18800 | 0.00532 | 0.03957 | 5 | neurons3 | Unfolded Protein Response |
| GO:0070509 | calcium ion import | 4/151 | 91/18800 | 0.00623 | 0.04389 | 4 | neurons3 | Calcium transport |
| GO:1903351 | cellular response to dopamine | 4/151 | 91/18800 | 0.00623 | 0.04389 | 4 | neurons3 | Dopamine metabolism |
| GO:0051284 | positive regulation of sequestering of calcium ion | 2/151 | 15/18800 | 0.00628 | 0.04389 | 2 | neurons3 | Calcium transport |
| GO:1903350 | response to dopamine | 4/151 | 92/18800 | 0.00647 | 0.04481 | 4 | neurons3 | Dopamine metabolism |
| GO:0006091 | generation of precursor metabolites and energy | 10/151 | 494/18800 | 0.00666 | 0.04585 | 10 | neurons3 | Energy production |
| GO:0050807 | regulation of synapse organization | 6/151 | 209/18800 | 0.0069 | 0.0473 | 6 | neurons3 | Synapse assembly |
| GO:0055064 | chloride ion homeostasis | 2/151 | 16/18800 | 0.00714 | 0.04789 | 2 | neurons3 | Ion transport |
| GO:1901379 | regulation of potassium ion transmembrane transport | 4/151 | 95/18800 | 0.00724 | 0.04812 | 4 | neurons3 | Ion transport |
| GO:0099054 | presynapse assembly | 3/151 | 50/18800 | 0.00755 | 0.04945 | 3 | neurons3 | Synapse assembly |
| GO:0050803 | regulation of synapse structure or activity | 6/151 | 215/18800 | 0.00789 | 0.05085 | 6 | neurons3 | Synapse assembly |
| GO:0015872 | dopamine transport | 3/151 | 51/18800 | 0.00798 | 0.05085 | 3 | neurons3 | Dopamine metabolism |
| GO:0043268 | positive regulation of potassium ion transport | 3/151 | 51/18800 | 0.00798 | 0.05085 | 3 | neurons3 | Ion transport |
| GO:0071732 | cellular response to nitric oxide | 2/151 | 17/18800 | 0.00805 | 0.05085 | 2 | neurons3 | Oxidative stress |
| GO:0099172 | presynapse organization | 3/151 | 52/18800 | 0.00842 | 0.05289 | 3 | neurons3 | Synapse assembly |
| GO:1903169 | regulation of calcium ion transmembrane transport | 5/151 | 156/18800 | 0.00856 | 0.05353 | 5 | neurons3 | Calcium transport |
| GO:0043217 | myelin maintenance | 2/151 | 18/18800 | 0.00901 | 0.0555 | 2 | neurons3 | Enhanced myelination |
| GO:0010524 | positive regulation of calcium ion transport into cytosol | 3/151 | 54/18800 | 0.00934 | 0.0571 | 3 | neurons3 | Calcium transport |
| GO:0051963 | regulation of synapse assembly | 4/151 | 103/18800 | 0.00957 | 0.05806 | 4 | neurons3 | Synapse assembly |
| GO:0016082 | synaptic vesicle priming | 2/151 | 19/18800 | 0.01002 | 0.06051 | 2 | neurons3 | Vesicle |
| GO:2000300 | regulation of synaptic vesicle exocytosis | 3/151 | 56/18800 | 0.01032 | 0.06171 | 3 | neurons3 | Vesicle |
| GO:0008637 | apoptotic mitochondrial changes | 4/151 | 107/18800 | 0.0109 | 0.06455 | 4 | neurons3 | Apoptosis |
| GO:0042053 | regulation of dopamine metabolic process | 2/151 | 20/18800 | 0.01107 | 0.06468 | 2 | neurons3 | Dopamine metabolism |
| GO:2000649 | regulation of sodium ion transmembrane transporter activity | 3/151 | 58/18800 | 0.01135 | 0.06554 | 3 | neurons3 | Ion transport |
| GO:0006826 | iron ion transport | 3/151 | 59/18800 | 0.01189 | 0.06818 | 3 | neurons3 | Ion transport |

| | | | | | | | | |
|---|---|---|---|---|---|---|---|---|
| GO:0016050 | vesicle organization | 7/151 | 306/18800 | 0.01194 | 0.06829 | 7 | neurons3 | Vesicle |
| GO:0043266 | regulation of potassium ion transport | 4/151 | 110/18800 | 0.01197 | 0.06831 | 4 | neurons3 | Ion transport |
| GO:0031629 | synaptic vesicle fusion to presynaptic active zone membrane | 2/151 | 21/18800 | 0.01218 | 0.06871 | 2 | neurons3 | Vesicle |
| GO:0050849 | negative regulation of calcium-mediated signaling | 2/151 | 21/18800 | 0.01218 | 0.06871 | 2 | neurons3 | Calcium transport |
| GO:0071731 | response to nitric oxide | 2/151 | 21/18800 | 0.01218 | 0.06871 | 2 | neurons3 | Oxidative stress |
| GO:0045428 | regulation of nitric oxide biosynthetic process | 3/151 | 61/18800 | 0.01301 | 0.07277 | 3 | neurons3 | Oxidative stress |
| GO:0010881 | regulation of cardiac muscle contraction by regulation of the release of sequestered calcium ion | 2/151 | 22/18800 | 0.01332 | 0.07352 | 2 | neurons3 | Muscle contraction |
| GO:0035584 | calcium-mediated signaling using intracellular calcium source | 2/151 | 22/18800 | 0.01332 | 0.07352 | 2 | neurons3 | Calcium transport |
| GO:0099500 | vesicle fusion to plasma membrane | 2/151 | 22/18800 | 0.01332 | 0.07352 | 2 | neurons3 | Vesicle |
| GO:0046916 | cellular transition metal ion homeostasis | 4/151 | 115/18800 | 0.0139 | 0.07636 | 4 | neurons3 | Ion transport |
| GO:0051965 | positive regulation of synapse assembly | 3/151 | 63/18800 | 0.01419 | 0.07739 | 3 | neurons3 | Synapse assembly |
| GO:0080164 | regulation of nitric oxide metabolic process | 3/151 | 64/18800 | 0.01481 | 0.07924 | 3 | neurons3 | Oxidative stress |
| GO:0006879 | cellular iron ion homeostasis | 3/151 | 66/18800 | 0.01607 | 0.08369 | 3 | neurons3 | Iron transport |
| GO:0051209 | release of sequestered calcium ion into cytosol | 4/151 | 121/18800 | 0.01647 | 0.08473 | 4 | neurons3 | Calcium transport |
| GO:1901016 | regulation of potassium ion transmembrane transporter activity | 3/151 | 67/18800 | 0.01673 | 0.0855 | 3 | neurons3 | Ion transport |
| GO:0051283 | negative regulation of sequestering of calcium ion | 4/151 | 122/18800 | 0.01692 | 0.08618 | 4 | neurons3 | Calcium transport |
| GO:0050860 | negative regulation of T cell receptor signaling pathway | 2/151 | 25/18800 | 0.01703 | 0.08618 | 2 | neurons3 | T cell activity |
| GO:0098719 | sodium ion import across plasma membrane | 2/151 | 25/18800 | 0.01703 | 0.08618 | 2 | neurons3 | Ion transport |
| GO:0051282 | regulation of sequestering of calcium ion | 4/151 | 124/18800 | 0.01786 | 0.08924 | 4 | neurons3 | Calcium transport |
| GO:0003143 | embryonic heart tube morphogenesis | 3/151 | 70/18800 | 0.01879 | 0.09279 | 3 | neurons3 | Morphogenesis |
| GO:0051208 | sequestering of calcium ion | 4/151 | 128/18800 | 0.01982 | 0.09578 | 4 | neurons3 | Calcium transport |
| GO:0031346 | positive regulation of cell projection organization | 7/151 | 341/18800 | 0.02041 | 0.09826 | 7 | neurons3 | Cell projection organization |
| GO:0097470 | ribbon synapse | 3/157 | 10/19594 | 6E-05 | 0.00038 | 3 | neurons3 | Synapse assembly |
| GO:0030285 | integral component of synaptic vesicle membrane | 4/157 | 28/19594 | 7E-05 | 0.00044 | 4 | neurons3 | Vesicle |
| GO:0060198 | clathrin-sculpted vesicle | 3/157 | 12/19594 | 0.00011 | 0.00065 | 3 | neurons3 | Vesicle |
| GO:0030672 | synaptic vesicle membrane | 6/157 | 103/19594 | 0.00018 | 0.00104 | 6 | neurons3 | Vesicle |

| | | | | | | | | |
|---|---|---|---|---|---|---|---|---|
| GO:0099501 | exocytic vesicle membrane | 6/157 | 103/19594 | 0.00018 | 0.00104 | 6 | neurons3 | Vesicle |
| GO:0098688 | parallel fiber to Purkinje cell synapse | 3/157 | 15/19594 | 0.00021 | 0.00114 | 3 | neurons3 | Synapse assembly |
| GO:0098685 | Schaffer collateral - CA1 synapse | 5/157 | 72/19594 | 0.00028 | 0.00144 | 5 | neurons3 | Synapse assembly |
| GO:0098563 | intrinsic component of synaptic vesicle membrane | 4/157 | 40/19594 | 0.00029 | 0.00146 | 4 | neurons3 | Vesicle |
| GO:0060205 | cytoplasmic vesicle lumen | 9/157 | 325/19594 | 0.00126 | 0.00477 | 9 | neurons3 | Vesicle |
| GO:0031983 | vesicle lumen | 9/157 | 327/19594 | 0.00131 | 0.00492 | 9 | neurons3 | Vesicle |
| GO:0030665 | clathrin-coated vesicle membrane | 5/157 | 111/19594 | 0.00201 | 0.0072 | 5 | neurons3 | Vesicle |
| GO:0032280 | symmetric synapse | 2/157 | 10/19594 | 0.00275 | 0.00909 | 2 | neurons3 | Synapse assembly |
| GO:0030136 | clathrin-coated vesicle | 6/157 | 192/19594 | 0.00456 | 0.0141 | 6 | neurons3 | Vesicle |
| GO:0030135 | coated vesicle | 7/157 | 290/19594 | 0.00898 | 0.02609 | 7 | neurons3 | Vesicle |
| GO:0030662 | coated vesicle membrane | 5/157 | 176/19594 | 0.01379 | 0.03747 | 5 | neurons3 | Vesicle |
| GO:0071682 | endocytic vesicle lumen | 2/157 | 23/19594 | 0.01445 | 0.03897 | 2 | neurons3 | Vesicle |
| GO:0015079 | potassium ion transmembrane transporter activity | 9/152 | 154/18410 | 1E-05 | 2E-04 | 9 | neurons3 | Ion transport |
| GO:0005237 | inhibitory extracellular ligand-gated ion channel activity | 4/152 | 15/18410 | 1E-05 | 2E-04 | 4 | neurons3 | Ion transport |
| GO:0015085 | calcium ion transmembrane transporter activity | 8/152 | 135/18410 | 2E-05 | 0.00042 | 8 | neurons3 | Calcium transport |
| GO:0005244 | voltage-gated ion channel activity | 9/152 | 201/18410 | 5E-05 | 0.00077 | 9 | neurons3 | Ion transport |
| GO:0022824 | transmitter-gated ion channel activity | 5/152 | 60/18410 | 0.00014 | 0.0017 | 5 | neurons3 | Ion transport |
| GO:0022851 | GABA-gated chloride ion channel activity | 3/152 | 13/18410 | 0.00015 | 0.00175 | 3 | neurons3 | Ion transport |
| GO:0099186 | structural constituent of postsynapse | 3/152 | 13/18410 | 0.00015 | 0.00175 | 3 | neurons3 | Synapse assembly |
| GO:0015276 | ligand-gated ion channel activity | 7/152 | 145/18410 | 2E-04 | 0.0022 | 7 | neurons3 | Ion transport |
| GO:0005230 | extracellular ligand-gated ion channel activity | 5/152 | 73/18410 | 0.00034 | 0.00301 | 5 | neurons3 | Ion transport |
| GO:0098918 | structural constituent of synapse | 3/152 | 19/18410 | 0.00049 | 0.00392 | 3 | neurons3 | Synapse assembly |
| GO:0051082 | unfolded protein binding | 6/152 | 121/18410 | 0.00051 | 4 | 6 | neurons3 | Unfolded Protein Response |
| GO:1904315 | transmitter-gated ion channel activity involved in regulation of postsynaptic membrane potential | 4/152 | 46/18410 | 0.00056 | 0.00417 | 4 | neurons3 | Ion transport |
| GO:0044183 | protein folding chaperone | 3/152 | 43/18410 | 0.00535 | 0.02395 | 3 | neurons3 | Unfolded Protein Response |
| GO:0050780 | dopamine receptor binding | 2/152 | 15/18410 | 0.00663 | 0.02874 | 2 | neurons3 | Dopamine metabolism |
| GO:0022821 | potassium ion antiporter activity | 2/152 | 16/18410 | 0.00753 | 0.03103 | 2 | neurons3 | Ion transport |

| | | | | | | | | |
|---|---|---|---|---|---|---|---|---|
| GO:0099604 | ligand-gated calcium channel activity | 2/152 | 26/18410 | 0.01933 | 0.06582 | 2 | neurons3 | Calcium transport |
| hsa04020 | Calcium signaling pathway | 11/89 | 240/8170 | 5E-05 | 0.00119 | 11 | neurons3 | Calcium transport |
| hsa04721 | Synaptic vesicle cycle | 6/89 | 78/8170 | 0.00019 | 0.00319 | 6 | neurons3 | Vesicle |
| hsa04713 | Circadian entrainment | 6/89 | 97/8170 | 0.00063 | 0.0078 | 6 | neurons3 | Circadian rhythm |
| hsa04260 | Cardiac muscle contraction | 4/89 | 87/8170 | 0.01475 | 0.07893 | 4 | neurons3 | Muscle contraction |
| GO:0010812 | negative regulation of cell-substrate adhesion | 11/472 | 66/18800 | 0 | 0.00013 | 11 | neurons4 | Cell adhesion |
| GO:0001953 | negative regulation of cell-matrix adhesion | 8/472 | 40/18800 | 1E-05 | 0.00066 | 8 | neurons4 | Cell adhesion |
| GO:0007162 | negative regulation of cell adhesion | 21/472 | 305/18800 | 3E-05 | 0.0029 | 21 | neurons4 | Cell adhesion |
| GO:0007409 | axonogenesis | 26/472 | 430/18800 | 4E-05 | 0.00317 | 26 | neurons4 | Axonogenesis |
| GO:0031346 | positive regulation of cell projection organization | 22/472 | 341/18800 | 6E-05 | 0.00449 | 22 | neurons4 | Cell projection organization |
| GO:0050808 | synapse organization | 25/472 | 419/18800 | 6E-05 | 0.00484 | 25 | neurons4 | Synapse assembly |
| GO:0099173 | postsynapse organization | 14/472 | 163/18800 | 7E-05 | 0.00484 | 14 | neurons4 | Synapse assembly |
| GO:0042552 | myelination | 12/472 | 138/18800 | 0.00019 | 0.00986 | 12 | neurons4 | Enhanced myelination |
| GO:0007272 | ensheathment of neurons | 12/472 | 140/18800 | 0.00022 | 11 | 12 | neurons4 | Enhanced myelination |
| GO:0008366 | axon ensheathment | 12/472 | 140/18800 | 0.00022 | 11 | 12 | neurons4 | Enhanced myelination |
| GO:0097120 | receptor localization to synapse | 7/472 | 56/18800 | 0.00048 | 0.01912 | 7 | neurons4 | Synapse assembly |
| GO:0051895 | negative regulation of focal adhesion assembly | 4/472 | 17/18800 | 0.00072 | 0.02528 | 4 | neurons4 | Cell adhesion |
| GO:0048488 | synaptic vesicle endocytosis | 7/472 | 61/18800 | 0.00081 | 0.02773 | 7 | neurons4 | Vesicle |
| GO:0031345 | negative regulation of cell projection organization | 13/472 | 188/18800 | 0.00096 | 0.03069 | 13 | neurons4 | Cell projection organization |
| GO:1903421 | regulation of synaptic vesicle recycling | 4/472 | 20/18800 | 0.00138 | 0.03948 | 4 | neurons4 | Vesicle |
| GO:0032228 | regulation of synaptic transmission, GABAergic | 5/472 | 34/18800 | 0.00149 | 0.04171 | 5 | neurons4 | GABAergic synapse |
| GO:0051056 | regulation of small GTPase mediated signal transduction | 17/472 | 299/18800 | 0.00155 | 0.04248 | 17 | neurons4 | GTPase activity |
| GO:2000392 | regulation of lamellipodium morphogenesis | 3/472 | 11/18800 | 0.00223 | 0.05338 | 3 | neurons4 | Morphogenesis |
| GO:2001241 | positive regulation of extrinsic apoptotic signaling pathway in absence of ligand | 3/472 | 11/18800 | 0.00223 | 0.05338 | 3 | neurons4 | Apoptosis |
| GO:0036465 | synaptic vesicle recycling | 7/472 | 73/18800 | 0.00235 | 0.05492 | 7 | neurons4 | Vesicle |
| GO:0010769 | regulation of cell morphogenesis involved in differentiation | 8/472 | 96/18800 | 0.00287 | 0.06319 | 8 | neurons4 | Morphogenesis |
| GO:0002664 | regulation of T cell tolerance induction | 3/472 | 13/18800 | 0.00373 | 0.07224 | 3 | neurons4 | T cell activity |

| | | | | | | | | |
|---|---|---|---|---|---|---|---|---|
| GO:1900242 | regulation of synaptic vesicle endocytosis | 3/472 | 13/18800 | 0.00373 | 0.07224 | 3 | neurons4 | Vesicle |
| GO:0051894 | positive regulation of focal adhesion assembly | 4/472 | 26/18800 | 0.00378 | 0.07262 | 4 | neurons4 | Cell adhesion |
| GO:2001239 | regulation of extrinsic apoptotic signaling pathway in absence of ligand | 5/472 | 44/18800 | 0.00474 | 0.08088 | 5 | neurons4 | Apoptosis |
| GO:0016236 | macroautophagy | 16/472 | 306/18800 | 0.00477 | 0.08088 | 16 | neurons4 | Autophagy |
| GO:0050771 | negative regulation of axonogenesis | 6/472 | 64/18800 | 0.00532 | 0.08564 | 6 | neurons4 | Axonogenesis |
| GO:0002517 | T cell tolerance induction | 3/472 | 15/18800 | 0.00571 | 0.0902 | 3 | neurons4 | T cell activity |
| GO:0099175 | regulation of postsynapse organization | 7/472 | 86/18800 | 0.00587 | 0.09169 | 7 | neurons4 | Synapse assembly |
| GO:0070588 | calcium ion transmembrane transport | 16/472 | 314/18800 | 0.00608 | 0.09403 | 16 | neurons4 | Calcium transport |
| GO:0098978 | glutamatergic synapse | 20/476 | 319/19594 | 0.00011 | 0.00252 | 20 | neurons4 | Synapse assembly |
| GO:0098793 | presynapse | 25/476 | 492/19594 | 0.00045 | 0.00678 | 25 | neurons4 | Synapse assembly |
| GO:0005925 | focal adhesion | 22/476 | 419/19594 | 0.00064 | 0.00896 | 22 | neurons4 | Cell adhesion |
| GO:0098562 | cytoplasmic side of membrane | 13/476 | 193/19594 | 0.00091 | 0.01171 | 13 | neurons4 | Plasma membrane |
| GO:0031234 | extrinsic component of cytoplasmic side of plasma membrane | 8/476 | 99/19594 | 0.00284 | 0.02779 | 8 | neurons4 | Plasma membrane |
| GO:0009898 | cytoplasmic side of plasma membrane | 11/476 | 169/19594 | 0.00288 | 0.02779 | 11 | neurons4 | Plasma membrane |
| GO:0030695 | GTPase regulator activity | 29/478 | 488/18410 | 3E-05 | 0.00382 | 29 | neurons4 | GTPase activity |
| GO:0005096 | GTPase activator activity | 17/478 | 274/18410 | 0.00086 | 0.03155 | 17 | neurons4 | GTPase activity |
| GO:0008199 | ferric iron binding | 3/478 | 11/18410 | 0.00246 | 0.06114 | 3 | neurons4 | Iron transport |
| GO:0005217 | intracellular ligand-gated ion channel activity | 4/478 | 30/18410 | 0.0072 | 0.09885 | 4 | neurons4 | Ion transport |
| hsa04670 | Leukocyte transendothelial migration | 12/208 | 114/8170 | 3E-05 | 0.0039 | 12 | neurons4 | Leukocytes |
| hsa04724 | Glutamatergic synapse | 9/208 | 114/8170 | 0.00243 | 0.03113 | 9 | neurons4 | Synapse assembly |
| hsa04510 | Focal adhesion | 12/208 | 201/8170 | 0.00518 | 0.05102 | 12 | neurons4 | Cell adhesion |
| hsa04725 | Cholinergic synapse | 8/208 | 113/8170 | 0.00804 | 0.06429 | 8 | neurons4 | Synapse assembly |
| GO:0098703 | calcium ion import across plasma membrane | 9/696 | 32/18800 | 0 | 0.00018 | 9 | neurons5 | Calcium transport |
| GO:1902656 | calcium ion import into cytosol | 9/696 | 33/18800 | 0 | 0.00023 | 9 | neurons5 | Calcium transport |
| GO:0032228 | regulation of synaptic transmission, GABAergic | 9/696 | 34/18800 | 0 | 0.00028 | 9 | neurons5 | GABAergic synapse |
| GO:0034765 | regulation of ion transmembrane transport | 39/696 | 476/18800 | 0 | 3E-04 | 39 | neurons5 | Ion transport |
| GO:0048854 | brain morphogenesis | 9/696 | 36/18800 | 0 | 0.00041 | 9 | neurons5 | Morphogenesis |
| GO:0097120 | receptor localization to synapse | 11/696 | 56/18800 | 1E-05 | 0.00045 | 11 | neurons5 | Synapse assembly |

| | | | | | | | | |
|---|---|---|---|---|---|---|---|---|
| GO:0098742 | cell-cell adhesion via plasma-membrane adhesion molecules | 27/696 | 279/18800 | 1E-05 | 0.00045 | 27 | neurons5 | Cell adhesion |
| GO:0099643 | signal release from synapse | 18/696 | 145/18800 | 1E-05 | 0.00055 | 18 | neurons5 | Synapse assembly |
| GO:0006816 | calcium ion transport | 35/696 | 424/18800 | 1E-05 | 0.00065 | 35 | neurons5 | Calcium transport |
| GO:0051932 | synaptic transmission, GABAergic | 10/696 | 51/18800 | 1E-05 | 0.00096 | 10 | neurons5 | GABAergic synapse |
| GO:0099504 | synaptic vesicle cycle | 20/696 | 183/18800 | 2E-05 | 0.00099 | 20 | neurons5 | Vesicle |
| GO:0097553 | calcium ion transmembrane import into cytosol | 18/696 | 154/18800 | 2E-05 | 0.00103 | 18 | neurons5 | Calcium transport |
| GO:0060402 | calcium ion transport into cytosol | 19/696 | 171/18800 | 2E-05 | 0.00122 | 19 | neurons5 | Calcium transport |
| GO:0048814 | regulation of dendrite morphogenesis | 11/696 | 64/18800 | 2E-05 | 0.00124 | 11 | neurons5 | Morphogenesis |
| GO:0099560 | synaptic membrane adhesion | 7/696 | 25/18800 | 2E-05 | 0.00142 | 7 | neurons5 | Cell adhesion |
| GO:0060401 | cytosolic calcium ion transport | 20/696 | 190/18800 | 3E-05 | 0.00148 | 20 | neurons5 | Calcium transport |
| GO:0070509 | calcium ion import | 13/696 | 91/18800 | 3E-05 | 0.00157 | 13 | neurons5 | Calcium transport |
| GO:0035249 | synaptic transmission, glutamatergic | 13/696 | 94/18800 | 4E-05 | 0.00213 | 13 | neurons5 | Glutamatergic synapse |
| GO:0099003 | vesicle-mediated transport in synapse | 20/696 | 197/18800 | 5E-05 | 0.00227 | 20 | neurons5 | Vesicle |
| GO:0050770 | regulation of axonogenesis | 17/696 | 154/18800 | 6E-05 | 0.00284 | 17 | neurons5 | Axonogenesis |
| GO:0099068 | postsynapse assembly | 7/696 | 29/18800 | 7E-05 | 0.00341 | 7 | neurons5 | Synapse assembly |
| GO:0051592 | response to calcium ion | 16/696 | 144/18800 | 9E-05 | 0.0042 | 16 | neurons5 | Calcium transport |
| GO:0097091 | synaptic vesicle clustering | 5/696 | 14/18800 | 1E-04 | 0.0048 | 5 | neurons5 | Vesicle |
| GO:0035725 | sodium ion transmembrane transport | 18/696 | 177/18800 | 1E-04 | 0.0048 | 18 | neurons5 | Ion transport |
| GO:1990806 | ligand-gated ion channel signaling pathway | 6/696 | 22/18800 | 0.00011 | 0.00495 | 6 | neurons5 | Ion transport |
| GO:0050772 | positive regulation of axonogenesis | 11/696 | 77/18800 | 0.00012 | 0.00518 | 11 | neurons5 | Axonogenesis |
| GO:0001952 | regulation of cell-matrix adhesion | 14/696 | 123/18800 | 0.00019 | 0.0073 | 14 | neurons5 | Cell adhesion |
| GO:0048041 | focal adhesion assembly | 11/696 | 83/18800 | 0.00024 | 0.00897 | 11 | neurons5 | Cell adhesion |
| GO:0021955 | central nervous system neuron axonogenesis | 7/696 | 35/18800 | 0.00025 | 0.00938 | 7 | neurons5 | Axonogenesis |
| GO:0060997 | dendritic spine morphogenesis | 9/696 | 58/18800 | 0.00026 | 0.00957 | 9 | neurons5 | Morphogenesis |
| GO:0090630 | activation of GTPase activity | 13/696 | 114/18800 | 0.00031 | 0.01076 | 13 | neurons5 | GTPase activity |
| GO:0099566 | regulation of postsynaptic cytosolic calcium ion concentration | 4/696 | 10/18800 | 0.00033 | 0.01142 | 4 | neurons5 | Calcium transport |
| GO:0006814 | sodium ion transport | 21/696 | 249/18800 | 4E-04 | 0.01385 | 21 | neurons5 | Ion transport |
| GO:0099587 | inorganic ion import across plasma membrane | 13/696 | 118/18800 | 0.00043 | 0.01427 | 13 | neurons5 | Ion transport |

| | | | | | | | | |
|---|---|---|---|---|---|---|---|---|
| GO:0099054 | presynapse assembly | 8/696 | 50/18800 | 0.00046 | 0.01505 | 8 | neurons5 | Synapse assembly |
| GO:0002028 | regulation of sodium ion transport | 11/696 | 90/18800 | 0.00048 | 0.01518 | 11 | neurons5 | Ion transport |
| GO:0051893 | regulation of focal adhesion assembly | 9/696 | 63/18800 | 0.00049 | 0.01518 | 9 | neurons5 | Cell adhesion |
| GO:0010959 | regulation of metal ion transport | 29/696 | 403/18800 | 0.00053 | 0.01579 | 29 | neurons5 | Ion transport |
| GO:0016079 | synaptic vesicle exocytosis | 12/696 | 107/18800 | 6E-04 | 0.01727 | 12 | neurons5 | Vesicle |
| GO:0099172 | presynapse organization | 8/696 | 52/18800 | 6E-04 | 0.01727 | 8 | neurons5 | Synapse assembly |
| GO:0031345 | negative regulation of cell projection organization | 17/696 | 188/18800 | 0.00064 | 0.01797 | 17 | neurons5 | Cell projection organization |
| GO:0001738 | morphogenesis of a polarized epithelium | 11/696 | 94/18800 | 7E-04 | 0.01902 | 11 | neurons5 | Morphogenesis |
| GO:1901379 | regulation of potassium ion transmembrane transport | 11/696 | 95/18800 | 0.00077 | 0.02005 | 11 | neurons5 | Ion transport |
| GO:0035418 | protein localization to synapse | 9/696 | 67/18800 | 0.00078 | 0.0201 | 9 | neurons5 | Synapse assembly |
| GO:0051480 | regulation of cytosolic calcium ion concentration | 26/696 | 356/18800 | 0.00081 | 0.02064 | 26 | neurons5 | Calcium transport |
| GO:0032412 | regulation of ion transmembrane transporter activity | 21/696 | 263/18800 | 0.00082 | 0.02081 | 21 | neurons5 | Ion transport |
| GO:0099558 | maintenance of synapse structure | 5/696 | 21/18800 | 0.00085 | 0.02104 | 5 | neurons5 | Synapse assembly |
| GO:0048512 | circadian behavior | 7/696 | 43/18800 | 0.00093 | 0.02223 | 7 | neurons5 | Circadian rhythm |
| GO:0061001 | regulation of dendritic spine morphogenesis | 7/696 | 44/18800 | 0.00107 | 0.02426 | 7 | neurons5 | Morphogenesis |
| GO:0031589 | cell-substrate adhesion | 26/696 | 364/18800 | 0.00112 | 0.02519 | 26 | neurons5 | Cell adhesion |
| GO:0010810 | regulation of cell-substrate adhesion | 18/696 | 217/18800 | 0.00123 | 0.02697 | 18 | neurons5 | Cell adhesion |
| GO:0071805 | potassium ion transmembrane transport | 18/696 | 219/18800 | 0.00137 | 0.02858 | 18 | neurons5 | Ion transport |
| GO:0019722 | calcium-mediated signaling | 17/696 | 202/18800 | 0.00142 | 0.02905 | 17 | neurons5 | Calcium transport |
| GO:0007156 | homophilic cell adhesion via plasma membrane adhesion molecules | 15/696 | 168/18800 | 0.00148 | 0.02999 | 15 | neurons5 | Cell adhesion |
| GO:0050775 | positive regulation of dendrite morphogenesis | 6/696 | 35/18800 | 0.00163 | 0.03158 | 6 | neurons5 | Morphogenesis |
| GO:0007623 | circadian rhythm | 17/696 | 205/18800 | 0.00167 | 0.03179 | 17 | neurons5 | Circadian rhythm |
| GO:0032414 | positive regulation of ion transmembrane transporter activity | 11/696 | 106/18800 | 0.00189 | 0.03426 | 11 | neurons5 | Ion transport |
| GO:0062237 | protein localization to postsynapse | 6/696 | 36/18800 | 0.0019 | 0.03426 | 6 | neurons5 | Synapse assembly |
| GO:1904861 | excitatory synapse assembly | 5/696 | 25/18800 | 0.00197 | 0.03502 | 5 | neurons5 | Synapse assembly |
| GO:1901021 | positive regulation of calcium ion transmembrane transporter activity | 6/696 | 37/18800 | 0.0022 | 0.03804 | 6 | neurons5 | Calcium transport |

| | | | | | | | | |
|---|---|---|---|---|---|---|---|---|
| GO:0002433 | immune response-regulating cell surface receptor signaling pathway involved in phagocytosis | 5/696 | 26/18800 | 0.00236 | 0.03928 | 5 | neurons5 | Immune response |
| GO:0021952 | central nervous system projection neuron axonogenesis | 5/696 | 26/18800 | 0.00236 | 0.03928 | 5 | neurons5 | Axonogenesis |
| GO:0043266 | regulation of potassium ion transport | 11/696 | 110/18800 | 0.00254 | 0.04128 | 11 | neurons5 | Ion transport |
| GO:0043268 | positive regulation of potassium ion transport | 7/696 | 51/18800 | 0.0026 | 0.04188 | 7 | neurons5 | Ion transport |
| GO:0055074 | calcium ion homeostasis | 30/696 | 468/18800 | 0.00262 | 0.04217 | 30 | neurons5 | Calcium transport |
| GO:0017156 | calcium-ion regulated exocytosis | 8/696 | 65/18800 | 0.00266 | 0.04257 | 8 | neurons5 | Calcium transport |
| GO:0002220 | innate immune response activating cell surface receptor signaling pathway | 5/696 | 27/18800 | 0.00281 | 0.04387 | 5 | neurons5 | Immune response |
| GO:0010812 | negative regulation of cell-substrate adhesion | 8/696 | 66/18800 | 0.00293 | 0.04527 | 8 | neurons5 | Cell adhesion |
| GO:0061577 | calcium ion transmembrane transport via high voltage-gated calcium channel | 4/696 | 17/18800 | 0.00301 | 0.04577 | 4 | neurons5 | Calcium transport |
| GO:2000027 | regulation of animal organ morphogenesis | 12/696 | 129/18800 | 0.00304 | 0.04592 | 12 | neurons5 | Morphogenesis |
| GO:1902305 | regulation of sodium ion transmembrane transport | 8/696 | 67/18800 | 0.00322 | 0.04799 | 8 | neurons5 | Ion transport |
| GO:0002758 | innate immune response-activating signal transduction | 5/696 | 28/18800 | 0.00332 | 0.04861 | 5 | neurons5 | Immune response |
| GO:0006874 | cellular calcium ion homeostasis | 29/696 | 456/18800 | 0.00343 | 0.0495 | 29 | neurons5 | Calcium transport |
| GO:0071277 | cellular response to calcium ion | 9/696 | 84/18800 | 0.00384 | 0.05444 | 9 | neurons5 | Calcium transport |
| GO:0006813 | potassium ion transport | 18/696 | 243/18800 | 0.00421 | 0.05893 | 18 | neurons5 | Ion transport |
| GO:0048592 | eye morphogenesis | 13/696 | 154/18800 | 0.00482 | 0.06444 | 13 | neurons5 | Morphogenesis |
| GO:0010522 | regulation of calcium ion transport into cytosol | 10/696 | 103/18800 | 0.00484 | 0.06456 | 10 | neurons5 | Calcium transport |
| GO:0007204 | positive regulation of cytosolic calcium ion concentration | 22/696 | 325/18800 | 0.00495 | 0.06529 | 22 | neurons5 | Calcium transport |
| GO:0010749 | regulation of nitric oxide mediated signal transduction | 3/696 | 10/18800 | 0.00499 | 0.06529 | 3 | neurons5 | Oxidative stress |
| GO:0010880 | regulation of release of sequestered calcium ion into cytosol by sarcoplasmic reticulum | 5/696 | 31/18800 | 0.00524 | 0.06739 | 5 | neurons5 | Calcium transport |
| GO:1901381 | positive regulation of potassium ion transmembrane transport | 6/696 | 44/18800 | 0.00536 | 0.06814 | 6 | neurons5 | Ion transport |
| GO:1903169 | regulation of calcium ion transmembrane transport | 13/696 | 156/18800 | 0.00537 | 0.06814 | 13 | neurons5 | Calcium transport |
| GO:0002011 | morphogenesis of an epithelial sheet | 7/696 | 58/18800 | 0.0054 | 0.06814 | 7 | neurons5 | Morphogenesis |

| | | | | | | | | |
|---|---|---|---|---|---|---|---|---|
| GO:1902473 | regulation of protein localization to synapse | 4/696 | 20/18800 | 0.00562 | 0.07012 | 4 | neurons5 | Synapse assembly |
| GO:0010171 | body morphogenesis | 6/696 | 45/18800 | 6 | 0.07406 | 6 | neurons5 | Morphogenesis |
| GO:0006941 | striated muscle contraction | 14/696 | 178/18800 | 0.00655 | 0.07902 | 14 | neurons5 | Muscle contraction |
| GO:0007160 | cell-matrix adhesion | 17/696 | 235/18800 | 0.00671 | 0.07907 | 17 | neurons5 | Cell adhesion |
| GO:0034767 | positive regulation of ion transmembrane transport | 13/696 | 163/18800 | 0.0077 | 0.08846 | 13 | neurons5 | Ion transport |
| GO:0050850 | positive regulation of calcium-mediated signaling | 5/696 | 34/18800 | 0.00783 | 0.08846 | 5 | neurons5 | Calcium transport |
| GO:0010881 | regulation of cardiac muscle contraction by regulation of the release of sequestered calcium ion | 4/696 | 22/18800 | 8 | 0.09016 | 4 | neurons5 | Muscle contraction |
| GO:0014808 | release of sequestered calcium ion into cytosol by sarcoplasmic reticulum | 5/696 | 35/18800 | 0.00886 | 0.09536 | 5 | neurons5 | Calcium transport |
| GO:0010769 | regulation of cell morphogenesis involved in differentiation | 9/696 | 96/18800 | 0.00917 | 0.09794 | 9 | neurons5 | Morphogenesis |
| GO:0005891 | voltage-gated calcium channel complex | 9/722 | 44/19594 | 3E-05 | 0.00034 | 9 | neurons5 | Calcium transport |
| GO:0060077 | inhibitory synapse | 6/722 | 19/19594 | 4E-05 | 5E-04 | 6 | neurons5 | Synapse assembly |
| GO:0098685 | Schaffer collateral - CA1 synapse | 11/722 | 72/19594 | 6E-05 | 0.00068 | 11 | neurons5 | Synapse assembly |
| GO:0060076 | excitatory synapse | 9/722 | 51/19594 | 9E-05 | 0.00096 | 9 | neurons5 | Synapse assembly |
| GO:0030672 | synaptic vesicle membrane | 13/722 | 103/19594 | 1E-04 | 0.00105 | 13 | neurons5 | Vesicle |
| GO:0099501 | exocytic vesicle membrane | 13/722 | 103/19594 | 1E-04 | 0.00105 | 13 | neurons5 | Vesicle |
| GO:0008021 | synaptic vesicle | 19/722 | 196/19594 | 0.00012 | 0.00119 | 19 | neurons5 | Vesicle |
| GO:0098686 | hippocampal mossy fiber to CA3 synapse | 7/722 | 34/19594 | 2E-04 | 0.0019 | 7 | neurons5 | Synapse assembly |
| GO:0070382 | exocytic vesicle | 19/722 | 214/19594 | 0.00038 | 0.00336 | 19 | neurons5 | Vesicle |
| GO:0098688 | parallel fiber to Purkinje cell synapse | 4/722 | 15/19594 | 0.0018 | 0.01267 | 4 | neurons5 | Synapse assembly |
| GO:0030658 | transport vesicle membrane | 16/722 | 205/19594 | 0.00393 | 0.02205 | 16 | neurons5 | Vesicle |
| GO:1990454 | L-type voltage-gated calcium channel complex | 3/722 | 11/19594 | 0.00659 | 0.0331 | 3 | neurons5 | Calcium transport |
| GO:0098992 | neuronal dense core vesicle | 3/722 | 13/19594 | 0.01081 | 0.0504 | 3 | neurons5 | Vesicle |
| GO:0030136 | clathrin-coated vesicle | 14/722 | 192/19594 | 0.01188 | 0.05489 | 14 | neurons5 | Vesicle |
| GO:0009898 | cytoplasmic side of plasma membrane | 12/722 | 169/19594 | 0.02282 | 0.0949 | 12 | neurons5 | Plasma membrane |
| GO:0046873 | metal ion transmembrane transporter activity | 37/710 | 428/18410 | 0 | 2E-04 | 37 | neurons5 | Ion transport |
| GO:0005245 | voltage-gated calcium channel activity | 10/710 | 45/18410 | 1E-05 | 0.00026 | 10 | neurons5 | Calcium transport |
| GO:0008331 | high voltage-gated calcium channel activity | 5/710 | 11/18410 | 3E-05 | 0.00113 | 5 | neurons5 | Calcium transport |
| GO:0005244 | voltage-gated ion channel activity | 21/710 | 201/18410 | 3E-05 | 0.00113 | 21 | neurons5 | Ion transport |

| | | | | | | | | |
|---|---|---|---|---|---|---|---|---|
| GO:0022824 | transmitter-gated ion channel activity | 10/710 | 60/18410 | 9E-05 | 0.00234 | 10 | neurons5 | Ion transport |
| GO:0009931 | calcium-dependent protein serine/threonine kinase activity | 6/710 | 22/18410 | 0.00014 | 0.00333 | 6 | neurons5 | Calcium transport |
| GO:0010857 | calcium-dependent protein kinase activity | 6/710 | 23/18410 | 0.00019 | 0.00409 | 6 | neurons5 | Calcium transport |
| GO:0005230 | extracellular ligand-gated ion channel activity | 10/710 | 73/18410 | 0.00047 | 0.00821 | 10 | neurons5 | Ion transport |
| GO:0098918 | structural constituent of synapse | 5/710 | 19/18410 | 0.00062 | 0.00966 | 5 | neurons5 | Synapse assembly |
| GO:0015276 | ligand-gated ion channel activity | 14/710 | 145/18410 | 0.00146 | 0.01974 | 14 | neurons5 | Ion transport |
| GO:0004698 | calcium-dependent protein kinase C activity | 4/710 | 16/18410 | 0.00275 | 0.03292 | 4 | neurons5 | Calcium transport |
| GO:0098632 | cell-cell adhesion mediator activity | 7/710 | 54/18410 | 0.00451 | 0.04632 | 7 | neurons5 | Cell adhesion |
| GO:0005432 | calcium:sodium antiporter activity | 3/710 | 10/18410 | 0.00559 | 0.05228 | 3 | neurons5 | Calcium transport |
| GO:0015368 | calcium:cation antiporter activity | 3/710 | 11/18410 | 0.00747 | 0.06334 | 3 | neurons5 | Calcium transport |
| GO:0031267 | small GTPase binding | 19/710 | 267/18410 | 0.00782 | 0.06427 | 19 | neurons5 | GTPase activity |
| GO:0098631 | cell adhesion mediator activity | 7/710 | 64/18410 | 0.01138 | 0.08469 | 7 | neurons5 | Cell adhesion |
| GO:0051020 | GTPase binding | 20/710 | 298/18410 | 0.0119 | 0.08674 | 20 | neurons5 | GTPase activity |
| GO:0099186 | structural constituent of postsynapse | 3/710 | 13/18410 | 0.01223 | 0.08717 | 3 | neurons5 | Synapse assembly |
| GO:0015081 | sodium ion transmembrane transporter activity | 12/710 | 150/18410 | 0.01345 | 0.0935 | 12 | neurons5 | Ion transport |
| hsa04510 | Focal adhesion | 20/279 | 201/8170 | 2E-05 | 0.00015 | 20 | neurons5 | Cell adhesion |
| hsa04726 | Serotonergic synapse | 14/279 | 115/8170 | 3E-05 | 0.00027 | 14 | neurons5 | Synapse assembly |
| hsa04270 | Vascular smooth muscle contraction | 15/279 | 134/8170 | 5E-05 | 0.00036 | 15 | neurons5 | Muscle contraction |
| hsa04660 | T cell receptor signaling pathway | 12/279 | 104/8170 | 0.00021 | 0.00135 | 12 | neurons5 | T cell activity |
| hsa04961 | Endocrine and other factor-regulated calcium reabsorption | 8/279 | 53/8170 | 0.00039 | 0.00241 | 8 | neurons5 | Calcium transport |
| hsa04514 | Cell adhesion molecules | 11/279 | 157/8170 | 0.01846 | 0.05446 | 11 | neurons5 | Cell adhesion |
| hsa04140 | Autophagy - animal | 10/279 | 141/8170 | 0.02233 | 0.06274 | 10 | neurons5 | Autophagy |
| hsa04215 | Apoptosis - multiple species | 4/279 | 32/8170 | 0.02256 | 0.06274 | 4 | neurons5 | Apoptosis |
| hsa04260 | Cardiac muscle contraction | 7/279 | 87/8170 | 0.02877 | 0.07691 | 7 | neurons5 | Muscle contraction |
| GO:0030695 | GTPase regulator activity | 5/20 | 488/18410 | 0.00014 | 0.00311 | 5 | astrocytes0 | GTPase activity |
| GO:0005096 | GTPase activator activity | 4/20 | 274/18410 | 0.00019 | 0.00335 | 4 | astrocytes0 | GTPase activity |
| GO:0003924 | GTPase activity | 2/20 | 336/18410 | 0.05083 | 0.09214 | 2 | astrocytes0 | GTPase activity |
| GO:0006936 | muscle contraction | 9/93 | 349/18800 | 6E-05 | 0.0409 | 9 | astrocytes1 | Muscle contraction |

| | | | | | | | | |
|---|---|---|---|---|---|---|---|---|
| hsa04270 | Vascular smooth muscle contraction | 5/45 | 134/8166 | 8E-04 | 0.06332 | 5 | astrocytes1 | Muscle contraction |
| hsa04713 | Circadian entrainment | 4/45 | 97/8166 | 0.00192 | 0.07631 | 4 | astrocytes1 | Circadian rhythm |
| GO:0007409 | axonogenesis | 12/98 | 430/18800 | 0 | 0.00019 | 12 | astrocytes2 | Axonogenesis |
| GO:0015872 | dopamine transport | 5/98 | 51/18800 | 1E-05 | 0.00047 | 5 | astrocytes2 | Dopamine metabolism |
| GO:0046034 | ATP metabolic process | 9/98 | 273/18800 | 1E-05 | 0.00078 | 9 | astrocytes2 | Energy production |
| GO:0042416 | dopamine biosynthetic process | 3/98 | 12/18800 | 3E-05 | 0.0015 | 3 | astrocytes2 | Dopamine metabolism |
| GO:0010882 | regulation of cardiac muscle contraction by calcium ion signaling | 3/98 | 26/18800 | 0.00033 | 0.00831 | 3 | astrocytes2 | Muscle contraction |
| GO:0001963 | synaptic transmission, dopaminergic | 3/98 | 27/18800 | 0.00037 | 0.00882 | 3 | astrocytes2 | Dopamine metabolism |
| GO:0014046 | dopamine secretion | 3/98 | 37/18800 | 0.00094 | 0.0172 | 3 | astrocytes2 | Dopamine metabolism |
| GO:0014059 | regulation of dopamine secretion | 3/98 | 37/18800 | 0.00094 | 0.0172 | 3 | astrocytes2 | Dopamine metabolism |
| GO:1903578 | regulation of ATP metabolic process | 4/98 | 89/18800 | 0.00121 | 0.0191 | 4 | astrocytes2 | Energy production |
| GO:0050770 | regulation of axonogenesis | 5/98 | 154/18800 | 0.00127 | 0.01972 | 5 | astrocytes2 | Axonogenesis |
| GO:0042417 | dopamine metabolic process | 3/98 | 42/18800 | 0.00136 | 0.02053 | 3 | astrocytes2 | Dopamine metabolism |
| GO:0051583 | dopamine uptake involved in synaptic transmission | 2/98 | 12/18800 | 0.00172 | 0.02303 | 2 | astrocytes2 | Dopamine metabolism |
| GO:0090494 | dopamine uptake | 2/98 | 17/18800 | 0.00348 | 0.03445 | 2 | astrocytes2 | Dopamine metabolism |
| GO:0006091 | generation of precursor metabolites and energy | 8/98 | 494/18800 | 0.00429 | 0.03939 | 8 | astrocytes2 | Energy production |
| GO:0043467 | regulation of generation of precursor metabolites and energy | 4/98 | 134/18800 | 0.00532 | 0.04461 | 4 | astrocytes2 | Energy production |
| GO:0010881 | regulation of cardiac muscle contraction by regulation of the release of sequestered calcium ion | 2/98 | 22/18800 | 0.00581 | 0.04606 | 2 | astrocytes2 | Muscle contraction |
| GO:0055117 | regulation of cardiac muscle contraction | 3/98 | 77/18800 | 0.0076 | 0.05394 | 3 | astrocytes2 | Muscle contraction |
| GO:0090151 | establishment of protein localization to mitochondrial membrane | 2/98 | 30/18800 | 0.01064 | 0.06398 | 2 | astrocytes2 | Mitochondrial changes |
| GO:1903579 | negative regulation of ATP metabolic process | 2/98 | 30/18800 | 0.01064 | 0.06398 | 2 | astrocytes2 | Energy production |
| GO:0006942 | regulation of striated muscle contraction | 3/98 | 96/18800 | 0.01384 | 0.07395 | 3 | astrocytes2 | Muscle contraction |
| GO:0021955 | central nervous system neuron axonogenesis | 2/98 | 35/18800 | 0.01431 | 0.07527 | 2 | astrocytes2 | Axonogenesis |
| GO:0007212 | dopamine receptor signaling pathway | 2/98 | 40/18800 | 0.01845 | 0.08748 | 2 | astrocytes2 | Dopamine metabolism |
| GO:0045429 | positive regulation of nitric oxide biosynthetic process | 2/98 | 41/18800 | 0.01933 | 0.08961 | 2 | astrocytes2 | Oxidative stress |

| | | | | | | | | |
|---|---|---|---|---|---|---|---|---|
| GO:0048512 | circadian behavior | 2/98 | 43/18800 | 0.02114 | 0.09497 | 2 | astrocytes2 | Circadian rhythm |
| GO:1904407 | positive regulation of nitric oxide metabolic process | 2/98 | 43/18800 | 0.02114 | 0.09497 | 2 | astrocytes2 | Oxidative stress |
| GO:0098691 | dopaminergic synapse | 2/98 | 11/19594 | 0.00132 | 0.00895 | 2 | astrocytes2 | Dopamine metabolism |
| GO:0044183 | protein folding chaperone | 3/97 | 43/18410 | 0.0015 | 0.01642 | 3 | astrocytes2 | Unfolded Protein Response |
| GO:0003924 | GTPase activity | 7/97 | 336/18410 | 2 | 0.01718 | 7 | astrocytes2 | GTPase activity |
| GO:0050998 | nitric-oxide synthase binding | 2/97 | 13/18410 | 0.00206 | 0.01718 | 2 | astrocytes2 | Oxidative stress |
| GO:0016620 | oxidoreductase activity, acting on the aldehyde or oxo group of donors, NAD or NADP as acceptor | 2/97 | 38/18410 | 0.01708 | 0.06543 | 2 | astrocytes2 | Oxidative stress |
| GO:0016903 | oxidoreductase activity, acting on the aldehyde or oxo group of donors | 2/97 | 46/18410 | 0.02447 | 0.08673 | 2 | astrocytes2 | Oxidative stress |
| GO:0051082 | unfolded protein binding | 3/97 | 121/18410 | 0.02612 | 0.08905 | 3 | astrocytes2 | Unfolded Protein Response |
| hsa04713 | Circadian entrainment | 7/74 | 97/8166 | 3E-05 | 0.00047 | 7 | astrocytes2 | Circadian rhythm |
| hsa04210 | Apoptosis | 8/74 | 136/8166 | 3E-05 | 5E-04 | 8 | astrocytes2 | Apoptosis |
| hsa04270 | Vascular smooth muscle contraction | 5/74 | 134/8166 | 0.0072 | 0.04026 | 5 | astrocytes2 | Muscle contraction |
| GO:0061077 | chaperone-mediated protein folding | 5/54 | 70/18800 | 0 | 0.00235 | 5 | astrocytes3 | Unfolded Protein Response |
| GO:0006457 | protein folding | 6/54 | 212/18800 | 3E-05 | 0.02184 | 6 | astrocytes3 | Unfolded Protein Response |
| GO:0006986 | response to unfolded protein | 5/54 | 139/18800 | 5E-05 | 0.02245 | 5 | astrocytes3 | Unfolded Protein Response |
| GO:0035966 | response to topologically incorrect protein | 5/54 | 160/18800 | 9E-05 | 0.02904 | 5 | astrocytes3 | Unfolded Protein Response |
| GO:0051085 | chaperone cofactor-dependent protein refolding | 3/54 | 32/18800 | 1E-04 | 0.02904 | 3 | astrocytes3 | Unfolded Protein Response |
| GO:0051084 | 'de novo' post-translational protein folding | 3/54 | 37/18800 | 0.00016 | 0.03678 | 3 | astrocytes3 | Unfolded Protein Response |
| GO:0006458 | 'de novo' protein folding | 3/54 | 41/18800 | 0.00022 | 0.03678 | 3 | astrocytes3 | Unfolded Protein Response |
| GO:0031998 | regulation of fatty acid beta-oxidation | 2/54 | 20/18800 | 0.00149 | 0.09824 | 2 | astrocytes3 | Oxidative stress |
| GO:0098869 | cellular oxidant detoxification | 2/14 | 100/18800 | 0.00245 | 0.0288 | 2 | astrocytes4 | Oxidative stress |
| GO:0051341 | regulation of oxidoreductase activity | 2/14 | 106/18800 | 0.00274 | 0.03035 | 2 | astrocytes4 | Oxidative stress |
| GO:0008637 | apoptotic mitochondrial changes | 2/14 | 107/18800 | 0.00279 | 0.03035 | 2 | astrocytes4 | Apoptosis |
| GO:0043619 | regulation of transcription from RNA polymerase II promoter in response to oxidative stress | 1/14 | 11/18800 | 0.00816 | 0.05809 | 1 | astrocytes4 | Oxidative stress |
| GO:1901030 | positive regulation of mitochondrial outer membrane permeabilization involved in apoptotic signaling pathway | 1/14 | 11/18800 | 0.00816 | 0.05809 | 1 | astrocytes4 | Apoptosis |

| | | | | | | | | |
|---|---|---|---|---|---|---|---|---|
| GO:0045088 | regulation of innate immune response | 2/14 | 231/18800 | 0.01241 | 0.05809 | 2 | astrocytes4 | Immune response |
| GO:0071732 | cellular response to nitric oxide | 1/14 | 17/18800 | 0.01259 | 0.05809 | 1 | astrocytes4 | Oxidative stress |
| GO:0002281 | macrophage activation involved in immune response | 1/14 | 18/18800 | 0.01333 | 0.05811 | 1 | astrocytes4 | Immune response |
| GO:0010310 | regulation of hydrogen peroxide metabolic process | 1/14 | 20/18800 | 0.0148 | 0.06018 | 1 | astrocytes4 | Oxidative stress |
| GO:0061760 | antifungal innate immune response | 1/14 | 21/18800 | 0.01553 | 0.06205 | 1 | astrocytes4 | Immune response |
| GO:0071731 | response to nitric oxide | 1/14 | 21/18800 | 0.01553 | 0.06205 | 1 | astrocytes4 | Oxidative stress |
| GO:0051000 | positive regulation of nitric-oxide synthase activity | 1/14 | 22/18800 | 0.01626 | 0.06224 | 1 | astrocytes4 | Oxidative stress |
| GO:0061081 | positive regulation of myeloid leukocyte cytokine production involved in immune response | 1/14 | 22/18800 | 0.01626 | 0.06224 | 1 | astrocytes4 | Leukocytes |
| GO:0019430 | removal of superoxide radicals | 1/14 | 23/18800 | 17 | 0.06266 | 1 | astrocytes4 | Oxidative stress |
| GO:1901028 | regulation of mitochondrial outer membrane permeabilization involved in apoptotic signaling pathway | 1/14 | 24/18800 | 0.01773 | 0.06266 | 1 | astrocytes4 | Apoptosis |
| GO:0071451 | cellular response to superoxide | 1/14 | 25/18800 | 0.01846 | 0.06367 | 1 | astrocytes4 | Oxidative stress |
| GO:0051204 | protein insertion into mitochondrial membrane | 1/14 | 26/18800 | 0.0192 | 0.06521 | 1 | astrocytes4 | Mitochondrial changes |
| GO:0002220 | innate immune response activating cell surface receptor signaling pathway | 1/14 | 27/18800 | 0.01993 | 0.0662 | 1 | astrocytes4 | Immune response |
| GO:0007263 | nitric oxide mediated signal transduction | 1/14 | 27/18800 | 0.01993 | 0.0662 | 1 | astrocytes4 | Oxidative stress |
| GO:0000303 | response to superoxide | 1/14 | 28/18800 | 0.02066 | 0.06621 | 1 | astrocytes4 | Oxidative stress |
| GO:0002758 | innate immune response-activating signal transduction | 1/14 | 28/18800 | 0.02066 | 0.06621 | 1 | astrocytes4 | Immune response |
| GO:0002313 | mature B cell differentiation involved in immune response | 1/14 | 29/18800 | 0.02139 | 0.06621 | 1 | astrocytes4 | Immune response |
| GO:0042744 | hydrogen peroxide catabolic process | 1/14 | 30/18800 | 0.02212 | 0.06621 | 1 | astrocytes4 | Oxidative stress |
| GO:0090151 | establishment of protein localization to mitochondrial membrane | 1/14 | 30/18800 | 0.02212 | 0.06621 | 1 | astrocytes4 | Mitochondrial changes |
| GO:0051354 | negative regulation of oxidoreductase activity | 1/14 | 35/18800 | 0.02576 | 0.0708 | 1 | astrocytes4 | Oxidative stress |
| GO:0097345 | mitochondrial outer membrane permeabilization | 1/14 | 35/18800 | 0.02576 | 0.0708 | 1 | astrocytes4 | Mitochondrial changes |
| GO:0045429 | positive regulation of nitric oxide biosynthetic process | 1/14 | 41/18800 | 0.03011 | 0.07608 | 1 | astrocytes4 | Oxidative stress |
| GO:1902110 | positive regulation of mitochondrial membrane permeability involved in apoptotic process | 1/14 | 41/18800 | 0.03011 | 0.07608 | 1 | astrocytes4 | Apoptosis |
| GO:0050999 | regulation of nitric-oxide synthase activity | 1/14 | 42/18800 | 0.03084 | 0.07676 | 1 | astrocytes4 | Oxidative stress |

| | | | | | | | | |
|---|---|---|---|---|---|---|---|---|
| GO:1902686 | mitochondrial outer membrane permeabilization involved in programmed cell death | 1/14 | 43/18800 | 0.03156 | 0.07737 | 1 | astrocytes4 | Mitochondrial changes |
| GO:1904407 | positive regulation of nitric oxide metabolic process | 1/14 | 43/18800 | 0.03156 | 0.07737 | 1 | astrocytes4 | Oxidative stress |
| GO:0002253 | activation of immune response | 2/14 | 386/18800 | 0.03251 | 0.07801 | 2 | astrocytes4 | Immune response |
| GO:0035794 | positive regulation of mitochondrial membrane permeability | 1/14 | 46/18800 | 0.03373 | 0.07871 | 1 | astrocytes4 | Mitochondrial changes |
| GO:1902108 | regulation of mitochondrial membrane permeability involved in apoptotic process | 1/14 | 48/18800 | 0.03517 | 0.07955 | 1 | astrocytes4 | Apoptosis |
| GO:0007409 | axonogenesis | 2/14 | 430/18800 | 0.03961 | 0.08355 | 2 | astrocytes4 | Axonogenesis |
| GO:0006979 | response to oxidative stress | 2/14 | 433/18800 | 0.04011 | 0.08382 | 2 | astrocytes4 | Oxidative stress |
| GO:0042743 | hydrogen peroxide metabolic process | 1/14 | 55/18800 | 0.0402 | 0.08382 | 1 | astrocytes4 | Oxidative stress |
| GO:0002218 | activation of innate immune response | 1/14 | 59/18800 | 0.04307 | 0.08604 | 1 | astrocytes4 | Immune response |
| GO:0051353 | positive regulation of oxidoreductase activity | 1/14 | 59/18800 | 0.04307 | 0.08604 | 1 | astrocytes4 | Oxidative stress |
| GO:0045428 | regulation of nitric oxide biosynthetic process | 1/14 | 61/18800 | 0.04449 | 0.0866 | 1 | astrocytes4 | Oxidative stress |
| GO:0046902 | regulation of mitochondrial membrane permeability | 1/14 | 62/18800 | 0.04521 | 0.08724 | 1 | astrocytes4 | Mitochondrial changes |
| GO:0050771 | negative regulation of axonogenesis | 1/14 | 64/18800 | 0.04663 | 0.08776 | 1 | astrocytes4 | Axonogenesis |
| GO:0080164 | regulation of nitric oxide metabolic process | 1/14 | 64/18800 | 0.04663 | 0.08776 | 1 | astrocytes4 | Oxidative stress |
| GO:0002720 | positive regulation of cytokine production involved in immune response | 1/14 | 68/18800 | 0.04948 | 0.0894 | 1 | astrocytes4 | Immune response |
| GO:0061077 | chaperone-mediated protein folding | 1/14 | 70/18800 | 0.0509 | 0.0907 | 1 | astrocytes4 | Unfolded Protein Response |
| GO:0006801 | superoxide metabolic process | 1/14 | 72/18800 | 0.05232 | 0.09143 | 1 | astrocytes4 | Oxidative stress |
| GO:0006809 | nitric oxide biosynthetic process | 1/14 | 75/18800 | 0.05444 | 0.09388 | 1 | astrocytes4 | Oxidative stress |
| GO:0046209 | nitric oxide metabolic process | 1/14 | 81/18800 | 0.05868 | 0.09967 | 1 | astrocytes4 | Oxidative stress |
| GO:0016209 | antioxidant activity | 2/15 | 85/18410 | 0.00213 | 0.02644 | 2 | astrocytes4 | Oxidative stress |
| GO:0050771 | negative regulation of axonogenesis | 9/344 | 64/18800 | 0 | 0.00046 | 9 | astrocytes5 | Axonogenesis |
| GO:0050770 | regulation of axonogenesis | 13/344 | 154/18800 | 1E-05 | 0.00093 | 13 | astrocytes5 | Axonogenesis |
| GO:0043087 | regulation of GTPase activity | 19/344 | 364/18800 | 5E-05 | 0.00489 | 19 | astrocytes5 | GTPase activity |
| GO:0051056 | regulation of small GTPase mediated signal transduction | 15/344 | 299/18800 | 0.00043 | 0.01768 | 15 | astrocytes5 | GTPase activity |
| GO:0043547 | positive regulation of GTPase activity | 13/344 | 272/18800 | 0.00158 | 0.03568 | 13 | astrocytes5 | GTPase activity |
| GO:0010818 | T cell chemotaxis | 4/344 | 28/18800 | 0.00159 | 0.03568 | 4 | astrocytes5 | T cell activity |

| | | | | | | | | |
|---|---|---|---|---|---|---|---|---|
| GO:2000406 | positive regulation of T cell migration | 4/344 | 31/18800 | 0.00235 | 0.04518 | 4 | astrocytes5 | T cell activity |
| GO:0010820 | positive regulation of T cell chemotaxis | 3/344 | 16/18800 | 0.00285 | 0.05108 | 3 | astrocytes5 | T cell activity |
| GO:0010819 | regulation of T cell chemotaxis | 3/344 | 17/18800 | 0.00341 | 0.0564 | 3 | astrocytes5 | T cell activity |
| GO:0034599 | cellular response to oxidative stress | 12/344 | 284/18800 | 0.00631 | 0.07947 | 12 | astrocytes5 | Oxidative stress |
| GO:0006979 | response to oxidative stress | 16/344 | 433/18800 | 0.00638 | 0.07988 | 16 | astrocytes5 | Oxidative stress |
| GO:0072678 | T cell migration | 5/344 | 67/18800 | 0.00762 | 0.08741 | 5 | astrocytes5 | T cell activity |
| GO:2000404 | regulation of T cell migration | 4/344 | 44/18800 | 0.00841 | 0.0927 | 4 | astrocytes5 | T cell activity |
| GO:1905360 | GTPase complex | 3/353 | 36/19594 | 0.0267 | 0.0992 | 3 | astrocytes5 | GTPase activity |
| GO:0030695 | GTPase regulator activity | 22/350 | 488/18410 | 0.00018 | 0.01209 | 22 | astrocytes5 | GTPase activity |
| GO:0050670 | regulation of lymphocyte proliferation | 3/6 | 230/18800 | 4E-05 | 0.00656 | 3 | microglia0 | Lymphocyte |
| GO:0070663 | regulation of leukocyte proliferation | 3/6 | 254/18800 | 5E-05 | 0.00656 | 3 | microglia0 | Leukocytes |
| GO:0046651 | lymphocyte proliferation | 3/6 | 296/18800 | 7E-05 | 0.00718 | 3 | microglia0 | Lymphocyte |
| GO:0070661 | leukocyte proliferation | 3/6 | 330/18800 | 1E-04 | 0.00817 | 3 | microglia0 | Leukocytes |
| GO:0050671 | positive regulation of lymphocyte proliferation | 2/6 | 141/18800 | 0.00082 | 0.02468 | 2 | microglia0 | Lymphocyte |
| GO:0070665 | positive regulation of leukocyte proliferation | 2/6 | 158/18800 | 0.00103 | 0.02721 | 2 | microglia0 | Leukocytes |
| GO:0042129 | regulation of T cell proliferation | 2/6 | 174/18800 | 0.00125 | 0.02928 | 2 | microglia0 | T cell activity |
| GO:0031345 | negative regulation of cell projection organization | 2/6 | 188/18800 | 0.00145 | 0.03102 | 2 | microglia0 | Cell projection organization |
| GO:0042098 | T cell proliferation | 2/6 | 204/18800 | 0.00171 | 0.03328 | 2 | microglia0 | T cell activity |
| GO:0045088 | regulation of innate immune response | 2/6 | 231/18800 | 0.00218 | 0.03328 | 2 | microglia0 | Immune response |
| GO:0060560 | developmental growth involved in morphogenesis | 2/6 | 234/18800 | 0.00224 | 0.03328 | 2 | microglia0 | Morphogenesis |
| GO:1902237 | positive regulation of endoplasmic reticulum stress-induced intrinsic apoptotic signaling pathway | 1/6 | 10/18800 | 0.00319 | 0.03328 | 1 | microglia0 | Apoptosis |
| GO:0006707 | cholesterol catabolic process | 1/6 | 11/18800 | 0.00351 | 0.03328 | 1 | microglia0 | Cholesterol metabolism |
| GO:1903897 | regulation of PERK-mediated unfolded protein response | 1/6 | 11/18800 | 0.00351 | 0.03328 | 1 | microglia0 | Unfolded Protein Response |
| GO:0001771 | immunological synapse formation | 1/6 | 14/18800 | 0.00446 | 0.03328 | 1 | microglia0 | Synapse assembly |
| GO:0010872 | regulation of cholesterol esterification | 1/6 | 14/18800 | 0.00446 | 0.03328 | 1 | microglia0 | Cholesterol metabolism |
| GO:0072578 | neurotransmitter-gated ion channel clustering | 1/6 | 14/18800 | 0.00446 | 0.03328 | 1 | microglia0 | Ion transport |
| GO:0050863 | regulation of T cell activation | 2/6 | 342/18800 | 0.00472 | 0.03328 | 2 | microglia0 | T cell activity |
| GO:1903037 | regulation of leukocyte cell-cell adhesion | 2/6 | 344/18800 | 0.00477 | 0.03328 | 2 | microglia0 | Leukocytes |

| | | | | | | | | |
|---|---|---|---|---|---|---|---|---|
| GO:1900102 | negative regulation of endoplasmic reticulum unfolded protein response | 1/6 | 15/18800 | 0.00478 | 0.03328 | 1 | microglia0 | Unfolded Protein Response |
| GO:0002921 | negative regulation of humoral immune response | 1/6 | 16/18800 | 0.0051 | 0.03328 | 1 | microglia0 | Immune response |
| GO:0051251 | positive regulation of lymphocyte activation | 2/6 | 371/18800 | 0.00553 | 0.03383 | 2 | microglia0 | Lymphocyte |
| GO:0034435 | cholesterol esterification | 1/6 | 18/18800 | 0.00573 | 0.03383 | 1 | microglia0 | Cholesterol metabolism |
| GO:0036499 | PERK-mediated unfolded protein response | 1/6 | 18/18800 | 0.00573 | 0.03383 | 1 | microglia0 | Unfolded Protein Response |
| GO:0007159 | leukocyte cell-cell adhesion | 2/6 | 381/18800 | 0.00582 | 0.03383 | 2 | microglia0 | Leukocytes |
| GO:0043691 | reverse cholesterol transport | 1/6 | 20/18800 | 0.00637 | 0.03398 | 1 | microglia0 | Cholesterol metabolism |
| GO:0051000 | positive regulation of nitric-oxide synthase activity | 1/6 | 22/18800 | 7 | 0.03531 | 1 | microglia0 | Oxidative stress |
| GO:0002696 | positive regulation of leukocyte activation | 2/6 | 421/18800 | 0.00707 | 0.03531 | 2 | microglia0 | Leukocytes |
| GO:0045540 | regulation of cholesterol biosynthetic process | 1/6 | 23/18800 | 0.00732 | 0.03531 | 1 | microglia0 | Cholesterol metabolism |
| GO:0007409 | axonogenesis | 2/6 | 430/18800 | 0.00737 | 0.03531 | 2 | microglia0 | Axonogenesis |
| GO:0071577 | zinc ion transmembrane transport | 1/6 | 24/18800 | 0.00764 | 0.03531 | 1 | microglia0 | Ion transport |
| GO:0022407 | regulation of cell-cell adhesion | 2/6 | 456/18800 | 0.00825 | 0.03553 | 2 | microglia0 | Cell adhesion |
| GO:0007263 | nitric oxide mediated signal transduction | 1/6 | 27/18800 | 0.00859 | 0.03553 | 1 | microglia0 | Oxidative stress |
| GO:0010875 | positive regulation of cholesterol efflux | 1/6 | 27/18800 | 0.00859 | 0.03553 | 1 | microglia0 | Cholesterol metabolism |
| GO:0006829 | zinc ion transport | 1/6 | 28/18800 | 0.0089 | 0.03553 | 1 | microglia0 | Ion transport |
| GO:0007271 | synaptic transmission, cholinergic | 1/6 | 29/18800 | 0.00922 | 0.03553 | 1 | microglia0 | Cholinergic synapse |
| GO:1900101 | regulation of endoplasmic reticulum unfolded protein response | 1/6 | 30/18800 | 0.00954 | 0.03553 | 1 | microglia0 | Unfolded Protein Response |
| GO:1902235 | regulation of endoplasmic reticulum stress-induced intrinsic apoptotic signaling pathway | 1/6 | 32/18800 | 0.01017 | 0.03615 | 1 | microglia0 | Apoptosis |
| GO:0090181 | regulation of cholesterol metabolic process | 1/6 | 37/18800 | 0.01175 | 0.0377 | 1 | microglia0 | Cholesterol metabolism |
| GO:0050999 | regulation of nitric-oxide synthase activity | 1/6 | 42/18800 | 0.01333 | 0.04088 | 1 | microglia0 | Oxidative stress |
| GO:0002920 | regulation of humoral immune response | 1/6 | 45/18800 | 0.01428 | 0.04215 | 1 | microglia0 | Immune response |
| GO:0010874 | regulation of cholesterol efflux | 1/6 | 52/18800 | 0.01648 | 0.04463 | 1 | microglia0 | Cholesterol metabolism |
| GO:0006695 | cholesterol biosynthetic process | 1/6 | 58/18800 | 0.01837 | 0.04656 | 1 | microglia0 | Cholesterol metabolism |
| GO:0051353 | positive regulation of oxidoreductase activity | 1/6 | 59/18800 | 0.01869 | 0.04692 | 1 | microglia0 | Oxidative stress |

| | | | | | | | | |
|---|---|---|---|---|---|---|---|---|
| GO:2001244 | positive regulation of intrinsic apoptotic signaling pathway | 1/6 | 61/18800 | 0.01931 | 0.04764 | 1 | microglia0 | Apoptosis |
| GO:0070059 | intrinsic apoptotic signaling pathway in response to endoplasmic reticulum stress | 1/6 | 63/18800 | 0.01994 | 0.04812 | 1 | microglia0 | Apoptosis |
| GO:0050771 | negative regulation of axonogenesis | 1/6 | 64/18800 | 0.02026 | 0.04866 | 1 | microglia0 | Axonogenesis |
| GO:0042130 | negative regulation of T cell proliferation | 1/6 | 68/18800 | 0.02151 | 0.04953 | 1 | microglia0 | T cell activity |
| GO:0033344 | cholesterol efflux | 1/6 | 69/18800 | 0.02182 | 0.04964 | 1 | microglia0 | Cholesterol metabolism |
| GO:0045824 | negative regulation of innate immune response | 1/6 | 73/18800 | 0.02308 | 0.05102 | 1 | microglia0 | Immune response |
| GO:0030968 | endoplasmic reticulum unfolded protein response | 1/6 | 76/18800 | 0.02401 | 0.05206 | 1 | microglia0 | Unfolded Protein Response |
| GO:0030301 | cholesterol transport | 1/6 | 79/18800 | 0.02495 | 0.05326 | 1 | microglia0 | Cholesterol metabolism |
| GO:0048844 | artery morphogenesis | 1/6 | 79/18800 | 0.02495 | 0.05326 | 1 | microglia0 | Morphogenesis |
| GO:0050672 | negative regulation of lymphocyte proliferation | 1/6 | 84/18800 | 0.02651 | 0.0547 | 1 | microglia0 | Lymphocyte |
| GO:0099175 | regulation of postsynapse organization | 1/6 | 86/18800 | 0.02714 | 0.05497 | 1 | microglia0 | Synapse assembly |
| GO:0070664 | negative regulation of leukocyte proliferation | 1/6 | 91/18800 | 0.0287 | 0.05648 | 1 | microglia0 | Leukocytes |
| GO:0034620 | cellular response to unfolded protein | 1/6 | 98/18800 | 0.03088 | 0.05849 | 1 | microglia0 | Unfolded Protein Response |
| GO:0042632 | cholesterol homeostasis | 1/6 | 98/18800 | 0.03088 | 0.05849 | 1 | microglia0 | Cholesterol metabolism |
| GO:0098869 | cellular oxidant detoxification | 1/6 | 100/18800 | 0.0315 | 0.05904 | 1 | microglia0 | Oxidative stress |
| GO:0000041 | transition metal ion transport | 1/6 | 101/18800 | 0.03181 | 0.05904 | 1 | microglia0 | Ion transport |
| GO:0042102 | positive regulation of T cell proliferation | 1/6 | 103/18800 | 0.03243 | 0.0594 | 1 | microglia0 | T cell activity |
| GO:0002444 | myeloid leukocyte mediated immunity | 1/6 | 104/18800 | 0.03274 | 0.05958 | 1 | microglia0 | Leukocytes |
| GO:0051341 | regulation of oxidoreductase activity | 1/6 | 106/18800 | 0.03336 | 0.05992 | 1 | microglia0 | Oxidative stress |
| GO:0002698 | negative regulation of immune effector process | 1/6 | 112/18800 | 0.03522 | 0.06245 | 1 | microglia0 | Immune response |
| GO:0035967 | cellular response to topologically incorrect protein | 1/6 | 117/18800 | 0.03677 | 0.06437 | 1 | microglia0 | Unfolded Protein Response |
| GO:0050868 | negative regulation of T cell activation | 1/6 | 125/18800 | 0.03924 | 0.06681 | 1 | microglia0 | T cell activity |
| GO:0001909 | leukocyte mediated cytotoxicity | 1/6 | 131/18800 | 0.04109 | 0.0689 | 1 | microglia0 | Leukocytes |
| GO:2001235 | positive regulation of apoptotic signaling pathway | 1/6 | 134/18800 | 0.04202 | 0.06982 | 1 | microglia0 | Apoptosis |
| GO:0006986 | response to unfolded protein | 1/6 | 139/18800 | 0.04356 | 0.07126 | 1 | microglia0 | Unfolded Protein Response |
| GO:0008203 | cholesterol metabolic process | 1/6 | 139/18800 | 0.04356 | 0.07126 | 1 | microglia0 | Cholesterol metabolism |

| | | | | | | | | |
|---|---|---|---|---|---|---|---|---|
| GO:1903038 | negative regulation of leukocyte cell-cell adhesion | 1/6 | 144/18800 | 0.04509 | 0.07138 | 1 | microglia0 | Leukocytes |
| GO:0043524 | negative regulation of neuron apoptotic process | 1/6 | 145/18800 | 0.0454 | 0.07138 | 1 | microglia0 | Apoptosis |
| GO:0050770 | regulation of axonogenesis | 1/6 | 154/18800 | 0.04816 | 0.07384 | 1 | microglia0 | Axonogenesis |
| GO:0035966 | response to topologically incorrect protein | 1/6 | 160/18800 | 0.05 | 0.07581 | 1 | microglia0 | Unfolded Protein Response |
| GO:0051250 | negative regulation of lymphocyte activation | 1/6 | 161/18800 | 0.0503 | 0.07586 | 1 | microglia0 | Lymphocyte |
| GO:0099173 | postsynapse organization | 1/6 | 163/18800 | 0.05091 | 0.07637 | 1 | microglia0 | Synapse assembly |
| GO:2001242 | regulation of intrinsic apoptotic signaling pathway | 1/6 | 171/18800 | 0.05336 | 0.07806 | 1 | microglia0 | Apoptosis |
| GO:0050777 | negative regulation of immune response | 1/6 | 179/18800 | 0.05579 | 0.07919 | 1 | microglia0 | Immune response |
| GO:0002695 | negative regulation of leukocyte activation | 1/6 | 193/18800 | 0.06004 | 0.08269 | 1 | microglia0 | Leukocytes |
| GO:0022408 | negative regulation of cell-cell adhesion | 1/6 | 199/18800 | 0.06186 | 0.08354 | 1 | microglia0 | Cell adhesion |
| GO:0043523 | regulation of neuron apoptotic process | 1/6 | 207/18800 | 0.06428 | 0.08596 | 1 | microglia0 | Apoptosis |
| GO:0050807 | regulation of synapse organization | 1/6 | 209/18800 | 0.06488 | 0.08615 | 1 | microglia0 | Synapse assembly |
| GO:0050803 | regulation of synapse structure or activity | 1/6 | 215/18800 | 0.06669 | 0.08769 | 1 | microglia0 | Synapse assembly |
| GO:0016064 | immunoglobulin mediated immune response | 1/6 | 216/18800 | 0.06699 | 0.08769 | 1 | microglia0 | Immune response |
| GO:0050870 | positive regulation of T cell activation | 1/6 | 223/18800 | 0.0691 | 0.08919 | 1 | microglia0 | T cell activity |
| GO:0002274 | myeloid leukocyte activation | 1/6 | 232/18800 | 0.0718 | 0.09116 | 1 | microglia0 | Leukocytes |
| GO:0051402 | neuron apoptotic process | 1/6 | 241/18800 | 0.0745 | 0.09355 | 1 | microglia0 | Apoptosis |
| GO:1903039 | positive regulation of leukocyte cell-cell adhesion | 1/6 | 245/18800 | 0.0757 | 0.09441 | 1 | microglia0 | Leukocytes |
| GO:0071682 | endocytic vesicle lumen | 1/6 | 23/19594 | 0.00702 | 0.03503 | 1 | microglia0 | Vesicle |
| GO:0030285 | integral component of synaptic vesicle membrane | 1/6 | 28/19594 | 0.00854 | 0.03503 | 1 | microglia0 | Vesicle |
| GO:0098563 | intrinsic component of synaptic vesicle membrane | 1/6 | 40/19594 | 0.01219 | 0.03569 | 1 | microglia0 | Vesicle |
| GO:0030669 | clathrin-coated endocytic vesicle membrane | 1/6 | 72/19594 | 0.02185 | 0.05972 | 1 | microglia0 | Vesicle |
| GO:0045334 | clathrin-coated endocytic vesicle | 1/6 | 91/19594 | 0.02755 | 0.07059 | 1 | microglia0 | Vesicle |
| GO:0030672 | synaptic vesicle membrane | 1/6 | 103/19594 | 0.03113 | 0.07091 | 1 | microglia0 | Vesicle |
| GO:0099501 | exocytic vesicle membrane | 1/6 | 103/19594 | 0.03113 | 0.07091 | 1 | microglia0 | Vesicle |
| GO:0030665 | clathrin-coated vesicle membrane | 1/6 | 111/19594 | 0.03352 | 0.07232 | 1 | microglia0 | Vesicle |
| GO:0030662 | coated vesicle membrane | 1/6 | 176/19594 | 0.0527 | 0.09601 | 1 | microglia0 | Vesicle |
| GO:0030136 | clathrin-coated vesicle | 1/6 | 192/19594 | 0.05738 | 0.09601 | 1 | microglia0 | Vesicle |
| GO:0030666 | endocytic vesicle membrane | 1/6 | 194/19594 | 0.05796 | 0.09601 | 1 | microglia0 | Vesicle |

| | | | | | | | | |
|---|---|---|---|---|---|---|---|---|
| GO:0008021 | synaptic vesicle | 1/6 | 196/19594 | 0.05854 | 0.09601 | 1 | microglia0 | Vesicle |
| GO:0030658 | transport vesicle membrane | 1/6 | 205/19594 | 0.06116 | 0.09645 | 1 | microglia0 | Vesicle |
| GO:0070382 | exocytic vesicle | 1/6 | 214/19594 | 0.06377 | 0.09684 | 1 | microglia0 | Vesicle |
| GO:0120020 | cholesterol transfer activity | 1/6 | 22/18410 | 0.00715 | 0.0342 | 1 | microglia0 | Cholesterol metabolism |
| GO:0016209 | antioxidant activity | 1/6 | 85/18410 | 0.02739 | 0.0447 | 1 | microglia0 | Oxidative stress |
| GO:0140375 | immune receptor activity | 1/6 | 148/18410 | 0.04728 | 0.06923 | 1 | microglia0 | Immune response |
| GO:0099003 | vesicle-mediated transport in synapse | 12/86 | 197/18800 | 0 | 0 | 12 | microglia1 | Vesicle |
| GO:0050808 | synapse organization | 15/86 | 419/18800 | 0 | 0 | 15 | microglia1 | Synapse assembly |
| GO:0099504 | synaptic vesicle cycle | 11/86 | 183/18800 | 0 | 0 | 11 | microglia1 | Vesicle |
| GO:0007409 | axonogenesis | 15/86 | 430/18800 | 0 | 0 | 15 | microglia1 | Axonogenesis |
| GO:0048278 | vesicle docking | 5/86 | 57/18800 | 1E-05 | 0.00061 | 5 | microglia1 | Vesicle |
| GO:0050770 | regulation of axonogenesis | 7/86 | 154/18800 | 1E-05 | 0.00063 | 7 | microglia1 | Axonogenesis |
| GO:0099173 | postsynapse organization | 7/86 | 163/18800 | 1E-05 | 0.00084 | 7 | microglia1 | Synapse assembly |
| GO:0050807 | regulation of synapse organization | 7/86 | 209/18800 | 5E-05 | 0.00271 | 7 | microglia1 | Synapse assembly |
| GO:0061684 | chaperone-mediated autophagy | 3/86 | 16/18800 | 5E-05 | 0.00271 | 3 | microglia1 | Autophagy |
| GO:0099643 | signal release from synapse | 6/86 | 145/18800 | 5E-05 | 0.00271 | 6 | microglia1 | Synapse assembly |
| GO:0050803 | regulation of synapse structure or activity | 7/86 | 215/18800 | 6E-05 | 0.00272 | 7 | microglia1 | Synapse assembly |
| GO:0015872 | dopamine transport | 4/86 | 51/18800 | 9E-05 | 0.00358 | 4 | microglia1 | Dopamine metabolism |
| GO:0060560 | developmental growth involved in morphogenesis | 7/86 | 234/18800 | 1E-04 | 0.00377 | 7 | microglia1 | Morphogenesis |
| GO:0031629 | synaptic vesicle fusion to presynaptic active zone membrane | 3/86 | 21/18800 | 0.00012 | 0.00415 | 3 | microglia1 | Vesicle |
| GO:0016079 | synaptic vesicle exocytosis | 5/86 | 107/18800 | 0.00013 | 0.00459 | 5 | microglia1 | Vesicle |
| GO:0099500 | vesicle fusion to plasma membrane | 3/86 | 22/18800 | 0.00013 | 0.0046 | 3 | microglia1 | Vesicle |
| GO:0007416 | synapse assembly | 6/86 | 180/18800 | 0.00018 | 0.00579 | 6 | microglia1 | Synapse assembly |
| GO:0048499 | synaptic vesicle membrane organization | 3/86 | 26/18800 | 0.00022 | 0.00697 | 3 | microglia1 | Vesicle |
| GO:0036465 | synaptic vesicle recycling | 4/86 | 73/18800 | 0.00035 | 0.00913 | 4 | microglia1 | Vesicle |
| GO:0050772 | positive regulation of axonogenesis | 4/86 | 77/18800 | 0.00043 | 0.01079 | 4 | microglia1 | Axonogenesis |
| GO:0006904 | vesicle docking involved in exocytosis | 3/86 | 36/18800 | 0.00059 | 0.01383 | 3 | microglia1 | Vesicle |
| GO:0051402 | neuron apoptotic process | 6/86 | 241/18800 | 0.00083 | 0.01775 | 6 | microglia1 | Apoptosis |
| GO:0042416 | dopamine biosynthetic process | 2/86 | 12/18800 | 0.00133 | 0.02359 | 2 | microglia1 | Dopamine metabolism |

| | | | | | | | | |
|---|---|---|---|---|---|---|---|---|
| GO:0005513 | detection of calcium ion | 2/86 | 13/18800 | 0.00156 | 0.02687 | 2 | microglia1 | Calcium transport |
| GO:0043523 | regulation of neuron apoptotic process | 5/86 | 207/18800 | 0.00261 | 0.03768 | 5 | microglia1 | Apoptosis |
| GO:0045956 | positive regulation of calcium ion-dependent exocytosis | 2/86 | 17/18800 | 0.00269 | 0.03779 | 2 | microglia1 | Calcium transport |
| GO:0048791 | calcium ion-regulated exocytosis of neurotransmitter | 2/86 | 17/18800 | 0.00269 | 0.03779 | 2 | microglia1 | Calcium transport |
| GO:0045428 | regulation of nitric oxide biosynthetic process | 3/86 | 61/18800 | 0.00275 | 0.03782 | 3 | microglia1 | Oxidative stress |
| GO:0048488 | synaptic vesicle endocytosis | 3/86 | 61/18800 | 0.00275 | 0.03782 | 3 | microglia1 | Vesicle |
| GO:0050771 | negative regulation of axonogenesis | 3/86 | 64/18800 | 0.00315 | 0.04115 | 3 | microglia1 | Axonogenesis |
| GO:0080164 | regulation of nitric oxide metabolic process | 3/86 | 64/18800 | 0.00315 | 0.04115 | 3 | microglia1 | Oxidative stress |
| GO:0017156 | calcium-ion regulated exocytosis | 3/86 | 65/18800 | 0.00329 | 0.04273 | 3 | microglia1 | Calcium transport |
| GO:0016082 | synaptic vesicle priming | 2/86 | 19/18800 | 0.00336 | 0.04286 | 2 | microglia1 | Vesicle |
| GO:1904427 | positive regulation of calcium ion transmembrane transport | 3/86 | 71/18800 | 0.00422 | 0.04843 | 3 | microglia1 | Calcium transport |
| GO:0051592 | response to calcium ion | 4/86 | 144/18800 | 0.00431 | 0.04893 | 4 | microglia1 | Calcium transport |
| GO:0043524 | negative regulation of neuron apoptotic process | 4/86 | 145/18800 | 0.00442 | 0.04893 | 4 | microglia1 | Apoptosis |
| GO:0031338 | regulation of vesicle fusion | 2/86 | 22/18800 | 0.0045 | 0.04893 | 2 | microglia1 | Vesicle |
| GO:0031346 | positive regulation of cell projection organization | 6/86 | 341/18800 | 0.00475 | 0.05107 | 6 | microglia1 | Cell projection organization |
| GO:0006809 | nitric oxide biosynthetic process | 3/86 | 75/18800 | 0.00492 | 0.05213 | 3 | microglia1 | Oxidative stress |
| GO:0055117 | regulation of cardiac muscle contraction | 3/86 | 77/18800 | 0.0053 | 0.05577 | 3 | microglia1 | Muscle contraction |
| GO:1903169 | regulation of calcium ion transmembrane transport | 4/86 | 156/18800 | 0.00572 | 0.05807 | 4 | microglia1 | Calcium transport |
| GO:0060074 | synapse maturation | 2/86 | 25/18800 | 0.0058 | 0.05807 | 2 | microglia1 | Synapse assembly |
| GO:1902107 | positive regulation of leukocyte differentiation | 4/86 | 157/18800 | 0.00585 | 0.05807 | 4 | microglia1 | Leukocytes |
| GO:0051279 | regulation of release of sequestered calcium ion into cytosol | 3/86 | 80/18800 | 0.00589 | 0.05822 | 3 | microglia1 | Calcium transport |
| GO:0046209 | nitric oxide metabolic process | 3/86 | 81/18800 | 0.0061 | 0.05969 | 3 | microglia1 | Oxidative stress |
| GO:0003007 | heart morphogenesis | 5/86 | 254/18800 | 0.0062 | 0.06033 | 5 | microglia1 | Morphogenesis |
| GO:0010882 | regulation of cardiac muscle contraction by calcium ion signaling | 2/86 | 26/18800 | 0.00626 | 0.06033 | 2 | microglia1 | Muscle contraction |
| GO:0060314 | regulation of ryanodine-sensitive calcium-release channel activity | 2/86 | 26/18800 | 0.00626 | 0.06033 | 2 | microglia1 | Calcium transport |
| GO:0001963 | synaptic transmission, dopaminergic | 2/86 | 27/18800 | 0.00674 | 0.06298 | 2 | microglia1 | Dopamine metabolism |

| | | | | | | | | |
|---|---|---|---|---|---|---|---|---|
| GO:0099175 | regulation of postsynapse organization | 3/86 | 86/18800 | 0.0072 | 0.06613 | 3 | microglia1 | Synapse assembly |
| GO:0099068 | postsynapse assembly | 2/86 | 29/18800 | 0.00775 | 0.06928 | 2 | microglia1 | Synapse assembly |
| GO:0043270 | positive regulation of ion transport | 5/86 | 273/18800 | 0.00834 | 0.07297 | 5 | microglia1 | Ion transport |
| GO:1901019 | regulation of calcium ion transmembrane transporter activity | 3/86 | 91/18800 | 0.00841 | 0.07324 | 3 | microglia1 | Calcium transport |
| GO:0010880 | regulation of release of sequestered calcium ion into cytosol by sarcoplasmic reticulum | 2/86 | 31/18800 | 0.00882 | 0.07593 | 2 | microglia1 | Calcium transport |
| GO:0034109 | homotypic cell-cell adhesion | 3/86 | 93/18800 | 0.00892 | 0.07612 | 3 | microglia1 | Cell adhesion |
| GO:0035249 | synaptic transmission, glutamatergic | 3/86 | 94/18800 | 0.00918 | 0.07805 | 3 | microglia1 | Glutamatergic synapse |
| GO:0006942 | regulation of striated muscle contraction | 3/86 | 96/18800 | 0.00973 | 0.08098 | 3 | microglia1 | Muscle contraction |
| GO:0042063 | gliogenesis | 5/86 | 291/18800 | 0.01079 | 0.08568 | 5 | microglia1 | Gliogenesis |
| GO:0031345 | negative regulation of cell projection organization | 4/86 | 188/18800 | 0.01088 | 0.08568 | 4 | microglia1 | Cell projection organization |
| GO:0014808 | release of sequestered calcium ion into cytosol by sarcoplasmic reticulum | 2/86 | 35/18800 | 0.01116 | 0.08568 | 2 | microglia1 | Calcium transport |
| GO:0021955 | central nervous system neuron axonogenesis | 2/86 | 35/18800 | 0.01116 | 0.08568 | 2 | microglia1 | Axonogenesis |
| GO:0010522 | regulation of calcium ion transport into cytosol | 3/86 | 103/18800 | 0.01177 | 0.0875 | 3 | microglia1 | Calcium transport |
| GO:0051963 | regulation of synapse assembly | 3/86 | 103/18800 | 0.01177 | 0.0875 | 3 | microglia1 | Synapse assembly |
| GO:1903514 | release of sequestered calcium ion into cytosol by endoplasmic reticulum | 2/86 | 36/18800 | 0.01178 | 0.0875 | 2 | microglia1 | Calcium transport |
| GO:0006906 | vesicle fusion | 3/86 | 105/18800 | 0.01239 | 0.08997 | 3 | microglia1 | Vesicle |
| GO:0014046 | dopamine secretion | 2/86 | 37/18800 | 0.01242 | 0.08997 | 2 | microglia1 | Dopamine metabolism |
| GO:0014059 | regulation of dopamine secretion | 2/86 | 37/18800 | 0.01242 | 0.08997 | 2 | microglia1 | Dopamine metabolism |
| GO:1901021 | positive regulation of calcium ion transmembrane transporter activity | 2/86 | 37/18800 | 0.01242 | 0.08997 | 2 | microglia1 | Calcium transport |
| GO:0019722 | calcium-mediated signaling | 4/86 | 202/18800 | 0.01386 | 0.09733 | 4 | microglia1 | Calcium transport |
| GO:0007212 | dopamine receptor signaling pathway | 2/86 | 40/18800 | 0.01441 | 0.09955 | 2 | microglia1 | Dopamine metabolism |
| GO:0017158 | regulation of calcium ion-dependent exocytosis | 2/86 | 40/18800 | 0.01441 | 0.09955 | 2 | microglia1 | Calcium transport |
| GO:0051281 | positive regulation of release of sequestered calcium ion into cytosol | 2/86 | 40/18800 | 0.01441 | 0.09955 | 2 | microglia1 | Calcium transport |
| GO:0098793 | presynapse | 16/86 | 492/19594 | 0 | 0 | 16 | microglia1 | Synapse assembly |
| GO:0060198 | clathrin-sculpted vesicle | 3/86 | 12/19594 | 2E-05 | 0.00045 | 3 | microglia1 | Vesicle |

| | | | | | | | | |
|---|---|---|---|---|---|---|---|---|
| GO:0008021 | synaptic vesicle | 7/86 | 196/19594 | 2E-05 | 0.00053 | 7 | microglia1 | Vesicle |
| GO:0070382 | exocytic vesicle | 7/86 | 214/19594 | 4E-05 | 0.00082 | 7 | microglia1 | Vesicle |
| GO:0060076 | excitatory synapse | 4/86 | 51/19594 | 7E-05 | 0.00118 | 4 | microglia1 | Synapse assembly |
| GO:0098978 | glutamatergic synapse | 8/86 | 319/19594 | 8E-05 | 0.0012 | 8 | microglia1 | Synapse assembly |
| GO:0030672 | synaptic vesicle membrane | 5/86 | 103/19594 | 9E-05 | 0.00122 | 5 | microglia1 | Vesicle |
| GO:0099501 | exocytic vesicle membrane | 5/86 | 103/19594 | 9E-05 | 0.00122 | 5 | microglia1 | Vesicle |
| GO:0098685 | Schaffer collateral - CA1 synapse | 4/86 | 72/19594 | 0.00028 | 0.00281 | 4 | microglia1 | Synapse assembly |
| GO:0030658 | transport vesicle membrane | 6/86 | 205/19594 | 0.00029 | 0.00281 | 6 | microglia1 | Vesicle |
| GO:0030133 | transport vesicle | 8/86 | 402/19594 | 0.00038 | 0.00352 | 8 | microglia1 | Vesicle |
| GO:0098563 | intrinsic component of synaptic vesicle membrane | 3/86 | 40/19594 | 0.00072 | 0.00574 | 3 | microglia1 | Vesicle |
| GO:0098984 | neuron to neuron synapse | 7/86 | 347/19594 | 0.00083 | 0.00631 | 7 | microglia1 | Synapse assembly |
| GO:0098691 | dopaminergic synapse | 2/86 | 11/19594 | 0.00102 | 0.00747 | 2 | microglia1 | Dopamine metabolism |
| GO:0098992 | neuronal dense core vesicle | 2/86 | 13/19594 | 0.00144 | 0.00921 | 2 | microglia1 | Vesicle |
| GO:0098562 | cytoplasmic side of membrane | 5/86 | 193/19594 | 0.00161 | 0.01004 | 5 | microglia1 | Plasma membrane |
| GO:0032279 | asymmetric synapse | 6/86 | 323/19594 | 0.00298 | 0.0166 | 6 | microglia1 | Synapse assembly |
| GO:0030139 | endocytic vesicle | 6/86 | 342/19594 | 0.00395 | 0.02062 | 6 | microglia1 | Vesicle |
| GO:0009898 | cytoplasmic side of plasma membrane | 4/86 | 169/19594 | 0.00655 | 0.02843 | 4 | microglia1 | Plasma membrane |
| GO:0030285 | integral component of synaptic vesicle membrane | 2/86 | 28/19594 | 0.00668 | 0.02852 | 2 | microglia1 | Vesicle |
| GO:0031234 | extrinsic component of cytoplasmic side of plasma membrane | 3/86 | 99/19594 | 0.00946 | 0.03536 | 3 | microglia1 | Plasma membrane |
| GO:0030665 | clathrin-coated vesicle membrane | 3/86 | 111/19594 | 0.01288 | 0.04518 | 3 | microglia1 | Vesicle |
| GO:0060205 | cytoplasmic vesicle lumen | 5/86 | 325/19594 | 0.01422 | 0.04719 | 5 | microglia1 | Vesicle |
| GO:0031983 | vesicle lumen | 5/86 | 327/19594 | 0.01456 | 0.04719 | 5 | microglia1 | Vesicle |
| GO:0043209 | myelin sheath | 2/86 | 45/19594 | 0.01668 | 0.05234 | 2 | microglia1 | Enhanced myelination |
| GO:0099186 | structural constituent of postsynapse | 3/86 | 13/18410 | 3E-05 | 0.00103 | 3 | microglia1 | Synapse assembly |
| GO:0098918 | structural constituent of synapse | 3/86 | 19/18410 | 9E-05 | 0.0022 | 3 | microglia1 | Synapse assembly |
| GO:0048306 | calcium-dependent protein binding | 4/86 | 87/18410 | 0.00074 | 0.01568 | 4 | microglia1 | Calcium transport |
| GO:0044183 | protein folding chaperone | 3/86 | 43/18410 | 0.00106 | 0.0201 | 3 | microglia1 | Unfolded Protein Response |
| GO:0003924 | GTPase activity | 6/86 | 336/18410 | 0.00489 | 0.04631 | 6 | microglia1 | GTPase activity |
| GO:0015079 | potassium ion transmembrane transporter activity | 4/86 | 154/18410 | 0.00588 | 0.05276 | 4 | microglia1 | Ion transport |

| | | | | | | | | |
|---|---|---|---|---|---|---|---|---|
| hsa04728 | Dopaminergic synapse | 8/54 | 132/8170 | 0 | 0.00019 | 8 | microglia1 | Dopamine metabolism |
| hsa04721 | Synaptic vesicle cycle | 5/54 | 78/8170 | 0.00015 | 0.00456 | 5 | microglia1 | Vesicle |
| hsa04713 | Circadian entrainment | 4/54 | 97/8170 | 0.00375 | 0.03949 | 4 | microglia1 | Circadian rhythm |
| hsa04726 | Serotonergic synapse | 4/54 | 115/8170 | 0.00686 | 0.06819 | 4 | microglia1 | Synapse assembly |
| hsa04210 | Apoptosis | 4/54 | 136/8170 | 0.01222 | 0.0982 | 4 | microglia1 | Apoptosis |
| GO:0030695 | GTPase regulator activity | 7/33 | 488/18410 | 2E-05 | 0.00089 | 7 | microglia2 | GTPase activity |
| GO:0005096 | GTPase activator activity | 4/33 | 274/18410 | 0.0014 | 0.03006 | 4 | microglia2 | GTPase activity |
| GO:0010885 | regulation of cholesterol storage | 4/131 | 19/18800 | 1E-05 | 0.00942 | 4 | microglia3 | Cholesterol metabolism |
| GO:0010878 | cholesterol storage | 4/131 | 21/18800 | 1E-05 | 0.00942 | 4 | microglia3 | Cholesterol metabolism |
| GO:0051056 | regulation of small GTPase mediated signal transduction | 10/131 | 299/18800 | 5E-05 | 0.0217 | 10 | microglia3 | GTPase activity |
| GO:0099173 | postsynapse organization | 7/131 | 163/18800 | 0.00015 | 0.04222 | 7 | microglia3 | Synapse assembly |
| GO:0030050 | vesicle transport along actin filament | 3/131 | 19/18800 | 3E-04 | 0.04675 | 3 | microglia3 | Vesicle |
| GO:0043087 | regulation of GTPase activity | 9/131 | 364/18800 | 0.00103 | 0.0565 | 9 | microglia3 | GTPase activity |
| GO:0021602 | cranial nerve morphogenesis | 3/131 | 29/18800 | 0.00106 | 0.0565 | 3 | microglia3 | Morphogenesis |
| GO:0060411 | cardiac septum morphogenesis | 4/131 | 71/18800 | 0.00153 | 0.06746 | 4 | microglia3 | Morphogenesis |
| GO:0099518 | vesicle cytoskeletal trafficking | 4/131 | 71/18800 | 0.00153 | 0.06746 | 4 | microglia3 | Vesicle |
| GO:0007435 | salivary gland morphogenesis | 3/131 | 34/18800 | 0.00169 | 0.07132 | 3 | microglia3 | Morphogenesis |
| GO:0022612 | gland morphogenesis | 5/131 | 123/18800 | 0.00171 | 0.07132 | 5 | microglia3 | Morphogenesis |
| GO:0060562 | epithelial tube morphogenesis | 8/131 | 328/18800 | 0.00213 | 0.07134 | 8 | microglia3 | Morphogenesis |
| GO:0003151 | outflow tract morphogenesis | 4/131 | 80/18800 | 0.00237 | 0.07134 | 4 | microglia3 | Morphogenesis |
| GO:0006707 | cholesterol catabolic process | 2/131 | 11/18800 | 0.00254 | 0.07134 | 2 | microglia3 | Cholesterol metabolism |
| GO:0010887 | negative regulation of cholesterol storage | 2/131 | 11/18800 | 0.00254 | 0.07134 | 2 | microglia3 | Cholesterol metabolism |
| GO:0021610 | facial nerve morphogenesis | 2/131 | 11/18800 | 0.00254 | 0.07134 | 2 | microglia3 | Morphogenesis |
| GO:0050808 | synapse organization | 9/131 | 419/18800 | 0.00269 | 0.07465 | 9 | microglia3 | Synapse assembly |
| GO:0099175 | regulation of postsynapse organization | 4/131 | 86/18800 | 0.00308 | 0.07781 | 4 | microglia3 | Synapse assembly |
| GO:0002573 | myeloid leukocyte differentiation | 6/131 | 210/18800 | 0.00356 | 0.07981 | 6 | microglia3 | Leukocytes |
| GO:0030139 | endocytic vesicle | 11/135 | 342/19594 | 3E-05 | 0.00616 | 11 | microglia3 | Vesicle |
| GO:0030666 | endocytic vesicle membrane | 7/135 | 194/19594 | 4E-04 | 0.01937 | 7 | microglia3 | Vesicle |
| GO:0009898 | cytoplasmic side of plasma membrane | 5/135 | 169/19594 | 0.00634 | 0.09454 | 5 | microglia3 | Plasma membrane |
| GO:0030695 | GTPase regulator activity | 16/133 | 488/18410 | 0 | 1E-04 | 16 | microglia3 | GTPase activity |
| GO:0005096 | GTPase activator activity | 11/133 | 274/18410 | 0 | 0.00067 | 11 | microglia3 | GTPase activity |
| GO:0031267 | small GTPase binding | 9/133 | 267/18410 | 0.00014 | 0.00521 | 9 | microglia3 | GTPase activity |
| GO:0051020 | GTPase binding | 9/133 | 298/18410 | 0.00032 | 0.00996 | 9 | microglia3 | GTPase activity |

| | | | | | | | | |
|---|---|---|---|---|---|---|---|---|
| GO:0005095 | GTPase inhibitor activity | 2/133 | 12/18410 | 0.00326 | 0.04844 | 2 | microglia3 | GTPase activity |
| hsa04979 | Cholesterol metabolism | 4/67 | 51/8170 | 0.00077 | 0.04114 | 4 | microglia3 | Cholesterol metabolism |
| GO:0002443 | leukocyte mediated immunity | 24/177 | 457/18800 | 0 | 0 | 24 | microglia4 | Leukocytes |
| GO:0016064 | immunoglobulin mediated immune response | 17/177 | 216/18800 | 0 | 0 | 17 | microglia4 | Immune response |
| GO:0002699 | positive regulation of immune effector process | 16/177 | 248/18800 | 0 | 0 | 16 | microglia4 | Immune response |
| GO:0002449 | lymphocyte mediated immunity | 19/177 | 365/18800 | 0 | 0 | 19 | microglia4 | Lymphocyte |
| GO:0002460 | adaptive immune response based on somatic recombination of immune receptors built from immunoglobulin superfamily domains | 19/177 | 370/18800 | 0 | 0 | 19 | microglia4 | Immune response |
| GO:1903037 | regulation of leukocyte cell-cell adhesion | 18/177 | 344/18800 | 0 | 0 | 18 | microglia4 | Leukocytes |
| GO:0002697 | regulation of immune effector process | 18/177 | 353/18800 | 0 | 0 | 18 | microglia4 | Immune response |
| GO:0002705 | positive regulation of leukocyte mediated immunity | 12/177 | 138/18800 | 0 | 0 | 12 | microglia4 | Leukocytes |
| GO:0070661 | leukocyte proliferation | 17/177 | 330/18800 | 0 | 0 | 17 | microglia4 | Leukocytes |
| GO:0002696 | positive regulation of leukocyte activation | 19/177 | 421/18800 | 0 | 0 | 19 | microglia4 | Leukocytes |
| GO:0070663 | regulation of leukocyte proliferation | 15/177 | 254/18800 | 0 | 0 | 15 | microglia4 | Leukocytes |
| GO:0007159 | leukocyte cell-cell adhesion | 18/177 | 381/18800 | 0 | 0 | 18 | microglia4 | Leukocytes |
| GO:0050863 | regulation of T cell activation | 17/177 | 342/18800 | 0 | 0 | 17 | microglia4 | T cell activity |
| GO:0098883 | synapse pruning | 5/177 | 11/18800 | 0 | 0 | 5 | microglia4 | Synapse assembly |
| GO:0002703 | regulation of leukocyte mediated immunity | 14/177 | 236/18800 | 0 | 0 | 14 | microglia4 | Leukocytes |
| GO:1903039 | positive regulation of leukocyte cell-cell adhesion | 14/177 | 245/18800 | 0 | 0 | 14 | microglia4 | Leukocytes |
| GO:0002253 | activation of immune response | 17/177 | 386/18800 | 0 | 1E-05 | 17 | microglia4 | Immune response |
| GO:0050870 | positive regulation of T cell activation | 13/177 | 223/18800 | 0 | 1E-05 | 13 | microglia4 | T cell activity |
| GO:0050670 | regulation of lymphocyte proliferation | 13/177 | 230/18800 | 0 | 1E-05 | 13 | microglia4 | Lymphocyte |
| GO:0001916 | positive regulation of T cell mediated cytotoxicity | 6/177 | 30/18800 | 0 | 1E-05 | 6 | microglia4 | T cell activity |
| GO:0022407 | regulation of cell-cell adhesion | 18/177 | 456/18800 | 0 | 1E-05 | 18 | microglia4 | Cell adhesion |
| GO:0022409 | positive regulation of cell-cell adhesion | 14/177 | 291/18800 | 0 | 3E-05 | 14 | microglia4 | Cell adhesion |
| GO:0046651 | lymphocyte proliferation | 14/177 | 296/18800 | 0 | 3E-05 | 14 | microglia4 | Lymphocyte |
| GO:0002824 | positive regulation of adaptive immune response based on somatic recombination of immune receptors built from | 9/177 | 110/18800 | 0 | 4E-05 | 9 | microglia4 | Immune response |

| | | | | | | | | |
|---|---|---|---|---|---|---|---|---|
| | immunoglobulin superfamily domains | | | | | | | |
| GO:0001912 | positive regulation of leukocyte mediated cytotoxicity | 7/177 | 58/18800 | 0 | 4E-05 | 7 | microglia4 | Leukocytes |
| GO:0002711 | positive regulation of T cell mediated immunity | 7/177 | 59/18800 | 0 | 5E-05 | 7 | microglia4 | T cell activity |
| GO:0002821 | positive regulation of adaptive immune response | 9/177 | 115/18800 | 0 | 5E-05 | 9 | microglia4 | Immune response |
| GO:0002708 | positive regulation of lymphocyte mediated immunity | 9/177 | 117/18800 | 0 | 6E-05 | 9 | microglia4 | Lymphocyte |
| GO:0001914 | regulation of T cell mediated cytotoxicity | 6/177 | 40/18800 | 0 | 6E-05 | 6 | microglia4 | T cell activity |
| GO:0051251 | positive regulation of lymphocyte activation | 15/177 | 371/18800 | 0 | 8E-05 | 15 | microglia4 | Lymphocyte |
| GO:0070665 | positive regulation of leukocyte proliferation | 10/177 | 158/18800 | 0 | 8E-05 | 10 | microglia4 | Leukocytes |
| GO:0002683 | negative regulation of immune system process | 16/177 | 425/18800 | 0 | 9E-05 | 16 | microglia4 | Immune response |
| GO:0002720 | positive regulation of cytokine production involved in immune response | 7/177 | 68/18800 | 0 | 0.00011 | 7 | microglia4 | Immune response |
| GO:0042098 | T cell proliferation | 11/177 | 204/18800 | 0 | 0.00011 | 11 | microglia4 | T cell activity |
| GO:0002718 | regulation of cytokine production involved in immune response | 8/177 | 98/18800 | 0 | 0.00012 | 8 | microglia4 | Immune response |
| GO:0001909 | leukocyte mediated cytotoxicity | 9/177 | 131/18800 | 0 | 0.00012 | 9 | microglia4 | Leukocytes |
| GO:0002367 | cytokine production involved in immune response | 8/177 | 100/18800 | 0 | 0.00013 | 8 | microglia4 | Immune response |
| GO:0042129 | regulation of T cell proliferation | 10/177 | 174/18800 | 1E-05 | 0.00017 | 10 | microglia4 | T cell activity |
| GO:0001913 | T cell mediated cytotoxicity | 6/177 | 50/18800 | 1E-05 | 0.00019 | 6 | microglia4 | T cell activity |
| GO:0050671 | positive regulation of lymphocyte proliferation | 9/177 | 141/18800 | 1E-05 | 2E-04 | 9 | microglia4 | Lymphocyte |
| GO:0002440 | production of molecular mediator of immune response | 13/177 | 312/18800 | 1E-05 | 0.00022 | 13 | microglia4 | Immune response |
| GO:0006959 | humoral immune response | 13/177 | 317/18800 | 1E-05 | 0.00025 | 13 | microglia4 | Immune response |
| GO:0046034 | ATP metabolic process | 12/177 | 273/18800 | 1E-05 | 0.00027 | 12 | microglia4 | Energy production |
| GO:0045088 | regulation of innate immune response | 11/177 | 231/18800 | 1E-05 | 0.00029 | 11 | microglia4 | Immune response |
| GO:0002274 | myeloid leukocyte activation | 11/177 | 232/18800 | 1E-05 | 0.00029 | 11 | microglia4 | Leukocytes |
| GO:0002764 | immune response-regulating signaling pathway | 16/177 | 482/18800 | 1E-05 | 0.00031 | 16 | microglia4 | Immune response |
| GO:0001910 | regulation of leukocyte mediated cytotoxicity | 7/177 | 85/18800 | 2E-05 | 0.00035 | 7 | microglia4 | Leukocytes |
| GO:0002709 | regulation of T cell mediated immunity | 7/177 | 88/18800 | 2E-05 | 0.00043 | 7 | microglia4 | T cell activity |

| | | | | | | | | |
|---|---|---|---|---|---|---|---|---|
| GO:0045785 | positive regulation of cell adhesion | 15/177 | 446/18800 | 2E-05 | 0.00047 | 15 | microglia4 | Cell adhesion |
| GO:0002888 | positive regulation of myeloid leukocyte mediated immunity | 4/177 | 19/18800 | 3E-05 | 0.00055 | 4 | microglia4 | Leukocytes |
| GO:0032930 | positive regulation of superoxide anion generation | 4/177 | 19/18800 | 3E-05 | 0.00055 | 4 | microglia4 | Oxidative stress |
| GO:0050852 | T cell receptor signaling pathway | 8/177 | 127/18800 | 3E-05 | 0.00056 | 8 | microglia4 | T cell activity |
| GO:0006457 | protein folding | 10/177 | 212/18800 | 3E-05 | 0.00068 | 10 | microglia4 | Unfolded Protein Response |
| GO:2001242 | regulation of intrinsic apoptotic signaling pathway | 9/177 | 171/18800 | 4E-05 | 0.00071 | 9 | microglia4 | Apoptosis |
| GO:0002822 | regulation of adaptive immune response based on somatic recombination of immune receptors built from immunoglobulin superfamily domains | 9/177 | 173/18800 | 4E-05 | 0.00075 | 9 | microglia4 | Immune response |
| GO:0002706 | regulation of lymphocyte mediated immunity | 9/177 | 175/18800 | 4E-05 | 0.00081 | 9 | microglia4 | Lymphocyte |
| GO:0032928 | regulation of superoxide anion generation | 4/177 | 22/18800 | 5E-05 | 9E-04 | 4 | microglia4 | Oxidative stress |
| GO:0042102 | positive regulation of T cell proliferation | 7/177 | 103/18800 | 5E-05 | 0.00099 | 7 | microglia4 | T cell activity |
| GO:0002444 | myeloid leukocyte mediated immunity | 7/177 | 104/18800 | 6E-05 | 0.00102 | 7 | microglia4 | Leukocytes |
| GO:0002381 | immunoglobulin production involved in immunoglobulin-mediated immune response | 6/177 | 73/18800 | 6E-05 | 0.00112 | 6 | microglia4 | Immune response |
| GO:0045824 | negative regulation of innate immune response | 6/177 | 73/18800 | 6E-05 | 0.00112 | 6 | microglia4 | Immune response |
| GO:0002768 | immune response-regulating cell surface receptor signaling pathway | 12/177 | 328/18800 | 7E-05 | 0.00115 | 12 | microglia4 | Immune response |
| GO:0002819 | regulation of adaptive immune response | 9/177 | 188/18800 | 7E-05 | 0.00126 | 9 | microglia4 | Immune response |
| GO:1902105 | regulation of leukocyte differentiation | 11/177 | 283/18800 | 8E-05 | 0.00132 | 11 | microglia4 | Leukocytes |
| GO:0002366 | leukocyte activation involved in immune response | 11/177 | 285/18800 | 8E-05 | 0.00138 | 11 | microglia4 | Leukocytes |
| GO:0002456 | T cell mediated immunity | 7/177 | 112/18800 | 9E-05 | 0.00149 | 7 | microglia4 | T cell activity |
| GO:0002263 | cell activation involved in immune response | 11/177 | 289/18800 | 1E-04 | 0.00154 | 11 | microglia4 | Immune response |
| GO:0002269 | leukocyte activation involved in inflammatory response | 5/177 | 50/18800 | 0.00011 | 0.00167 | 5 | microglia4 | Leukocytes |
| GO:0097193 | intrinsic apoptotic signaling pathway | 11/177 | 295/18800 | 0.00011 | 0.00179 | 11 | microglia4 | Apoptosis |
| GO:0002761 | regulation of myeloid leukocyte differentiation | 7/177 | 117/18800 | 0.00012 | 0.00185 | 7 | microglia4 | Leukocytes |
| GO:0034975 | protein folding in endoplasmic reticulum | 3/177 | 11/18800 | 0.00013 | 0.00195 | 3 | microglia4 | Unfolded Protein Response |
| GO:0002429 | immune response-activating cell surface receptor signaling pathway | 11/177 | 300/18800 | 0.00013 | 0.00196 | 11 | microglia4 | Immune response |

| | | | | | | | | |
|---|---|---|---|---|---|---|---|---|
| GO:0002757 | immune response-activating signal transduction | 11/177 | 300/18800 | 0.00013 | 0.00196 | 11 | microglia4 | Immune response |
| GO:0002702 | positive regulation of production of molecular mediator of immune response | 7/177 | 120/18800 | 0.00014 | 0.00208 | 7 | microglia4 | Immune response |
| GO:0002573 | myeloid leukocyte differentiation | 9/177 | 210/18800 | 0.00017 | 0.00246 | 9 | microglia4 | Leukocytes |
| GO:0002700 | regulation of production of molecular mediator of immune response | 8/177 | 167/18800 | 0.00019 | 0.00262 | 8 | microglia4 | Immune response |
| GO:0002886 | regulation of myeloid leukocyte mediated immunity | 5/177 | 59/18800 | 0.00023 | 0.00311 | 5 | microglia4 | Leukocytes |
| GO:0002275 | myeloid cell activation involved in immune response | 6/177 | 93/18800 | 0.00025 | 0.00329 | 6 | microglia4 | Immune response |
| GO:0090322 | regulation of superoxide metabolic process | 4/177 | 34/18800 | 0.00028 | 0.00367 | 4 | microglia4 | Oxidative stress |
| GO:0050777 | negative regulation of immune response | 8/177 | 179/18800 | 3E-04 | 0.00387 | 8 | microglia4 | Immune response |
| GO:0046635 | positive regulation of alpha-beta T cell activation | 5/177 | 69/18800 | 0.00048 | 0.00569 | 5 | microglia4 | T cell activity |
| GO:0061077 | chaperone-mediated protein folding | 5/177 | 70/18800 | 0.00052 | 0.00593 | 5 | microglia4 | Unfolded Protein Response |
| GO:0002281 | macrophage activation involved in immune response | 3/177 | 18/18800 | 6E-04 | 0.00685 | 3 | microglia4 | Immune response |
| GO:0002698 | negative regulation of immune effector process | 6/177 | 112/18800 | 0.00067 | 0.00752 | 6 | microglia4 | Immune response |
| GO:0042554 | superoxide anion generation | 4/177 | 43/18800 | 7E-04 | 0.00779 | 4 | microglia4 | Oxidative stress |
| GO:0002283 | neutrophil activation involved in immune response | 3/177 | 19/18800 | 0.00071 | 0.00781 | 3 | microglia4 | Immune response |
| GO:0050856 | regulation of T cell receptor signaling pathway | 4/177 | 44/18800 | 0.00077 | 0.00837 | 4 | microglia4 | T cell activity |
| GO:0002286 | T cell activation involved in immune response | 6/177 | 116/18800 | 0.00081 | 0.00872 | 6 | microglia4 | T cell activity |
| GO:0045591 | positive regulation of regulatory T cell differentiation | 3/177 | 20/18800 | 0.00083 | 0.00881 | 3 | microglia4 | T cell activity |
| GO:0051250 | negative regulation of lymphocyte activation | 7/177 | 161/18800 | 0.00084 | 0.00884 | 7 | microglia4 | Lymphocyte |
| GO:0002455 | humoral immune response mediated by circulating immunoglobulin | 6/177 | 121/18800 | 0.00101 | 0.01022 | 6 | microglia4 | Immune response |
| GO:0061081 | positive regulation of myeloid leukocyte cytokine production involved in immune response | 3/177 | 22/18800 | 0.00111 | 0.01117 | 3 | microglia4 | Leukocytes |
| GO:0050868 | negative regulation of T cell activation | 6/177 | 125/18800 | 0.00119 | 0.01191 | 6 | microglia4 | T cell activity |
| GO:0046641 | positive regulation of alpha-beta T cell proliferation | 3/177 | 23/18800 | 0.00126 | 0.01252 | 3 | microglia4 | T cell activity |
| GO:0097529 | myeloid leukocyte migration | 8/177 | 229/18800 | 0.0015 | 0.01458 | 8 | microglia4 | Leukocytes |
| GO:0001911 | negative regulation of leukocyte mediated cytotoxicity | 3/177 | 25/18800 | 0.00162 | 0.01542 | 3 | microglia4 | Leukocytes |

| | | | | | | | | |
|---|---|---|---|---|---|---|---|---|
| GO:0070664 | negative regulation of leukocyte proliferation | 5/177 | 91/18800 | 0.00169 | 0.01599 | 5 | microglia4 | Leukocytes |
| GO:0002763 | positive regulation of myeloid leukocyte differentiation | 4/177 | 55/18800 | 0.00178 | 0.01678 | 4 | microglia4 | Leukocytes |
| GO:0002726 | positive regulation of T cell cytokine production | 3/177 | 26/18800 | 0.00182 | 0.01701 | 3 | microglia4 | T cell activity |
| GO:0006986 | response to unfolded protein | 6/177 | 139/18800 | 0.00205 | 0.01862 | 6 | microglia4 | Unfolded Protein Response |
| GO:0006826 | iron ion transport | 4/177 | 59/18800 | 0.00231 | 0.02033 | 4 | microglia4 | Ion transport |
| GO:0002695 | negative regulation of leukocyte activation | 7/177 | 193/18800 | 0.00238 | 0.02092 | 7 | microglia4 | Leukocytes |
| GO:1903038 | negative regulation of leukocyte cell-cell adhesion | 6/177 | 144/18800 | 0.00244 | 0.02126 | 6 | microglia4 | Leukocytes |
| GO:0061082 | myeloid leukocyte cytokine production | 3/177 | 29/18800 | 0.0025 | 0.02149 | 3 | microglia4 | Leukocytes |
| GO:2001244 | positive regulation of intrinsic apoptotic signaling pathway | 4/177 | 61/18800 | 0.00261 | 0.02226 | 4 | microglia4 | Apoptosis |
| GO:2001233 | regulation of apoptotic signaling pathway | 10/177 | 370/18800 | 0.00272 | 0.02298 | 10 | microglia4 | Apoptosis |
| GO:1900101 | regulation of endoplasmic reticulum unfolded protein response | 3/177 | 30/18800 | 0.00276 | 0.02325 | 3 | microglia4 | Unfolded Protein Response |
| GO:0002285 | lymphocyte activation involved in immune response | 7/177 | 201/18800 | 0.00299 | 0.02472 | 7 | microglia4 | Lymphocyte |
| GO:0046634 | regulation of alpha-beta T cell activation | 5/177 | 105/18800 | 0.00316 | 0.02591 | 5 | microglia4 | T cell activity |
| GO:0051085 | chaperone cofactor-dependent protein refolding | 3/177 | 32/18800 | 0.00333 | 0.02712 | 3 | microglia4 | Unfolded Protein Response |
| GO:0030217 | T cell differentiation | 8/177 | 263/18800 | 0.00353 | 0.02832 | 8 | microglia4 | T cell activity |
| GO:0045589 | regulation of regulatory T cell differentiation | 3/177 | 33/18800 | 0.00364 | 0.02884 | 3 | microglia4 | T cell activity |
| GO:0002704 | negative regulation of leukocyte mediated immunity | 4/177 | 67/18800 | 0.00367 | 0.02891 | 4 | microglia4 | Leukocytes |
| GO:1902107 | positive regulation of leukocyte differentiation | 6/177 | 157/18800 | 0.00375 | 0.02891 | 6 | microglia4 | Leukocytes |
| GO:2000425 | regulation of apoptotic cell clearance | 2/177 | 10/18800 | 0.00377 | 0.02891 | 2 | microglia4 | Apoptosis |
| GO:0033344 | cholesterol efflux | 4/177 | 69/18800 | 0.00408 | 0.0309 | 4 | microglia4 | Cholesterol metabolism |
| GO:0035966 | response to topologically incorrect protein | 6/177 | 160/18800 | 0.00411 | 0.03105 | 6 | microglia4 | Unfolded Protein Response |
| GO:0006122 | mitochondrial electron transport, ubiquinol to cytochrome c | 2/177 | 11/18800 | 0.00458 | 0.03351 | 2 | microglia4 | Mitochondrial changes |
| GO:1903897 | regulation of PERK-mediated unfolded protein response | 2/177 | 11/18800 | 0.00458 | 0.03351 | 2 | microglia4 | Unfolded Protein Response |
| GO:0045066 | regulatory T cell differentiation | 3/177 | 36/18800 | 0.00466 | 0.03381 | 3 | microglia4 | T cell activity |
| GO:0006801 | superoxide metabolic process | 4/177 | 72/18800 | 0.00475 | 0.03426 | 4 | microglia4 | Oxidative stress |
| GO:0046640 | regulation of alpha-beta T cell proliferation | 3/177 | 37/18800 | 0.00504 | 0.03597 | 3 | microglia4 | T cell activity |

| | | | | | | | | |
|---|---|---|---|---|---|---|---|---|
| GO:0051084 | 'de novo' post-translational protein folding | 3/177 | 37/18800 | 0.00504 | 0.03597 | 3 | microglia4 | Unfolded Protein Response |
| GO:0002369 | T cell cytokine production | 3/177 | 38/18800 | 0.00543 | 0.03732 | 3 | microglia4 | T cell activity |
| GO:0002724 | regulation of T cell cytokine production | 3/177 | 38/18800 | 0.00543 | 0.03732 | 3 | microglia4 | T cell activity |
| GO:1902229 | regulation of intrinsic apoptotic signaling pathway in response to DNA damage | 3/177 | 38/18800 | 0.00543 | 0.03732 | 3 | microglia4 | Apoptosis |
| GO:0002424 | T cell mediated immune response to tumor cell | 2/177 | 12/18800 | 0.00547 | 0.03732 | 2 | microglia4 | T cell activity |
| GO:0043380 | regulation of memory T cell differentiation | 2/177 | 12/18800 | 0.00547 | 0.03732 | 2 | microglia4 | T cell activity |
| GO:0044650 | adhesion of symbiont to host cell | 2/177 | 12/18800 | 0.00547 | 0.03732 | 2 | microglia4 | Cell adhesion |
| GO:0140052 | cellular response to oxidised low-density lipoprotein particle stimulus | 2/177 | 12/18800 | 0.00547 | 0.03732 | 2 | microglia4 | Oxidative stress |
| GO:2000516 | positive regulation of CD4-positive, alpha-beta T cell activation | 3/177 | 39/18800 | 0.00585 | 0.03935 | 3 | microglia4 | T cell activity |
| GO:0046633 | alpha-beta T cell proliferation | 3/177 | 40/18800 | 0.00628 | 0.04183 | 3 | microglia4 | T cell activity |
| GO:0033077 | T cell differentiation in thymus | 4/177 | 78/18800 | 0.00631 | 0.04191 | 4 | microglia4 | T cell activity |
| GO:0043379 | memory T cell differentiation | 2/177 | 13/18800 | 0.00642 | 0.04197 | 2 | microglia4 | T cell activity |
| GO:0050808 | synapse organization | 10/177 | 419/18800 | 0.00647 | 0.04219 | 10 | microglia4 | Synapse assembly |
| GO:0030301 | cholesterol transport | 4/177 | 79/18800 | 0.00659 | 0.04289 | 4 | microglia4 | Cholesterol metabolism |
| GO:0002891 | positive regulation of immunoglobulin mediated immune response | 3/177 | 41/18800 | 0.00673 | 0.04313 | 3 | microglia4 | Immune response |
| GO:0006458 | 'de novo' protein folding | 3/177 | 41/18800 | 0.00673 | 0.04313 | 3 | microglia4 | Unfolded Protein Response |
| GO:0030595 | leukocyte chemotaxis | 7/177 | 236/18800 | 0.00712 | 0.04553 | 7 | microglia4 | Leukocytes |
| GO:0010872 | regulation of cholesterol esterification | 2/177 | 14/18800 | 0.00745 | 0.04632 | 2 | microglia4 | Cholesterol metabolism |
| GO:0045059 | positive thymic T cell selection | 2/177 | 14/18800 | 0.00745 | 0.04632 | 2 | microglia4 | T cell activity |
| GO:0051402 | neuron apoptotic process | 7/177 | 241/18800 | 0.00794 | 0.04886 | 7 | microglia4 | Apoptosis |
| GO:0050672 | negative regulation of lymphocyte proliferation | 4/177 | 84/18800 | 0.00817 | 0.05 | 4 | microglia4 | Lymphocyte |
| GO:0055072 | iron ion homeostasis | 4/177 | 85/18800 | 0.00851 | 0.05067 | 4 | microglia4 | Ion transport |
| GO:0044406 | adhesion of symbiont to host | 2/177 | 15/18800 | 0.00854 | 0.05067 | 2 | microglia4 | Cell adhesion |
| GO:1900102 | negative regulation of endoplasmic reticulum unfolded protein response | 2/177 | 15/18800 | 0.00854 | 0.05067 | 2 | microglia4 | Unfolded Protein Response |
| GO:0002920 | regulation of humoral immune response | 3/177 | 45/18800 | 0.00871 | 0.05159 | 3 | microglia4 | Immune response |
| GO:0050862 | positive regulation of T cell receptor signaling pathway | 2/177 | 16/18800 | 0.0097 | 0.05549 | 2 | microglia4 | T cell activity |
| GO:1903578 | regulation of ATP metabolic process | 4/177 | 89/18800 | 0.00997 | 0.05621 | 4 | microglia4 | Energy production |

| | | | | | | | | |
|---|---|---|---|---|---|---|---|---|
| GO:0006119 | oxidative phosphorylation | 5/177 | 139/18800 | 0.01021 | 0.05733 | 5 | microglia4 | Oxidative stress |
| GO:0055076 | transition metal ion homeostasis | 5/177 | 139/18800 | 0.01021 | 0.05733 | 5 | microglia4 | Ion transport |
| GO:0030098 | lymphocyte differentiation | 9/177 | 382/18800 | 0.01039 | 0.05821 | 9 | microglia4 | Lymphocyte |
| GO:0050900 | leukocyte migration | 9/177 | 384/18800 | 0.01072 | 0.05993 | 9 | microglia4 | Leukocytes |
| GO:0002830 | positive regulation of type 2 immune response | 2/177 | 17/18800 | 0.01092 | 0.06059 | 2 | microglia4 | Immune response |
| GO:0042775 | mitochondrial ATP synthesis coupled electron transport | 4/177 | 92/18800 | 0.01116 | 0.06129 | 4 | microglia4 | Mitochondrial changes |
| GO:0022408 | negative regulation of cell-cell adhesion | 6/177 | 199/18800 | 0.01151 | 0.06294 | 6 | microglia4 | Cell adhesion |
| GO:0045582 | positive regulation of T cell differentiation | 4/177 | 94/18800 | 0.01201 | 0.06489 | 4 | microglia4 | T cell activity |
| GO:0006123 | mitochondrial electron transport, cytochrome c to oxygen | 2/177 | 18/18800 | 0.01221 | 0.06495 | 2 | microglia4 | Mitochondrial changes |
| GO:0034435 | cholesterol esterification | 2/177 | 18/18800 | 0.01221 | 0.06495 | 2 | microglia4 | Cholesterol metabolism |
| GO:0036499 | PERK-mediated unfolded protein response | 2/177 | 18/18800 | 0.01221 | 0.06495 | 2 | microglia4 | Unfolded Protein Response |
| GO:0043277 | apoptotic cell clearance | 3/177 | 51/18800 | 0.01227 | 0.06514 | 3 | microglia4 | Apoptosis |
| GO:0051900 | regulation of mitochondrial depolarization | 2/177 | 19/18800 | 0.01357 | 0.07089 | 2 | microglia4 | Mitochondrial changes |
| GO:0098780 | response to mitochondrial depolarisation | 2/177 | 19/18800 | 0.01357 | 0.07089 | 2 | microglia4 | Mitochondrial changes |
| GO:2001185 | regulation of CD8-positive, alpha-beta T cell activation | 2/177 | 19/18800 | 0.01357 | 0.07089 | 2 | microglia4 | T cell activity |
| GO:0034620 | cellular response to unfolded protein | 4/177 | 98/18800 | 0.01382 | 0.07165 | 4 | microglia4 | Unfolded Protein Response |
| GO:0045580 | regulation of T cell differentiation | 5/177 | 151/18800 | 0.01421 | 0.07341 | 5 | microglia4 | T cell activity |
| GO:0010506 | regulation of autophagy | 8/177 | 336/18800 | 0.01447 | 0.07459 | 8 | microglia4 | Autophagy |
| GO:0098869 | cellular oxidant detoxification | 4/177 | 100/18800 | 0.01479 | 0.07609 | 4 | microglia4 | Oxidative stress |
| GO:0002707 | negative regulation of lymphocyte mediated immunity | 3/177 | 55/18800 | 0.01505 | 0.07613 | 3 | microglia4 | Lymphocyte |
| GO:0000041 | transition metal ion transport | 4/177 | 101/18800 | 0.01529 | 0.07694 | 4 | microglia4 | Ion transport |
| GO:0008630 | intrinsic apoptotic signaling pathway in response to DNA damage | 4/177 | 101/18800 | 0.01529 | 0.07694 | 4 | microglia4 | Apoptosis |
| GO:1902106 | negative regulation of leukocyte differentiation | 4/177 | 102/18800 | 0.0158 | 0.07892 | 4 | microglia4 | Leukocytes |
| GO:2001243 | negative regulation of intrinsic apoptotic signaling pathway | 4/177 | 102/18800 | 0.0158 | 0.07892 | 4 | microglia4 | Apoptosis |
| GO:0051882 | mitochondrial depolarization | 2/177 | 21/18800 | 0.01646 | 0.08101 | 2 | microglia4 | Mitochondrial changes |
| GO:0046631 | alpha-beta T cell activation | 5/177 | 158/18800 | 0.01697 | 0.08292 | 5 | microglia4 | T cell activity |
| GO:0045061 | thymic T cell selection | 2/177 | 22/18800 | 0.01799 | 0.08576 | 2 | microglia4 | T cell activity |
| GO:0045621 | positive regulation of lymphocyte differentiation | 4/177 | 107/18800 | 0.01851 | 0.08763 | 4 | microglia4 | Lymphocyte |

| | | | | | | | | |
|---|---|---|---|---|---|---|---|---|
| GO:0099173 | postsynapse organization | 5/177 | 163/18800 | 0.01915 | 0.09033 | 5 | microglia4 | Synapse assembly |
| GO:0045428 | regulation of nitric oxide biosynthetic process | 3/177 | 61/18800 | 0.01982 | 0.09202 | 3 | microglia4 | Oxidative stress |
| GO:0002889 | regulation of immunoglobulin mediated immune response | 3/177 | 62/18800 | 0.02068 | 95 | 3 | microglia4 | Immune response |
| GO:0002922 | positive regulation of humoral immune response | 2/177 | 24/18800 | 0.02124 | 0.09535 | 2 | microglia4 | Immune response |
| GO:0070059 | intrinsic apoptotic signaling pathway in response to endoplasmic reticulum stress | 3/177 | 63/18800 | 0.02157 | 0.09606 | 3 | microglia4 | Apoptosis |
| GO:0006979 | response to oxidative stress | 9/177 | 433/18800 | 0.02172 | 0.09639 | 9 | microglia4 | Oxidative stress |
| GO:0080164 | regulation of nitric oxide metabolic process | 3/177 | 64/18800 | 0.02248 | 0.09873 | 3 | microglia4 | Oxidative stress |
| GO:0043302 | positive regulation of leukocyte degranulation | 2/177 | 25/18800 | 0.02294 | 0.09939 | 2 | microglia4 | Leukocytes |
| GO:0050860 | negative regulation of T cell receptor signaling pathway | 2/177 | 25/18800 | 0.02294 | 0.09939 | 2 | microglia4 | T cell activity |
| GO:0005925 | focal adhesion | 52/180 | 419/19594 | 0 | 0 | 52 | microglia4 | Cell adhesion |
| GO:0030666 | endocytic vesicle membrane | 20/180 | 194/19594 | 0 | 0 | 20 | microglia4 | Vesicle |
| GO:0030139 | endocytic vesicle | 21/180 | 342/19594 | 0 | 0 | 21 | microglia4 | Vesicle |
| GO:0012507 | ER to Golgi transport vesicle membrane | 11/180 | 62/19594 | 0 | 0 | 11 | microglia4 | Vesicle |
| GO:0030662 | coated vesicle membrane | 14/180 | 176/19594 | 0 | 0 | 14 | microglia4 | Vesicle |
| GO:0030134 | COPII-coated ER to Golgi transport vesicle | 11/180 | 94/19594 | 0 | 0 | 11 | microglia4 | Vesicle |
| GO:0030669 | clathrin-coated endocytic vesicle membrane | 9/180 | 72/19594 | 0 | 0 | 9 | microglia4 | Vesicle |
| GO:0030670 | phagocytic vesicle membrane | 9/180 | 77/19594 | 0 | 0 | 9 | microglia4 | Vesicle |
| GO:0032279 | asymmetric synapse | 16/180 | 323/19594 | 0 | 0 | 16 | microglia4 | Synapse assembly |
| GO:0098984 | neuron to neuron synapse | 16/180 | 347/19594 | 0 | 0 | 16 | microglia4 | Synapse assembly |
| GO:0045334 | clathrin-coated endocytic vesicle | 9/180 | 91/19594 | 0 | 0 | 9 | microglia4 | Vesicle |
| GO:0030135 | coated vesicle | 14/180 | 290/19594 | 0 | 0 | 14 | microglia4 | Vesicle |
| GO:0030665 | clathrin-coated vesicle membrane | 9/180 | 111/19594 | 0 | 1E-05 | 9 | microglia4 | Vesicle |
| GO:0030658 | transport vesicle membrane | 11/180 | 205/19594 | 0 | 3E-05 | 11 | microglia4 | Vesicle |
| GO:0045335 | phagocytic vesicle | 9/180 | 138/19594 | 1E-05 | 4E-05 | 9 | microglia4 | Vesicle |
| GO:0005741 | mitochondrial outer membrane | 10/180 | 205/19594 | 2E-05 | 0.00014 | 10 | microglia4 | Mitochondrial changes |
| GO:0030136 | clathrin-coated vesicle | 9/180 | 192/19594 | 7E-05 | 0.00044 | 9 | microglia4 | Vesicle |
| GO:0098800 | inner mitochondrial membrane protein complex | 8/180 | 155/19594 | 9E-05 | 0.00056 | 8 | microglia4 | Mitochondrial changes |
| GO:0060205 | cytoplasmic vesicle lumen | 11/180 | 325/19594 | 0.00022 | 0.00118 | 11 | microglia4 | Vesicle |
| GO:0031983 | vesicle lumen | 11/180 | 327/19594 | 0.00023 | 0.00122 | 11 | microglia4 | Vesicle |

| | | | | | | | | |
|---|---|---|---|---|---|---|---|---|
| GO:0098554 | cytoplasmic side of endoplasmic reticulum membrane | 3/180 | 14/19594 | 0.00026 | 0.00134 | 3 | microglia4 | Plasma membrane |
| GO:0030133 | transport vesicle | 12/180 | 402/19594 | 0.00036 | 0.00175 | 12 | microglia4 | Vesicle |
| GO:0005742 | mitochondrial outer membrane translocase complex | 3/180 | 21/19594 | 9E-04 | 0.00394 | 3 | microglia4 | Mitochondrial changes |
| GO:0098799 | outer mitochondrial membrane protein complex | 3/180 | 22/19594 | 0.00103 | 0.00447 | 3 | microglia4 | Mitochondrial changes |
| GO:0071682 | endocytic vesicle lumen | 3/180 | 23/19594 | 0.00118 | 0.00496 | 3 | microglia4 | Vesicle |
| GO:0098798 | mitochondrial protein-containing complex | 9/180 | 281/19594 | 0.00119 | 0.00496 | 9 | microglia4 | Mitochondrial changes |
| GO:0005750 | mitochondrial respiratory chain complex III | 2/180 | 14/19594 | 0.0071 | 0.02267 | 2 | microglia4 | Mitochondrial changes |
| GO:0001772 | immunological synapse | 3/180 | 44/19594 | 0.00766 | 0.02421 | 3 | microglia4 | Synapse assembly |
| GO:0005746 | mitochondrial respirasome | 4/180 | 94/19594 | 0.01106 | 0.03267 | 4 | microglia4 | Mitochondrial changes |
| GO:0005753 | mitochondrial proton-transporting ATP synthase complex | 2/180 | 21/19594 | 0.01572 | 0.04515 | 2 | microglia4 | Mitochondrial changes |
| GO:0005751 | mitochondrial respiratory chain complex IV | 2/180 | 25/19594 | 0.02192 | 0.06076 | 2 | microglia4 | Mitochondrial changes |
| GO:0098562 | cytoplasmic side of membrane | 5/180 | 193/19594 | 0.03301 | 0.08755 | 5 | microglia4 | Plasma membrane |
| GO:0005743 | mitochondrial inner membrane | 9/180 | 491/19594 | 0.03796 | 0.09753 | 9 | microglia4 | Mitochondrial changes |
| GO:0042608 | T cell receptor binding | 4/180 | 10/18410 | 0 | 6E-05 | 4 | microglia4 | T cell activity |
| GO:0044183 | protein folding chaperone | 6/180 | 43/18410 | 0 | 1E-04 | 6 | microglia4 | Unfolded Protein Response |
| GO:0051082 | unfolded protein binding | 8/180 | 121/18410 | 3E-05 | 0.00055 | 8 | microglia4 | Unfolded Protein Response |
| GO:0140375 | immune receptor activity | 8/180 | 148/18410 | 0.00011 | 0.00185 | 8 | microglia4 | Immune response |
| GO:0120020 | cholesterol transfer activity | 3/180 | 22/18410 | 0.00123 | 0.01627 | 3 | microglia4 | Cholesterol metabolism |
| GO:0008199 | ferric iron binding | 2/180 | 11/18410 | 0.00493 | 0.05057 | 2 | microglia4 | Iron transport |
| GO:0050998 | nitric-oxide synthase binding | 2/180 | 13/18410 | 0.00691 | 0.06536 | 2 | microglia4 | Oxidative stress |
| GO:0016209 | antioxidant activity | 4/180 | 85/18410 | 0.00968 | 0.08306 | 4 | microglia4 | Oxidative stress |
| hsa05320 | Autoimmune thyroid disease | 9/154 | 53/8170 | 0 | 1E-05 | 9 | microglia4 | Immune response |
| hsa05166 | Human T-cell leukemia virus 1 infection | 14/154 | 222/8170 | 7E-05 | 0.00084 | 14 | microglia4 | T cell activity |
| hsa04514 | Cell adhesion molecules | 10/154 | 157/8170 | 0.00074 | 0.00715 | 10 | microglia4 | Cell adhesion |
| hsa04672 | Intestinal immune network for IgA production | 5/154 | 49/8170 | 0.00218 | 0.01649 | 5 | microglia4 | Immune response |
| hsa00190 | Oxidative phosphorylation | 7/154 | 134/8170 | 0.0132 | 0.06958 | 7 | microglia4 | Oxidative stress |
| hsa04979 | Cholesterol metabolism | 4/154 | 51/8170 | 0.01527 | 0.07816 | 4 | microglia4 | Cholesterol metabolism |
| GO:0050808 | synapse organization | 30/301 | 419/18800 | 0 | 0 | 30 | microglia5 | Synapse assembly |

| | | | | | | | | |
|---|---|---|---|---|---|---|---|---|
| GO:0007409 | axonogenesis | 29/301 | 430/18800 | 0 | 0 | 29 | microglia5 | Axonogenesis |
| GO:0099173 | postsynapse organization | 15/301 | 163/18800 | 0 | 3E-05 | 15 | microglia5 | Synapse assembly |
| GO:0002028 | regulation of sodium ion transport | 11/301 | 90/18800 | 0 | 8E-05 | 11 | microglia5 | Ion transport |
| GO:0042552 | myelination | 13/301 | 138/18800 | 0 | 0.00011 | 13 | microglia5 | Enhanced myelination |
| GO:0007272 | ensheathment of neurons | 13/301 | 140/18800 | 0 | 0.00011 | 13 | microglia5 | Enhanced myelination |
| GO:0008366 | axon ensheathment | 13/301 | 140/18800 | 0 | 0.00011 | 13 | microglia5 | Enhanced myelination |
| GO:0007416 | synapse assembly | 14/301 | 180/18800 | 0 | 0.00029 | 14 | microglia5 | Synapse assembly |
| GO:0050807 | regulation of synapse organization | 15/301 | 209/18800 | 0 | 0.00031 | 15 | microglia5 | Synapse assembly |
| GO:0050803 | regulation of synapse structure or activity | 15/301 | 215/18800 | 0 | 0.00038 | 15 | microglia5 | Synapse assembly |
| GO:0006814 | sodium ion transport | 16/301 | 249/18800 | 0 | 0.00041 | 16 | microglia5 | Ion transport |
| GO:0007158 | neuron cell-cell adhesion | 5/301 | 17/18800 | 1E-05 | 7E-04 | 5 | microglia5 | Cell adhesion |
| GO:0010959 | regulation of metal ion transport | 20/301 | 403/18800 | 1E-05 | 0.00096 | 20 | microglia5 | Ion transport |
| GO:1902305 | regulation of sodium ion transmembrane transport | 8/301 | 67/18800 | 1E-05 | 0.00123 | 8 | microglia5 | Ion transport |
| GO:0010765 | positive regulation of sodium ion transport | 6/301 | 33/18800 | 1E-05 | 0.00123 | 6 | microglia5 | Ion transport |
| GO:0048813 | dendrite morphogenesis | 11/301 | 139/18800 | 2E-05 | 0.00137 | 11 | microglia5 | Morphogenesis |
| GO:0055117 | regulation of cardiac muscle contraction | 8/301 | 77/18800 | 3E-05 | 0.00238 | 8 | microglia5 | Muscle contraction |
| GO:0031346 | positive regulation of cell projection organization | 17/301 | 341/18800 | 4E-05 | 0.00274 | 17 | microglia5 | Cell projection organization |
| GO:0034765 | regulation of ion transmembrane transport | 20/301 | 476/18800 | 9E-05 | 0.00533 | 20 | microglia5 | Ion transport |
| GO:0032412 | regulation of ion transmembrane transporter activity | 14/301 | 263/18800 | 9E-05 | 0.00533 | 14 | microglia5 | Ion transport |
| GO:2000651 | positive regulation of sodium ion transmembrane transporter activity | 4/301 | 16/18800 | 1E-04 | 0.00545 | 4 | microglia5 | Ion transport |
| GO:0051650 | establishment of vesicle localization | 11/301 | 173/18800 | 0.00011 | 0.00576 | 11 | microglia5 | Vesicle |
| GO:0010880 | regulation of release of sequestered calcium ion into cytosol by sarcoplasmic reticulum | 5/301 | 31/18800 | 0.00012 | 0.00589 | 5 | microglia5 | Calcium transport |
| GO:0022604 | regulation of cell morphogenesis | 15/301 | 305/18800 | 0.00013 | 0.00589 | 15 | microglia5 | Morphogenesis |
| GO:0035725 | sodium ion transmembrane transport | 11/301 | 177/18800 | 0.00014 | 0.0061 | 11 | microglia5 | Ion transport |
| GO:1901379 | regulation of potassium ion transmembrane transport | 8/301 | 95/18800 | 0.00014 | 0.00612 | 8 | microglia5 | Ion transport |
| GO:0006942 | regulation of striated muscle contraction | 8/301 | 96/18800 | 0.00015 | 0.00642 | 8 | microglia5 | Muscle contraction |

| | | | | | | | | |
|---|---|---|---|---|---|---|---|---|
| GO:0050770 | regulation of axonogenesis | 10/301 | 154/18800 | 0.00019 | 0.00768 | 10 | microglia5 | Axonogenesis |
| GO:0014808 | release of sequestered calcium ion into cytosol by sarcoplasmic reticulum | 5/301 | 35/18800 | 0.00022 | 0.00841 | 5 | microglia5 | Calcium transport |
| GO:0071805 | potassium ion transmembrane transport | 12/301 | 219/18800 | 0.00022 | 0.00841 | 12 | microglia5 | Ion transport |
| GO:0051648 | vesicle localization | 11/301 | 189/18800 | 0.00024 | 0.00892 | 11 | microglia5 | Vesicle |
| GO:1902307 | positive regulation of sodium ion transmembrane transport | 4/301 | 20/18800 | 0.00025 | 0.00898 | 4 | microglia5 | Ion transport |
| GO:1903514 | release of sequestered calcium ion into cytosol by endoplasmic reticulum | 5/301 | 36/18800 | 0.00026 | 0.00898 | 5 | microglia5 | Calcium transport |
| GO:0042063 | gliogenesis | 14/301 | 291/18800 | 0.00027 | 0.00927 | 14 | microglia5 | Gliogenesis |
| GO:0060042 | retina morphogenesis in camera-type eye | 6/301 | 57/18800 | 0.00029 | 0.01003 | 6 | microglia5 | Morphogenesis |
| GO:2000649 | regulation of sodium ion transmembrane transporter activity | 6/301 | 58/18800 | 0.00032 | 0.01067 | 6 | microglia5 | Ion transport |
| GO:0051056 | regulation of small GTPase mediated signal transduction | 14/301 | 299/18800 | 0.00035 | 0.01117 | 14 | microglia5 | GTPase activity |
| GO:0032288 | myelin assembly | 4/301 | 22/18800 | 0.00038 | 0.01173 | 4 | microglia5 | Enhanced myelination |
| GO:0043266 | regulation of potassium ion transport | 8/301 | 110/18800 | 0.00039 | 0.01194 | 8 | microglia5 | Ion transport |
| GO:0006937 | regulation of muscle contraction | 10/301 | 170/18800 | 0.00043 | 0.01238 | 10 | microglia5 | Muscle contraction |
| GO:0099175 | regulation of postsynapse organization | 7/301 | 86/18800 | 0.00046 | 0.0126 | 7 | microglia5 | Synapse assembly |
| GO:0051965 | positive regulation of synapse assembly | 6/301 | 63/18800 | 0.00051 | 0.01346 | 6 | microglia5 | Synapse assembly |
| GO:0070296 | sarcoplasmic reticulum calcium ion transport | 5/301 | 42/18800 | 0.00053 | 0.01406 | 5 | microglia5 | Calcium transport |
| GO:0070588 | calcium ion transmembrane transport | 14/301 | 314/18800 | 0.00057 | 0.01447 | 14 | microglia5 | Calcium transport |
| GO:0006813 | potassium ion transport | 12/301 | 243/18800 | 0.00057 | 0.01447 | 12 | microglia5 | Ion transport |
| GO:0099560 | synaptic membrane adhesion | 4/301 | 25/18800 | 0.00062 | 0.01526 | 4 | microglia5 | Cell adhesion |
| GO:1901019 | regulation of calcium ion transmembrane transporter activity | 7/301 | 91/18800 | 0.00065 | 0.01558 | 7 | microglia5 | Calcium transport |
| GO:0035418 | protein localization to synapse | 6/301 | 67/18800 | 7E-04 | 0.01641 | 6 | microglia5 | Synapse assembly |
| GO:1901016 | regulation of potassium ion transmembrane transporter activity | 6/301 | 67/18800 | 7E-04 | 0.01641 | 6 | microglia5 | Ion transport |
| GO:0010882 | regulation of cardiac muscle contraction by calcium ion signaling | 4/301 | 26/18800 | 0.00073 | 0.01668 | 4 | microglia5 | Muscle contraction |
| GO:0051057 | positive regulation of small GTPase mediated signal transduction | 6/301 | 70/18800 | 0.00089 | 0.01894 | 6 | microglia5 | GTPase activity |
| GO:0051966 | regulation of synaptic transmission, glutamatergic | 6/301 | 70/18800 | 0.00089 | 0.01894 | 6 | microglia5 | Glutamatergic synapse |

| | | | | | | | | |
|---|---|---|---|---|---|---|---|---|
| GO:0099518 | vesicle cytoskeletal trafficking | 6/301 | 71/18800 | 0.00096 | 0.01993 | 6 | microglia5 | Vesicle |
| GO:0022011 | myelination in peripheral nervous system | 4/301 | 28/18800 | 0.00097 | 0.01993 | 4 | microglia5 | Enhanced myelination |
| GO:0032292 | peripheral nervous system axon ensheathment | 4/301 | 28/18800 | 0.00097 | 0.01993 | 4 | microglia5 | Enhanced myelination |
| GO:0099068 | postsynapse assembly | 4/301 | 29/18800 | 0.00112 | 0.02178 | 4 | microglia5 | Synapse assembly |
| GO:0099054 | presynapse assembly | 5/301 | 50/18800 | 0.0012 | 0.02308 | 5 | microglia5 | Synapse assembly |
| GO:0030007 | cellular potassium ion homeostasis | 3/301 | 14/18800 | 0.0013 | 0.02448 | 3 | microglia5 | Ion transport |
| GO:0007157 | heterophilic cell-cell adhesion via plasma membrane cell adhesion molecules | 5/301 | 51/18800 | 0.00131 | 0.02455 | 5 | microglia5 | Cell adhesion |
| GO:0051963 | regulation of synapse assembly | 7/301 | 103/18800 | 0.00134 | 0.02508 | 7 | microglia5 | Synapse assembly |
| GO:0006892 | post-Golgi vesicle-mediated transport | 7/301 | 104/18800 | 0.00142 | 0.02619 | 7 | microglia5 | Vesicle |
| GO:0099172 | presynapse organization | 5/301 | 52/18800 | 0.00143 | 0.02619 | 5 | microglia5 | Synapse assembly |
| GO:0060560 | developmental growth involved in morphogenesis | 11/301 | 234/18800 | 0.00143 | 0.02619 | 11 | microglia5 | Morphogenesis |
| GO:0043270 | positive regulation of ion transport | 12/301 | 273/18800 | 0.00157 | 0.02778 | 12 | microglia5 | Ion transport |
| GO:0006936 | muscle contraction | 14/301 | 349/18800 | 0.00157 | 0.02778 | 14 | microglia5 | Muscle contraction |
| GO:0060048 | cardiac muscle contraction | 8/301 | 138/18800 | 0.00173 | 0.02925 | 8 | microglia5 | Muscle contraction |
| GO:0061684 | chaperone-mediated autophagy | 3/301 | 16/18800 | 0.00195 | 0.0315 | 3 | microglia5 | Autophagy |
| GO:0097120 | receptor localization to synapse | 5/301 | 56/18800 | 0.00199 | 0.03192 | 5 | microglia5 | Synapse assembly |
| GO:0048041 | focal adhesion assembly | 6/301 | 83/18800 | 0.00215 | 0.03358 | 6 | microglia5 | Cell adhesion |
| GO:0006941 | striated muscle contraction | 9/301 | 178/18800 | 0.00234 | 0.03497 | 9 | microglia5 | Muscle contraction |
| GO:1903580 | positive regulation of ATP metabolic process | 4/301 | 37/18800 | 0.00281 | 0.03998 | 4 | microglia5 | Energy production |
| GO:0030050 | vesicle transport along actin filament | 3/301 | 19/18800 | 0.00326 | 0.04488 | 3 | microglia5 | Vesicle |
| GO:0051893 | regulation of focal adhesion assembly | 5/301 | 63/18800 | 0.00335 | 0.04539 | 5 | microglia5 | Cell adhesion |
| GO:0006816 | calcium ion transport | 15/301 | 424/18800 | 0.00362 | 0.04792 | 15 | microglia5 | Calcium transport |
| GO:1903169 | regulation of calcium ion transmembrane transport | 8/301 | 156/18800 | 0.0037 | 0.04864 | 8 | microglia5 | Calcium transport |
| GO:1902473 | regulation of protein localization to synapse | 3/301 | 20/18800 | 0.00379 | 0.04879 | 3 | microglia5 | Synapse assembly |
| GO:0035249 | synaptic transmission, glutamatergic | 6/301 | 94/18800 | 0.00402 | 0.0512 | 6 | microglia5 | Glutamatergic synapse |
| GO:0099558 | maintenance of synapse structure | 3/301 | 21/18800 | 0.00436 | 0.05439 | 3 | microglia5 | Synapse assembly |

| | | | | | | | | |
|---|---|---|---|---|---|---|---|---|
| GO:0010769 | regulation of cell morphogenesis involved in differentiation | 6/301 | 96/18800 | 0.00446 | 0.05532 | 6 | microglia5 | Morphogenesis |
| GO:0034767 | positive regulation of ion transmembrane transport | 8/301 | 163/18800 | 0.00482 | 0.05897 | 8 | microglia5 | Ion transport |
| GO:0006883 | cellular sodium ion homeostasis | 3/301 | 22/18800 | 0.00499 | 0.05918 | 3 | microglia5 | Ion transport |
| GO:0010881 | regulation of cardiac muscle contraction by regulation of the release of sequestered calcium ion | 3/301 | 22/18800 | 0.00499 | 0.05918 | 3 | microglia5 | Muscle contraction |
| GO:0019722 | calcium-mediated signaling | 9/301 | 202/18800 | 0.00538 | 0.06226 | 9 | microglia5 | Calcium transport |
| GO:0098742 | cell-cell adhesion via plasma-membrane adhesion molecules | 11/301 | 279/18800 | 0.00551 | 0.0635 | 11 | microglia5 | Cell adhesion |
| GO:0022010 | central nervous system myelination | 3/301 | 24/18800 | 0.00641 | 0.06917 | 3 | microglia5 | Enhanced myelination |
| GO:0032291 | axon ensheathment in central nervous system | 3/301 | 24/18800 | 0.00641 | 0.06917 | 3 | microglia5 | Enhanced myelination |
| GO:0098876 | vesicle-mediated transport to the plasma membrane | 7/301 | 139/18800 | 0.00713 | 0.07421 | 7 | microglia5 | Vesicle |
| GO:0032414 | positive regulation of ion transmembrane transporter activity | 6/301 | 106/18800 | 0.00719 | 0.07421 | 6 | microglia5 | Ion transport |
| GO:0032469 | endoplasmic reticulum calcium ion homeostasis | 3/301 | 25/18800 | 0.0072 | 0.07421 | 3 | microglia5 | Calcium transport |
| GO:1904861 | excitatory synapse assembly | 3/301 | 25/18800 | 0.0072 | 0.07421 | 3 | microglia5 | Synapse assembly |
| GO:0031641 | regulation of myelination | 4/301 | 48/18800 | 0.00721 | 0.07421 | 4 | microglia5 | Enhanced myelination |
| GO:0050772 | positive regulation of axonogenesis | 5/301 | 77/18800 | 0.00787 | 0.07744 | 5 | microglia5 | Axonogenesis |
| GO:0060314 | regulation of ryanodine-sensitive calcium-release channel activity | 3/301 | 26/18800 | 0.00804 | 0.07851 | 3 | microglia5 | Calcium transport |
| GO:0002269 | leukocyte activation involved in inflammatory response | 4/301 | 50/18800 | 0.00832 | 0.08029 | 4 | microglia5 | Leukocytes |
| GO:1990573 | potassium ion import across plasma membrane | 4/301 | 50/18800 | 0.00832 | 0.08029 | 4 | microglia5 | Ion transport |
| GO:0010810 | regulation of cell-substrate adhesion | 9/301 | 217/18800 | 0.00846 | 0.08114 | 9 | microglia5 | Cell adhesion |
| GO:0051592 | response to calcium ion | 7/301 | 144/18800 | 0.00859 | 0.08215 | 7 | microglia5 | Calcium transport |
| GO:0055074 | calcium ion homeostasis | 15/301 | 468/18800 | 0.00874 | 0.08272 | 15 | microglia5 | Calcium transport |
| GO:0043268 | positive regulation of potassium ion transport | 4/301 | 51/18800 | 0.00892 | 0.08344 | 4 | microglia5 | Ion transport |
| GO:1901018 | positive regulation of potassium ion transmembrane transporter activity | 3/301 | 27/18800 | 0.00894 | 0.08344 | 3 | microglia5 | Ion transport |

| | | | | | | | | |
|---|---|---|---|---|---|---|---|---|
| GO:0051279 | regulation of release of sequestered calcium ion into cytosol | 5/301 | 80/18800 | 0.00921 | 0.08567 | 5 | microglia5 | Calcium transport |
| GO:0051968 | positive regulation of synaptic transmission, glutamatergic | 3/301 | 29/18800 | 0.01091 | 0.09623 | 3 | microglia5 | Glutamatergic synapse |
| GO:0031345 | negative regulation of cell projection organization | 8/301 | 188/18800 | 0.01101 | 0.09657 | 8 | microglia5 | Cell projection organization |
| GO:0071277 | cellular response to calcium ion | 5/301 | 84/18800 | 0.01123 | 0.09803 | 5 | microglia5 | Calcium transport |
| GO:0098984 | neuron to neuron synapse | 26/307 | 347/19594 | 0 | 0 | 26 | microglia5 | Synapse assembly |
| GO:0032279 | asymmetric synapse | 25/307 | 323/19594 | 0 | 0 | 25 | microglia5 | Synapse assembly |
| GO:0098978 | glutamatergic synapse | 21/307 | 319/19594 | 0 | 0 | 21 | microglia5 | Synapse assembly |
| GO:0043209 | myelin sheath | 8/307 | 45/19594 | 0 | 1E-05 | 8 | microglia5 | Enhanced myelination |
| GO:0098793 | presynapse | 21/307 | 492/19594 | 4E-05 | 0.00043 | 21 | microglia5 | Synapse assembly |
| GO:0030665 | clathrin-coated vesicle membrane | 6/307 | 111/19594 | 0.00808 | 0.04104 | 6 | microglia5 | Vesicle |
| GO:0030136 | clathrin-coated vesicle | 8/307 | 192/19594 | 11 | 0.05218 | 8 | microglia5 | Vesicle |
| GO:0098562 | cytoplasmic side of membrane | 8/307 | 193/19594 | 0.01132 | 0.05218 | 8 | microglia5 | Plasma membrane |
| GO:0043218 | compact myelin | 2/307 | 12/19594 | 0.01456 | 0.05957 | 2 | microglia5 | Enhanced myelination |
| GO:0009898 | cytoplasmic side of plasma membrane | 7/307 | 169/19594 | 0.01738 | 0.06834 | 7 | microglia5 | Plasma membrane |
| GO:0050998 | nitric-oxide synthase binding | 3/299 | 13/18410 | 0.00107 | 0.03595 | 3 | microglia5 | Oxidative stress |
| GO:0051020 | GTPase binding | 12/299 | 298/18410 | 0.00361 | 0.08292 | 12 | microglia5 | GTPase activity |
| GO:0015662 | P-type ion transporter activity | 3/299 | 20/18410 | 0.00394 | 0.08292 | 3 | microglia5 | Ion transport |
| GO:0031267 | small GTPase binding | 11/299 | 267/18410 | 0.00442 | 0.08378 | 11 | microglia5 | GTPase activity |
| GO:0005096 | GTPase activator activity | 11/299 | 274/18410 | 0.00535 | 0.09288 | 11 | microglia5 | GTPase activity |
| GO:0030695 | GTPase regulator activity | 16/299 | 488/18410 | 0.00636 | 0.09456 | 16 | microglia5 | GTPase activity |
| hsa04514 | Cell adhesion molecules | 13/135 | 157/8170 | 0 | 0.00038 | 13 | microglia5 | Cell adhesion |
| hsa04961 | Endocrine and other factor-regulated calcium reabsorption | 6/135 | 53/8170 | 0.00022 | 0.00972 | 6 | microglia5 | Calcium transport |
| hsa04727 | GABAergic synapse | 6/135 | 89/8170 | 0.00345 | 0.05959 | 6 | microglia5 | Synapse assembly |
| GO:0031345 | negative regulation of cell projection organization | 5/30 | 188/18800 | 1E-05 | 0.01019 | 5 | oligos0 | Cell projection organization |
| GO:0042063 | gliogenesis | 4/30 | 291/18800 | 0.00112 | 0.05773 | 4 | oligos0 | Gliogenesis |
| GO:0051056 | regulation of small GTPase mediated signal transduction | 4/30 | 299/18800 | 0.00124 | 0.05773 | 4 | oligos0 | GTPase activity |
| GO:0007162 | negative regulation of cell adhesion | 4/30 | 305/18800 | 0.00133 | 0.05773 | 4 | oligos0 | Cell adhesion |
| GO:0048813 | dendrite morphogenesis | 3/30 | 139/18800 | 0.00139 | 0.05773 | 3 | oligos0 | Morphogenesis |
| GO:0099173 | postsynapse organization | 3/30 | 163/18800 | 0.00219 | 0.06399 | 3 | oligos0 | Synapse assembly |

| | | | | | | | | |
|---|---|---|---|---|---|---|---|---|
| GO:0042733 | embryonic digit morphogenesis | 2/30 | 58/18800 | 0.00385 | 0.07114 | 2 | oligos0 | Morphogenesis |
| GO:0060997 | dendritic spine morphogenesis | 2/30 | 58/18800 | 0.00385 | 0.07114 | 2 | oligos0 | Morphogenesis |
| GO:0050808 | synapse organization | 4/30 | 419/18800 | 0.00421 | 0.0748 | 4 | oligos0 | Synapse assembly |
| GO:0010812 | negative regulation of cell-substrate adhesion | 2/30 | 66/18800 | 0.00496 | 0.08034 | 2 | oligos0 | Cell adhesion |
| GO:0051057 | positive regulation of small GTPase mediated signal transduction | 2/30 | 70/18800 | 0.00556 | 0.08559 | 2 | oligos0 | GTPase activity |
| GO:0006814 | sodium ion transport | 3/30 | 249/18800 | 0.00716 | 0.09371 | 3 | oligos0 | Ion transport |
| GO:0051056 | regulation of small GTPase mediated signal transduction | 12/161 | 299/18800 | 1E-05 | 0.02276 | 12 | oligos1 | GTPase activity |
| GO:0051058 | negative regulation of small GTPase mediated signal transduction | 5/161 | 57/18800 | 0.00013 | 0.04199 | 5 | oligos1 | GTPase activity |
| GO:0097479 | synaptic vesicle localization | 4/161 | 52/18800 | 0.00102 | 0.08093 | 4 | oligos1 | Vesicle |
| GO:0043087 | regulation of GTPase activity | 10/161 | 364/18800 | 0.0012 | 0.08712 | 10 | oligos1 | GTPase activity |
| GO:0022604 | regulation of cell morphogenesis | 9/161 | 305/18800 | 0.00129 | 0.08892 | 9 | oligos1 | Morphogenesis |
| hsa04510 | Focal adhesion | 8/74 | 201/8166 | 0.00044 | 0.08558 | 8 | oligos1 | Cell adhesion |
| GO:0007409 | axonogenesis | 10/54 | 430/18800 | 0 | 2E-04 | 10 | oligos2 | Axonogenesis |
| GO:0015872 | dopamine transport | 4/54 | 51/18800 | 1E-05 | 0.00265 | 4 | oligos2 | Dopamine metabolism |
| GO:0099003 | vesicle-mediated transport in synapse | 6/54 | 197/18800 | 2E-05 | 0.00301 | 6 | oligos2 | Vesicle |
| GO:0010765 | positive regulation of sodium ion transport | 3/54 | 33/18800 | 0.00011 | 0.00738 | 3 | oligos2 | Ion transport |
| GO:0099504 | synaptic vesicle cycle | 5/54 | 183/18800 | 0.00018 | 0.0107 | 5 | oligos2 | Vesicle |
| GO:0050808 | synapse organization | 7/54 | 419/18800 | 0.00019 | 0.0107 | 7 | oligos2 | Synapse assembly |
| GO:0031346 | positive regulation of cell projection organization | 6/54 | 341/18800 | 0.00042 | 0.01955 | 6 | oligos2 | Cell projection organization |
| GO:0042416 | dopamine biosynthetic process | 2/54 | 12/18800 | 0.00052 | 0.02166 | 2 | oligos2 | Dopamine metabolism |
| GO:0048278 | vesicle docking | 3/54 | 57/18800 | 0.00059 | 0.02335 | 3 | oligos2 | Vesicle |
| GO:0006813 | potassium ion transport | 5/54 | 243/18800 | 0.00065 | 0.02361 | 5 | oligos2 | Ion transport |
| GO:0098876 | vesicle-mediated transport to the plasma membrane | 4/54 | 139/18800 | 0.00068 | 0.02366 | 4 | oligos2 | Vesicle |
| GO:0050770 | regulation of axonogenesis | 4/54 | 154/18800 | 1 | 0.02808 | 4 | oligos2 | Axonogenesis |
| GO:0043270 | positive regulation of ion transport | 5/54 | 273/18800 | 0.0011 | 0.02893 | 5 | oligos2 | Ion transport |
| GO:0050772 | positive regulation of axonogenesis | 3/54 | 77/18800 | 0.00141 | 0.0326 | 3 | oligos2 | Axonogenesis |
| GO:0031629 | synaptic vesicle fusion to presynaptic active zone membrane | 2/54 | 21/18800 | 0.00164 | 0.03451 | 2 | oligos2 | Vesicle |
| GO:0099500 | vesicle fusion to plasma membrane | 2/54 | 22/18800 | 0.0018 | 0.03489 | 2 | oligos2 | Vesicle |

| | | | | | | | | |
|---|---|---|---|---|---|---|---|---|
| GO:0002028 | regulation of sodium ion transport | 3/54 | 90/18800 | 0.00221 | 0.03984 | 3 | oligos2 | Ion transport |
| GO:0034109 | homotypic cell-cell adhesion | 3/54 | 93/18800 | 0.00242 | 0.04307 | 3 | oligos2 | Cell adhesion |
| GO:0048499 | synaptic vesicle membrane organization | 2/54 | 26/18800 | 0.00252 | 0.0441 | 2 | oligos2 | Vesicle |
| GO:0001963 | synaptic transmission, dopaminergic | 2/54 | 27/18800 | 0.00271 | 0.04548 | 2 | oligos2 | Dopamine metabolism |
| GO:1901017 | negative regulation of potassium ion transmembrane transporter activity | 2/54 | 28/18800 | 0.00292 | 0.04819 | 2 | oligos2 | Ion transport |
| GO:0071805 | potassium ion transmembrane transport | 4/54 | 219/18800 | 0.00359 | 0.0569 | 4 | oligos2 | Ion transport |
| GO:1901380 | negative regulation of potassium ion transmembrane transport | 2/54 | 32/18800 | 0.0038 | 0.05858 | 2 | oligos2 | Ion transport |
| GO:0034110 | regulation of homotypic cell-cell adhesion | 2/54 | 35/18800 | 0.00453 | 0.06569 | 2 | oligos2 | Cell adhesion |
| GO:0014046 | dopamine secretion | 2/54 | 37/18800 | 0.00506 | 0.06877 | 2 | oligos2 | Dopamine metabolism |
| GO:0014059 | regulation of dopamine secretion | 2/54 | 37/18800 | 0.00506 | 0.06877 | 2 | oligos2 | Dopamine metabolism |
| GO:0043267 | negative regulation of potassium ion transport | 2/54 | 39/18800 | 0.00561 | 0.07105 | 2 | oligos2 | Ion transport |
| GO:0006814 | sodium ion transport | 4/54 | 249/18800 | 0.00566 | 0.07105 | 4 | oligos2 | Ion transport |
| GO:0007212 | dopamine receptor signaling pathway | 2/54 | 40/18800 | 0.00589 | 0.07105 | 2 | oligos2 | Dopamine metabolism |
| GO:0010959 | regulation of metal ion transport | 5/54 | 403/18800 | 0.0059 | 0.07105 | 5 | oligos2 | Ion transport |
| GO:0003007 | heart morphogenesis | 4/54 | 254/18800 | 0.00606 | 0.07152 | 4 | oligos2 | Morphogenesis |
| GO:0042417 | dopamine metabolic process | 2/54 | 42/18800 | 0.00648 | 0.07488 | 2 | oligos2 | Dopamine metabolism |
| GO:0099643 | signal release from synapse | 3/54 | 145/18800 | 0.00836 | 0.0879 | 3 | oligos2 | Synapse assembly |
| GO:0098793 | presynapse | 11/54 | 492/19594 | 0 | 0 | 11 | oligos2 | Synapse assembly |
| GO:0098562 | cytoplasmic side of membrane | 5/54 | 193/19594 | 0.00019 | 0.00463 | 5 | oligos2 | Plasma membrane |
| GO:0008021 | synaptic vesicle | 5/54 | 196/19594 | 2E-04 | 0.00463 | 5 | oligos2 | Vesicle |
| GO:0070382 | exocytic vesicle | 5/54 | 214/19594 | 3E-04 | 0.00625 | 5 | oligos2 | Vesicle |
| GO:0098984 | neuron to neuron synapse | 6/54 | 347/19594 | 0.00037 | 0.00693 | 6 | oligos2 | Synapse assembly |
| GO:0098691 | dopaminergic synapse | 2/54 | 11/19594 | 4E-04 | 0.00693 | 2 | oligos2 | Dopamine metabolism |
| GO:0060198 | clathrin-sculpted vesicle | 2/54 | 12/19594 | 0.00048 | 0.00766 | 2 | oligos2 | Vesicle |
| GO:0030133 | transport vesicle | 6/54 | 402/19594 | 0.00081 | 0.00958 | 6 | oligos2 | Vesicle |
| GO:0009898 | cytoplasmic side of plasma membrane | 4/54 | 169/19594 | 0.00121 | 0.01309 | 4 | oligos2 | Plasma membrane |
| GO:0032279 | asymmetric synapse | 5/54 | 323/19594 | 0.00193 | 0.01804 | 5 | oligos2 | Synapse assembly |
| GO:0030658 | transport vesicle membrane | 4/54 | 205/19594 | 0.00245 | 0.01912 | 4 | oligos2 | Vesicle |

| | | | | | | | | |
|---|---|---|---|---|---|---|---|---|
| GO:0031234 | extrinsic component of cytoplasmic side of plasma membrane | 3/54 | 99/19594 | 0.00257 | 0.01912 | 3 | oligos2 | Plasma membrane |
| GO:0030285 | integral component of synaptic vesicle membrane | 2/54 | 28/19594 | 0.00269 | 0.01912 | 2 | oligos2 | Vesicle |
| GO:0030672 | synaptic vesicle membrane | 3/54 | 103/19594 | 0.00288 | 0.01914 | 3 | oligos2 | Vesicle |
| GO:0099501 | exocytic vesicle membrane | 3/54 | 103/19594 | 0.00288 | 0.01914 | 3 | oligos2 | Vesicle |
| GO:0098563 | intrinsic component of synaptic vesicle membrane | 2/54 | 40/19594 | 0.00544 | 32 | 2 | oligos2 | Vesicle |
| GO:0098978 | glutamatergic synapse | 4/54 | 319/19594 | 0.01151 | 0.0593 | 4 | oligos2 | Synapse assembly |
| GO:0098685 | Schaffer collateral - CA1 synapse | 2/54 | 72/19594 | 0.01685 | 0.07922 | 2 | oligos2 | Synapse assembly |
| GO:0099186 | structural constituent of postsynapse | 2/53 | 13/18410 | 0.00062 | 0.01998 | 2 | oligos2 | Synapse assembly |
| GO:0098918 | structural constituent of synapse | 2/53 | 19/18410 | 0.00135 | 0.02527 | 2 | oligos2 | Synapse assembly |
| GO:0048306 | calcium-dependent protein binding | 3/53 | 87/18410 | 0.00201 | 0.0302 | 3 | oligos2 | Calcium transport |
| GO:0022853 | active ion transmembrane transporter activity | 4/53 | 254/18410 | 0.0061 | 0.06071 | 4 | oligos2 | Ion transport |
| GO:0046873 | metal ion transmembrane transporter activity | 5/53 | 428/18410 | 0.00762 | 0.06877 | 5 | oligos2 | Ion transport |
| GO:0015079 | potassium ion transmembrane transporter activity | 3/53 | 154/18410 | 0.00991 | 0.07686 | 3 | oligos2 | Ion transport |
| GO:0098632 | cell-cell adhesion mediator activity | 2/53 | 54/18410 | 0.01058 | 0.0788 | 2 | oligos2 | Cell adhesion |
| GO:0098631 | cell adhesion mediator activity | 2/53 | 64/18410 | 0.01463 | 0.09404 | 2 | oligos2 | Cell adhesion |
| GO:0003924 | GTPase activity | 4/53 | 336/18410 | 0.01582 | 0.09889 | 4 | oligos2 | GTPase activity |
| hsa04721 | Synaptic vesicle cycle | 6/35 | 78/8166 | 0 | 6E-05 | 6 | oligos2 | Vesicle |
| hsa04728 | Dopaminergic synapse | 7/35 | 132/8166 | 0 | 6E-05 | 7 | oligos2 | Dopamine metabolism |
| GO:0007409 | axonogenesis | 16/87 | 430/18800 | 0 | 0 | 16 | oligos3 | Axonogenesis |
| GO:0048813 | dendrite morphogenesis | 8/87 | 139/18800 | 0 | 9E-05 | 8 | oligos3 | Morphogenesis |
| GO:0031346 | positive regulation of cell projection organization | 11/87 | 341/18800 | 0 | 0.00012 | 11 | oligos3 | Cell projection organization |
| GO:0031345 | negative regulation of cell projection organization | 8/87 | 188/18800 | 0 | 0.00043 | 8 | oligos3 | Cell projection organization |
| GO:0050808 | synapse organization | 11/87 | 419/18800 | 0 | 0.00056 | 11 | oligos3 | Synapse assembly |
| GO:0050770 | regulation of axonogenesis | 7/87 | 154/18800 | 1E-05 | 0.00095 | 7 | oligos3 | Axonogenesis |
| GO:0007158 | neuron cell-cell adhesion | 3/87 | 17/18800 | 6E-05 | 0.00472 | 3 | oligos3 | Cell adhesion |
| GO:0060997 | dendritic spine morphogenesis | 4/87 | 58/18800 | 0.00015 | 0.00974 | 4 | oligos3 | Morphogenesis |
| GO:0048814 | regulation of dendrite morphogenesis | 4/87 | 64/18800 | 0.00022 | 0.01375 | 4 | oligos3 | Morphogenesis |
| GO:0031589 | cell-substrate adhesion | 8/87 | 364/18800 | 0.00028 | 0.01504 | 8 | oligos3 | Cell adhesion |

| | | | | | | | | |
|---|---|---|---|---|---|---|---|---|
| GO:0034765 | regulation of ion transmembrane transport | 9/87 | 476/18800 | 0.00035 | 0.01659 | 9 | oligos3 | Ion transport |
| GO:0050807 | regulation of synapse organization | 6/87 | 209/18800 | 0.00042 | 0.01817 | 6 | oligos3 | Synapse assembly |
| GO:0050772 | positive regulation of axonogenesis | 4/87 | 77/18800 | 0.00045 | 0.01857 | 4 | oligos3 | Axonogenesis |
| GO:0051056 | regulation of small GTPase mediated signal transduction | 7/87 | 299/18800 | 0.00047 | 0.01857 | 7 | oligos3 | GTPase activity |
| GO:0050803 | regulation of synapse structure or activity | 6/87 | 215/18800 | 0.00049 | 0.01871 | 6 | oligos3 | Synapse assembly |
| GO:0099175 | regulation of postsynapse organization | 4/87 | 86/18800 | 0.00068 | 0.02344 | 4 | oligos3 | Synapse assembly |
| GO:0060560 | developmental growth involved in morphogenesis | 6/87 | 234/18800 | 0.00076 | 0.02465 | 6 | oligos3 | Morphogenesis |
| GO:0001738 | morphogenesis of a polarized epithelium | 4/87 | 94/18800 | 0.00095 | 0.02615 | 4 | oligos3 | Morphogenesis |
| GO:0099173 | postsynapse organization | 5/87 | 163/18800 | 0.00096 | 0.02615 | 5 | oligos3 | Synapse assembly |
| GO:0061001 | regulation of dendritic spine morphogenesis | 3/87 | 44/18800 | 0.00111 | 0.02696 | 3 | oligos3 | Morphogenesis |
| GO:2000651 | positive regulation of sodium ion transmembrane transporter activity | 2/87 | 16/18800 | 0.00244 | 0.04149 | 2 | oligos3 | Ion transport |
| GO:0003184 | pulmonary valve morphogenesis | 2/87 | 17/18800 | 0.00275 | 0.04467 | 2 | oligos3 | Morphogenesis |
| GO:2000027 | regulation of animal organ morphogenesis | 4/87 | 129/18800 | 0.00304 | 0.04744 | 4 | oligos3 | Morphogenesis |
| GO:0010810 | regulation of cell-substrate adhesion | 5/87 | 217/18800 | 0.00336 | 0.05201 | 5 | oligos3 | Cell adhesion |
| GO:0071805 | potassium ion transmembrane transport | 5/87 | 219/18800 | 0.00349 | 0.0536 | 5 | oligos3 | Ion transport |
| GO:1901016 | regulation of potassium ion transmembrane transporter activity | 3/87 | 67/18800 | 0.0037 | 0.05484 | 3 | oligos3 | Ion transport |
| GO:1902307 | positive regulation of sodium ion transmembrane transport | 2/87 | 20/18800 | 0.00381 | 0.05544 | 2 | oligos3 | Ion transport |
| GO:0007204 | positive regulation of cytosolic calcium ion concentration | 6/87 | 325/18800 | 0.00398 | 0.05625 | 6 | oligos3 | Calcium transport |
| GO:0051057 | positive regulation of small GTPase mediated signal transduction | 3/87 | 70/18800 | 0.00419 | 0.05625 | 3 | oligos3 | GTPase activity |
| GO:0051592 | response to calcium ion | 4/87 | 144/18800 | 0.00449 | 0.0589 | 4 | oligos3 | Calcium transport |
| GO:0010881 | regulation of cardiac muscle contraction by regulation of the release of sequestered calcium ion | 2/87 | 22/18800 | 0.0046 | 0.0589 | 2 | oligos3 | Muscle contraction |
| GO:0007160 | cell-matrix adhesion | 5/87 | 235/18800 | 0.00471 | 0.05974 | 5 | oligos3 | Cell adhesion |
| GO:0002689 | negative regulation of leukocyte chemotaxis | 2/87 | 23/18800 | 0.00503 | 0.06287 | 2 | oligos3 | Leukocytes |
| GO:0006874 | cellular calcium ion homeostasis | 7/87 | 456/18800 | 0.0052 | 0.06457 | 7 | oligos3 | Calcium transport |

| | | | | | | | | |
|---|---|---|---|---|---|---|---|---|
| GO:0006813 | potassium ion transport | 5/87 | 243/18800 | 0.00542 | 0.06457 | 5 | oligos3 | Ion transport |
| GO:0048754 | branching morphogenesis of an epithelial tube | 4/87 | 153/18800 | 0.00557 | 0.06478 | 4 | oligos3 | Morphogenesis |
| GO:0097553 | calcium ion transmembrane import into cytosol | 4/87 | 154/18800 | 0.00569 | 0.06538 | 4 | oligos3 | Calcium transport |
| GO:0055074 | calcium ion homeostasis | 7/87 | 468/18800 | 0.00598 | 0.06639 | 7 | oligos3 | Calcium transport |
| GO:0051279 | regulation of release of sequestered calcium ion into cytosol | 3/87 | 80/18800 | 0.00608 | 0.06669 | 3 | oligos3 | Calcium transport |
| GO:0051480 | regulation of cytosolic calcium ion concentration | 6/87 | 356/18800 | 0.00617 | 0.06719 | 6 | oligos3 | Calcium transport |
| GO:0010882 | regulation of cardiac muscle contraction by calcium ion signaling | 2/87 | 26/18800 | 0.0064 | 0.06753 | 2 | oligos3 | Muscle contraction |
| GO:0021952 | central nervous system projection neuron axonogenesis | 2/87 | 26/18800 | 0.0064 | 0.06753 | 2 | oligos3 | Axonogenesis |
| GO:0071277 | cellular response to calcium ion | 3/87 | 84/18800 | 0.00696 | 0.07256 | 3 | oligos3 | Calcium transport |
| GO:1901017 | negative regulation of potassium ion transmembrane transporter activity | 2/87 | 28/18800 | 0.0074 | 0.07571 | 2 | oligos3 | Ion transport |
| GO:0032412 | regulation of ion transmembrane transporter activity | 5/87 | 263/18800 | 0.00751 | 0.07637 | 5 | oligos3 | Ion transport |
| GO:0060402 | calcium ion transport into cytosol | 4/87 | 171/18800 | 0.00819 | 0.07993 | 4 | oligos3 | Calcium transport |
| GO:0010880 | regulation of release of sequestered calcium ion into cytosol by sarcoplasmic reticulum | 2/87 | 31/18800 | 0.00902 | 0.08509 | 2 | oligos3 | Calcium transport |
| GO:0098742 | cell-cell adhesion via plasma-membrane adhesion molecules | 5/87 | 279/18800 | 0.00955 | 0.08755 | 5 | oligos3 | Cell adhesion |
| GO:0003180 | aortic valve morphogenesis | 2/87 | 32/18800 | 0.0096 | 0.08755 | 2 | oligos3 | Morphogenesis |
| GO:1901380 | negative regulation of potassium ion transmembrane transport | 2/87 | 32/18800 | 0.0096 | 0.08755 | 2 | oligos3 | Ion transport |
| GO:1901379 | regulation of potassium ion transmembrane transport | 3/87 | 95/18800 | 0.00975 | 0.08759 | 3 | oligos3 | Ion transport |
| GO:0010769 | regulation of cell morphogenesis involved in differentiation | 3/87 | 96/18800 | 0.01004 | 0.08917 | 3 | oligos3 | Morphogenesis |
| GO:0010765 | positive regulation of sodium ion transport | 2/87 | 33/18800 | 0.01018 | 0.08953 | 2 | oligos3 | Ion transport |
| GO:0061138 | morphogenesis of a branching epithelium | 4/87 | 185/18800 | 0.01072 | 0.09197 | 4 | oligos3 | Morphogenesis |
| GO:0032228 | regulation of synaptic transmission, GABAergic | 2/87 | 34/18800 | 0.01079 | 0.09197 | 2 | oligos3 | GABAergic synapse |
| GO:0014808 | release of sequestered calcium ion into cytosol by sarcoplasmic reticulum | 2/87 | 35/18800 | 0.01141 | 0.09394 | 2 | oligos3 | Calcium transport |

| | | | | | | | | |
|---|---|---|---|---|---|---|---|---|
| GO:0021955 | central nervous system neuron axonogenesis | 2/87 | 35/18800 | 0.01141 | 0.09394 | 2 | oligos3 | Axonogenesis |
| GO:0060401 | cytosolic calcium ion transport | 4/87 | 190/18800 | 0.01173 | 0.09566 | 4 | oligos3 | Calcium transport |
| GO:1903514 | release of sequestered calcium ion into cytosol by endoplasmic reticulum | 2/87 | 36/18800 | 0.01204 | 0.09666 | 2 | oligos3 | Calcium transport |
| GO:0010522 | regulation of calcium ion transport into cytosol | 3/87 | 103/18800 | 0.01214 | 0.09666 | 3 | oligos3 | Calcium transport |
| GO:0071542 | dopaminergic neuron differentiation | 2/87 | 37/18800 | 0.01269 | 0.09784 | 2 | oligos3 | Dopamine metabolism |
| GO:0032414 | positive regulation of ion transmembrane transporter activity | 3/87 | 106/18800 | 0.01311 | 0.09925 | 3 | oligos3 | Ion transport |
| GO:0032279 | asymmetric synapse | 12/90 | 323/19594 | 0 | 0 | 12 | oligos3 | Synapse assembly |
| GO:0098984 | neuron to neuron synapse | 12/90 | 347/19594 | 0 | 0 | 12 | oligos3 | Synapse assembly |
| GO:0098793 | presynapse | 9/90 | 492/19594 | 0.00043 | 0.00434 | 9 | oligos3 | Synapse assembly |
| GO:0098978 | glutamatergic synapse | 6/90 | 319/19594 | 0.00352 | 0.02001 | 6 | oligos3 | Synapse assembly |
| GO:0009898 | cytoplasmic side of plasma membrane | 4/90 | 169/19594 | 0.00768 | 0.03638 | 4 | oligos3 | Plasma membrane |
| GO:0031234 | extrinsic component of cytoplasmic side of plasma membrane | 3/90 | 99/19594 | 0.0107 | 0.04531 | 3 | oligos3 | Plasma membrane |
| GO:0098562 | cytoplasmic side of membrane | 4/90 | 193/19594 | 0.01208 | 0.05008 | 4 | oligos3 | Plasma membrane |
| GO:0008021 | synaptic vesicle | 4/90 | 196/19594 | 0.01272 | 0.05166 | 4 | oligos3 | Vesicle |
| GO:0070382 | exocytic vesicle | 4/90 | 214/19594 | 0.01704 | 0.06279 | 4 | oligos3 | Vesicle |
| GO:0005244 | voltage-gated ion channel activity | 5/85 | 201/18410 | 0.00239 | 0.06419 | 5 | oligos3 | Ion transport |
| GO:0009898 | cytoplasmic side of plasma membrane | 3/24 | 169/19594 | 0.00112 | 0.06867 | 3 | oligos4 | Plasma membrane |
| GO:0098562 | cytoplasmic side of membrane | 3/24 | 193/19594 | 0.00164 | 0.06867 | 3 | oligos4 | Plasma membrane |
| GO:2001242 | regulation of intrinsic apoptotic signaling pathway | 13/215 | 171/18800 | 0 | 2E-05 | 13 | oligos5 | Apoptosis |
| GO:2001244 | positive regulation of intrinsic apoptotic signaling pathway | 8/215 | 61/18800 | 0 | 9E-05 | 8 | oligos5 | Apoptosis |
| GO:0098869 | cellular oxidant detoxification | 9/215 | 100/18800 | 0 | 0.00031 | 9 | oligos5 | Oxidative stress |
| GO:0042552 | myelination | 10/215 | 138/18800 | 0 | 0.00056 | 10 | oligos5 | Enhanced myelination |
| GO:0007272 | ensheathment of neurons | 10/215 | 140/18800 | 0 | 0.00056 | 10 | oligos5 | Enhanced myelination |
| GO:0008366 | axon ensheathment | 10/215 | 140/18800 | 0 | 0.00056 | 10 | oligos5 | Enhanced myelination |
| GO:0097193 | intrinsic apoptotic signaling pathway | 14/215 | 295/18800 | 1E-05 | 0.00082 | 14 | oligos5 | Apoptosis |
| GO:0046034 | ATP metabolic process | 13/215 | 273/18800 | 2E-05 | 0.00125 | 13 | oligos5 | Energy production |

| | | | | | | | | |
|---|---|---|---|---|---|---|---|---|
| GO:2001235 | positive regulation of apoptotic signaling pathway | 9/215 | 134/18800 | 2E-05 | 0.00177 | 9 | oligos5 | Apoptosis |
| GO:2001233 | regulation of apoptotic signaling pathway | 15/215 | 370/18800 | 2E-05 | 0.00177 | 15 | oligos5 | Apoptosis |
| GO:0042063 | gliogenesis | 13/215 | 291/18800 | 3E-05 | 0.00224 | 13 | oligos5 | Gliogenesis |
| GO:0006979 | response to oxidative stress | 16/215 | 433/18800 | 4E-05 | 0.0027 | 16 | oligos5 | Oxidative stress |
| GO:0007409 | axonogenesis | 15/215 | 430/18800 | 0.00013 | 0.00791 | 15 | oligos5 | Axonogenesis |
| GO:0010506 | regulation of autophagy | 13/215 | 336/18800 | 0.00014 | 0.00794 | 13 | oligos5 | Autophagy |
| GO:0022010 | central nervous system myelination | 4/215 | 24/18800 | 0.00015 | 0.00814 | 4 | oligos5 | Enhanced myelination |
| GO:0032291 | axon ensheathment in central nervous system | 4/215 | 24/18800 | 0.00015 | 0.00814 | 4 | oligos5 | Enhanced myelination |
| GO:0006091 | generation of precursor metabolites and energy | 16/215 | 494/18800 | 0.00019 | 0.00992 | 16 | oligos5 | Energy production |
| GO:0006119 | oxidative phosphorylation | 8/215 | 139/18800 | 2E-04 | 0.01002 | 8 | oligos5 | Oxidative stress |
| GO:0055076 | transition metal ion homeostasis | 8/215 | 139/18800 | 2E-04 | 0.01002 | 8 | oligos5 | Ion transport |
| GO:0046916 | cellular transition metal ion homeostasis | 7/215 | 115/18800 | 0.00036 | 0.01579 | 7 | oligos5 | Ion transport |
| GO:1902235 | regulation of endoplasmic reticulum stress-induced intrinsic apoptotic signaling pathway | 4/215 | 32/18800 | 0.00047 | 0.01924 | 4 | oligos5 | Apoptosis |
| GO:0035966 | response to topologically incorrect protein | 8/215 | 160/18800 | 0.00052 | 0.02111 | 8 | oligos5 | Unfolded Protein Response |
| GO:0006826 | iron ion transport | 5/215 | 59/18800 | 0.00057 | 0.02236 | 5 | oligos5 | Ion transport |
| GO:0006457 | protein folding | 9/215 | 212/18800 | 0.00077 | 0.02787 | 9 | oligos5 | Unfolded Protein Response |
| GO:0034620 | cellular response to unfolded protein | 6/215 | 98/18800 | 0.00092 | 0.0307 | 6 | oligos5 | Unfolded Protein Response |
| GO:0000041 | transition metal ion transport | 6/215 | 101/18800 | 0.00107 | 0.03501 | 6 | oligos5 | Ion transport |
| GO:0006986 | response to unfolded protein | 7/215 | 139/18800 | 0.00111 | 0.03531 | 7 | oligos5 | Unfolded Protein Response |
| GO:0015980 | energy derivation by oxidation of organic compounds | 11/215 | 321/18800 | 0.00122 | 0.03743 | 11 | oligos5 | Oxidative stress |
| GO:0061077 | chaperone-mediated protein folding | 5/215 | 70/18800 | 0.00124 | 0.03761 | 5 | oligos5 | Unfolded Protein Response |
| GO:1904427 | positive regulation of calcium ion transmembrane transport | 5/215 | 71/18800 | 0.00132 | 0.03918 | 5 | oligos5 | Calcium transport |
| GO:1902236 | negative regulation of endoplasmic reticulum stress-induced intrinsic apoptotic signaling pathway | 3/215 | 20/18800 | 0.00146 | 0.04101 | 3 | oligos5 | Apoptosis |
| GO:0032288 | myelin assembly | 3/215 | 22/18800 | 0.00193 | 0.04786 | 3 | oligos5 | Enhanced myelination |
| GO:0072332 | intrinsic apoptotic signaling pathway by p53 class mediator | 5/215 | 79/18800 | 0.00212 | 0.05054 | 5 | oligos5 | Apoptosis |
| GO:0019430 | removal of superoxide radicals | 3/215 | 23/18800 | 0.00221 | 0.05078 | 3 | oligos5 | Oxidative stress |

| | | | | | | | | |
|---|---|---|---|---|---|---|---|---|
| GO:0035967 | cellular response to topologically incorrect protein | 6/215 | 117/18800 | 0.00228 | 0.05151 | 6 | oligos5 | Unfolded Protein Response |
| GO:0043281 | regulation of cysteine-type endopeptidase activity involved in apoptotic process | 8/215 | 204/18800 | 0.00246 | 0.0517 | 8 | oligos5 | Apoptosis |
| GO:0051928 | positive regulation of calcium ion transport | 6/215 | 119/18800 | 0.00248 | 0.0517 | 6 | oligos5 | Calcium transport |
| GO:1901028 | regulation of mitochondrial outer membrane permeabilization involved in apoptotic signaling pathway | 3/215 | 24/18800 | 0.0025 | 0.0517 | 3 | oligos5 | Apoptosis |
| GO:0071451 | cellular response to superoxide | 3/215 | 25/18800 | 0.00282 | 0.05484 | 3 | oligos5 | Oxidative stress |
| GO:0055072 | iron ion homeostasis | 5/215 | 85/18800 | 0.00292 | 0.05528 | 5 | oligos5 | Ion transport |
| GO:0002726 | positive regulation of T cell cytokine production | 3/215 | 26/18800 | 0.00316 | 0.05726 | 3 | oligos5 | T cell activity |
| GO:0000303 | response to superoxide | 3/215 | 28/18800 | 0.00391 | 0.06316 | 3 | oligos5 | Oxidative stress |
| GO:0042775 | mitochondrial ATP synthesis coupled electron transport | 5/215 | 92/18800 | 0.0041 | 0.06391 | 5 | oligos5 | Mitochondrial changes |
| GO:2001234 | negative regulation of apoptotic signaling pathway | 8/215 | 230/18800 | 0.00508 | 0.07346 | 8 | oligos5 | Apoptosis |
| GO:0010039 | response to iron ion | 3/215 | 31/18800 | 0.00524 | 0.07452 | 3 | oligos5 | Iron transport |
| GO:0046902 | regulation of mitochondrial membrane permeability | 4/215 | 62/18800 | 0.00553 | 0.0748 | 4 | oligos5 | Mitochondrial changes |
| GO:0034599 | cellular response to oxidative stress | 9/215 | 284/18800 | 0.00554 | 0.0748 | 9 | oligos5 | Oxidative stress |
| GO:0070059 | intrinsic apoptotic signaling pathway in response to endoplasmic reticulum stress | 4/215 | 63/18800 | 0.00585 | 0.07706 | 4 | oligos5 | Apoptosis |
| GO:0008630 | intrinsic apoptotic signaling pathway in response to DNA damage | 5/215 | 101/18800 | 0.00609 | 0.0794 | 5 | oligos5 | Apoptosis |
| GO:2001243 | negative regulation of intrinsic apoptotic signaling pathway | 5/215 | 102/18800 | 0.00634 | 0.08192 | 5 | oligos5 | Apoptosis |
| GO:1901030 | positive regulation of mitochondrial outer membrane permeabilization involved in apoptotic signaling pathway | 2/215 | 11/18800 | 0.00669 | 0.08392 | 2 | oligos5 | Apoptosis |
| GO:0010959 | regulation of metal ion transport | 11/215 | 403/18800 | 0.00687 | 0.08529 | 11 | oligos5 | Ion transport |
| GO:0006879 | cellular iron ion homeostasis | 4/215 | 66/18800 | 0.0069 | 0.08529 | 4 | oligos5 | Iron transport |
| GO:0097345 | mitochondrial outer membrane permeabilization | 3/215 | 35/18800 | 0.00737 | 0.08953 | 3 | oligos5 | Mitochondrial changes |
| GO:0002720 | positive regulation of cytokine production involved in immune response | 4/215 | 68/18800 | 0.00765 | 0.0921 | 4 | oligos5 | Immune response |
| GO:0008637 | apoptotic mitochondrial changes | 5/215 | 107/18800 | 0.00774 | 0.09261 | 5 | oligos5 | Apoptosis |
| GO:0044650 | adhesion of symbiont to host cell | 2/215 | 12/18800 | 0.00797 | 0.09261 | 2 | oligos5 | Cell adhesion |
| GO:1901029 | negative regulation of mitochondrial outer membrane permeabilization | 2/215 | 12/18800 | 0.00797 | 0.09261 | 2 | oligos5 | Apoptosis |

| | | | | | | | | |
|---|---|---|---|---|---|---|---|---|
| | involved in apoptotic signaling pathway | | | | | | | |
| GO:0051924 | regulation of calcium ion transport | 8/215 | 251/18800 | 0.00844 | 0.09615 | 8 | oligos5 | Calcium transport |
| GO:0046640 | regulation of alpha-beta T cell proliferation | 3/215 | 37/18800 | 0.00861 | 0.09615 | 3 | oligos5 | T cell activity |
| GO:1901021 | positive regulation of calcium ion transmembrane transporter activity | 3/215 | 37/18800 | 0.00861 | 0.09615 | 3 | oligos5 | Calcium transport |
| GO:0016236 | macroautophagy | 9/215 | 306/18800 | 0.00884 | 0.09778 | 9 | oligos5 | Autophagy |
| GO:0051881 | regulation of mitochondrial membrane potential | 4/215 | 71/18800 | 0.00889 | 0.09778 | 4 | oligos5 | Mitochondrial changes |
| GO:0002369 | T cell cytokine production | 3/215 | 38/18800 | 0.00927 | 0.09778 | 3 | oligos5 | T cell activity |
| GO:0002724 | regulation of T cell cytokine production | 3/215 | 38/18800 | 0.00927 | 0.09778 | 3 | oligos5 | T cell activity |
| GO:0010927 | cellular component assembly involved in morphogenesis | 5/215 | 112/18800 | 0.00933 | 0.09778 | 5 | oligos5 | Morphogenesis |
| GO:0006801 | superoxide metabolic process | 4/215 | 72/18800 | 0.00934 | 0.09778 | 4 | oligos5 | Oxidative stress |
| GO:0005925 | focal adhesion | 35/222 | 419/19594 | 0 | 0 | 35 | oligos5 | Cell adhesion |
| GO:0043209 | myelin sheath | 10/222 | 45/19594 | 0 | 0 | 10 | oligos5 | Enhanced myelination |
| GO:0032279 | asymmetric synapse | 15/222 | 323/19594 | 0 | 0.00012 | 15 | oligos5 | Synapse assembly |
| GO:0043218 | compact myelin | 4/222 | 12/19594 | 1E-05 | 0.00016 | 4 | oligos5 | Enhanced myelination |
| GO:0098984 | neuron to neuron synapse | 15/222 | 347/19594 | 1E-05 | 0.00021 | 15 | oligos5 | Synapse assembly |
| GO:0005741 | mitochondrial outer membrane | 11/222 | 205/19594 | 2E-05 | 0.00043 | 11 | oligos5 | Mitochondrial changes |
| GO:0030666 | endocytic vesicle membrane | 10/222 | 194/19594 | 8E-05 | 0.00111 | 10 | oligos5 | Vesicle |
| GO:0060205 | cytoplasmic vesicle lumen | 12/222 | 325/19594 | 0.00035 | 0.00427 | 12 | oligos5 | Vesicle |
| GO:0031983 | vesicle lumen | 12/222 | 327/19594 | 0.00037 | 0.00437 | 12 | oligos5 | Vesicle |
| GO:0098800 | inner mitochondrial membrane protein complex | 8/222 | 155/19594 | 4E-04 | 0.00451 | 8 | oligos5 | Mitochondrial changes |
| GO:0005753 | mitochondrial proton-transporting ATP synthase complex | 3/222 | 21/19594 | 0.00164 | 0.01469 | 3 | oligos5 | Mitochondrial changes |
| GO:0030139 | endocytic vesicle | 11/222 | 342/19594 | 0.00187 | 0.01611 | 11 | oligos5 | Vesicle |
| GO:0005743 | mitochondrial inner membrane | 13/222 | 491/19594 | 0.00409 | 0.03142 | 13 | oligos5 | Mitochondrial changes |
| GO:0005746 | mitochondrial respirasome | 5/222 | 94/19594 | 0.00433 | 0.03192 | 5 | oligos5 | Mitochondrial changes |
| GO:0098798 | mitochondrial protein-containing complex | 9/222 | 281/19594 | 0.00489 | 0.03535 | 9 | oligos5 | Mitochondrial changes |
| GO:0000276 | mitochondrial proton-transporting ATP synthase complex, coupling factor F(o) | 2/222 | 11/19594 | 0.00657 | 0.04575 | 2 | oligos5 | Mitochondrial changes |
| GO:0030670 | phagocytic vesicle membrane | 4/222 | 77/19594 | 0.0114 | 0.06801 | 4 | oligos5 | Vesicle |
| GO:0005747 | mitochondrial respiratory chain complex I | 3/222 | 49/19594 | 0.01803 | 0.09418 | 3 | oligos5 | Mitochondrial changes |

| | | | | | | | | |
|---|---|---|---|---|---|---|---|---|
| GO:0016209 | antioxidant activity | 9/219 | 85/18410 | 0 | 6E-05 | 9 | oligos5 | Oxidative stress |
| GO:0008199 | ferric iron binding | 3/219 | 11/18410 | 0.00026 | 0.00945 | 3 | oligos5 | Iron transport |
| GO:0051082 | unfolded protein binding | 7/219 | 121/18410 | 0.00061 | 0.01924 | 7 | oligos5 | Unfolded Protein Response |
| GO:0015453 | oxidoreduction-driven active transmembrane transporter activity | 5/219 | 72/18410 | 0.00167 | 0.04002 | 5 | oligos5 | Oxidative stress |
| GO:0044183 | protein folding chaperone | 4/219 | 43/18410 | 0.00167 | 0.04002 | 4 | oligos5 | Unfolded Protein Response |
| GO:0004602 | glutathione peroxidase activity | 3/219 | 22/18410 | 0.00216 | 0.04403 | 3 | oligos5 | Oxidative stress |
| GO:0004601 | peroxidase activity | 4/219 | 52/18410 | 0.00337 | 0.05522 | 4 | oligos5 | Oxidative stress |
| GO:0008198 | ferrous iron binding | 3/219 | 26/18410 | 0.00353 | 0.05522 | 3 | oligos5 | Iron transport |
| GO:0016684 | oxidoreductase activity, acting on peroxide as acceptor | 4/219 | 56/18410 | 0.00441 | 0.06143 | 4 | oligos5 | Oxidative stress |
| GO:0016655 | oxidoreductase activity, acting on NAD(P)H, quinone or similar compound as acceptor | 4/219 | 57/18410 | 0.0047 | 0.06143 | 4 | oligos5 | Oxidative stress |
| hsa00190 | Oxidative phosphorylation | 9/142 | 134/8166 | 0.00052 | 0.01189 | 9 | oligos5 | Oxidative stress |

**Table 7:** Differentially expressed genes in PD vs Control conditions per neuronal subtypes, states of astrocytes, states of microglia, states of oligodendrocytes.

| Gene | PD/Control coef | P value | Z score | Adjusted p value (BH correction) | subtype |
|---|---|---|---|---|---|
| OLFM1 | -0.26672 | 0 | -5.47691 | 0.00027 | neurons0 |
| CHST1 | -0.28907 | 0 | -5.0146 | 0.00165 | neurons0 |
| KBTBD6 | -0.29538 | 0 | -4.65221 | 0.00583 | neurons0 |
| ATCAY | -0.23614 | 0 | -4.62478 | 0.00583 | neurons0 |
| KIAA0586 | 0.18133 | 1E-05 | 4.53091 | 0.0073 | neurons0 |
| LSS | 0.17348 | 2E-05 | 4.27356 | 0.01993 | neurons0 |
| PLPPR4 | 0.1876 | 4E-05 | 4.09646 | 0.03726 | neurons0 |
| COL12A1 | 0.16096 | 5E-05 | 4.04658 | 0.04039 | neurons0 |
| N4BP2L2 | 0.10388 | 8E-05 | 3.95952 | 0.04816 | neurons0 |
| MFSD12 | -0.22816 | 8E-05 | -3.9521 | 0.04816 | neurons0 |
| KIFC3 | 0.10765 | 0.00011 | 3.86664 | 0.05632 | neurons0 |
| OGA | -0.15426 | 0.00013 | -3.83608 | 0.05632 | neurons0 |
| KIAA1549 | 0.12134 | 0.00013 | 3.83528 | 0.05632 | neurons0 |
| COG4 | -0.21418 | 0.00013 | -3.82149 | 0.05632 | neurons0 |
| CDIPT | -0.23319 | 0.00014 | -3.81555 | 0.05632 | neurons0 |
| RALGDS | 0.13355 | 0.00023 | 3.68738 | 0.08414 | neurons0 |
| CCNI | -0.22117 | 0.00025 | -3.65973 | 0.08414 | neurons0 |

| | | | | | |
|---|---|---|---|---|---|
| SMIM12 | -0.19965 | 0.00026 | -3.65469 | 0.08414 | neurons0 |
| PLEKHA6 | 0.14284 | 0.00026 | 3.65134 | 0.08414 | neurons0 |
| RAB6B | -0.1408 | 0.00027 | -3.64184 | 0.08414 | neurons0 |
| CAMK2G | 0.32289 | 1E-05 | 4.39478 | 0.06894 | neurons2 |
| NEAT1 | 0.74493 | 1E-05 | 4.35672 | 0.05952 | neurons3 |
| CA2 | 0.64802 | 2E-05 | 4.2746 | 0.05952 | neurons3 |
| SEPT8 | 0.78399 | 1E-05 | 4.53591 | 0.02647 | neurons4 |
| ANAPC16 | 0.76649 | 1E-05 | 4.45177 | 0.02647 | neurons4 |
| HP1BP3 | 0.56645 | 4E-05 | 4.12389 | 0.0772 | neurons4 |
| PLEKHA6 | 0.80128 | 1E-05 | 4.34902 | 0.05143 | neurons5 |
| HSPA1A | 0.47449 | 2E-05 | 4.22957 | 0.05143 | neurons5 |
| CSDC2 | 0.50572 | 2E-05 | 4.21644 | 0.05143 | neurons5 |
| NKIRAS1 | 0.37936 | 6E-05 | 4.00024 | 0.09835 | neurons5 |
| SAMD5 | -0.54796 | 0 | -4.8128 | 0.00601 | astrocytes0 |
| SLC9C2 | 0.26481 | 0 | 4.79957 | 0.00601 | astrocytes0 |
| ABCC4 | 0.22855 | 0 | 4.59018 | 0.01116 | astrocytes0 |
| FAM184A | -0.35513 | 1E-05 | -4.43513 | 0.01739 | astrocytes0 |
| ITGB5 | 141 | 4E-05 | 4.09984 | 0.05762 | astrocytes0 |
| KCNC4 | 0.13042 | 5E-05 | 4.07636 | 0.05762 | astrocytes0 |
| MUTYH | 0.12335 | 6E-05 | 4.0064 | 0.06023 | astrocytes0 |
| RGS9 | 0.24528 | 6E-05 | 3.99848 | 0.06023 | astrocytes0 |
| IFI27L1 | 0.10907 | 8E-05 | 3.94526 | 0.06694 | astrocytes0 |
| RHBDD1 | -0.1199 | 9E-05 | -3.91611 | 0.06801 | astrocytes0 |
| RSU1 | -0.12809 | 0.00011 | -3.86309 | 0.07087 | astrocytes0 |
| ITIH5 | 0.30553 | 0.00011 | 3.86186 | 0.07087 | astrocytes0 |
| FAM227A | 0.23401 | 0.00013 | 3.8188 | 0.07797 | astrocytes0 |
| SPRED1 | -0.18839 | 0.00014 | -3.7998 | 0.07818 | astrocytes0 |
| CRIP2 | 0.14677 | 0.00017 | 3.76171 | 0.08503 | astrocytes0 |
| NOL3 | -0.1695 | 0 | -4.89249 | 0.00753 | astrocytes1 |
| AL139158.2 | -0.14618 | 2E-05 | -4.23608 | 0.08595 | astrocytes1 |
| TMEM178B | -0.17256 | 4E-05 | -4.13535 | 0.08879 | astrocytes1 |
| CCDC3 | 0.31668 | 5E-05 | 4.0701 | 0.08879 | astrocytes1 |
| TTLL4 | -0.42099 | 0 | -4.59265 | 0.01702 | astrocytes2 |
| ABHD6 | 0.3516 | 0 | 4.58665 | 0.01702 | astrocytes2 |

| | | | | | |
|---|---|---|---|---|---|
| UST | -0.49356 | 2E-05 | -4.22894 | 0.05915 | astrocytes2 |
| MED14 | 0.30989 | 5E-05 | 4.04989 | 0.09682 | astrocytes2 |
| HSD17B12 | -0.31165 | 0 | -5.2138 | 0.0014 | astrocytes3 |
| MYO9A | -0.31693 | 0 | -4.87557 | 0.0041 | astrocytes3 |
| SYNE3 | -0.27603 | 1E-05 | -4.53623 | 0.01443 | astrocytes3 |
| SIK2 | -0.34147 | 1E-05 | -4.36434 | 0.02152 | astrocytes3 |
| CD38 | -0.28929 | 1E-05 | -4.34014 | 0.02152 | astrocytes3 |
| NUBPL | -0.30195 | 3E-05 | -4.14047 | 0.04366 | astrocytes3 |
| PPP2R2B | -0.62659 | 5E-05 | -4.063 | 0.04927 | astrocytes3 |
| AMZ1 | -0.30441 | 5E-05 | -4.04579 | 0.04927 | astrocytes3 |
| RBFOX2 | -0.35009 | 6E-05 | -4.01805 | 0.04928 | astrocytes3 |
| AC104596.1 | -0.29502 | 9E-05 | -3.92897 | 0.06448 | astrocytes3 |
| KLHL24 | 0.28695 | 0.00012 | 3.8519 | 0.0744 | astrocytes3 |
| FLG-AS1 | -0.48489 | 0.00012 | -3.85 | 0.0744 | astrocytes3 |
| POLR2A | -0.25021 | 0.00014 | -3.80811 | 0.08071 | astrocytes3 |
| PAAF1 | -0.22253 | 0.00015 | -3.79191 | 0.08071 | astrocytes3 |
| SPRED1 | -0.34877 | 0.00016 | -3.77267 | 0.08138 | astrocytes3 |
| SLTM | -0.2479 | 0.00019 | -3.73371 | 0.08512 | astrocytes3 |
| ACTN4 | 0.31108 | 2E-04 | 3.72359 | 0.08512 | astrocytes3 |
| BRINP2 | -0.28116 | 2E-04 | -3.71561 | 0.08512 | astrocytes3 |
| FAM217B | 0.30822 | 0.00023 | 3.68575 | 0.08963 | astrocytes3 |
| ANAPC16 | 0.42492 | 0.00024 | 3.67573 | 0.08963 | astrocytes3 |
| CSRP1 | 0.43871 | 0 | 5.59571 | 0.00017 | astrocytes4 |
| TP53BP2 | 0.2598 | 0 | 5.20475 | 0.00073 | astrocytes4 |
| ADORA1 | 0.28156 | 1E-05 | 4.33567 | 0.03661 | astrocytes4 |
| MATN2 | -0.24736 | 2E-05 | -4.22537 | 0.04507 | astrocytes4 |
| LIMK2 | -0.21737 | 5E-05 | -4.07577 | 0.06342 | astrocytes4 |
| PSD2 | 0.44303 | 7E-05 | 3.99352 | 0.06342 | astrocytes4 |
| 1 SOS | 0.18795 | 7E-05 | 3.98796 | 0.06342 | astrocytes4 |
| SERTAD2 | 0.19279 | 8E-05 | 3.94934 | 0.06342 | astrocytes4 |
| SCMH1 | -0.21422 | 8E-05 | -3.9345 | 0.06342 | astrocytes4 |
| RHBDD2 | 0.25137 | 1E-04 | 3.90276 | 0.06342 | astrocytes4 |
| SLC22A3 | 0.19068 | 1E-04 | 3.88381 | 0.06342 | astrocytes4 |
| NRSN1 | 0.24976 | 0.00011 | 3.86378 | 0.06342 | astrocytes4 |
| TMEM38A | 0.40441 | 0.00011 | 3.86122 | 0.06342 | astrocytes4 |
| MBOAT2 | 0.16859 | 0.00012 | 3.85133 | 0.06342 | astrocytes4 |
| SOX13 | 0.19235 | 0.00015 | 3.79501 | 0.07051 | astrocytes4 |
| PLOD3 | 0.22611 | 0.00015 | 3.7923 | 0.07051 | astrocytes4 |
| MIR34AHG | -0.27527 | 0.00017 | -3.76109 | 0.07521 | astrocytes4 |

| | | | | | |
|---|---|---|---|---|---|
| CLCN7 | 0.30673 | 0.00018 | 3.74568 | 0.07554 | astrocytes4 |
| LINC01301 | -0.84051 | 1E-05 | -4.37165 | 0.08794 | astrocytes5 |
| PTOV1 | 0.5708 | 2E-05 | 4.22326 | 0.08794 | astrocytes5 |
| CYP7B1 | -0.58807 | 5E-05 | -4.05885 | 0.08794 | astrocytes5 |
| RPS6KB1 | -0.48261 | 6E-05 | -4.02347 | 0.08794 | astrocytes5 |
| ZNF253 | 0.53904 | 6E-05 | 4.02009 | 0.08794 | astrocytes5 |
| PLXNA1 | 0.32742 | 0 | 5.51394 | 0.00022 | microglia1 |
| DLEU7-AS1 | -0.60573 | 0 | -5.33937 | 3E-04 | microglia1 |
| NCAM1 | 0.71476 | 0 | 4.92475 | 0.00181 | microglia1 |
| SLC24A2 | 0.27931 | 0 | 4.81228 | 0.00239 | microglia1 |
| MAP4K5 | 0.27615 | 0 | 4.73038 | 0.00287 | microglia1 |
| USP54 | 0.2552 | 0 | 4.66952 | 0.00312 | microglia1 |
| CCDC26 | -2.1687 | 0 | -4.64474 | 0.00312 | microglia1 |
| ATG10 | -0.33666 | 1E-05 | -4.5028 | 0.00537 | microglia1 |
| UTRN | 0.30394 | 1E-05 | 4.35941 | 0.00855 | microglia1 |
| SEC24C | 0.29088 | 1E-05 | 4.35463 | 0.00855 | microglia1 |
| TMBIM1 | 0.25421 | 1E-05 | 4.33011 | 0.00869 | microglia1 |
| STXBP3 | 0.23906 | 2E-05 | 4.25846 | 11 | microglia1 |
| SOCS5 | 0.26128 | 2E-05 | 4.22812 | 0.01162 | microglia1 |
| ABHD13 | 0.22609 | 3E-05 | 4.18863 | 0.01242 | microglia1 |
| LINC01004 | 0.23908 | 3E-05 | 4.1808 | 0.01242 | microglia1 |
| ZNF609 | -0.2808 | 5E-05 | -4.06872 | 0.0181 | microglia1 |
| JAK2 | -0.26623 | 5E-05 | -4.06518 | 0.0181 | microglia1 |
| TMOD2 | 0.27245 | 5E-05 | 4.04992 | 0.01813 | microglia1 |
| NACC2 | 0.24611 | 5E-05 | 4.03878 | 0.01813 | microglia1 |
| H2AFY | -0.32558 | 7E-05 | -3.96721 | 0.02331 | microglia1 |
| ZNRF2 | -0.30362 | 9E-05 | -3.92629 | 0.02522 | microglia1 |
| PHF3 | 0.23427 | 9E-05 | 3.92551 | 0.02522 | microglia1 |
| NDRG1 | 0.21139 | 1E-04 | 3.88946 | 0.02716 | microglia1 |
| SETBP1 | -0.39137 | 1E-04 | -3.88652 | 0.02716 | microglia1 |
| CRADD | 0.37713 | 0.00012 | 3.85174 | 0.02851 | microglia1 |
| POGLUT1 | 0.22349 | 0.00012 | 3.84699 | 0.02851 | microglia1 |
| NFKBIA | 0.35878 | 0.00012 | 3.84596 | 0.02851 | microglia1 |
| MNAT1 | -0.30939 | 0.00013 | -3.81971 | 0.03059 | microglia1 |
| TTYH3 | 0.22007 | 0.00014 | 3.80178 | 0.03172 | microglia1 |
| ZHX3 | 0.31845 | 0.00015 | 3.79365 | 0.03172 | microglia1 |
| KIAA0513 | 0.28326 | 0.00016 | 3.77682 | 0.03201 | microglia1 |
| XPR1 | 0.22121 | 0.00016 | 3.77538 | 0.03201 | microglia1 |
| MXI1 | 0.21683 | 0.00018 | 3.75237 | 0.03403 | microglia1 |

| | | | | | |
|---|---|---|---|---|---|
| GPM6B | 0.77696 | 0.00019 | 3.73857 | 0.0349 | microglia1 |
| UBAC2 | -0.24197 | 2E-04 | -3.72113 | 0.03633 | microglia1 |
| GAB1 | 0.26308 | 0.00021 | 3.70522 | 0.03653 | microglia1 |
| SORT1 | 0.21149 | 0.00021 | 3.70313 | 0.03653 | microglia1 |
| CHPT1 | 0.30577 | 0.00022 | 3.69891 | 0.03653 | microglia1 |
| PTBP1 | 0.2064 | 0.00026 | 3.65316 | 0.04258 | microglia1 |
| TBC1D9 | -0.24717 | 0.00027 | -3.64607 | 0.04263 | microglia1 |
| MIR222HG | -0.33024 | 0.00027 | -3.64002 | 0.04263 | microglia1 |
| SHOC2 | 0.21068 | 0.00035 | 3.57734 | 0.05298 | microglia1 |
| CLPTM1 | 0.24842 | 0.00039 | 3.54638 | 0.05823 | microglia1 |
| KIAA1551 | 0.40637 | 0.00041 | 3.53638 | 0.05911 | microglia1 |
| MRPL18 | 0.19482 | 0.00044 | 3.51664 | 0.06227 | microglia1 |
| WWP1 | 0.20854 | 0.00047 | 3.49974 | 0.06491 | microglia1 |
| HACD2 | 0.22345 | 0.00049 | 3.48453 | 0.06616 | microglia1 |
| AHDC1 | 0.21337 | 0.00051 | 3.47773 | 0.06616 | microglia1 |
| IL4R | -0.46619 | 0.00051 | -3.47708 | 0.06616 | microglia1 |
| SMARCA5 | 0.22175 | 0.00052 | 3.4723 | 0.06616 | microglia1 |
| CAMK1D | -0.33838 | 0.00053 | -3.46506 | 0.06642 | microglia1 |
| SP1 | 0.20032 | 0.00055 | 3.45623 | 0.06642 | microglia1 |
| CEBPG | 0.22948 | 0.00055 | 3.45558 | 0.06642 | microglia1 |
| SS18L1 | 249 | 0.00057 | 3.44765 | 0.06714 | microglia1 |
| EPS15 | 0.30877 | 0.00062 | 3.4246 | 0.07053 | microglia1 |
| C11orf80 | 0.26124 | 0.00062 | 3.42445 | 0.07053 | microglia1 |
| SEM1 | 0.24577 | 0.00065 | 3.41145 | 0.07268 | microglia1 |
| ST3GAL2 | 0.21566 | 7E-04 | 3.39028 | 0.07629 | microglia1 |
| IRS2 | 0.37506 | 7E-04 | 3.38876 | 0.07629 | microglia1 |
| ABCA1 | 0.34762 | 0.00074 | 3.37527 | 0.07778 | microglia1 |
| TOPORS | 0.20355 | 0.00074 | 3.37428 | 0.07778 | microglia1 |
| AK3 | 0.23244 | 0.00076 | 3.36747 | 0.07837 | microglia1 |
| MAPKAP1 | -0.19812 | 0.00078 | -3.36064 | 0.07837 | microglia1 |
| TMEM65 | 0.26591 | 8E-04 | 3.35283 | 0.07837 | microglia1 |
| EIF4A1 | 0.28029 | 0.00081 | 3.34886 | 0.07837 | microglia1 |
| NT5DC1 | 0.23808 | 0.00083 | 3.34317 | 0.07837 | microglia1 |
| NAF1 | 0.20368 | 0.00083 | 3.34276 | 0.07837 | microglia1 |
| ACTR2 | -0.2033 | 0.00084 | -3.34076 | 0.07837 | microglia1 |
| FARP1 | 0.42547 | 0.00084 | 3.33812 | 0.07837 | microglia1 |
| AP2A2 | 0.25694 | 0.00089 | 3.32407 | 0.08125 | microglia1 |
| KAT2B | 0.23219 | 0.00096 | 3.30133 | 0.08689 | microglia1 |
| DENND3 | -0.70845 | 0.00099 | -3.29306 | 0.08788 | microglia1 |
| MCM3AP | -0.2085 | 0.00101 | -3.28814 | 0.08788 | microglia1 |
| RNF19A | 0.23834 | 0.00101 | 3.28652 | 0.08788 | microglia1 |

| | | | | | |
|---|---|---|---|---|---|
| ARHGEF12 | 0.22063 | 0.00105 | 3.27613 | 0.08996 | microglia1 |
| ATF2 | 0.19078 | 0.00107 | 3.27181 | 0.09014 | microglia1 |
| KHDC4 | -0.23128 | 0.00109 | -3.2668 | 0.09056 | microglia1 |
| RALGAPA2 | 0.18573 | 0.00113 | 3.25689 | 0.09258 | microglia1 |
| AC012368.1 | 0.37485 | 0 | 5.59021 | 9E-05 | microglia3 |
| VPS53 | -0.34643 | 0 | -5.54677 | 9E-05 | microglia3 |
| PLCL1 | 0.71097 | 0 | 5.30359 | 0.00024 | microglia3 |
| CBLB | -0.47335 | 1E-05 | -4.44632 | 0.01092 | microglia3 |
| IRF2BP2 | 0.26187 | 1E-05 | 4.41512 | 0.01092 | microglia3 |
| DMD | -0.48102 | 1E-05 | -4.36217 | 0.01092 | microglia3 |
| RHOBTB3 | -0.50609 | 1E-05 | -4.35874 | 0.01092 | microglia3 |
| ST7 | -0.69111 | 1E-05 | -4.34384 | 0.01092 | microglia3 |
| RASAL2 | 0.5203 | 2E-05 | 4.32381 | 0.01092 | microglia3 |
| BCAT1 | -0.40645 | 2E-05 | -4.29234 | 0.01121 | microglia3 |
| TMCC3 | 0.38601 | 2E-05 | 4.27357 | 0.01121 | microglia3 |
| PRKCH | 0.57601 | 2E-05 | 4.24338 | 0.01176 | microglia3 |
| XPR1 | 0.26169 | 4E-05 | 4.13725 | 0.01629 | microglia3 |
| DSCAM | 0.74437 | 4E-05 | 4.13454 | 0.01629 | microglia3 |
| OXR1 | 0.48946 | 5E-05 | 4.05766 | 0.02002 | microglia3 |
| AKT3 | 0.38058 | 5E-05 | 4.05577 | 0.02002 | microglia3 |
| DLST | 0.18565 | 7E-05 | 3.96211 | 0.02802 | microglia3 |
| EMB | -0.55562 | 8E-05 | -3.93723 | 0.02936 | microglia3 |
| XPO1 | -0.24241 | 9E-05 | -3.90965 | 0.03119 | microglia3 |
| PYGL | -0.62151 | 1E-04 | -3.88367 | 0.03293 | microglia3 |
| KANSL1L | 0.30447 | 0.00011 | 3.87215 | 0.03293 | microglia3 |
| VWA8 | -0.47626 | 0.00014 | -3.81401 | 0.03984 | microglia3 |
| ASPH | -0.38205 | 0.00016 | -3.78105 | 0.04236 | microglia3 |
| SKP2 | -0.44947 | 0.00016 | -3.77724 | 0.04236 | microglia3 |
| ITPKB | 0.37446 | 0.00018 | 3.74016 | 0.04716 | microglia3 |
| PARVG | 0.29498 | 0.00023 | 3.68276 | 0.04877 | microglia3 |
| ICE1 | -0.21472 | 0.00024 | -3.66891 | 0.04877 | microglia3 |
| GALC | -0.24618 | 0.00025 | -3.66545 | 0.04877 | microglia3 |
| UBR5 | -0.20431 | 0.00025 | -3.66071 | 0.04877 | microglia3 |
| MICU3 | 0.21746 | 0.00025 | 3.65726 | 0.04877 | microglia3 |
| PELI1 | 0.52035 | 0.00026 | 3.65315 | 0.04877 | microglia3 |
| H3F3A | -0.26083 | 0.00026 | -3.65026 | 0.04877 | microglia3 |
| AC087500.1 | 0.27814 | 0.00027 | 3.64543 | 0.04877 | microglia3 |
| MARCH3 | 0.32579 | 0.00027 | 3.63897 | 0.04877 | microglia3 |
| ANK2 | -0.60537 | 0.00028 | -3.63378 | 0.04877 | microglia3 |
| FRG1-DT | 0.28253 | 0.00029 | 3.61985 | 0.04877 | microglia3 |

| | | | | | |
|---|---|---|---|---|---|
| NIPA2 | -0.18431 | 3E-04 | -3.61601 | 0.04877 | microglia3 |
| RFTN1 | -0.25219 | 3E-04 | -3.61543 | 0.04877 | microglia3 |
| TMEM65 | 0.30009 | 0.00031 | 3.60692 | 0.04877 | microglia3 |
| PDE4DIP | 0.38464 | 0.00032 | 3.60172 | 0.04877 | microglia3 |
| SMAD7 | 0.20234 | 0.00032 | 3.59899 | 0.04877 | microglia3 |
| AGTPBP1 | -0.42559 | 0.00032 | -3.59897 | 0.04877 | microglia3 |
| YWHAG | -0.31624 | 0.00033 | -3.58765 | 0.04889 | microglia3 |
| ITGAM | -0.23126 | 0.00034 | -3.58128 | 0.04889 | microglia3 |
| LPIN2 | 0.2405 | 0.00035 | 3.57696 | 0.04889 | microglia3 |
| ACSL3 | -0.21786 | 0.00035 | -3.57459 | 0.04889 | microglia3 |
| ASAP1 | -0.88432 | 0.00039 | -3.5492 | 0.05271 | microglia3 |
| SIGLEC8 | 0.35164 | 0.00041 | 3.53083 | 0.05371 | microglia3 |
| CLEC9A | -0.11673 | 0.00042 | -3.52471 | 0.05371 | microglia3 |
| EML1 | 0.22283 | 0.00045 | 3.51085 | 0.05371 | microglia3 |
| VASH1 | 0.2245 | 0.00045 | 3.51074 | 0.05371 | microglia3 |
| GAB2 | 0.27694 | 0.00045 | 3.50673 | 0.05371 | microglia3 |
| AGFG1 | -1.15215 | 0.00046 | -3.50169 | 0.05371 | microglia3 |
| MS4A14 | 0.35579 | 0.00047 | 3.49796 | 0.05371 | microglia3 |
| FCHSD2 | -0.78002 | 0.00047 | -3.4948 | 0.05371 | microglia3 |
| CAPZA1 | -0.2003 | 0.00048 | -3.49336 | 0.05371 | microglia3 |
| 1,00 PGK | -0.16679 | 0.00048 | -3.49306 | 0.05371 | microglia3 |
| TANC2 | 0.70644 | 0.00051 | 3.47334 | 0.05682 | microglia3 |
| CD2AP | -0.2129 | 0.00059 | -3.43738 | 0.06312 | microglia3 |
| CMTR2 | 0.1736 | 0.00059 | 3.43584 | 0.06312 | microglia3 |
| METTL7A | 0.20867 | 0.00061 | 3.42498 | 0.06413 | microglia3 |
| HBP1 | 0.19807 | 0.00062 | 3.42264 | 0.06413 | microglia3 |
| TMEM106A | -0.19835 | 0.00066 | -3.40644 | 0.06651 | microglia3 |
| MAML2 | 0.36112 | 0.00067 | 3.39988 | 0.06651 | microglia3 |
| EIF4G3 | -0.19468 | 0.00067 | -3.39981 | 0.06651 | microglia3 |
| ARHGDIB | 0.19355 | 7E-04 | 3.38913 | 0.06811 | microglia3 |
| STAB1 | -0.82052 | 0.00076 | -3.36623 | 0.07292 | microglia3 |
| CXCR4 | 0.50172 | 0.00081 | 3.34912 | 0.07556 | microglia3 |
| SKIL | 0.22627 | 0.00081 | 3.34824 | 0.07556 | microglia3 |
| OSBPL3 | 0.44288 | 0.00085 | 3.33741 | 0.07745 | microglia3 |
| BICRAL | 0.2219 | 0.00087 | 3.32829 | 0.0789 | microglia3 |
| SSBP1 | 0.17004 | 9E-04 | 3.31935 | 0.08034 | microglia3 |
| SORL1 | 447 | 0.00093 | 3.31091 | 0.08085 | microglia3 |
| RALGAPA2 | 0.47568 | 0.00093 | 3.30991 | 0.08085 | microglia3 |
| LY86 | 0.26719 | 0.00097 | 3.29979 | 0.08167 | microglia3 |
| RABEP2 | -0.16284 | 0.00097 | -3.29962 | 0.08167 | microglia3 |
| MTSS1 | -1.61208 | 0.00103 | -3.28255 | 0.08565 | microglia3 |

| | | | | | |
|---|---|---|---|---|---|
| FAM149A | 0.37105 | 0.00114 | 3.25268 | 0.09369 | microglia3 |
| ACPP | -0.17066 | 0.00117 | -3.24714 | 0.09369 | microglia3 |
| MYO5A | -0.90459 | 0.00118 | -3.24288 | 0.09369 | microglia3 |
| RAD23B | -0.18051 | 0.00121 | -3.23733 | 0.09369 | microglia3 |
| STON2 | -0.79218 | 0.00122 | -3.23444 | 0.09369 | microglia3 |
| DGKD | -0.2641 | 0.00123 | -3.23228 | 0.09369 | microglia3 |
| WDFY2 | -0.33013 | 0.00124 | -3.22982 | 0.09369 | microglia3 |
| TAF2 | -0.19117 | 0.00124 | -3.22902 | 0.09369 | microglia3 |
| UGCG | 0.42289 | 0.00128 | 3.21988 | 0.0956 | microglia3 |
| FAM126B | -0.19878 | 0.0013 | -3.21613 | 0.09574 | microglia3 |
| PILRA | 0.26224 | 0.00132 | 3.2109 | 0.09606 | microglia3 |
| RHOA | 0.16229 | 0.00134 | 3.20717 | 0.09606 | microglia3 |
| CSNK1A1 | -0.24804 | 0.00135 | -3.20546 | 0.09606 | microglia3 |
| DOCK1 | -0.68487 | 0.0014 | -3.19481 | 0.09858 | microglia3 |
| ST6GAL1 | -0.60245 | 0 | -4.71453 | 0.01553 | microglia5 |
| CCDC191 | -0.10931 | 0 | -5.01488 | 0.0037 | oligos0 |
| CTDSP2 | 0.19471 | 0 | 4.8858 | 0.0037 | oligos0 |
| SPESP1 | -0.10686 | 0 | -4.73482 | 0.00525 | oligos0 |
| KCNH1 | -0.1129 | 1E-05 | -4.54567 | 0.00983 | oligos0 |
| HS6ST1 | 0.15691 | 1E-05 | 4.47395 | 0.01103 | oligos0 |
| MRPS30-DT | -0.15653 | 2E-05 | -4.31855 | 0.01852 | oligos0 |
| SNX2 | 0.1079 | 2E-05 | 4.28782 | 0.01852 | oligos0 |
| GMDS-DT | -0.18796 | 2E-05 | -4.24877 | 0.0193 | oligos0 |
| CCDC32 | -0.12379 | 3E-05 | -4.14604 | 0.02627 | oligos0 |
| NDRG3 | 0.10286 | 4E-05 | 4.12814 | 0.02627 | oligos0 |
| METTL15 | -0.11497 | 5E-05 | -4.05742 | 0.0324 | oligos0 |
| USP40 | -0.19442 | 6E-05 | -4.02866 | 0.03291 | oligos0 |
| GOLGA1 | -0.11689 | 6E-05 | -4.01238 | 0.03291 | oligos0 |
| VGLL4 | 0.21794 | 7E-05 | 3.99364 | 0.03291 | oligos0 |
| ZNF420 | 0.09812 | 7E-05 | 3.97668 | 0.03291 | oligos0 |
| FRG1-DT | 0.13706 | 7E-05 | 3.96526 | 0.03291 | oligos0 |
| PARP4 | -0.09105 | 9E-05 | -3.92159 | 0.03583 | oligos0 |
| PTPDC1 | -0.2137 | 9E-05 | -3.91662 | 0.03583 | oligos0 |
| GGA3 | 0.10112 | 0.00012 | 3.84729 | 0.04297 | oligos0 |
| AAGAB | -0.11478 | 0.00012 | -3.84683 | 0.04297 | oligos0 |
| G3BP2 | -0.12592 | 0.00017 | -3.76396 | 0.05139 | oligos0 |
| CCDC102B | -0.19494 | 0.00017 | -3.76096 | 0.05139 | oligos0 |
| MIR181A2HG | -0.15928 | 0.00017 | -3.76023 | 0.05139 | oligos0 |
| GPRC5B | 0.34611 | 0.00017 | 3.75738 | 0.05139 | oligos0 |

| | | | | | |
|---|---|---|---|---|---|
| IKBKB | 0.0944 | 0.00018 | 3.74324 | 0.05182 | oligos0 |
| SUN2 | 0.37533 | 0.00019 | 3.73517 | 0.05182 | oligos0 |
| SCLT1 | -0.09996 | 0.00022 | -3.69579 | 0.05832 | oligos0 |
| LIX1-AS1 | -0.15237 | 0.00025 | -3.66523 | 0.06339 | oligos0 |
| SLC9B1 | -0.19787 | 0.00026 | -3.6527 | 0.06427 | oligos0 |
| ZNF143 | 0.09599 | 0.00028 | 3.63516 | 0.06588 | oligos0 |
| UNC79 | -0.36911 | 3E-04 | -3.61264 | 0.06588 | oligos0 |
| CES4A | -0.21239 | 0.00031 | -3.60995 | 0.06588 | oligos0 |
| ZMAT1 | -0.0979 | 0.00031 | -3.6095 | 0.06588 | oligos0 |
| S100PBP | -0.13195 | 0.00031 | -3.60526 | 0.06588 | oligos0 |
| ETFA | -0.14051 | 0.00037 | -3.56127 | 0.07517 | oligos0 |
| JARID2 | -0.12869 | 0.00038 | -3.55587 | 0.07517 | oligos0 |
| DNAJB14 | -0.11862 | 0.00042 | -3.52449 | 0.08237 | oligos0 |
| LRP4 | 0.37835 | 0.00049 | 3.48486 | 0.08817 | oligos0 |
| PITRM1 | -0.07455 | 0.00053 | -3.46735 | 0.08817 | oligos0 |
| POLR3G | -0.10461 | 0.00053 | -3.46672 | 0.08817 | oligos0 |
| CYP20A1 | 0.14689 | 0.00053 | 3.46284 | 0.08817 | oligos0 |
| CREBRF | 0.26314 | 0.00053 | 3.46277 | 0.08817 | oligos0 |
| PAXBP1-AS1 | -0.12311 | 0.00054 | -3.46194 | 0.08817 | oligos0 |
| EBF4 | 0.15717 | 0.00054 | 3.46003 | 0.08817 | oligos0 |
| MEIS1 | -0.18234 | 0.00057 | -3.44773 | 0.09024 | oligos0 |
| MDN1 | -0.10118 | 6E-04 | -3.4327 | 0.09332 | oligos0 |
| MSN | -0.08441 | 0.00065 | -3.4115 | 0.09694 | oligos0 |
| ATF1 | 0.12079 | 0.00066 | 3.40446 | 0.09694 | oligos0 |
| IRS2 | 0.17172 | 0.00066 | 3.40386 | 0.09694 | oligos0 |
| TIAM2 | 0.19023 | 7E-04 | 3.38964 | 0.09694 | oligos0 |
| NSRP1 | -0.10101 | 0.00071 | -3.38744 | 0.09694 | oligos0 |
| GPR75-ASB3 | 0.29619 | 0.00071 | 3.38632 | 0.09694 | oligos0 |
| POM121C | 0.07809 | 0.00072 | 3.38095 | 0.09694 | oligos0 |
| AC096677.1 | 0.15601 | 0.00073 | 3.37852 | 0.09694 | oligos0 |
| LARP1 | 0.16933 | 0.00076 | 3.3681 | 0.09813 | oligos0 |
| ZMAT3 | 0.08282 | 0.00077 | 3.36515 | 0.09813 | oligos0 |
| DZIP3 | -0.12132 | 0.00079 | -3.35466 | 0.09842 | oligos0 |
| FAM196B | -0.23226 | 0.00079 | -3.35462 | 0.09842 | oligos0 |
| CCDC7 | -0.18142 | 0.00083 | -3.34271 | 0.09978 | oligos0 |
| MAZ | 0.11087 | 0.00083 | 3.34144 | 0.09978 | oligos0 |
| CTDSP2 | 0.21654 | 0 | 5.62037 | 0.00011 | oligos1 |
| MEIS1 | -0.29298 | 0 | -5.54513 | 0.00011 | oligos1 |
| ICA1L | -0.24442 | 0 | -5.11026 | 0.00077 | oligos1 |
| SRBD1 | -0.15468 | 0 | -4.89253 | 0.00154 | oligos1 |

| | | | | | |
|---|---|---|---|---|---|
| SNX27 | 0.11794 | 0 | 4.87741 | 0.00154 | oligos1 |
| AMER2 | 0.46083 | 0 | 4.77161 | 0.00219 | oligos1 |
| TRAK2 | -0.16784 | 0 | -4.56958 | 0.00501 | oligos1 |
| IRS2 | 0.38168 | 1E-05 | 4.51348 | 0.00504 | oligos1 |
| EP300 | 0.12705 | 1E-05 | 4.51063 | 0.00504 | oligos1 |
| PHLPP2 | -0.10004 | 1E-05 | -4.49312 | 0.00504 | oligos1 |
| BOK | 0.40974 | 1E-05 | 4.45449 | 0.00533 | oligos1 |
| HS3ST5 | -0.32354 | 1E-05 | -4.44218 | 0.00533 | oligos1 |
| AC233296.1 | -0.58724 | 1E-05 | -4.41952 | 0.00547 | oligos1 |
| MNAT1 | -0.18912 | 1E-05 | -4.39131 | 0.00578 | oligos1 |
| MYO5A | -0.25951 | 1E-05 | -4.33176 | 0.00708 | oligos1 |
| DNAH17 | -0.41982 | 2E-05 | -4.30772 | 0.00741 | oligos1 |
| YPEL3 | 0.25165 | 2E-05 | 4.24581 | 0.00848 | oligos1 |
| POU2F1 | -0.10871 | 2E-05 | -4.22882 | 0.00848 | oligos1 |
| RHOBTB3 | 0.27766 | 2E-05 | 4.22838 | 0.00848 | oligos1 |
| PNPLA7 | 0.22779 | 2E-05 | 4.22777 | 0.00848 | oligos1 |
| ZCCHC17 | -0.22199 | 3E-05 | -4.20643 | 0.00887 | oligos1 |
| MIR181A2HG | -0.13784 | 3E-05 | -4.15776 | 0.01049 | oligos1 |
| SKP2 | -0.12017 | 4E-05 | -4.10316 | 0.01203 | oligos1 |
| TAF4 | 0.12648 | 4E-05 | 4.09429 | 0.01203 | oligos1 |
| 1,00 AMD | 0.35334 | 4E-05 | 4.08591 | 0.01203 | oligos1 |
| AGPAT4 | -0.73548 | 5E-05 | -4.07692 | 0.01203 | oligos1 |
| GLS | -0.2323 | 5E-05 | -4.07553 | 0.01203 | oligos1 |
| TRIP12 | -0.15357 | 5E-05 | -4.07065 | 0.01203 | oligos1 |
| PITRM1 | -0.13547 | 5E-05 | -4.05158 | 0.0126 | oligos1 |
| AGAP3 | 0.16443 | 6E-05 | 4.02629 | 0.01357 | oligos1 |
| MED28 | 0.08691 | 6E-05 | 4.00906 | 0.01383 | oligos1 |
| ZC3H12B | -0.15097 | 6E-05 | -4.00176 | 0.01383 | oligos1 |
| RASAL2 | -0.31537 | 6E-05 | -3.99928 | 0.01383 | oligos1 |
| USH1C | -0.87453 | 7E-05 | -3.98817 | 0.01407 | oligos1 |
| ST3GAL5 | -0.27061 | 9E-05 | -3.9215 | 0.01806 | oligos1 |
| PCBD2 | -0.17291 | 1E-04 | -3.9024 | 0.019 | oligos1 |
| ZNF358 | 0.14066 | 0.00012 | 3.85045 | 0.02289 | oligos1 |
| SLC9B1 | -0.26183 | 0.00012 | -3.83823 | 0.02343 | oligos1 |
| CDH11 | -0.3767 | 0.00014 | -3.80909 | 0.02414 | oligos1 |
| GLMN | -0.14446 | 0.00014 | -3.80891 | 0.02414 | oligos1 |
| SOX10 | 0.21778 | 0.00014 | 3.80761 | 0.02414 | oligos1 |
| LINC00320 | 0.24182 | 0.00014 | 3.8061 | 0.02414 | oligos1 |
| DDHD2 | -0.1622 | 0.00015 | -3.78773 | 0.02515 | oligos1 |
| DHX57 | -0.10736 | 0.00016 | -3.78063 | 0.02515 | oligos1 |
| LINC01170 | -0.86279 | 0.00016 | -3.77882 | 0.02515 | oligos1 |

| | | | | | |
|---|---|---|---|---|---|
| WDR26 | 0.09592 | 0.00016 | 3.76854 | 0.02564 | oligos1 |
| DCBLD2 | 0.14253 | 0.00018 | 3.74506 | 0.02641 | oligos1 |
| MORC4 | 0.13005 | 0.00018 | 3.74432 | 0.02641 | oligos1 |
| PNPLA6 | 0.18691 | 0.00018 | 3.74246 | 0.02641 | oligos1 |
| RPS6KA3 | 0.18138 | 0.00018 | 3.74024 | 0.02641 | oligos1 |
| HNRNPF | 0.12919 | 2E-04 | 3.72275 | 0.02701 | oligos1 |
| ZNF24 | 0.11874 | 2E-04 | 3.72205 | 0.02701 | oligos1 |
| KLF7 | -0.21081 | 2E-04 | -3.7199 | 0.02701 | oligos1 |
| LTBP3 | 0.13592 | 0.00021 | 3.71111 | 0.02745 | oligos1 |
| NSRP1 | -0.10391 | 0.00021 | -3.70608 | 0.02749 | oligos1 |
| UBE2J1 | -0.08303 | 0.00021 | -3.70155 | 0.02749 | oligos1 |
| SLC25A12 | -0.16611 | 0.00023 | -3.68372 | 0.02897 | oligos1 |
| TRPC1 | -0.22806 | 0.00024 | -3.67246 | 0.02975 | oligos1 |
| AIG1 | -0.29236 | 0.00025 | -3.6625 | 0.02997 | oligos1 |
| ANKRD52 | 0.08357 | 0.00025 | 3.66191 | 0.02997 | oligos1 |
| AC007364.1 | -0.40697 | 0.00027 | -3.64651 | 0.0313 | oligos1 |
| LINC01004 | 0.13748 | 0.00027 | 3.63954 | 0.03164 | oligos1 |
| DMXL1 | -0.21805 | 0.00029 | -3.62632 | 0.03278 | oligos1 |
| FBXW8 | -0.13608 | 0.00031 | -3.60781 | 0.03466 | oligos1 |
| KLF15 | 0.29287 | 0.00032 | 3.59614 | 0.03474 | oligos1 |
| OGT | 0.14844 | 0.00032 | 3.59522 | 0.03474 | oligos1 |
| PRKCE | 0.37865 | 0.00033 | 3.58939 | 0.03474 | oligos1 |
| MTFR1 | 0.15588 | 0.00033 | 3.58822 | 0.03474 | oligos1 |
| ANAPC16 | 0.33326 | 0.00033 | 3.58761 | 0.03474 | oligos1 |
| LINC01505 | -2.16783 | 0.00034 | -3.5813 | 0.03508 | oligos1 |
| MTR | -0.13063 | 0.00038 | -3.55271 | 0.03857 | oligos1 |
| AL445472.1 | -0.19941 | 0.00039 | -3.54449 | 0.03924 | oligos1 |
| HP1BP3 | 0.20082 | 0.00041 | 3.53333 | 0.04038 | oligos1 |
| MAVS | 0.14315 | 0.00042 | 3.52963 | 0.04039 | oligos1 |
| CACNG7 | 0.14924 | 0.00043 | 3.51812 | 0.0404 | oligos1 |
| CALCOCO1 | 0.10838 | 0.00044 | 3.51262 | 0.0404 | oligos1 |
| ZBTB7A | 0.29004 | 0.00045 | 3.51107 | 0.0404 | oligos1 |
| RELN | 0.20241 | 0.00045 | 3.50877 | 0.0404 | oligos1 |
| RAB9A | 0.13323 | 0.00047 | 3.49969 | 0.0404 | oligos1 |
| APOLD1 | 0.3016 | 0.00047 | 3.49733 | 0.0404 | oligos1 |
| NAPG | -0.09818 | 0.00047 | -3.49702 | 0.0404 | oligos1 |
| UACA | -0.20861 | 0.00047 | -3.4953 | 0.0404 | oligos1 |
| CAB39L | 0.32865 | 0.00048 | 3.49381 | 0.0404 | oligos1 |
| SMARCD1 | -0.09092 | 0.00048 | -3.49266 | 0.0404 | oligos1 |
| ZNF518A | -0.28002 | 0.00049 | -3.4862 | 0.0404 | oligos1 |
| RPAIN | -0.10346 | 0.00049 | -3.48618 | 0.0404 | oligos1 |

| | | | | | |
|---|---|---|---|---|---|
| LIMA1 | -0.2314 | 0.00049 | -3.48366 | 0.0404 | oligos1 |
| ARSB | 0.13511 | 0.00049 | 3.48351 | 0.0404 | oligos1 |
| CCNH | 0.17217 | 5E-04 | 3.48041 | 0.04041 | oligos1 |
| TTC13 | -0.11563 | 0.00051 | -3.47476 | 0.04055 | oligos1 |
| FTCDNL1 | -0.31831 | 0.00052 | -3.47163 | 0.04055 | oligos1 |
| LMO4 | 0.15877 | 0.00052 | 3.46817 | 0.04055 | oligos1 |
| DOPEY1 | -0.14498 | 0.00053 | -3.46612 | 0.04055 | oligos1 |
| LINC00511 | -633 | 0.00053 | -3.46481 | 0.04055 | oligos1 |
| CBX7 | 0.10778 | 0.00055 | 3.45538 | 0.04151 | oligos1 |
| GPRC5B | 0.31879 | 0.00056 | 3.45054 | 0.04151 | oligos1 |
| PPP1R12B | -0.34273 | 0.00056 | -3.45002 | 0.04151 | oligos1 |
| CEP97 | 0.13988 | 0.00057 | 3.44364 | 42 | oligos1 |
| EML4 | -0.10916 | 0.00058 | -3.44093 | 42 | oligos1 |
| PTOV1 | 0.18442 | 0.00058 | 3.43863 | 42 | oligos1 |
| GDPD1 | 0.09882 | 6E-04 | 3.43285 | 0.04204 | oligos1 |
| ZNF107 | -0.14731 | 6E-04 | -3.43158 | 0.04204 | oligos1 |
| JADE2 | 0.28196 | 6E-04 | 3.43035 | 0.04204 | oligos1 |
| CREBBP | 0.16894 | 0.00061 | 3.4271 | 0.04214 | oligos1 |
| CSNK1E | 0.13149 | 0.00065 | 3.40824 | 0.04439 | oligos1 |
| ARAP1 | 0.08439 | 0.00066 | 3.40774 | 0.04439 | oligos1 |
| ZNF226 | -0.08329 | 0.00066 | -3.40463 | 0.04448 | oligos1 |
| SORT1 | 0.24869 | 0.00068 | 3.3964 | 0.0449 | oligos1 |
| ETFA | -0.19502 | 0.00068 | -3.39575 | 0.0449 | oligos1 |
| ZFP36L2 | 0.14436 | 0.00069 | 3.39449 | 0.0449 | oligos1 |
| CDK19 | 0.29654 | 0.00071 | 3.38718 | 0.0457 | oligos1 |
| C2CD6 | -0.32821 | 0.00071 | -3.38453 | 0.04573 | oligos1 |
| UBE2E1 | -0.1301 | 0.00073 | -3.37899 | 0.04625 | oligos1 |
| RGS11 | -0.19725 | 0.00075 | -3.37209 | 0.04696 | oligos1 |
| RAD17 | -0.07305 | 0.00075 | -3.36938 | 0.04696 | oligos1 |
| AL645568.1 | -0.12435 | 0.00076 | -3.36621 | 0.04696 | oligos1 |
| GREB1 | -0.21335 | 0.00077 | -3.36402 | 0.04696 | oligos1 |
| CPLANE1 | -0.11009 | 0.00077 | -3.36186 | 0.04696 | oligos1 |
| MAP6 | -0.15833 | 0.00078 | -3.3605 | 0.04696 | oligos1 |
| STARD7 | 0.18966 | 0.00079 | 3.3566 | 0.04723 | oligos1 |
| BCAS1 | -0.26502 | 8E-04 | -3.35397 | 0.04729 | oligos1 |
| ZNF793 | -0.07267 | 0.00082 | -3.34613 | 0.04822 | oligos1 |
| SEC23IP | -0.0709 | 0.00083 | -3.34403 | 0.04822 | oligos1 |
| TIAM1 | 0.23831 | 0.00084 | 3.33824 | 0.04884 | oligos1 |
| FAM69A | 0.14341 | 0.00088 | 3.32719 | 0.05 | oligos1 |
| SLC13A3 | -0.35567 | 0.00088 | -3.32547 | 0.05 | oligos1 |
| RANBP3 | 0.06811 | 0.00088 | 3.32503 | 0.05 | oligos1 |

| | | | | | |
|---|---|---|---|---|---|
| CEP164 | -0.11306 | 9E-04 | -3.32145 | 0.05005 | oligos1 |
| RFTN2 | 0.28956 | 9E-04 | 3.31942 | 0.05005 | oligos1 |
| KCNJ2 | 0.26321 | 0.00091 | 3.31826 | 0.05005 | oligos1 |
| ZBTB40 | 0.12091 | 0.00092 | 3.31282 | 0.05064 | oligos1 |
| ATG10 | -0.28871 | 0.00093 | -3.30973 | 0.05082 | oligos1 |
| ACVR2B | 0.09537 | 0.00096 | 3.30105 | 0.05202 | oligos1 |
| POLR3B | -0.08898 | 0.00098 | -3.29626 | 0.05251 | oligos1 |
| MAPKBP1 | 0.16901 | 0.00099 | 3.29424 | 0.05251 | oligos1 |
| TMEM232 | -0.31253 | 1 | -3.29152 | 0.05263 | oligos1 |
| GTF2H3 | -0.07101 | 0.00106 | -3.27499 | 0.0551 | oligos1 |
| MAZ | 0.16571 | 0.00106 | 3.27445 | 0.0551 | oligos1 |
| RRNAD1 | 0.08655 | 0.00108 | 3.26895 | 0.05544 | oligos1 |
| MGAT4A | 0.10173 | 0.00109 | 3.26689 | 0.05544 | oligos1 |
| WDR11-AS1 | -0.14277 | 0.00109 | -3.26667 | 0.05544 | oligos1 |
| JUP | 0.09248 | 0.0011 | 3.26389 | 0.05559 | oligos1 |
| LIN54 | -0.21145 | 0.00113 | -3.2555 | 0.05686 | oligos1 |
| PTPN12 | -0.16411 | 0.00125 | -3.2264 | 0.0614 | oligos1 |
| TMPO | 0.10252 | 0.00126 | 3.22461 | 0.0614 | oligos1 |
| GEMIN8 | -0.20271 | 0.00127 | -3.22223 | 0.0614 | oligos1 |
| UBE2Q2 | -0.16514 | 0.00127 | -3.222 | 0.0614 | oligos1 |
| EIF3J-DT | 0.11886 | 0.00128 | 3.22092 | 0.0614 | oligos1 |
| CHIC1 | -0.09422 | 0.00128 | -3.22014 | 0.0614 | oligos1 |
| PLEKHG1 | -0.62639 | 0.00129 | -3.21819 | 0.0614 | oligos1 |
| PDE4B | -0.26156 | 0.0013 | -3.21572 | 0.0614 | oligos1 |
| DECR1 | -0.07275 | 0.00131 | -3.21408 | 0.0614 | oligos1 |
| NEK3 | -0.16156 | 0.00131 | -3.21308 | 0.0614 | oligos1 |
| CCDC191 | -0.17268 | 0.00132 | -3.21195 | 0.0614 | oligos1 |
| SH3GL1 | 0.09603 | 0.00133 | 3.20899 | 0.0614 | oligos1 |
| MIER3 | 0.09756 | 0.00133 | 3.20862 | 0.0614 | oligos1 |
| VLDLR | -0.14976 | 0.00134 | -3.20684 | 0.0614 | oligos1 |
| NOS1AP | -0.22217 | 0.00139 | -3.1967 | 0.0632 | oligos1 |
| NTRK2 | 0.20592 | 0.00142 | 3.19109 | 0.06332 | oligos1 |
| MAP2K7 | 0.10763 | 0.00142 | 3.191 | 0.06332 | oligos1 |
| GMDS-DT | -0.20336 | 0.00143 | -3.18756 | 0.06332 | oligos1 |
| GCN1 | 0.08689 | 0.00144 | 3.18624 | 0.06332 | oligos1 |
| BAZ1A | 146 | 0.00145 | 3.18355 | 0.06332 | oligos1 |
| CPQ | 0.32319 | 0.00146 | 3.18179 | 0.06332 | oligos1 |
| INTU | -0.37512 | 0.00147 | -3.18021 | 0.06332 | oligos1 |
| USP33 | 0.07646 | 0.00148 | 3.17852 | 0.06332 | oligos1 |
| PPM1D | 0.13036 | 0.00149 | 3.17666 | 0.06332 | oligos1 |
| TNFRSF21 | -0.40815 | 0.0015 | -3.17386 | 0.06332 | oligos1 |

| | | | | | |
|---|---|---|---|---|---|
| TPCN1 | 0.24818 | 0.0015 | 3.17382 | 0.06332 | oligos1 |
| PLOD3 | 392 | 0.00151 | 3.17314 | 0.06332 | oligos1 |
| ULK1 | 0.12673 | 0.00151 | 3.17223 | 0.06332 | oligos1 |
| ZDHHC8 | 0.07628 | 0.00152 | 3.17005 | 0.06332 | oligos1 |
| OXCT1 | -0.14616 | 0.00153 | -3.16898 | 0.06332 | oligos1 |
| SPANXA2-OT1 | -0.23267 | 0.00153 | -3.16819 | 0.06332 | oligos1 |
| RBM23 | 0.13601 | 0.00155 | 3.16528 | 0.06359 | oligos1 |
| PIGN | -0.21478 | 0.00156 | -3.16305 | 0.06372 | oligos1 |
| INPP5A | 0.17925 | 0.0016 | 3.15619 | 0.06462 | oligos1 |
| SGK2 | -0.20044 | 0.0016 | -3.15568 | 0.06462 | oligos1 |
| HABP4 | -0.12629 | 0.00165 | -3.14735 | 0.06612 | oligos1 |
| CREBRF | 0.22636 | 0.00167 | 3.14283 | 0.06677 | oligos1 |
| GDE1 | 0.18749 | 0.00169 | 3.13987 | 0.06708 | oligos1 |
| NPEPPS | 0.13448 | 0.00171 | 3.13673 | 0.0672 | oligos1 |
| MTMR7 | -0.25235 | 0.00171 | -3.13575 | 0.0672 | oligos1 |
| GBE1 | -0.14736 | 0.00172 | -3.13453 | 0.0672 | oligos1 |
| SREK1IP1 | 0.10309 | 0.00174 | 3.13191 | 0.06735 | oligos1 |
| NBEA | 0.20163 | 0.00175 | 3.13016 | 0.06735 | oligos1 |
| DPYD | -0.46125 | 0.00176 | -3.12737 | 0.06735 | oligos1 |
| PUM2 | 0.07917 | 0.00177 | 3.12655 | 0.06735 | oligos1 |
| TMED7-TICAM2 | 0.14163 | 0.00178 | 3.12515 | 0.06735 | oligos1 |
| SIRT5 | -0.13015 | 0.0018 | -3.1219 | 0.06735 | oligos1 |
| PNRC1 | 0.17097 | 0.00181 | 3.11908 | 0.06735 | oligos1 |
| PPP2R2C | -0.33328 | 0.00181 | -3.11899 | 0.06735 | oligos1 |
| AC012593.1 | -0.57647 | 0.00183 | -3.11678 | 0.06735 | oligos1 |
| SMAP1 | -0.09964 | 0.00183 | -3.11642 | 0.06735 | oligos1 |
| ECHDC1 | -0.13643 | 0.00183 | -3.11619 | 0.06735 | oligos1 |
| FIRRE | -0.14174 | 0.00184 | -3.11528 | 0.06735 | oligos1 |
| IRF2BP2 | 0.16788 | 0.00187 | 3.1106 | 0.06808 | oligos1 |
| LEMD2 | 84 | 0.00191 | 3.10444 | 0.06916 | oligos1 |
| PEPD | 0.13681 | 0.00193 | 3.10027 | 0.06979 | oligos1 |
| CHRAC1 | 0.09237 | 0.00198 | 3.09367 | 0.071 | oligos1 |
| FKTN | -0.13051 | 0.00199 | -3.09102 | 0.07128 | oligos1 |
| GPR75-ASB3 | 0.17468 | 0.00202 | 3.08687 | 0.07154 | oligos1 |
| SHROOM1 | 0.19883 | 0.00203 | 3.08624 | 0.07154 | oligos1 |
| PIN4 | 0.10025 | 0.00204 | 3.08473 | 0.07154 | oligos1 |
| DHPS | 0.09775 | 0.00205 | 3.08299 | 0.07154 | oligos1 |
| LINC01102 | -0.28213 | 0.00205 | -3.08264 | 0.07154 | oligos1 |
| TNRC6B | 0.20184 | 0.00207 | 3.08006 | 0.07182 | oligos1 |
| AC096677.1 | 0.11497 | 0.00212 | 3.0734 | 0.07308 | oligos1 |

| | | | | | |
|---|---|---|---|---|---|
| AGO1 | -0.07755 | 0.00215 | -3.06893 | 0.0737 | oligos1 |
| CRIM1 | 0.12236 | 0.00215 | 3.06803 | 0.0737 | oligos1 |
| TRIM8 | 0.1328 | 0.00221 | 3.06055 | 0.07506 | oligos1 |
| CCDC82 | -0.20835 | 0.00222 | -3.05973 | 0.07506 | oligos1 |
| GTF2F2 | 0.10615 | 0.00225 | 3.05542 | 0.07568 | oligos1 |
| SCGB2B2 | -0.17081 | 0.00225 | -3.05444 | 0.07568 | oligos1 |
| ZRANB3 | -0.13466 | 0.00228 | -3.05092 | 0.07614 | oligos1 |
| PDK4 | 0.47077 | 0.00229 | 3.04983 | 0.07614 | oligos1 |
| MID1IP1 | 0.55121 | 0.0023 | 3.04789 | 0.07628 | oligos1 |
| GLTP | 0.19561 | 0.00233 | 3.04469 | 0.07649 | oligos1 |
| FAM66D | -0.31154 | 0.00234 | -3.04327 | 0.07649 | oligos1 |
| TMEM184B | -0.22046 | 0.00235 | -3.04258 | 0.07649 | oligos1 |
| RASGRF2 | -0.45616 | 0.00236 | -3.04044 | 0.07649 | oligos1 |
| ZNF33A | 0.14583 | 0.00237 | 3.0397 | 0.07649 | oligos1 |
| NUP93 | -0.11465 | 0.00237 | -3.03887 | 0.07649 | oligos1 |
| DUSP7 | 0.13448 | 0.00239 | 3.03673 | 0.0765 | oligos1 |
| ZNF318 | 0.08986 | 0.0024 | 3.03588 | 0.0765 | oligos1 |
| HS6ST1 | 0.15523 | 0.00241 | 3.0348 | 0.0765 | oligos1 |
| KANK4 | -0.49323 | 0.00253 | -3.02029 | 0.07964 | oligos1 |
| DENND4B | 0.10096 | 0.00253 | 3.01997 | 0.07964 | oligos1 |
| MFSD12 | 0.09286 | 0.00261 | 3.00986 | 0.08176 | oligos1 |
| ZCCHC8 | 0.08709 | 0.00262 | 3.00937 | 0.08176 | oligos1 |
| SPOP | 0.0847 | 0.00265 | 3.0054 | 0.08238 | oligos1 |
| CCDC112 | -0.14009 | 0.00267 | -3.00362 | 0.08238 | oligos1 |
| CLCN7 | 0.13746 | 0.00267 | 3.00314 | 0.08238 | oligos1 |
| TPPP | 0.24065 | 0.00269 | 3.00132 | 0.08245 | oligos1 |
| NLGN1 | 0.24086 | 0.0027 | 3.0002 | 0.08245 | oligos1 |
| FOCAD | -0.20159 | 0.00271 | -2.99896 | 0.08245 | oligos1 |
| ULK4 | -0.1133 | 0.00275 | -2.99438 | 0.08335 | oligos1 |
| PAK2 | 0.20977 | 0.00276 | 2.993 | 0.08338 | oligos1 |
| GIPC2 | 0.20724 | 0.0028 | 2.98856 | 0.08424 | oligos1 |
| TMCC2 | 0.18012 | 0.00286 | 2.98215 | 0.08488 | oligos1 |
| FAM13C | -0.19741 | 0.00288 | -2.9807 | 0.08488 | oligos1 |
| PPP1R2 | -0.09191 | 0.00288 | -2.9799 | 0.08488 | oligos1 |
| LINC00877 | -0.20768 | 0.00288 | -2.97985 | 0.08488 | oligos1 |
| USO1 | -0.15859 | 0.00289 | -2.97907 | 0.08488 | oligos1 |
| ZNF529 | -0.07255 | 0.0029 | -2.97865 | 0.08488 | oligos1 |
| MGAT1 | 0.1088 | 0.00294 | 2.97366 | 0.08593 | oligos1 |
| TRIM16L | -0.27331 | 0.00296 | -2.97206 | 0.08602 | oligos1 |
| PPP2R2B | -0.63353 | 0.00299 | -2.96921 | 0.08641 | oligos1 |
| GTPBP6 | 0.12788 | 3 | 2.9682 | 0.08641 | oligos1 |

| | | | | | |
|---|---|---|---|---|---|
| POLR3G | -0.22337 | 0.00306 | -2.96118 | 0.08805 | oligos1 |
| GFOD1 | -0.15683 | 0.00308 | -2.95966 | 0.08814 | oligos1 |
| DDHD1 | -0.17922 | 0.00311 | -2.95663 | 0.08861 | oligos1 |
| BDNF-AS | -0.1453 | 0.00312 | -2.95557 | 0.08861 | oligos1 |
| KLHL21 | 0.21846 | 0.00315 | 2.95318 | 0.08881 | oligos1 |
| PPP4R2 | 0.24312 | 0.00315 | 2.95244 | 0.08881 | oligos1 |
| ZNF326 | -0.09772 | 0.00319 | -2.9489 | 0.08948 | oligos1 |
| FBRSL1 | 0.11224 | 0.0032 | 2.94745 | 0.08949 | oligos1 |
| AZIN2 | -0.1791 | 0.00322 | -2.94614 | 0.08949 | oligos1 |
| ZNF333 | -0.07426 | 0.00323 | -2.94527 | 0.08949 | oligos1 |
| AC007376.2 | -0.24373 | 0.00326 | -2.94245 | 0.08996 | oligos1 |
| FBXW5 | 0.08235 | 0.00329 | 2.93918 | 0.09057 | oligos1 |
| PPP3CA | 0.14319 | 0.00336 | 2.93293 | 0.09206 | oligos1 |
| SLC25A53 | 0.11962 | 0.00344 | 2.92512 | 0.09404 | oligos1 |
| KIAA1324L | -0.22563 | 0.00349 | -2.92114 | 0.0948 | oligos1 |
| TBC1D32 | -0.16358 | 0.0035 | -2.92025 | 0.0948 | oligos1 |
| NCAM1 | 0.20701 | 0.00356 | 2.91516 | 0.09588 | oligos1 |
| TPRN | 0.14168 | 0.00356 | 2.91437 | 0.09588 | oligos1 |
| CLPB | 0.10344 | 0.00363 | 2.90846 | 0.09703 | oligos1 |
| COQ8A | 0.08973 | 0.00363 | 2.90832 | 0.09703 | oligos1 |
| KLHL23 | 0.12265 | 0.00371 | 2.90151 | 0.0988 | oligos1 |
| ECE1 | 0.17522 | 0.00374 | 2.8994 | 0.09905 | oligos1 |
| TOB1 | 0.14227 | 0.00375 | 2.89838 | 0.09905 | oligos1 |
| METTL7A | 0.43052 | 0.00376 | 2.89725 | 0.09905 | oligos1 |
| LIX1-AS1 | -0.22894 | 0.00379 | -2.89518 | 0.09934 | oligos1 |
| CDV3 | 0.20736 | 0 | 5.43156 | 0.00036 | oligos2 |
| UBE2E2 | -0.30967 | 0 | -5.32481 | 0.00036 | oligos2 |
| CDKN1B | 0.20562 | 0 | 5.15254 | 0.00052 | oligos2 |
| SSFA2 | 0.19613 | 0 | 5.13042 | 0.00052 | oligos2 |
| GIPC2 | 0.1982 | 0 | 4.70505 | 0.00365 | oligos2 |
| CDK14 | 0.26294 | 1E-05 | 4.39295 | 0.01186 | oligos2 |
| CAMK2G | 0.2368 | 1E-05 | 4.38573 | 0.01186 | oligos2 |
| NDRG3 | 0.19279 | 2E-05 | 4.31594 | 0.01218 | oligos2 |
| RHOBTB3 | 0.2713 | 2E-05 | 4.31452 | 0.01218 | oligos2 |
| TOMM40 | 0.18815 | 2E-05 | 4.30153 | 0.01218 | oligos2 |
| GPM6A | 0.81546 | 2E-05 | 4.26185 | 0.01324 | oligos2 |
| BIRC2 | 0.16621 | 3E-05 | 4.17604 | 0.01752 | oligos2 |
| ATP11C | 0.23694 | 3E-05 | 4.16085 | 0.01752 | oligos2 |
| CLK4 | 0.16699 | 4E-05 | 4.08846 | 0.0211 | oligos2 |
| AARS | 0.22386 | 4E-05 | 4.08503 | 0.0211 | oligos2 |

| | | | | | |
|---|---|---|---|---|---|
| FAM208A | 0.17419 | 5E-05 | 4.05277 | 0.02272 | oligos2 |
| ENSA | 0.17655 | 6E-05 | 4.00344 | 26 | oligos2 |
| POU3F3 | 0.14834 | 7E-05 | 3.98962 | 26 | oligos2 |
| ATF6B | 0.16164 | 7E-05 | 3.9771 | 26 | oligos2 |
| PLCH2 | -0.18191 | 7E-05 | -3.96825 | 26 | oligos2 |
| FBXO30 | 0.1668 | 8E-05 | 3.94197 | 0.0267 | oligos2 |
| ZNF281 | 0.15639 | 8E-05 | 3.93915 | 0.0267 | oligos2 |
| HNRNPU | 0.13868 | 9E-05 | 3.92506 | 0.02708 | oligos2 |
| TBC1D8 | 0.19958 | 9E-05 | 3.91322 | 0.02726 | oligos2 |
| THUMPD1 | 0.16432 | 1E-04 | 3.88743 | 0.02911 | oligos2 |
| ZNF549 | 0.15549 | 0.00011 | 3.85925 | 0.03142 | oligos2 |
| ALAD | 0.16895 | 0.00013 | 3.82793 | 0.03438 | oligos2 |
| TSPAN14 | 0.1472 | 0.00014 | 3.81211 | 0.0345 | oligos2 |
| TOPBP1 | 0.18651 | 0.00014 | 3.80942 | 0.0345 | oligos2 |
| LETMD1 | -0.19118 | 0.00015 | -3.79108 | 0.03591 | oligos2 |
| RAB9A | 0.16184 | 0.00016 | 3.76913 | 0.03796 | oligos2 |
| USP22 | 0.19625 | 0.00018 | 3.74048 | 0.0411 | oligos2 |
| FAM13A | 0.28945 | 0.00019 | 3.73351 | 0.0411 | oligos2 |
| MSH3 | -0.17997 | 2E-04 | -3.71517 | 0.04223 | oligos2 |
| RING1 | 0.16055 | 0.00021 | 3.71182 | 0.04223 | oligos2 |
| BROX | 0.14691 | 0.00022 | 3.69556 | 0.0436 | oligos2 |
| PRPF8 | 0.17309 | 0.00022 | 3.6896 | 0.0436 | oligos2 |
| TCOF1 | 0.1923 | 0.00025 | 3.6612 | 0.04679 | oligos2 |
| GPC5 | 0.77213 | 0.00025 | 3.65817 | 0.04679 | oligos2 |
| NONO | 0.14372 | 0.00027 | 3.64388 | 0.04719 | oligos2 |
| PPTC7 | 0.18154 | 0.00027 | 3.63999 | 0.04719 | oligos2 |
| TTC32 | 0.13639 | 0.00028 | 3.63691 | 0.04719 | oligos2 |
| PAQR3 | 0.14097 | 0.00029 | 3.62056 | 0.04911 | oligos2 |
| PLS3 | -0.28597 | 0.00033 | -3.5879 | 0.05442 | oligos2 |
| PRPF38A | 0.13758 | 0.00034 | 3.57992 | 0.05486 | oligos2 |
| THOC3 | 0.19918 | 0.00036 | 3.56853 | 0.05498 | oligos2 |
| SYVN1 | 0.12853 | 0.00036 | 3.56797 | 0.05498 | oligos2 |
| SATB2 | 0.19566 | 0.00055 | 3.45301 | 0.08296 | oligos2 |
| POLR1A | -0.15698 | 0.00058 | -3.44037 | 0.08516 | oligos2 |
| CDKN1C | 0.20622 | 0.00061 | 3.42584 | 0.0875 | oligos2 |
| MAP4K5 | 0.36465 | 0.00063 | 3.41663 | 0.0875 | oligos2 |
| TRNAU1AP | 0.14459 | 0.00064 | 3.41349 | 0.0875 | oligos2 |
| RBM22 | 0.13983 | 0.00065 | 3.41168 | 0.0875 | oligos2 |
| SLCO3A1 | -0.26184 | 0.00071 | -3.38678 | 0.09253 | oligos2 |
| BCAS3 | -0.23897 | 0.00071 | -3.38523 | 0.09253 | oligos2 |
| GABPB1 | 0.15441 | 0.00073 | 3.37688 | 0.09253 | oligos2 |

| | | | | | |
|---|---|---|---|---|---|
| LINC01088 | 0.38802 | 0.00073 | 3.37647 | 0.09253 | oligos2 |
| FMN2 | 0.79006 | 0.00076 | 3.36718 | 0.09405 | oligos2 |
| SRP19 | 0.13944 | 8E-04 | 3.35424 | 0.09689 | oligos2 |
| ARHGAP26 | 0.21618 | 0.00084 | 3.34073 | 0.09829 | oligos2 |
| AL035413.1 | 0.19705 | 0.00084 | 3.3395 | 0.09829 | oligos2 |
| ACAT2 | 0.13579 | 0.00085 | 3.33529 | 0.09829 | oligos2 |
| TMEM65 | 0.17658 | 0.00087 | 3.32975 | 0.09829 | oligos2 |
| WNK1 | 0.15289 | 0.00088 | 3.32768 | 0.09829 | oligos2 |
| AC105285.1 | 0.16835 | 0 | 4.991 | 0.00431 | oligos3 |
| HPS1 | 0.14125 | 0 | 4.75633 | 0.00485 | oligos3 |
| AL121761.1 | -0.18073 | 0 | -4.75105 | 0.00485 | oligos3 |
| MAP3K7 | 0.12679 | 1E-05 | 4.38395 | 0.02024 | oligos3 |
| AGO3 | 0.11354 | 1E-05 | 4.34244 | 0.02024 | oligos3 |
| IFIT1 | -0.0823 | 5E-05 | -4.062 | 0.05013 | oligos3 |
| MID1 | 0.32932 | 6E-05 | 4.02146 | 0.05013 | oligos3 |
| HECW2 | -0.29123 | 6E-05 | -4.01474 | 0.05013 | oligos3 |
| CPB2-AS1 | -0.54779 | 6E-05 | -4.00199 | 0.05013 | oligos3 |
| ANK3 | -0.7701 | 7E-05 | -3.97005 | 0.05162 | oligos3 |
| PNN | -0.12748 | 8E-05 | -3.93756 | 0.05375 | oligos3 |
| SUN2 | 0.36315 | 0.00012 | 3.85366 | 0.06965 | oligos3 |
| ST3GAL5 | -0.14988 | 0.00018 | -3.74508 | 0.09022 | oligos3 |
| NSF | -0.37066 | 0.00018 | -3.73978 | 0.09022 | oligos3 |
| ENOX2 | -0.35512 | 0.00019 | -3.72728 | 0.09022 | oligos3 |
| PSRC1 | -0.14018 | 2E-04 | -3.71782 | 0.09022 | oligos3 |
| ZDHHC8 | 0.09917 | 0.00025 | 3.66498 | 0.09885 | oligos3 |
| CNTLN | 0.30163 | 0.00025 | 3.65889 | 0.09885 | oligos3 |
| 1,00 AMD | 0.34269 | 0.00027 | 3.64621 | 0.09885 | oligos3 |
| ACSL4 | -0.2439 | 0.00028 | -3.63344 | 0.09885 | oligos3 |
| SCAMP2 | 0.16116 | 0.00029 | 3.61996 | 0.09885 | oligos3 |
| CES4A | -0.26971 | 0.00031 | -3.60517 | 0.09885 | oligos3 |
| PTPDC1 | -0.20312 | 0.00032 | -3.6014 | 0.09885 | oligos3 |
| MAP3K2 | 0.17676 | 0.00034 | 3.57987 | 0.09955 | oligos3 |
| MMS22L | 0.12736 | 0.00037 | 3.56171 | 0.09955 | oligos3 |
| TOX3 | 0.14069 | 0.00037 | 3.55865 | 0.09955 | oligos3 |
| KLHL32 | -0.41802 | 0.00037 | -3.55763 | 0.09955 | oligos3 |
| CNTLN | 0.22392 | 0 | 4.7708 | 0.01127 | oligos4 |
| RAB9A | 0.1398 | 0 | 4.61215 | 0.01127 | oligos4 |
| BOK | 0.22594 | 0 | 4.57752 | 0.01127 | oligos4 |
| NDUFA2 | 0.10238 | 1E-05 | 4.46859 | 0.01225 | oligos4 |

| | | | | | |
|---|---|---|---|---|---|
| ACAP3 | 0.27875 | 1E-05 | 4.45157 | 0.01225 | oligos4 |
| RAI14 | 0.18904 | 1E-05 | 4.34135 | 0.01695 | oligos4 |
| ATG10 | -0.32209 | 2E-05 | -4.22346 | 0.02469 | oligos4 |
| DENND4B | 0.15487 | 3E-05 | 4.17261 | 0.02704 | oligos4 |
| SAT2 | 0.08893 | 4E-05 | 4.11344 | 0.03111 | oligos4 |
| SEMA6A | -0.24587 | 5E-05 | -4.03589 | 0.03907 | oligos4 |
| SPANXA2-OT1 | -0.41165 | 6E-05 | -3.99496 | 0.04225 | oligos4 |
| FAM50A | 0.11343 | 8E-05 | 3.94671 | 0.04493 | oligos4 |
| HP1BP3 | 0.24337 | 8E-05 | 3.94048 | 0.04493 | oligos4 |
| NTM | 1.01881 | 9E-05 | 3.9166 | 0.04608 | oligos4 |
| CTPS2 | -0.16725 | 0.00012 | -3.8504 | 0.05412 | oligos4 |
| TMEM178B | -0.34837 | 0.00013 | -3.82789 | 0.05412 | oligos4 |
| VEZF1 | 0.10794 | 0.00013 | 3.82011 | 0.05412 | oligos4 |
| CORO2B | 0.27251 | 0.00014 | 3.8114 | 0.05412 | oligos4 |
| NUDT16L1 | 0.07458 | 0.00015 | 3.79475 | 0.05412 | oligos4 |
| SLC22A17 | 0.07947 | 0.00015 | 3.78995 | 0.05412 | oligos4 |
| CES4A | -0.25169 | 0.00018 | -3.74838 | 0.06088 | oligos4 |
| GDE1 | 0.10698 | 0.00021 | 3.70507 | 69 | oligos4 |
| ZMIZ2 | 0.14103 | 0.00032 | 3.59488 | 0.09731 | oligos4 |
| GPRC5B | 0.17621 | 0.00033 | 3.5944 | 0.09731 | oligos4 |
| AC015923.1 | -0.3268 | 0 | -5.49547 | 0.00028 | oligos5 |
| RALGDS | 0.26165 | 0 | 5.33951 | 0.00033 | oligos5 |
| CLIP4 | 0.24902 | 0 | 5.2253 | 0.00042 | oligos5 |
| FAM69A | 0.19359 | 0 | 4.94647 | 0.00136 | oligos5 |
| AL356599.1 | -0.20483 | 0 | -4.74381 | 0.00301 | oligos5 |
| TP53TG5 | -0.18099 | 1E-05 | -4.49506 | 0.00833 | oligos5 |
| PAPOLA | 0.15161 | 2E-05 | 4.23309 | 0.02101 | oligos5 |
| ZNF302 | -0.21417 | 3E-05 | -4.20799 | 0.02101 | oligos5 |
| RTF1 | -0.2059 | 3E-05 | -4.20315 | 0.02101 | oligos5 |
| PNRC1 | 0.17118 | 3E-05 | 4.16079 | 0.02278 | oligos5 |
| SCFD2 | -0.14508 | 5E-05 | -4.0682 | 0.03094 | oligos5 |
| TNKS2 | 0.12468 | 1E-04 | 3.89157 | 0.05962 | oligos5 |
| MAP2K4 | 0.12766 | 0.00011 | 3.86042 | 0.06254 | oligos5 |
| CDC37 | -0.20565 | 0.00013 | -3.82666 | 0.06316 | oligos5 |
| AEBP2 | 0.14922 | 0.00014 | 3.81465 | 0.06316 | oligos5 |
| LINC00877 | -0.13274 | 0.00015 | -3.78501 | 0.06316 | oligos5 |
| SH3GL3 | -0.17109 | 0.00015 | -3.78339 | 0.06316 | oligos5 |
| DDHD2 | -0.1254 | 0.00016 | -3.7777 | 0.06316 | oligos5 |
| CASC3 | -0.13376 | 0.00017 | -3.76206 | 0.06371 | oligos5 |

| | | | | | |
|---|---|---|---|---|---|
| WDR11-AS1 | -0.1561 | 0.00019 | -3.73069 | 0.0671 | oligos5 |
| SLC13A3 | -0.23127 | 0.00021 | -3.70293 | 0.0671 | oligos5 |
| HPRT1 | 0.12831 | 0.00021 | 3.70184 | 0.0671 | oligos5 |
| AZIN2 | -0.13976 | 0.00021 | -3.70086 | 0.0671 | oligos5 |
| AC009093.2 | -0.12642 | 0.00026 | -3.6504 | 0.07836 | oligos5 |

**Table 8:** Pathway (GO and KEGG) over-representation analysis on cell-type/cell-state specific genes up/down regulated in PD brain compared to controls as reported in **Table 7**.

| ID | Description | Gene Ratio | Background Ratio | P value | Adjusted P value (Benjamini-Hochberg correction) | geneID | Count | subtype | Group |
|---|---|---|---|---|---|---|---|---|---|
| GO:0006044 | N-acetylglucosamine metabolic process | 2/17 | 18/18800 | 0.00012 | 0.02087 | OGA/CHST1 | 2 | neurons0 | Sugar metabolism |
| GO:1901071 | glucosamine-containing compound metabolic process | 2/17 | 25/18800 | 0.00023 | 0.02087 | OGA/CHST1 | 2 | neurons0 | Sugar metabolism |
| GO:0006891 | intra-Golgi vesicle-mediated transport | 2/17 | 34/18800 | 0.00042 | 0.02589 | RAB6B/COG4 | 2 | neurons0 | Golgi vesicle |
| GO:0006040 | amino sugar metabolic process | 2/17 | 41/18800 | 0.00062 | 0.02828 | OGA/CHST1 | 2 | neurons0 | Sugar metabolism |
| GO:0006890 | retrograde vesicle-mediated transport, Golgi to endoplasmic reticulum | 2/17 | 52/18800 | 0.00099 | 0.03637 | RAB6B/COG4 | 2 | neurons0 | Golgi vesicle |
| hsa00564 | Glycerophospholipid metabolism | 1/6 | 98/8163 | 0.06993 | 0.09987 | CDIPT | 1 | neurons0 | Lipoprotein activity |
| GO:0051924 | regulation of calcium ion transport | 1/1 | 251/18800 | 0.01335 | 0.02499 | CAMK2G | 1 | neurons2 | Ion transport |
| GO:0010959 | regulation of metal ion transport | 1/1 | 403/18800 | 0.02144 | 0.02499 | CAMK2G | 1 | neurons2 | Ion transport |
| GO:0006816 | calcium ion transport | 1/1 | 424/18800 | 0.02255 | 0.02499 | CAMK2G | 1 | neurons2 | Ion transport |
| hsa04713 | Circadian entrainment | 1/1 | 97/8163 | 0.01188 | 0.02968 | CAMK2G | 1 | neurons2 | Circadian rhythm |

| | | | | | | | | | |
|---|---|---|---|---|---|---|---|---|---|
| hsa04750 | Inflammatory mediator regulation of TRP channels | 1/1 | 98/8163 | 0.01201 | 0.02968 | CAMK2G | 1 | neurons2 | Ion channels |
| GO:0000152 | nuclear ubiquitin ligase complex | 1/2 | 45/19594 | 0.00459 | 0.02982 | ANAPC16 | 1 | neurons4 | Ubiquitin activity |
| GO:0031461 | cullin-RING ubiquitin ligase complex | 1/2 | 175/19594 | 0.01778 | 0.03328 | ANAPC16 | 1 | neurons4 | Ubiquitin activity |
| GO:0000151 | ubiquitin ligase complex | 1/2 | 302/19594 | 0.03059 | 0.03615 | ANAPC16 | 1 | neurons4 | Ubiquitin activity |
| hsa04120 | Ubiquitin mediated proteolysis | 1/1 | 142/8163 | 0.0174 | 0.02174 | ANAPC16 | 1 | neurons4 | Ubiquitin activity |
| hsa05166 | Human T-cell leukemia virus 1 infection | 1/1 | 222/8163 | 0.0272 | 0.0272 | ANAPC16 | 1 | neurons4 | T cell activity |
| GO:0010968 | regulation of microtubule nucleation | 1/3 | 10/18800 | 0.00159 | 0.02125 | HSPA1A | 1 | neurons5 | Microtubule organization |
| GO:0051131 | chaperone-mediated protein complex assembly | 1/3 | 23/18800 | 0.00367 | 0.02376 | HSPA1A | 1 | neurons5 | Chaperone binding |
| GO:0051085 | chaperone cofactor-dependent protein refolding | 1/3 | 32/18800 | 0.0051 | 0.02564 | HSPA1A | 1 | neurons5 | Chaperone binding |
| GO:0031116 | positive regulation of microtubule polymerization | 1/3 | 33/18800 | 0.00526 | 0.02564 | HSPA1A | 1 | neurons5 | Microtubule organization |
| GO:0007020 | microtubule nucleation | 1/3 | 36/18800 | 0.00573 | 0.02564 | HSPA1A | 1 | neurons5 | Microtubule organization |
| GO:0031112 | positive regulation of microtubule polymerization or depolymerization | 1/3 | 37/18800 | 0.00589 | 0.02564 | HSPA1A | 1 | neurons5 | Microtubule organization |
| GO:0031113 | regulation of microtubule polymerization | 1/3 | 55/18800 | 0.00875 | 0.02803 | HSPA1A | 1 | neurons5 | Microtubule organization |
| GO:0061077 | chaperone-mediated protein folding | 1/3 | 70/18800 | 0.01113 | 0.03042 | HSPA1A | 1 | neurons5 | Chaperone binding |
| GO:0031397 | negative regulation of protein ubiquitination | 1/3 | 84/18800 | 0.01335 | 0.0342 | HSPA1A | 1 | neurons5 | Ubiquitin activity |
| GO:0046785 | microtubule polymerization | 1/3 | 84/18800 | 0.01335 | 0.0342 | HSPA1A | 1 | neurons5 | Microtubule organization |
| GO:0031110 | regulation of microtubule polymerization | 1/3 | 88/18800 | 0.01398 | 0.03538 | HSPA1A | 1 | neurons5 | Microtubule organization |

| | | | | | | | | | |
|---|---|---|---|---|---|---|---|---|---|
| | or depolymerization | | | | | | | | |
| GO:0032436 | positive regulation of proteasomal ubiquitin-dependent protein catabolic process | 1/3 | 92/18800 | 0.01461 | 0.03647 | HSPA1A | 1 | neurons5 | Ubiquitin activity |
| GO:2000060 | positive regulation of ubiquitin-dependent protein catabolic process | 1/3 | 108/18800 | 0.01714 | 0.03772 | HSPA1A | 1 | neurons5 | Ubiquitin activity |
| GO:0031109 | microtubule polymerization or depolymerization | 1/3 | 123/18800 | 0.0195 | 0.03951 | HSPA1A | 1 | neurons5 | Microtubule organization |
| GO:0032434 | regulation of proteasomal ubiquitin-dependent protein catabolic process | 1/3 | 137/18800 | 0.0217 | 0.0412 | HSPA1A | 1 | neurons5 | Ubiquitin activity |
| GO:1902850 | microtubule cytoskeleton organization involved in mitosis | 1/3 | 151/18800 | 0.0239 | 0.04254 | HSPA1A | 1 | neurons5 | Microtubule organization |
| GO:0070507 | regulation of microtubule cytoskeleton organization | 1/3 | 152/18800 | 0.02406 | 0.04254 | HSPA1A | 1 | neurons5 | Microtubule organization |
| GO:2000058 | regulation of ubiquitin-dependent protein catabolic process | 1/3 | 166/18800 | 0.02626 | 0.04376 | HSPA1A | 1 | neurons5 | Ubiquitin activity |
| GO:0052126 | movement in host environment | 1/3 | 177/18800 | 0.02798 | 0.04447 | HSPA1A | 1 | neurons5 | Iron transport |
| GO:0031396 | regulation of protein ubiquitination | 1/3 | 213/18800 | 0.03361 | 0.04655 | HSPA1A | 1 | neurons5 | Ubiquitin activity |
| GO:0032886 | regulation of microtubule-based process | 1/3 | 249/18800 | 0.03921 | 0.05186 | HSPA1A | 1 | neurons5 | Microtubule organization |
| GO:0034599 | cellular response to oxidative stress | 1/3 | 284/18800 | 0.04464 | 0.0548 | HSPA1A | 1 | neurons5 | Oxidative stress |
| GO:0043161 | proteasome-mediated ubiquitin-dependent | 1/3 | 414/18800 | 0.06462 | 0.06952 | HSPA1A | 1 | neurons5 | Ubiquitin activity |

| | | | | | | | | | |
|---|---|---|---|---|---|---|---|---|---|
| | protein catabolic process | | | | | | | | |
| GO:0006979 | response to oxidative stress | 1/3 | 433/18800 | 0.06752 | 0.07172 | HSPA1A | 1 | neurons5 | Oxidative stress |
| GO:0005925 | focal adhesion | 1/3 | 419/19594 | 0.06279 | 0.06411 | HSPA1A | 1 | neurons5 | Cell adhesion |
| GO:0032794 | GTPase activating protein binding | 1/4 | 15/18410 | 0.00326 | 0.05121 | NKIRAS1 | 1 | neurons5 | GTPase activity |
| GO:0044183 | protein folding chaperone | 1/4 | 43/18410 | 0.00931 | 0.05121 | HSPA1A | 1 | neurons5 | Chaperone binding |
| GO:0031625 | ubiquitin protein ligase binding | 1/4 | 298/18410 | 0.0632 | 0.08682 | HSPA1A | 1 | neurons5 | Ubiquitin activity |
| GO:0044389 | ubiquitin-like protein ligase binding | 1/4 | 317/18410 | 0.06712 | 0.08682 | HSPA1A | 1 | neurons5 | Ubiquitin activity |
| GO:0003924 | GTPase activity | 1/4 | 336/18410 | 0.07104 | 0.08682 | NKIRAS1 | 1 | neurons5 | GTPase activity |
| GO:0015079 | potassium ion transmembrane transporter activity | 2/13 | 154/18410 | 0.0051 | 0.06671 | SLC9C2/KCNC4 | 2 | astrocytes0 | Potassium transport |
| GO:0032356 | oxidized DNA binding | 1/13 | 10/18410 | 0.00704 | 0.06671 | MUTYH | 1 | astrocytes0 | Oxidative stress |
| GO:0015386 | potassium:proton antiporter activity | 1/13 | 11/18410 | 0.00774 | 0.06671 | SLC9C2 | 1 | astrocytes0 | Potassium transport |
| GO:0015385 | sodium:proton antiporter activity | 1/13 | 13/18410 | 0.00914 | 0.06671 | SLC9C2 | 1 | astrocytes0 | Sodium transport |
| GO:0022821 | potassium ion antiporter activity | 1/13 | 16/18410 | 0.01124 | 0.06671 | SLC9C2 | 1 | astrocytes0 | Potassium transport |
| GO:0022853 | active ion transmembrane transporter activity | 2/13 | 254/18410 | 0.01338 | 0.07176 | SLC9C2/ABCC4 | 2 | astrocytes0 | Ion transport |
| GO:0005251 | delayed rectifier potassium channel activity | 1/13 | 33/18410 | 0.02306 | 0.07501 | KCNC4 | 1 | astrocytes0 | Ion channels |
| GO:0051539 | 4 iron, 4 sulfur cluster binding | 1/13 | 41/18410 | 0.02858 | 0.08458 | MUTYH | 1 | astrocytes0 | Iron transport |
| GO:0046873 | metal ion transmembrane transporter activity | 2/13 | 428/18410 | 0.03551 | 0.08458 | SLC9C2/KCNC4 | 2 | astrocytes0 | Ion transport |
| GO:0051536 | iron-sulfur cluster binding | 1/13 | 67/18410 | 0.04631 | 0.09956 | MUTYH | 1 | astrocytes0 | Iron transport |
| hsa04510 | Focal adhesion | 1/4 | 201/8163 | 0.09493 | 0.11786 | ITGB5 | 1 | astrocytes0 | Cell adhesion |
| GO:1902176 | negative regulation of oxidative stress-induced | 1/2 | 20/18800 | 0.00213 | 0.01614 | NOL3 | 1 | astrocytes1 | Oxidative stress |

| | | | | | | | | | |
|---|---|---|---|---|---|---|---|---|---|
| | intrinsic apoptotic signaling pathway | | | | | | | | |
| GO:1902175 | regulation of oxidative stress-induced intrinsic apoptotic signaling pathway | 1/2 | 29/18800 | 0.00308 | 0.01729 | NOL3 | 1 | astrocytes1 | Oxidative stress |
| GO:0014808 | release of sequestered calcium ion into cytosol by sarcoplasmic reticulum | 1/2 | 35/18800 | 0.00372 | 0.01792 | NOL3 | 1 | astrocytes1 | Ion transport |
| GO:1903514 | release of sequestered calcium ion into cytosol by endoplasmic reticulum | 1/2 | 36/18800 | 0.00383 | 0.01792 | NOL3 | 1 | astrocytes1 | Ion transport |
| GO:0070296 | sarcoplasmic reticulum calcium ion transport | 1/2 | 42/18800 | 0.00446 | 0.01792 | NOL3 | 1 | astrocytes1 | Ion transport |
| GO:0008631 | intrinsic apoptotic signaling pathway in response to oxidative stress | 1/2 | 45/18800 | 0.00478 | 0.01792 | NOL3 | 1 | astrocytes1 | Oxidative stress |
| GO:1903202 | negative regulation of oxidative stress-induced cell death | 1/2 | 54/18800 | 0.00574 | 0.01805 | NOL3 | 1 | astrocytes1 | Oxidative stress |
| GO:0051055 | negative regulation of lipid biosynthetic process | 1/2 | 61/18800 | 0.00648 | 0.01931 | CCDC3 | 1 | astrocytes1 | Lipoprotein activity |
| GO:1903201 | regulation of oxidative stress-induced cell death | 1/2 | 77/18800 | 0.00817 | 0.02068 | NOL3 | 1 | astrocytes1 | Oxidative stress |
| GO:0046889 | positive regulation of lipid biosynthetic process | 1/2 | 85/18800 | 0.00902 | 0.02078 | CCDC3 | 1 | astrocytes1 | Lipoprotein activity |
| GO:1900407 | regulation of cellular response to oxidative stress | 1/2 | 92/18800 | 0.00976 | 0.0221 | NOL3 | 1 | astrocytes1 | Oxidative stress |
| GO:0036473 | cell death in response to oxidative stress | 1/2 | 98/18800 | 0.0104 | 0.02236 | NOL3 | 1 | astrocytes1 | Oxidative stress |

| | | | | | | | | | |
|---|---|---|---|---|---|---|---|---|---|
| GO:1902882 | regulation of response to oxidative stress | 1/2 | 101/18800 | 0.01072 | 0.02252 | NOL3 | 1 | astrocytes1 | Oxidative stress |
| GO:0045833 | negative regulation of lipid metabolic process | 1/2 | 105/18800 | 0.01114 | 0.02281 | CCDC3 | 1 | astrocytes1 | Lipoprotein activity |
| GO:0051209 | release of sequestered calcium ion into cytosol | 1/2 | 121/18800 | 0.01283 | 0.02419 | NOL3 | 1 | astrocytes1 | Ion transport |
| GO:0045834 | positive regulation of lipid metabolic process | 1/2 | 153/18800 | 0.01621 | 0.02664 | CCDC3 | 1 | astrocytes1 | Lipoprotein activity |
| GO:0097553 | calcium ion transmembrane import into cytosol | 1/2 | 154/18800 | 0.01632 | 0.02664 | NOL3 | 1 | astrocytes1 | Ion transport |
| GO:0060402 | calcium ion transport into cytosol | 1/2 | 171/18800 | 0.01811 | 0.02846 | NOL3 | 1 | astrocytes1 | Ion transport |
| GO:0046890 | regulation of lipid biosynthetic process | 1/2 | 175/18800 | 0.01853 | 0.02846 | CCDC3 | 1 | astrocytes1 | Lipoprotein activity |
| GO:0060401 | cytosolic calcium ion transport | 1/2 | 190/18800 | 0.02011 | 0.03017 | NOL3 | 1 | astrocytes1 | Ion transport |
| GO:0034599 | cellular response to oxidative stress | 1/2 | 284/18800 | 0.02999 | 0.03857 | NOL3 | 1 | astrocytes1 | Oxidative stress |
| GO:0070588 | calcium ion transmembrane transport | 1/2 | 314/18800 | 0.03313 | 0.0402 | NOL3 | 1 | astrocytes1 | Ion transport |
| GO:0007204 | positive regulation of cytosolic calcium ion concentration | 1/2 | 325/18800 | 0.03428 | 0.0402 | NOL3 | 1 | astrocytes1 | Ion transport |
| GO:0019216 | regulation of lipid metabolic process | 1/2 | 339/18800 | 0.03574 | 0.0408 | CCDC3 | 1 | astrocytes1 | Lipoprotein activity |
| GO:0051480 | regulation of cytosolic calcium ion concentration | 1/2 | 356/18800 | 0.03751 | 0.04172 | NOL3 | 1 | astrocytes1 | Ion transport |
| GO:0006816 | calcium ion transport | 1/2 | 424/18800 | 0.0446 | 0.04817 | NOL3 | 1 | astrocytes1 | Ion transport |
| GO:0006979 | response to oxidative stress | 1/2 | 433/18800 | 0.04553 | 0.04855 | NOL3 | 1 | astrocytes1 | Oxidative stress |
| GO:0006874 | cellular calcium ion homeostasis | 1/2 | 456/18800 | 0.04792 | 0.04906 | NOL3 | 1 | astrocytes1 | Ion transport |
| GO:0055074 | calcium ion homeostasis | 1/2 | 468/18800 | 0.04917 | 0.04994 | NOL3 | 1 | astrocytes1 | Ion transport |

| | | | | | | | | | |
|---|---|---|---|---|---|---|---|---|---|
| GO:0046461 | neutral lipid catabolic process | 1/4 | 41/18800 | 0.0087 | 0.05122 | ABHD6 | 1 | astrocytes2 | Lipoprotein activity |
| GO:0009395 | phospholipid catabolic process | 1/4 | 52/18800 | 0.01102 | 0.05122 | ABHD6 | 1 | astrocytes2 | Lipoprotein activity |
| GO:0046503 | glycerolipid catabolic process | 1/4 | 68/18800 | 0.01439 | 0.05122 | ABHD6 | 1 | astrocytes2 | Lipoprotein activity |
| GO:0046889 | positive regulation of lipid biosynthetic process | 1/4 | 85/18800 | 0.01796 | 0.05122 | ABHD6 | 1 | astrocytes2 | Lipoprotein activity |
| GO:0001676 | long-chain fatty acid metabolic process | 1/4 | 111/18800 | 0.02341 | 0.05122 | ABHD6 | 1 | astrocytes2 | Fatty acid metabolism |
| GO:0033559 | unsaturated fatty acid metabolic process | 1/4 | 115/18800 | 0.02425 | 0.05122 | ABHD6 | 1 | astrocytes2 | Fatty acid metabolism |
| GO:0006638 | neutral lipid metabolic process | 1/4 | 132/18800 | 0.02779 | 0.05122 | ABHD6 | 1 | astrocytes2 | Lipoprotein activity |
| GO:0045834 | positive regulation of lipid metabolic process | 1/4 | 153/18800 | 0.03216 | 0.05122 | ABHD6 | 1 | astrocytes2 | Lipoprotein activity |
| GO:0046890 | regulation of lipid biosynthetic process | 1/4 | 175/18800 | 0.03672 | 0.05452 | ABHD6 | 1 | astrocytes2 | Lipoprotein activity |
| GO:0044242 | cellular lipid catabolic process | 1/4 | 219/18800 | 0.04579 | 0.06319 | ABHD6 | 1 | astrocytes2 | Lipoprotein activity |
| GO:0006650 | glycerophospholipid metabolic process | 1/4 | 309/18800 | 0.06415 | 0.08679 | ABHD6 | 1 | astrocytes2 | Lipoprotein activity |
| GO:0016042 | lipid catabolic process | 1/4 | 327/18800 | 0.06779 | 0.08801 | ABHD6 | 1 | astrocytes2 | Lipoprotein activity |
| GO:0019216 | regulation of lipid metabolic process | 1/4 | 339/18800 | 0.07021 | 0.08801 | ABHD6 | 1 | astrocytes2 | Lipoprotein activity |
| GO:0006644 | phospholipid metabolic process | 1/4 | 388/18800 | 0.08004 | 0.08801 | ABHD6 | 1 | astrocytes2 | Lipoprotein activity |
| GO:0006631 | fatty acid metabolic process | 1/4 | 395/18800 | 0.08144 | 0.08801 | ABHD6 | 1 | astrocytes2 | Fatty acid metabolism |
| GO:0046486 | glycerolipid metabolic process | 1/4 | 396/18800 | 0.08164 | 0.08801 | ABHD6 | 1 | astrocytes2 | Lipoprotein activity |
| GO:0034703 | cation channel complex | 1/4 | 221/19594 | 0.04436 | 0.08362 | ABHD6 | 1 | astrocytes2 | Ion channels |
| GO:0034702 | ion channel complex | 1/4 | 294/19594 | 0.05869 | 0.08776 | ABHD6 | 1 | astrocytes2 | Ion channels |

| | | | | | | | | | |
|---|---|---|---|---|---|---|---|---|---|
| GO:0005874 | microtubule | 1/4 | 435/19594 | 0.0859 | 0.09173 | TTLL4 | 1 | astrocytes2 | Microtubule organization |
| GO:0034992 | microtubule organizing center attachment site | 1/18 | 11/19594 | 0.01006 | 0.12517 | SYNE3 | 1 | astrocytes3 | Microtubule organization |
| GO:0034993 | meiotic nuclear membrane microtubule tethering complex | 1/18 | 11/19594 | 0.01006 | 0.12517 | SYNE3 | 1 | astrocytes3 | Microtubule organization |
| GO:0106094 | nuclear membrane microtubule tethering complex | 1/18 | 11/19594 | 0.01006 | 0.12517 | SYNE3 | 1 | astrocytes3 | Microtubule organization |
| GO:0031461 | cullin-RING ubiquitin ligase complex | 2/18 | 175/19594 | 0.01105 | 0.12517 | ANAPC16/KLHL24 | 2 | astrocytes3 | Ubiquitin activity |
| GO:0005963 | magnesium-dependent protein serine/threonine phosphatase complex | 1/18 | 28/19594 | 0.02542 | 0.1316 | PPP2R2B | 1 | astrocytes3 | serine/threonine phosphatase complex |
| GO:0000151 | ubiquitin ligase complex | 2/18 | 302/19594 | 0.03079 | 0.14371 | ANAPC16/KLHL24 | 2 | astrocytes3 | Ubiquitin activity |
| GO:0031463 | Cul3-RING ubiquitin ligase complex | 1/18 | 36/19594 | 0.03257 | 0.14371 | KLHL24 | 1 | astrocytes3 | Ubiquitin activity |
| GO:0016620 | oxidoreductase activity, acting on the aldehyde or oxo group of donors, NAD or NADP as acceptor | 1/17 | 38/18410 | 0.03453 | 0.13646 | HSD17B12 | 1 | astrocytes3 | Oxidative stress |
| GO:0051539 | 4 iron, 4 sulfur cluster binding | 1/17 | 41/18410 | 0.03721 | 0.13646 | NUBPL | 1 | astrocytes3 | Iron transport |
| GO:0016903 | oxidoreductase activity, acting on the aldehyde or oxo group of donors | 1/17 | 46/18410 | 0.04166 | 0.13885 | HSD17B12 | 1 | astrocytes3 | Oxidative stress |
| GO:0045580 | regulation of T cell differentiation | 2/18 | 151/18800 | 0.00901 | 0.11738 | SOX13/SOS1 | 2 | astrocytes4 | T cell activity |
| GO:0030953 | astral microtubule organization | 1/18 | 10/18800 | 0.00954 | 0.11738 | LIMK2 | 1 | astrocytes4 | Microtubule organization |
| GO:0046643 | regulation of gamma-delta T cell activation | 1/18 | 11/18800 | 0.01048 | 0.11738 | SOX13 | 1 | astrocytes4 | T cell activity |
| GO:0042492 | gamma-delta T cell differentiation | 1/18 | 13/18800 | 0.01238 | 0.11738 | SOX13 | 1 | astrocytes4 | T cell activity |

| | | | | | | | | | |
|---|---|---|---|---|---|---|---|---|---|
| GO:0045619 | regulation of lymphocyte differentiation | 2/18 | 181/18800 | 0.01275 | 0.11738 | SOX13/SOS1 | 2 | astrocytes4 | Lymphocyte |
| GO:0042754 | negative regulation of circadian rhythm | 1/18 | 14/18800 | 0.01333 | 0.11738 | ADORA1 | 1 | astrocytes4 | Circadian rhythm |
| GO:0045187 | regulation of circadian sleep/wake cycle, sleep | 1/18 | 15/18800 | 0.01427 | 0.11738 | ADORA1 | 1 | astrocytes4 | Circadian rhythm |
| GO:0099509 | regulation of presynaptic cytosolic calcium ion concentration | 1/18 | 15/18800 | 0.01427 | 0.11738 | ADORA1 | 1 | astrocytes4 | Ion transport |
| GO:0050802 | circadian sleep/wake cycle, sleep | 1/18 | 17/18800 | 0.01616 | 0.11738 | ADORA1 | 1 | astrocytes4 | Circadian rhythm |
| GO:0055089 | fatty acid homeostasis | 1/18 | 17/18800 | 0.01616 | 0.11738 | ADORA1 | 1 | astrocytes4 | Fatty acid metabolism |
| GO:0042749 | regulation of circadian sleep/wake cycle | 1/18 | 19/18800 | 0.01804 | 0.11738 | ADORA1 | 1 | astrocytes4 | Circadian rhythm |
| GO:0022410 | circadian sleep/wake cycle process | 1/18 | 20/18800 | 0.01899 | 0.11738 | ADORA1 | 1 | astrocytes4 | Circadian rhythm |
| GO:0035813 | regulation of renal sodium excretion | 1/18 | 20/18800 | 0.01899 | 0.11738 | ADORA1 | 1 | astrocytes4 | Sodium transport |
| GO:0046629 | gamma-delta T cell activation | 1/18 | 20/18800 | 0.01899 | 0.11738 | SOX13 | 1 | astrocytes4 | T cell activity |
| GO:0035812 | renal sodium excretion | 1/18 | 22/18800 | 0.02086 | 0.11738 | ADORA1 | 1 | astrocytes4 | Sodium transport |
| GO:0042745 | circadian sleep/wake cycle | 1/18 | 23/18800 | 0.0218 | 0.11738 | ADORA1 | 1 | astrocytes4 | Circadian rhythm |
| GO:0097164 | ammonium ion metabolic process | 1/18 | 23/18800 | 0.0218 | 0.11738 | SLC22A3 | 1 | astrocytes4 | Ion transport |
| GO:0006813 | potassium ion transport | 2/18 | 243/18800 | 0.02221 | 0.11738 | TMEM38A/ADORA1 | 2 | astrocytes4 | Potassium transport |
| GO:0033081 | regulation of T cell differentiation in thymus | 1/18 | 25/18800 | 0.02368 | 0.11738 | 1 SOS | 1 | astrocytes4 | T cell activity |
| GO:0010882 | regulation of cardiac muscle contraction by calcium ion signaling | 1/18 | 26/18800 | 0.02461 | 0.11738 | TMEM38A | 1 | astrocytes4 | Ion transport |
| GO:0050995 | negative regulation of lipid catabolic process | 1/18 | 26/18800 | 0.02461 | 0.11738 | ADORA1 | 1 | astrocytes4 | Lipoprotein activity |
| GO:0030217 | T cell differentiation | 2/18 | 263/18800 | 0.02574 | 0.11738 | SOX13/SOS1 | 2 | astrocytes4 | T cell activity |

| | | | | | | | | | |
|---|---|---|---|---|---|---|---|---|---|
| GO:0050996 | positive regulation of lipid catabolic process | 1/18 | 28/18800 | 0.02648 | 0.11873 | ADORA1 | 1 | astrocytes4 | Lipoprotein activity |
| GO:0010880 | regulation of release of sequestered calcium ion into cytosol by sarcoplasmic reticulum | 1/18 | 31/18800 | 0.02928 | 0.12127 | TMEM38A | 1 | astrocytes4 | Ion transport |
| GO:1902105 | regulation of leukocyte differentiation | 2/18 | 283/18800 | 0.02948 | 0.12127 | SOX13/SOS1 | 2 | astrocytes4 | Leukocytes |
| GO:0051056 | regulation of small GTPase mediated signal transduction | 2/18 | 299/18800 | 0.03262 | 0.12127 | PSD2/SOS1 | 2 | astrocytes4 | GTPase activity |
| GO:0014808 | release of sequestered calcium ion into cytosol by sarcoplasmic reticulum | 1/18 | 35/18800 | 33 | 0.12127 | TMEM38A | 1 | astrocytes4 | Ion transport |
| GO:1903514 | release of sequestered calcium ion into cytosol by endoplasmic reticulum | 1/18 | 36/18800 | 0.03393 | 0.12127 | TMEM38A | 1 | astrocytes4 | Ion transport |
| GO:0005247 | voltage-gated chloride channel activity | 1/17 | 11/18410 | 0.01011 | 0.12796 | CLCN7 | 1 | astrocytes4 | Ion channels |
| GO:0008308 | voltage-gated anion channel activity | 1/17 | 17/18410 | 0.01559 | 0.12796 | CLCN7 | 1 | astrocytes4 | Ion channels |
| GO:0046527 | glucosyltransferase activity | 1/17 | 18/18410 | 0.0165 | 0.12796 | PLOD3 | 1 | astrocytes4 | Sugar metabolism |
| GO:1990381 | ubiquitin-specific protease binding | 1/17 | 21/18410 | 0.01922 | 0.12796 | RHBDD2 | 1 | astrocytes4 | Ubiquitin activity |
| GO:0071617 | lysophospholipid acyltransferase activity | 1/17 | 23/18410 | 0.02104 | 0.12796 | MBOAT2 | 1 | astrocytes4 | Lipoprotein activity |
| GO:0044539 | long-chain fatty acid import into cell | 1/4 | 17/18800 | 0.00361 | 0.05 | RPS6KB1 | 1 | astrocytes5 | Fatty acid metabolism |
| GO:0140354 | lipid import into cell | 1/4 | 18/18800 | 0.00382 | 0.05 | RPS6KB1 | 1 | astrocytes5 | Lipoprotein activity |
| GO:0071385 | cellular response to glucocorticoid stimulus | 1/4 | 53/18800 | 0.01123 | 0.05171 | RPS6KB1 | 1 | astrocytes5 | Sugar metabolism |
| GO:0046324 | regulation of glucose import | 1/4 | 60/18800 | 0.01271 | 0.05268 | RPS6KB1 | 1 | astrocytes5 | Sugar metabolism |

| | | | | | | | | | |
|---|---|---|---|---|---|---|---|---|---|
| GO:0048247 | lymphocyte chemotaxis | 1/4 | 64/18800 | 0.01355 | 0.05397 | CYP7B1 | 1 | astrocytes5 | Lymphocyte |
| GO:0015909 | long-chain fatty acid transport | 1/4 | 66/18800 | 0.01397 | 0.05397 | RPS6KB1 | 1 | astrocytes5 | Fatty acid metabolism |
| GO:0010827 | regulation of glucose transmembrane transport | 1/4 | 78/18800 | 0.01649 | 0.05795 | RPS6KB1 | 1 | astrocytes5 | Sugar metabolism |
| GO:0046323 | glucose import | 1/4 | 79/18800 | 0.0167 | 0.05795 | RPS6KB1 | 1 | astrocytes5 | Sugar metabolism |
| GO:0015908 | fatty acid transport | 1/4 | 99/18800 | 0.0209 | 0.05954 | RPS6KB1 | 1 | astrocytes5 | Fatty acid metabolism |
| GO:1904659 | glucose transmembrane transport | 1/4 | 113/18800 | 0.02383 | 0.05954 | RPS6KB1 | 1 | astrocytes5 | Sugar metabolism |
| GO:0072676 | lymphocyte migration | 1/4 | 119/18800 | 0.02508 | 0.05954 | CYP7B1 | 1 | astrocytes5 | Lymphocyte |
| GO:0051384 | response to glucocorticoid | 1/4 | 139/18800 | 0.02925 | 0.06076 | RPS6KB1 | 1 | astrocytes5 | Sugar metabolism |
| GO:0055088 | lipid homeostasis | 1/4 | 173/18800 | 0.03631 | 0.06265 | CYP7B1 | 1 | astrocytes5 | Lipoprotein activity |
| GO:0048017 | inositol lipid-mediated signaling | 1/4 | 181/18800 | 0.03796 | 0.06265 | RPS6KB1 | 1 | astrocytes5 | Lipoprotein activity |
| GO:0009749 | response to glucose | 1/4 | 189/18800 | 0.03961 | 0.06386 | RPS6KB1 | 1 | astrocytes5 | Sugar metabolism |
| GO:0030595 | leukocyte chemotaxis | 1/4 | 236/18800 | 0.04928 | 0.06923 | CYP7B1 | 1 | astrocytes5 | Leukocytes |
| GO:0032496 | response to lipopolysaccharide | 1/4 | 333/18800 | 69 | 0.08169 | RPS6KB1 | 1 | astrocytes5 | Lipoprotein activity |
| GO:0050900 | leukocyte migration | 1/4 | 384/18800 | 0.07924 | 0.08804 | CYP7B1 | 1 | astrocytes5 | Leukocytes |
| GO:0006869 | lipid transport | 1/4 | 391/18800 | 0.08064 | 0.08902 | RPS6KB1 | 1 | astrocytes5 | Lipoprotein activity |
| GO:0010876 | lipid localization | 1/4 | 446/18800 | 0.09158 | 0.09435 | RPS6KB1 | 1 | astrocytes5 | Lipoprotein activity |
| GO:0016712 | oxidoreductase activity, acting on paired donors, with incorporation or reduction of molecular oxygen, reduced flavin or flavoprotein as one donor, and incorporation of one atom of oxygen | 1/3 | 40/18410 | 0.0065 | 0.02971 | CYP7B1 | 1 | astrocytes5 | Oxidative stress |
| GO:0016709 | oxidoreductase activity, acting on paired donors, | 1/3 | 43/18410 | 0.00699 | 0.02971 | CYP7B1 | 1 | astrocytes5 | Oxidative stress |

| | | | | | | | | | |
|---|---|---|---|---|---|---|---|---|---|
| | with incorporation or reduction of molecular oxygen, NAD(P)H as one donor, and incorporation of one atom of oxygen | | | | | | | | |
| GO:0005506 | iron ion binding | 1/3 | 151/18410 | 0.02441 | 0.04149 | CYP7B1 | 1 | astrocytes5 | Iron transport |
| GO:0016705 | oxidoreductase activity, acting on paired donors, with incorporation or reduction of molecular oxygen | 1/3 | 177/18410 | 0.02857 | 0.04409 | CYP7B1 | 1 | astrocytes5 | Oxidative stress |
| GO:1905953 | negative regulation of lipid localization | 3/68 | 47/18800 | 0.00065 | 0.14502 | IRS2/NFKBIA/ABCA1 | 3 | microglia1 | Lipoprotein activity |
| GO:0051056 | regulation of small GTPase mediated signal transduction | 6/68 | 299/18800 | 0.00074 | 0.14502 | DENND3/ABCA1/MAPKAP1/RALGAPA2/ARHGEF12/SHOC2 | 6 | microglia1 | GTPase activity |
| GO:0004402 | histone acetyltransferase activity | 3/70 | 36/18410 | 0.00034 | 0.04925 | KAT2B/ATF2/MCM3AP | 3 | microglia1 | Acetyltransferase |
| GO:0061733 | peptide-lysine-N-acetyltransferase activity | 3/70 | 38/18410 | 4E-04 | 0.04925 | KAT2B/ATF2/MCM3AP | 3 | microglia1 | Acetyltransferase |
| GO:0034212 | peptide N-acetyltransferase activity | 3/70 | 46/18410 | 0.00071 | 0.05781 | KAT2B/ATF2/MCM3AP | 3 | microglia1 | Acetyltransferase |
| GO:0008080 | N-acetyltransferase activity | 3/70 | 65/18410 | 0.00194 | 0.11849 | KAT2B/ATF2/MCM3AP | 3 | microglia1 | Acetyltransferase |
| GO:0005096 | GTPase activator activity | 7/82 | 274/18410 | 0.00022 | 0.05102 | RALGAPA2/DOCK1/ARHGDIB/RABEP2/AGFG1/ASAP1/RASAL2 | 7 | microglia3 | GTPase activity |
| GO:0031267 | small GTPase binding | 6/82 | 267/18410 | 0.00123 | 0.07087 | DOCK1/MYO5A/SORL1/ARHGDIB/RHOBTB3/XPO1 | 6 | microglia3 | GTPase activity |
| GO:0034450 | ubiquitin-ubiquitin ligase activity | 2/82 | 13/18410 | 0.00148 | 0.07087 | PELI1/UBR5 | 2 | microglia3 | Ubiquitin activity |
| GO:0005041 | low-density lipoprotein particle receptor activity | 2/82 | 15/18410 | 0.00198 | 0.07087 | SORL1/STAB1 | 2 | microglia3 | Lipoprotein activity |
| GO:0051020 | GTPase binding | 6/82 | 298/18410 | 0.00214 | 0.07087 | DOCK1/MYO5A/SORL1/ARHGDIB/RHOBTB3/XPO1 | 6 | microglia3 | GTPase activity |

| | | | | | | | | | |
|---|---|---|---|---|---|---|---|---|---|
| GO:0030169 | low-density lipoprotein particle binding | 2/82 | 18/18410 | 0.00286 | 0.0738 | SORL1/STAB1 | 2 | microglia3 | Lipoprotein activity |
| GO:0030228 | lipoprotein particle receptor activity | 2/82 | 18/18410 | 0.00286 | 0.0738 | SORL1/STAB1 | 2 | microglia3 | Lipoprotein activity |
| GO:0030695 | GTPase regulator activity | 7/82 | 488/18410 | 0.00605 | 0.09362 | RALGAPA2/DOCK1/ARHGDIB/RABEP2/AGFG1/ASAP1/RASAL2 | 7 | microglia3 | GTPase activity |
| GO:0071813 | lipoprotein particle binding | 2/82 | 31/18410 | 0.00838 | 0.09722 | SORL1/STAB1 | 2 | microglia3 | Lipoprotein activity |
| GO:0071814 | protein-lipid complex binding | 2/82 | 31/18410 | 0.00838 | 0.09722 | SORL1/STAB1 | 2 | microglia3 | Lipoprotein activity |
| GO:0043130 | ubiquitin binding | 3/82 | 96/18410 | 0.00904 | 0.09986 | CXCR4/RAD23B/UBR5 | 3 | microglia3 | Ubiquitin activity |
| GO:0055029 | nuclear DNA-directed RNA polymerase complex | 7/233 | 100/19594 | 0.00019 | 0.02267 | GTF2F2/POLR3G/GTF2H3/CCNH/POLR3B/TAF4/MNAT1 | 7 | oligos1 | RNA polymerase complex |
| GO:0000428 | DNA-directed RNA polymerase complex | 7/233 | 101/19594 | 2E-04 | 0.02267 | GTF2F2/POLR3G/GTF2H3/CCNH/POLR3B/TAF4/MNAT1 | 7 | oligos1 | RNA polymerase complex |
| GO:0030880 | RNA polymerase complex | 7/233 | 105/19594 | 0.00026 | 0.02267 | GTF2F2/POLR3G/GTF2H3/CCNH/POLR3B/TAF4/MNAT1 | 7 | oligos1 | RNA polymerase complex |
| GO:0005963 | magnesium-dependent protein serine/threonine phosphatase complex | 4/233 | 28/19594 | 0.00032 | 0.02267 | PPP2R2B/PPP4R2/PPP2R2C/PPP3CA | 4 | oligos1 | serine/threonine phosphatase complex |
| GO:0000151 | ubiquitin ligase complex | 11/233 | 302/19594 | 0.00103 | 0.04685 | ANAPC16/FBXW8/SKP2/KLHL21/UBE2E1/USP33/FBXW5/CBX7/GLMN/SPOP/WDR26 | 11 | oligos1 | Ubiquitin activity |
| GO:0008287 | protein serine/threonine phosphatase complex | 4/233 | 50/19594 | 0.00292 | 0.08187 | PPP2R2B/PPP4R2/PPP2R2C/PPP3CA | 4 | oligos1 | serine/threonine phosphatase complex |
| GO:0000123 | histone acetyltransferase complex | 5/233 | 87/19594 | 0.00382 | 0.09865 | CREBBP/EP300/JADE2/TAF4/OGT | 5 | oligos1 | Acetyltransferase |
| GO:0008200 | ion channel inhibitor activity | 2/59 | 32/18410 | 0.00471 | 0.08726 | WNK1/ENSA | 2 | oligos2 | Ion channels |
| GO:0051087 | chaperone binding | 3/59 | 106/18410 | 0.00477 | 0.08726 | BIRC2/CDKN1B/SYVN1 | 3 | oligos2 | Chaperone binding |
| GO:0016248 | channel inhibitor activity | 2/59 | 33/18410 | 5 | 0.08726 | WNK1/ENSA | 2 | oligos2 | Ion channels |

| | | | | | | | | | |
|---|---|---|---|---|---|---|---|---|---|
| GO:0006892 | post-Golgi vesicle-mediated transport | 3/22 | 104/18800 | 0.00023 | 0.12762 | NSF/SCAMP2/ANK3 | 3 | oligos3 | Golgi vesicle |
| GO:0034992 | microtubule organizing center attachment site | 1/23 | 11/19594 | 0.01284 | 0.14866 | SUN2 | 1 | oligos3 | Microtubule organization |
| GO:0034993 | meiotic nuclear membrane microtubule tethering complex | 1/23 | 11/19594 | 0.01284 | 0.14866 | SUN2 | 1 | oligos3 | Microtubule organization |
| GO:0106094 | nuclear membrane microtubule tethering complex | 1/23 | 11/19594 | 0.01284 | 0.14866 | SUN2 | 1 | oligos3 | Microtubule organization |
| GO:0004709 | MAP kinase kinase kinase activity | 2/24 | 26/18410 | 0.00052 | 0.04155 | MAP3K7/MAP3K2 | 2 | oligos3 | MAP kinase activity |
| GO:0008017 | microtubule binding | 3/24 | 272/18410 | 0.00513 | 0.13193 | MID1/PSRC1/SUN2 | 3 | oligos3 | Microtubule organization |
| GO:0031957 | very long-chain fatty acid-CoA ligase activity | 1/24 | 10/18410 | 0.01296 | 0.13193 | ACSL4 | 1 | oligos3 | Fatty acid metabolism |
| GO:0004467 | long-chain fatty acid-CoA ligase activity | 1/24 | 15/18410 | 0.01938 | 0.13193 | ACSL4 | 1 | oligos3 | Fatty acid metabolism |
| GO:0004707 | MAP kinase activity | 1/24 | 15/18410 | 0.01938 | 0.13193 | MAP3K7 | 1 | oligos3 | MAP kinase activity |
| GO:0015645 | fatty acid ligase activity | 1/24 | 22/18410 | 0.02831 | 0.13193 | ACSL4 | 1 | oligos3 | Fatty acid metabolism |
| GO:0015036 | disulfide oxidoreductase activity | 1/24 | 42/18410 | 0.05337 | 0.13774 | ENOX2 | 1 | oligos3 | Oxidative stress |
| GO:0032482 | Rab protein signal transduction | 2/21 | 21/18800 | 0.00025 | 0.09683 | RAB9A/DENND4B | 2 | oligos4 | Rab protein signal transduction |
| GO:0034389 | lipid droplet organization | 1/17 | 26/18800 | 0.02326 | 0.12818 | DDHD2 | 1 | oligos5 | Lipoprotein activity |
| GO:0046461 | neutral lipid catabolic process | 1/17 | 41/18800 | 0.03645 | 0.12818 | DDHD2 | 1 | oligos5 | Lipoprotein activity |
| GO:0001913 | T cell mediated cytotoxicity | 1/17 | 50/18800 | 0.04428 | 0.13443 | HPRT1 | 1 | oligos5 | T cell activity |
| GO:0031122 | cytoplasmic microtubule organization | 1/17 | 57/18800 | 0.05033 | 0.13705 | CLIP4 | 1 | oligos5 | Microtubule organization |
| GO:0046503 | glycerolipid catabolic process | 1/17 | 68/18800 | 0.05977 | 0.14605 | DDHD2 | 1 | oligos5 | Lipoprotein activity |
| GO:0005793 | endoplasmic reticulum-Golgi | 2/18 | 131/19594 | 0.00633 | 0.1151 | DDHD2/AZIN2 | 2 | oligos5 | Golgi vesicle |

| | | | | | | | | | |
|---|---|---|---|---|---|---|---|---|---|
| | intermediate compartment | | | | | | | | |
| GO:0035371 | microtubule plus-end | 1/18 | 23/19594 | 0.02093 | 0.14127 | CLIP4 | 1 | oligos5 | Microtubule organization |
| GO:0004708 | MAP kinase kinase activity | 1/18 | 18/18410 | 0.01746 | 0.12481 | MAP2K4 | 1 | oligos5 | MAP kinase activity |
| GO:0000287 | magnesium ion binding | 2/18 | 222/18410 | 0.01951 | 0.12481 | PAPOLA/HPRT1 | 2 | oligos5 | Ion transport |
| GO:0051010 | microtubule plus-end binding | 1/18 | 21/18410 | 0.02034 | 0.12481 | CLIP4 | 1 | oligos5 | Microtubule organization |
| GO:0005343 | organic acid:sodium symporter activity | 1/18 | 30/18410 | 0.02894 | 0.12481 | SLC13A3 | 1 | oligos5 | Sodium transport |
