## Supplementary information for "Unravelling cell type specific response to Parkinson’s Disease at single cell resolution"

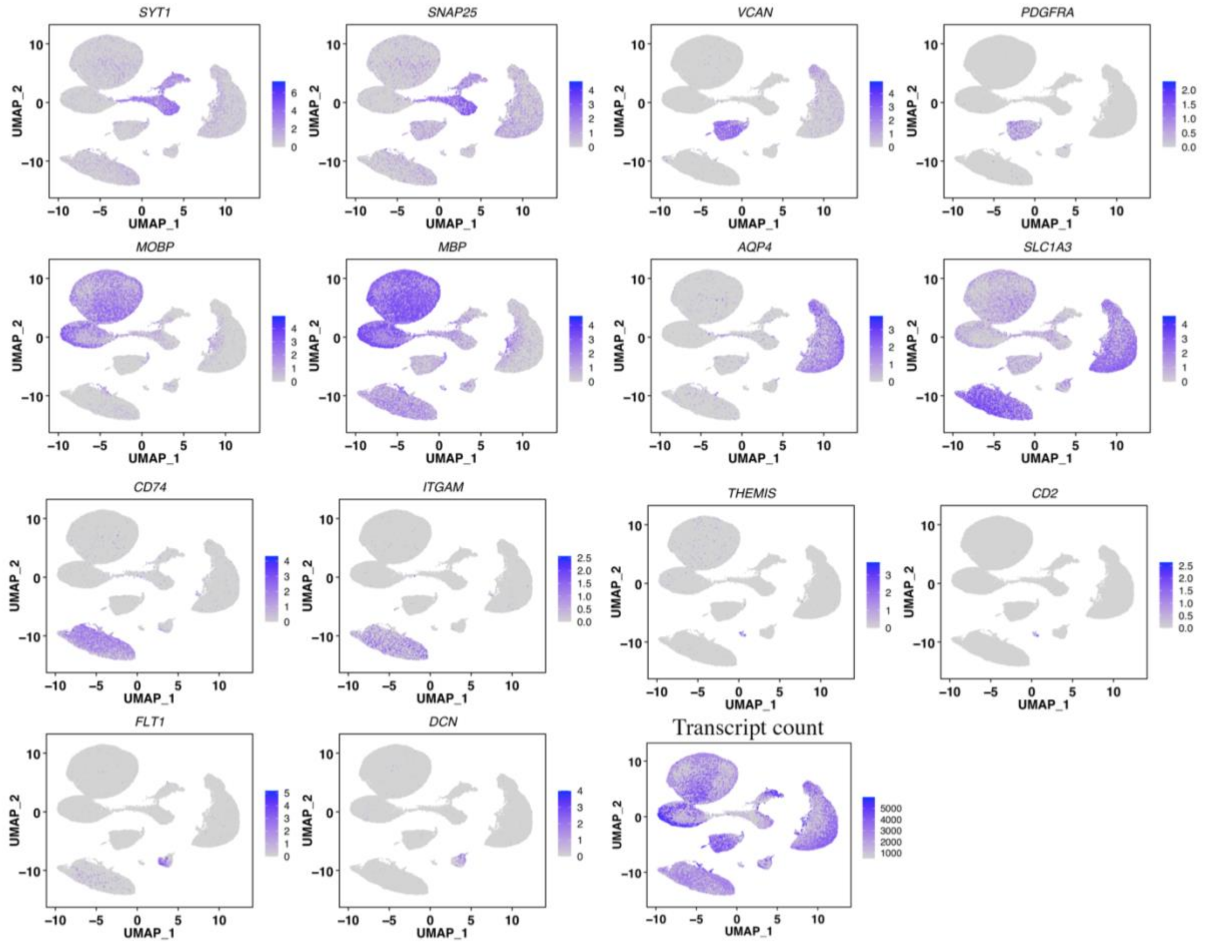


**Supplementary Figure 1:** High level cell type markers**.** Ln-normalized SCT counts for gene expression levels are shown.

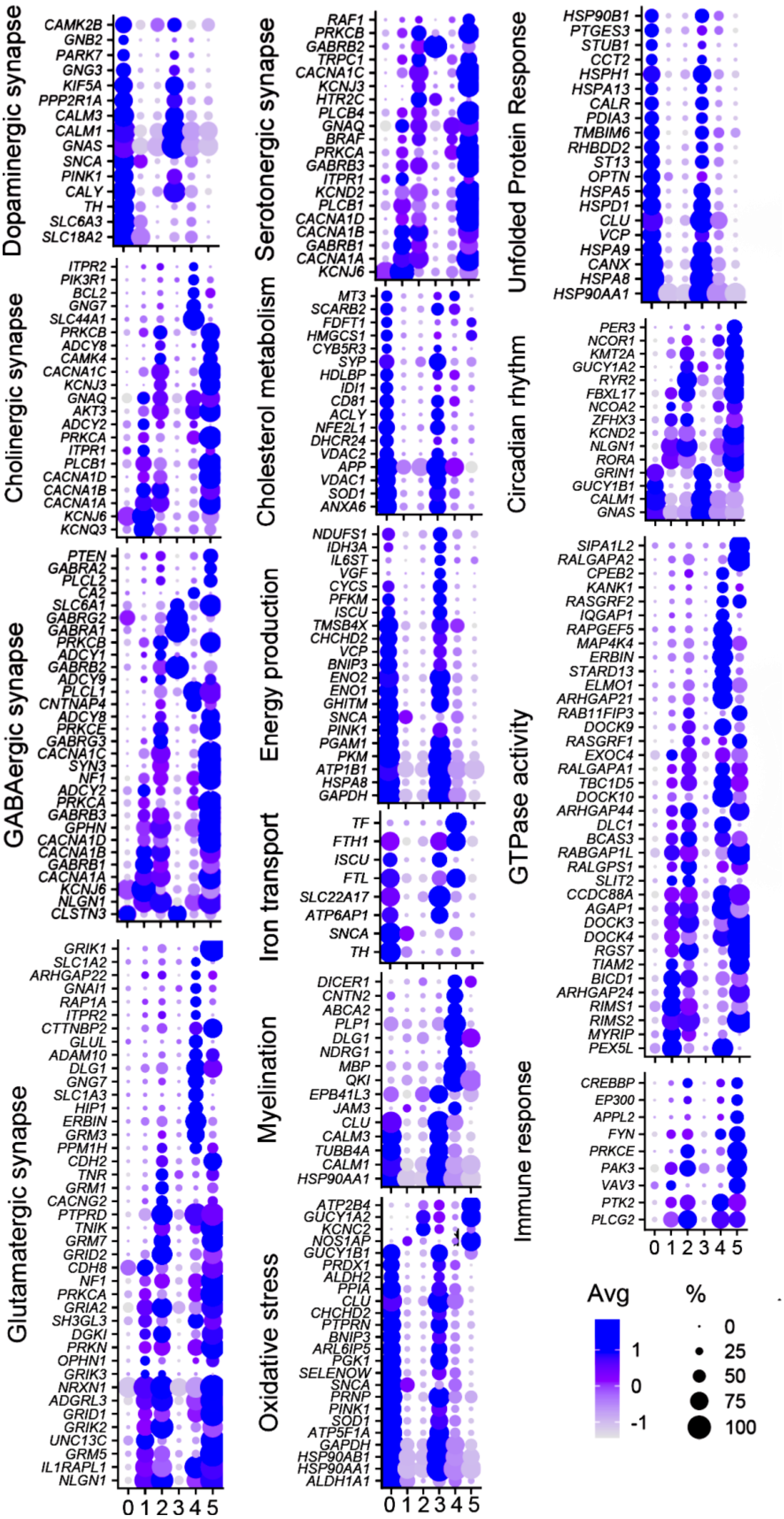


**Supplementary Figure 2:** Markers of neuronal populations grouped by enriched terms reported in **Table 6.**

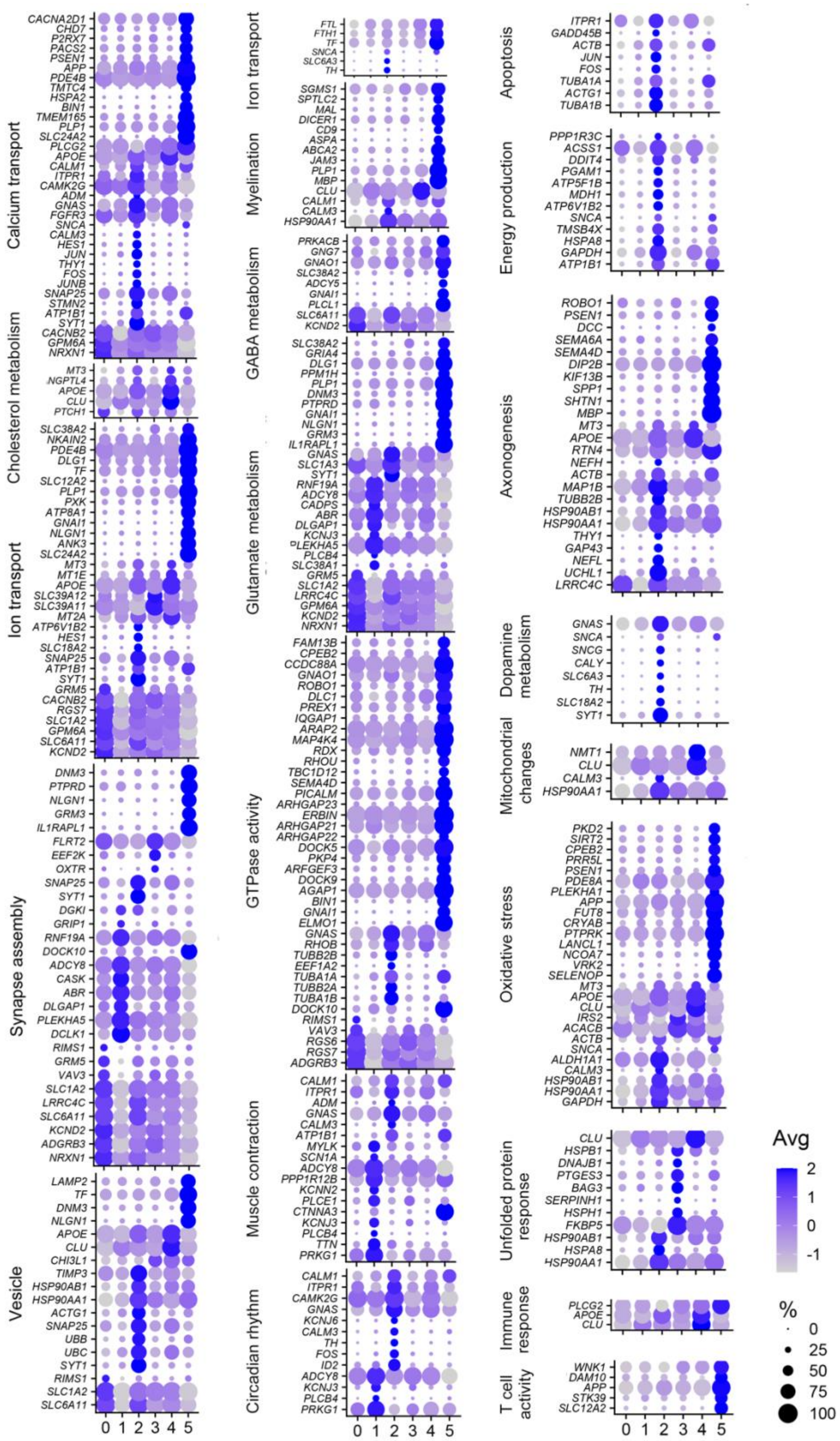


**Supplementary Figure 3:** Markers of astrocytic states grouped by enriched terms reported in **Table 6.**

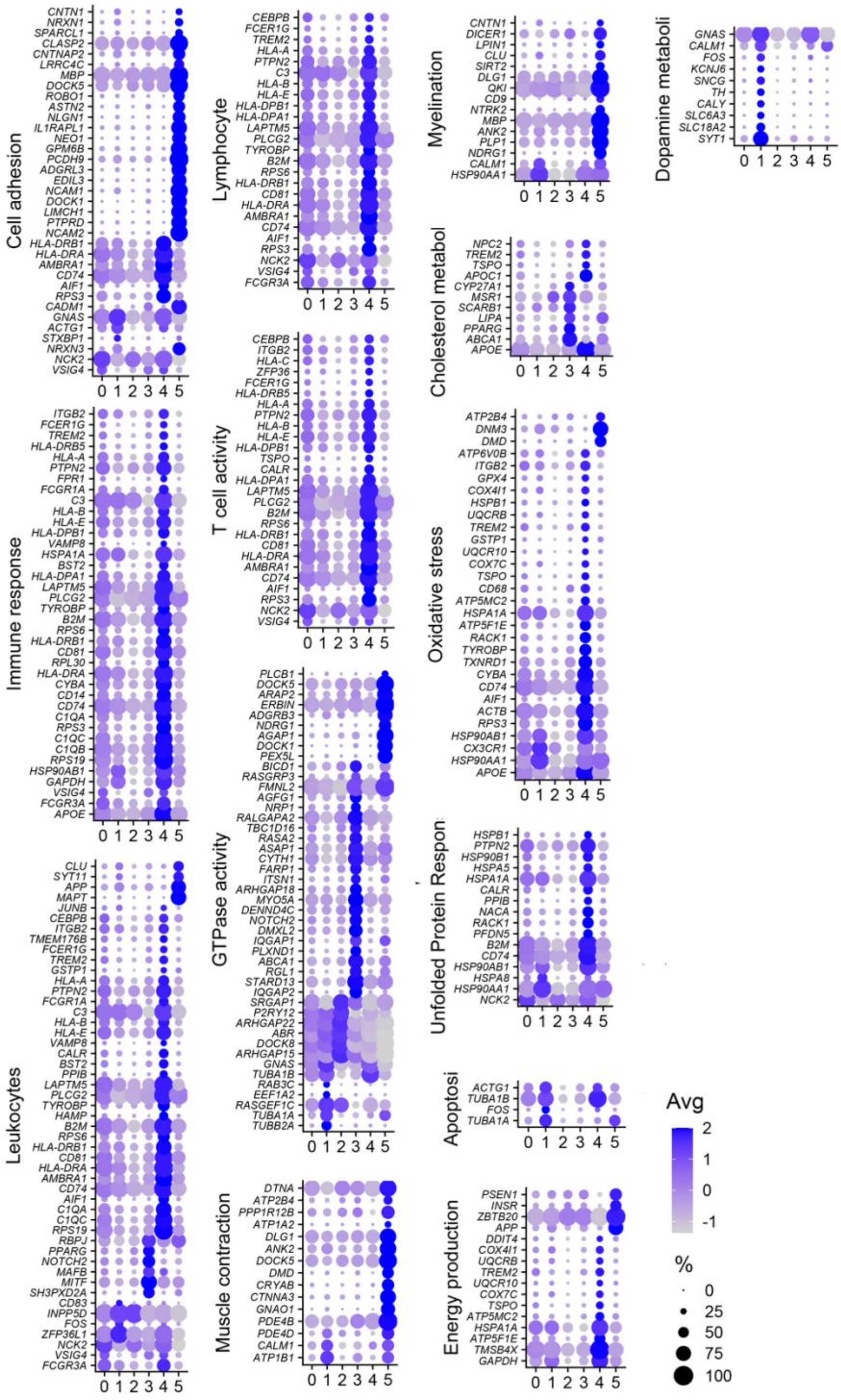


**Supplementary Figure 4:** Markers of microglial states grouped by enriched terms reported in **Table 6.**

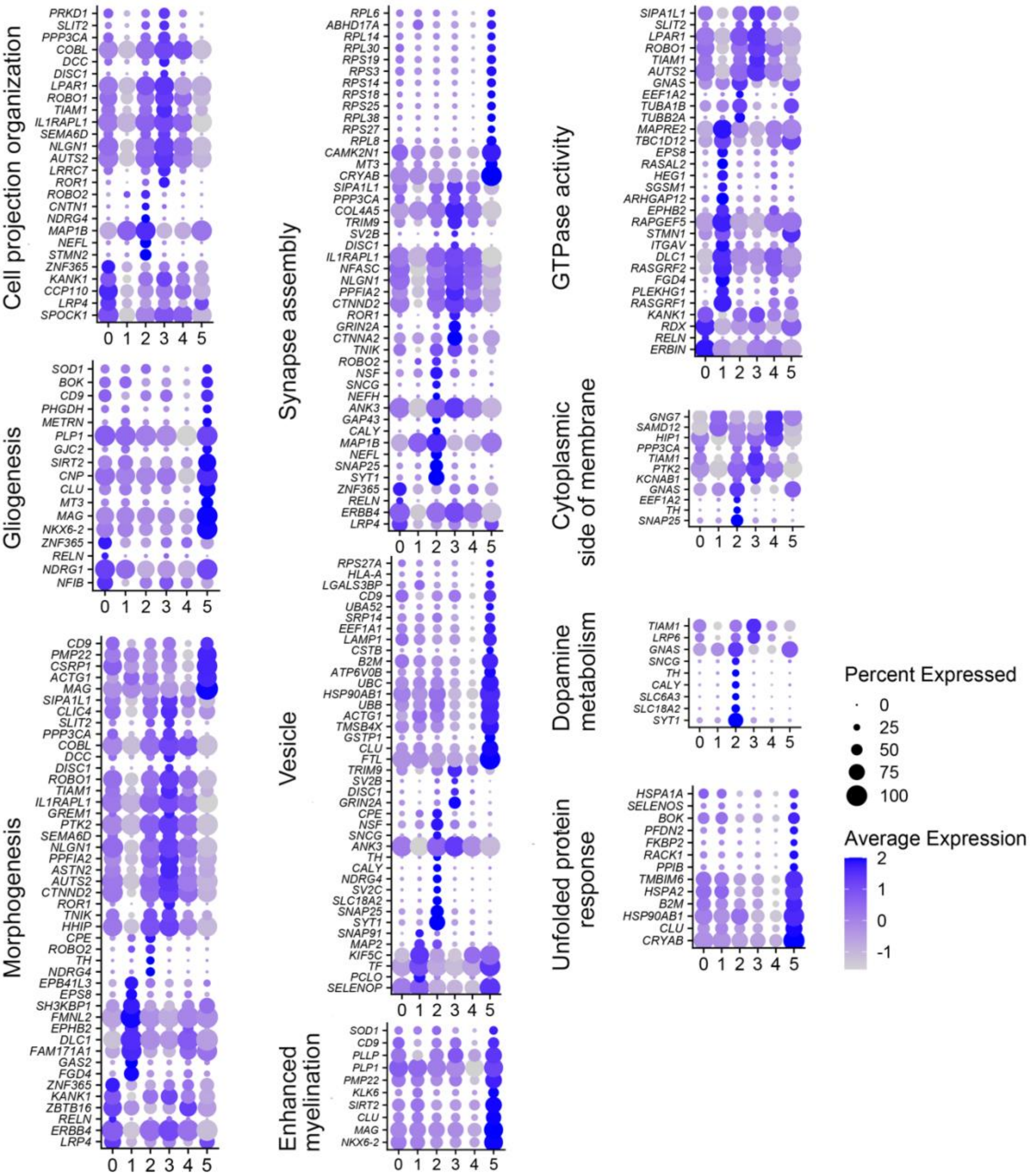


**Supplementary Figure 5:** Markers of oligodendrocytic states grouped by enriched terms reported in **Table 6.**

| N | Brain Bank ID | Clinical diagnosis | Estimated number of cells (1st run) | Estimated number of cells (2nd run) | Mean reads per cell (1st run) | Mean reads per cell (2nd run) | Median number of genes per cell (1st run) | Median number of genes per cell (2nd run) | Sequencing saturation (1st run) | Sequencing saturation (2nd run) | The total number of doublets detected by Scrublet algorithm |
|---|---|---|---|---|---|---|---|---|---|---|---|
| **1** | 2020 | Parkinson's | 5.790 | NA | 43.844 | NA | 2.341 | NA | 66,10% | NA | 88 |
| **2** | 2140 | Parkinson's | 11.147 | 13.032 | 19.470 | 44.400 | 2.141 | 2.704 | 42,10% | 63,20% | 414 |
| **3** | 2122 | Parkinson's | 3.927 | NA | 46.169 | NA | 1.543 | NA | 63,90% | NA | 49 |
| **4** | 2228 | Parkinson's | 5.821 | NA | 43.891 | NA | 2.666 | NA | 61,30% | NA | 51 |
| **5** | 2291 | Parkinson's | 5.398 | NA | 42.412 | NA | 1.668 | NA | 55,90% | NA | 61 |
| **6** | 2466 | Parkinson's | 4.679 | 4.781 | 43.159 | 51.359 | 1.812 | 1.879 | 57,80% | 62,10% | 58 |
| **7** | 2647 | Parkinson's | 3.817 | NA | 43.020 | NA | 1.531 | NA | 66,50% | NA | 26 |
| **8** | 2825 | Parkinson's | 4.672 | NA | 38.453 | NA | 1.516 | NA | 51,90% | NA | 43 |
| **9** | 2910 | Parkinson's | 6.533 | 7.890 | 26.199 | 42.793 | 1.653 | 1.527 | 48,10% | 64,30% | 107 |
| **10** | 3268 | Parkinson's | 2.135 | NA | 39.375 | NA | 706 | NA | 84,80% | NA | 24 |
| **11** | 3440 | Parkinson's | 4.156 | NA | 39.242 | NA | 944 | NA | 84,40% | NA | 129 |
| **12** | 3659 | Parkinson's | 4.773 | 6.080 | 23.801 | 37.682 | 1.509 | 1.462 | 38,90% | 54,20% | 56 |
| **13** | 3688 | Parkinson's | 3.783 | 4.373 | 33.745 | 44.313 | 1.781 | 1.763 | 43,30% | 52,30% | 40 |
| **14** | 3708 | Parkinson's | 1.497 | NA | 48.473 | NA | 2.181 | NA | 62,40% | NA | 20 |
| **15** | 3791 | Parkinson's | 2.876 | NA | 77.477 | NA | 584 | NA | 94,30% | NA | 45 |
| **16** | 2286 | Control | 4.509 | 5.038 | 33.044 | 46.336 | 1.259 | 1.289 | 47,80% | 57,60% | 33 |
| **17** | 2293 | Control | 3.871 | 5.583 | 15.642 | 36.638 | 940 | 1.427 | 21,90% | 40,30% | 41 |
| **18** | 2298 | Control | 5.527 | NA | 36.481 | NA | 1.883 | NA | 54,80% | NA | 76 |
| **19** | 2309 | Control | 5.396 | 7.403 | 7.607 | 35.357 | 775 | 2.146 | 18,00% | 42,20% | 142 |
| **20** | 2311 | Control | 6.446 | 9.976 | 11.846 | 31.295 | 1.217 | 1.995 | 22,50% | 45,60% | 128 |
| **21** | 2919 | Control | 5.473 | NA | 16.422 | NA | 801 | NA | 61,20% | NA | 95 |
| **22** | 2992 | Control | 10.098 | NA | 16.679 | NA | 975 | NA | 48,50% | NA | 2 |
| **23** | 3337 | Control | 4.275 | NA | 45.605 | NA | 2.371 | NA | 61,90% | NA | 50 |
| **24** | 3345 | Control | 7.381 | 8.613 | 29.089 | 42.926 | 2.096 | 1.994 | 51,70% | 64,00% | 220 |
| **25** | 3424 | Control | 4.061 | NA | 38.355 | NA | 1.665 | NA | 49,20% | NA | 30 |
| **26** | 3624 | Control | 5.810 | 8.129 | 19.912 | 36.322 | 1.525 | 1.620 | 32,90% | 52,10% | 64 |
| **27** | 3649 | Control | 7.594 | 11.032 | 17.879 | 32.349 | 1.471 | 1.425 | 38,40% | 61,60% | 180 |
| **28** | 3651 | Control | 7.444 | 8.279 | 32.152 | 46.032 | 1.946 | 1.894 | 54,20% | 64,20% | 186 |
| **29** | 3823 | Control | 3.465 | NA | 55.728 | NA | 783 | NA | 87,30% | NA | 38 |

**Supplementary Table 1:** Single nuclei RNA-seq quality assessment.
